# Stop guessing thinning distances: a data-driven framework for spatial thinning parametrization for ecological modelling

**DOI:** 10.64898/2026.10.23.759308

**Authors:** Héctor Ibáñez, Gorka Bidegain, Haitz Saéz de Ocáriz Borde, Jorge Mestre-Tomás, Heroen Verbruggen, Alexander Oliver Jueterbock, Mikel Aingeru Becerro

## Abstract

1. Spatially heterogeneous sampling and uneven publication effort in biodiversity repositories limit ecological niche characterization and habitat suitability modelling, affecting downstream inferences at regional and global scales. Spatial thinning offers a widely used strategy to mitigate these biases due to its simplicity and scalability. However, researchers typically select thinning parameters using heuristic or arbitrary criteria, often independent of species-specific characteristics, which introduces an additional, largely unquantified source of potential error.
2. Here, we propose a data-driven framework for spatial thinning parameterization that replaces heuristic decision-making with principled, reproducible criteria. We define optimality as the stability of thermal niche estimates derived from empirical data, rather than as an agreement with a known true distribution. The framework integrates two complementary approaches. The Last Stable Thinning Parameterization (LS-TP) identifies the largest thinning intensity that preserves the stability of species-specific niche descriptors. Biogeographic-size Dependent Thinning Parameterization (BD-TP) links thinning intensity to species’ geographic range size and scales spatial filtering across taxa with different distributional properties. Using seagrasses as a model system, we evaluated the framework across a global compilation of more than 100,000 occurrence records.
3. Our results showed that optimal thinning parameters varied systematically with species range size and that uniform thinning approaches led to either excessive information loss or reduction of spatial clustering. Furthermore, the framework identified species for which available data did not support stable niche characterization, as these species failed to meet stability criteria even at the smallest thinning parameters. We recommend explicitly acknowledging this limitation in future studies and prioritizing increased sampling effort for such undersampled taxa.
4. Our approaches provide a first step toward explicit, data-driven spatial thinning parameterization, with direct implications for ecological niche modelling, biodiversity assessment, and climate-driven distribution analyses.

## INTRODUCTION

The increasing availability of biological occurrence data in biodiversity repositories has expanded the application of ecological niche and habitat suitability models, enabling their use across broader spatial, taxonomic, and environmental contexts (Jones and Cheung, 2014; Araújo et al., 2019; Moore et al., 2023). Recent efforts have promoted more standardised and repeatable workflows for processing occurrence data, including data cleaning and bias correction procedures (Maldonado et al., 2015; Zizka et al., 2019; Moudrý et al., 2024; Xu et al., 2024). Despite this progress, several preprocessing steps still depend on subjective decisions, and researchers often lack explicit criteria to guide key data-handling choices. Developing transparent metrics and clear protocols therefore remains an important goal for ecological modelling (Yates et al., 2018).

The term “opportunistic data” is widely used to describe records collected through weakly structured or non-systematic sampling and compilation processes (Ball-Damerow et al., 2019; Cretois et al., 2021; Isaac et al., 2014; Reddy and Dávalos, 2003; Tulloch and Szabo, 2012). These data frequently exhibit non-uniform sampling effort (Cretois et al., 2021; Oliver et al., 2021; Reddy and Dávalos, 2003; Tulloch and Szabo, 2012), driven by spatial, temporal, taxonomic, and other observational biases (Isaac et al., 2014; Coron et al., 2018; Ball-Damerow et al., 2019; Erickson and Smith, 2021). Although large data volumes can partially offset these limitations and support macroecological analyses (Dickinson et al., 2010; Isaac et al., 2014; Bried and Siepielski, 2017; Gantchoff et al., 2022), biased occurrence data can still generate spurious ecological signals and misleading predictions when data quality and sampling bias are not explicitly accounted for (Ball-Damerow et al., 2019; Erickson and Smith, 2021). Indeed, Ball-Damerow et al. (2019) found that 31% of 501 analysed studies did not account for known data quality issues in opportunistic datasets, even though such issues can have downstream consequences for ecological inference (Yates et al., 2018).

Spatial thinning is widely used to reduce sampling bias in occurrence datasets (Suárez-Seoane et al., 2019; Hu et al., 2021; Sainz-Villegas et al., 2022; Moore et al., 2023; Huang et al., 2025), largely because it is simple, scalable, and implemented in standardised R workflows (Aiello-Lammens et al., 2015; Mestre-Tomás, 2026). Common approaches include grid-based thinning, which retains a single record per grid cell (Jones and Cheung, 2014; Chefaoui et al., 2018; Mestre-Tomás, 2026); distance-based thinning, which removes records within a predefined distance (Aiello-Lammens et al., 2015; Hu et al., 2021; Van Eupen et al., 2021; Xu et al., 2024; Mestre-Tomás, 2026); and density- or probability-based thinning, which preferentially reduces records in areas of high sampling density (Jueterbock et al., 2013; Verbruggen et al., 2013). In practice, thinning is commonly parameterised using fixed distances or grid resolutions applied uniformly across taxa, with parameter choices based on raster resolution or heuristic reasoning rather than species-specific data characteristics. Such choices can introduce an additional and unquantified source of uncertainty (Castellanos et al., 2019), because a single thinning intensity may remove critical information from narrowly distributed taxa while remaining insufficient to reduce spatial clustering in widespread species. Although recent work has evaluated thinning performance using virtual-species model validation (Lamboley and Fourcade, 2024), there is still limited guidance on how to parametrise thinning in real-world opportunistic datasets using internal properties of the data. Previous work has suggested that thinning performance depends on range size and environmental variability (Castellanos et al., 2019), but studies have not formally parametrised this relationship. As a result, thinning parameter selection remains largely subjective and may influence downstream ecological inference.

This limitation becomes particularly relevant in coastal systems, where many species exhibit narrow, linear, and fragmented ribbon-like distributions that differ from the broader spatial patterns typically analysed in terrestrial studies. Seagrasses provide an ideal model group to evaluate this problem. This polyphyletic group of high conservation importance (Duarte et al., 2008; Chefaoui et al., 2018) exhibits wide variation in geographic range size, endemicity, and sampling intensity, providing a strong gradient for testing thinning parameterization approaches. In this work, we focus on the thermal niches of seagrasses, which plays a central role in understanding climate vulnerability, range shifts, and conservation planning.

We address the lack of methodological procedures to parametrize thinning by introducing a data-driven framework for spatial thinning parameterization. Rather than relying on purely heuristic parameter choices, the framework uses explicit stability-based criteria to evaluate how thinning intensity affects thermal niche characterization. We conceptualize thinning parameter selection as a data-driven constrained selection problem, in which thinning intensity is increased up to the point at which stable niche characterization can no longer be supported. The framework includes two semi-independent approaches. Biogeographic-size-Dependent Thinning Parameterization (BD-TP), which relates thinning intensity to species range size using a classification approach, and Last Stable Thinning Parameterization (LS-TP), which identifies the maximum thinning level that preserves niche stability. Both methods rely on stability metrics derived from the convergence of thermal niche estimates across thinning gradients. The framework is evaluated using a global compilation of seagrass occurrence records (>100,000 observations) for all currently accepted species, assembled from biodiversity repositories, published datasets, and literature sources.

## METHODS

To implement this framework, we developed a workflow that links species-specific range characteristics with niche stability across a gradient of thinning intensities. The workflow combines three key components: (i) the estimation of species range size (with the species-specific range size descriptor, *D_s_*), (ii) the generation of thinned datasets across multiple thinning parameters (Θ), and (iii) the quantification of niche stability for each species-parameter combination (*s, θ*). The BD-TP approach explicitly links the optimal thinning parameter (*θ_opt,s_*) to the species’ range size (*D_s_*) using a classification framework, whereas the LS-TP approach selects *θ_opt,s_* as the largest thinning parameter for which niche estimates remain stable, independently of range size. Figure 1 summarizes the main steps of the workflow. All steps were carried out in R 4.5.1 (2025-06-13) (R Core Team, 2025), with use of basic (Wickham, 2016; Wickham et al., 2019) and more specialised packages cited through the text.

**Figure 1.**
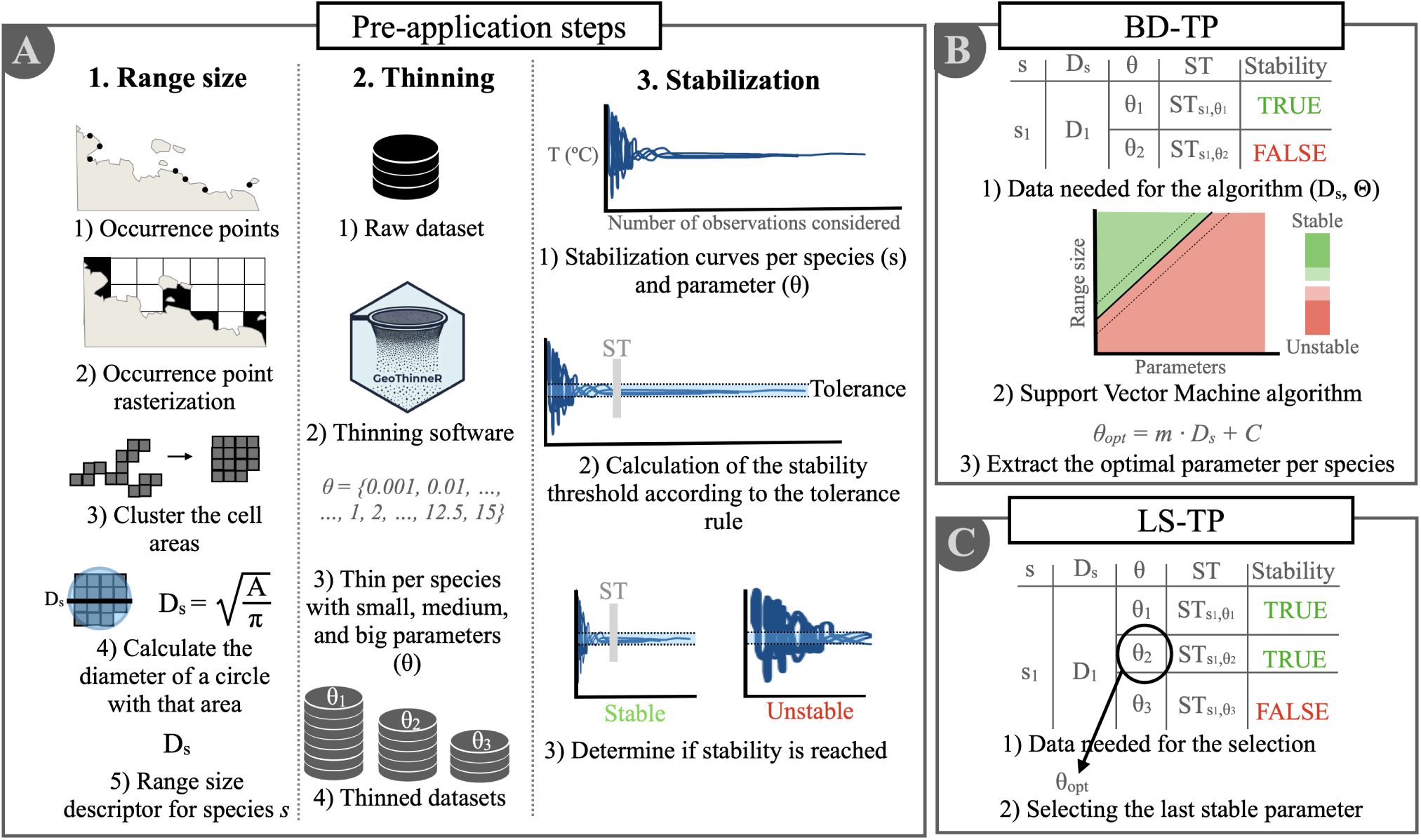
Summary of the main methodological steps in BD-TP and LS-TP. (A) Pre-processing pipeline, including (A1) generation of the range size descriptor, (A2) thinning the main dataset with different parameters, (A3) computation of the thermal stability trajectories and thresholds. (B) BD-TP: the thinning parametrization procedure which considers range size. Useful for datasets with several species with similar ecology (C) LS-TP: the thinning parametrization that maximises thinning regardless of range size. Useful when modelling with a small number of species.

### Data collection and preprocessing

The opportunistic seagrass dataset combined four main data sources: (1) global biodiversity repositories, GBIF and OBIS, queried on 3 September 2025 using the *rgbif* (v. 3.8.2) and *robis* (v. 2.11.3) packages (Chamberlain et al., 2017; Provoost and Bosch, 2022), respectively; (2) published global syntheses (UNEP-WCMC, 2020; Assis et al., 2024); (3) regional datasets targeting underrepresented areas and species (e.g., Sudo and Nakaoka (2020)); and (4) records extracted from the literature (see Supplementary Material 1: Extended Methods). Datasets obtained from GBIF and OBIS are listed in Supplementary Material 2. Records lacking an explicit reuse license or restricting derivative use were excluded. All seagrass species were considered, employing the query list shown in Supplementary Material 1, and considering a nomenclatural and taxonomical system with 54 seagrass species, as shown in Supplementary Material 3.

We retained records proximal to the 2000–2020 period to ensure consistency with the climatological raster data. To reduce spatial imprecision, we restricted records containing explicit uncertainty metadata to a maximum uncertainty of 2 km.

We cleaned occurrence records using established geospatial quality-control workflows (Coca de la Iglesia et al., 2023; Dorey et al., 2023), including CoordinateCleaner (v. 3.0.1) (Zizka et al., 2019). We removed duplicated records based on species identity, latitude, longitude, and decade. We excluded records showing common geospatial issues, including coordinates assigned to biodiversity institutions, country centroids, zero coordinates, equal latitude and longitude values, and invalid geographic positions. Taxonomic harmonization followed the taxonomic and nomenclatural notes summarised in Supplementary Material 3.

We extracted decadal mean Sea Surface Temperature (SST) values for each occurrence point from Bio-ORACLE v3 (Assis et al., 2024) (spatial resolution = 0.05 ° degrees; temporal resolution = decadal steps). To minimize data loss associated with raster coastal edges (such as points falling within estuaries, embayments, fjords, or harbours that do not have rasterised values), we applied a localised gap-filling procedure along raster boundaries (See Supplementary Material 1 for more details).

### Pre-processing pipeline for thinning parametrization

The two proposed spatial thinning approaches were applied on a seagrass global dataset compiled from sources of different coverage and scope. To later apply the two methods, the raw dataset was thinned with a parameter set and the niche stabilization of species and parameter pairs was characterised. For the BD-TP approach, the range size of each species was also calculated.

#### Pre-processing pipeline for thinning parametrization

We defined the range size of each species ‘*s*’ as the range size descriptor (*D_s_*), using the diameter of the clustered occupancy area. We rasterised occurrence records at a resolution of 0.5^◦^ and calculated occupancy areas as the number of occupied cells multiplied by the area of a single cell in km^2^. To ensure equal areas, spatial data was reprojected to EPSG:4326. To standardize comparisons among species with different distributional geometries, we calculated the diameter of a circle with an area equivalent to the occupancy area. *D_s_* was calculated only from the raw occurrence data and kept it constant across thinned datasets.

#### Generation of thinned datasets

Grid-based thinning (retention of a single random record per grid cell) was conducted with *GeoThinneR* v1 (Mestre-Tomás, 2026) to generate 11 thinned datasets using the following parameter set (in latitude degrees in the Equator):

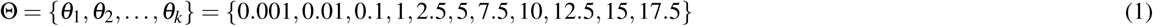

We selected Θ to include values that are clearly below and above the expected optimal thinning intensity for calibration purposes. We performed thinning by grouping records at the species level and iterated the thinning function across five trials, retaining the replicate that preserved the most records.

#### Niche stabilization analysis

We evaluated the stability of thermal niche characterization across species-parameter combinations (*s, θ* pairs). We conceptualised the niche optimum as one of its typical proxies, the SST median (Burrows et al., 2020) and quantified stabilization using cumulative moving medians across progressively larger random subsets of occurrence records. The stabilization procedure includes (i) the selection of settings, (ii) the calculation of replicate moving median trajectories, (iii) the computation of stability metrics, and (iv) summarization and visualization.

For each (*s, θ*) pair, cumulative moving medians of decadal SST average were computed with the *slider* package (v. 0.3.2) (Vaughan, 2024) for up to 500 randomly subsampled occurrence records from the thinned dataset. The process was iterated 10 times, resulting in 10 replicate trajectories. We aggregated replicate trajectories to derive the summary median estimate and the associated 95% confidence intervals.

We defined the stabilization threshold of each pair (*ST_s,θ_*) as the first occurrence index (*n_x_*) at which three conditions were simultaneously met: (i) the absolute difference between the cumulative median at observation *n_x_*(*m̃_n_x__*) and the terminal reference value (*m*_ref_) is less than or equal to 0.3 ^◦^C (- 4see Eqs. 2–4), (ii) the confidence interval width is less than or equal to 0.5 ^◦^C, and (iii) these conditions hold for at least 20 consecutive observations. The stability condition at observation *n_x_* is defined as:

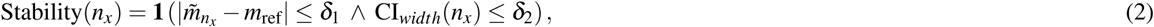

where *m̃_n_x* is the cumulative median across replicate trajectories at observation index *n_x_*, *δ*_1_ = 0.3^◦^C, *δ*_2_ = 0.5^◦^C, **1**(·) is the indicator function, and *m*_ref_ is defined as the mean of the last *K* cumulative median estimates:

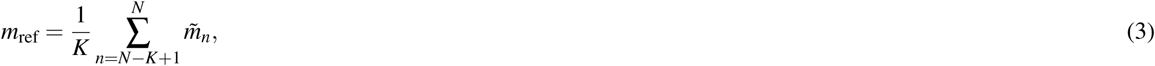

where *m̃_n_* denotes the cumulative median at observation index *n*, and *N* is the maximum number of observations considered for the corresponding (*s, θ*) pair. We used *K* = 20 terminal cumulative median estimates to calculate *m*_ref_.

The stabilization threshold is then defined as:

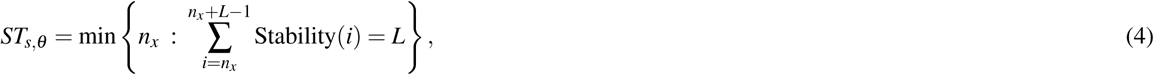

where *L* = 20 denotes the minimum number of consecutive observations required for stabilization.

The resulting threshold (*ST_s,θ_*) represents the minimum number of observations required to obtain a stable estimate of the thermal optimum. We compiled thresholds for all (*s, θ*) pairs and used them as inputs for the two data-driven thinning parameterization methods.

### Biogeographic-size dependent thinning parametrization (BD-TP)

In the biogeographic-size dependent thinning parameterization (BD-TP) method, for each species *s*, we conceptualised the relationship between range size (*D_s_*) and thermal niche stability across thinning parameters as:

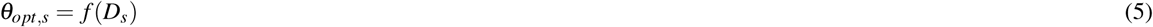

with *θ_opt,s_* represents the optimal thinning parameter predicted from species range size for a species *s*. The method is built under the assumption that bigger ranges will support a higher maximised thinning parameter, without turning the ecological signal unstable.

We classified each (*s, θ*) pair according to whether the stabilization threshold had been reached (*ST_s,θ_* ≤ *N_s,θ_*) or not (*ST_s,θ_ > N_s,θ_*), with *N_s,θ_* being the total number of observations for the pair. We modeled the function using a Support Vector Machine (SVM) algorithm (Vapnik, 1995). The SVM classified stability using *D_s_* and *θ* as predictors. We implemented the analysis using the svm() function from the *e1071* package (v. 1.7-16) (Meyer et al., 2024) with a linear kernel. We extracted the hyperplane coefficients analitically and fitted them to a linear model, resulting in the formula:

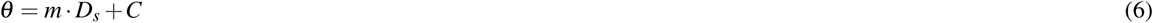

We capped the formula at a minimum thinning parameter of 0.001^◦^ to avoid theoretical negative values and ensure that extremely proximal points remained thinned.

### Last stable thinning parametrization (LS-TP)

The second method, last stable thinning parameterization (LS-TP), does not consider range size and defines the optimal thinning parameter as the largest value of *θ* that satisfies the stability condition:

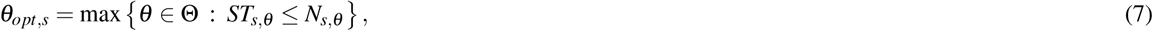

where Θ is the set of evaluated thinning parameters for species *s*, and *N_s,θ_* is the number of available observations after thinning. Thus, the LS-TP method attempts to identify the last tested thinning parameter for which there is stability. The formula was also capped at a minimum *θ* = 0.001.

### Testing the methods

Habitat suitability models (HSMs) and realised thermal niche (RTN) curves were computed for *Phyllospadix torreyi*, a seagrass endemic to the northeastern Pacific and the species with sufficient occurrence data for this analysis (178 original presence records). To assess the effect of spatial sampling bias and thinning, HSMs were fitted for four separate datasets: datasets spatially thinned using the parameters estimated by (1) BD-TP and (2) LS-TP, (3) the environmental raster resolution (0.05^◦^) and (4) no thinning. To evaluate the effects of strong spatial sampling bias, an additional ‘biased’ dataset was generated by adding 300 artificial occurrence points assigned randomly in close proximity to the three major cities along the distribution of *P. torreyii*, and thinned with the same four treatments (See Supplementary Material 1 for more details). HSMs were fitted using GLOSSA, an R Shiny application implementing Bayesian Additive Regression Trees (BART) for marine species distribution modelling (Mestre-Tomás et al., 2026). Five of the most common variables for seagrass HSM were used (Bertelli et al., 2022), including bathymetry, temperature, photosynthetically active radiation (PAR), slope, and salinity. Environmental rasters were downloaded from Bio-Oracle v3 (Assis et al., 2024). Pseudo-absences were generated randomly within a 0.2^◦^ buffer-restricted area from presence occurrence points using a 1:1 pseudo-absence-to-presence ratio. Model performance was assessed through 10-fold spatial block cross-validation, with block size determined from the spatial autocorrelation of model residuals, using all evaluation metrics available in GLOSSA.

RTN curves were estimated from the temperatures associated with *P. torreyi* occurrence records for both the original and artificially biased datasets and the four candidate parameters. Thermal niches were represented by kernel density estimates of the temperature values at occurrence sites, calculated using `ggplot2::geom_density()` with its default Gaussian kernel and optimal smoothing bandwidth selected with biased cross-validation (Scott and Terrell, 1987). To evaluate the sensitivity of the estimated thermal niche to the occurrence sample, a repeated resampling was performed 50 times, randomly assigning 60% of occurrence records to the training subset in each replicate.

## RESULTS

The two methods were tested on a global seagrass dataset with 109,150 occurrence records for 54 taxa. The number of raw occurrence records per species showed significant variability, with important deviations from the median value of 148.5 obs. (interquartile range [IQR] = 32.5 obs - 1920.5 obs.). *Nanozostera muelleri* was the most frequently recorded taxon with 20,813 occurrence records, whereas *Zostera pacifica* and *Halodule bermudensis* contained only three records. Six taxa were identified as upper-bound outliers in occurrence counts with the Tukey 1.5×IQR method: *N. muelleri*, *Halophila ovalis* group, *Zostera marina*, *Halodule uninervis*, *Thalassia testudinum*, and *Thalassia hemprichii*.

Species range sizes varied significantly as well, with important deviations from the median value of 314.96 km (IQR = 214.70 - 605.65 km). *Halodule bermudensis* showed the smallest range size (52.93 km), whereas *Zostera marina* displayed the largest range (1149.72 km).

### Stabilization curves and thresholds

We found that 31.48% of the species did not reach stabilization under any thinning parameter across the evaluated thinning gradient (Θ = [0.001 − 17.5]). The cumulative moving medians of decadal SST values followed markedly different stabilization trajectories across species-parameter combinations (*s, θ* pairs). The results for two species, *Thalassia hemprichii* with a broad Indo-Pacific distribution and the Australian endemic *Posidonia angustifolia*, are shown in Fig. 2. Outpus for all other species and parameters are available in the Supplementary materials.

**Figure 2.**
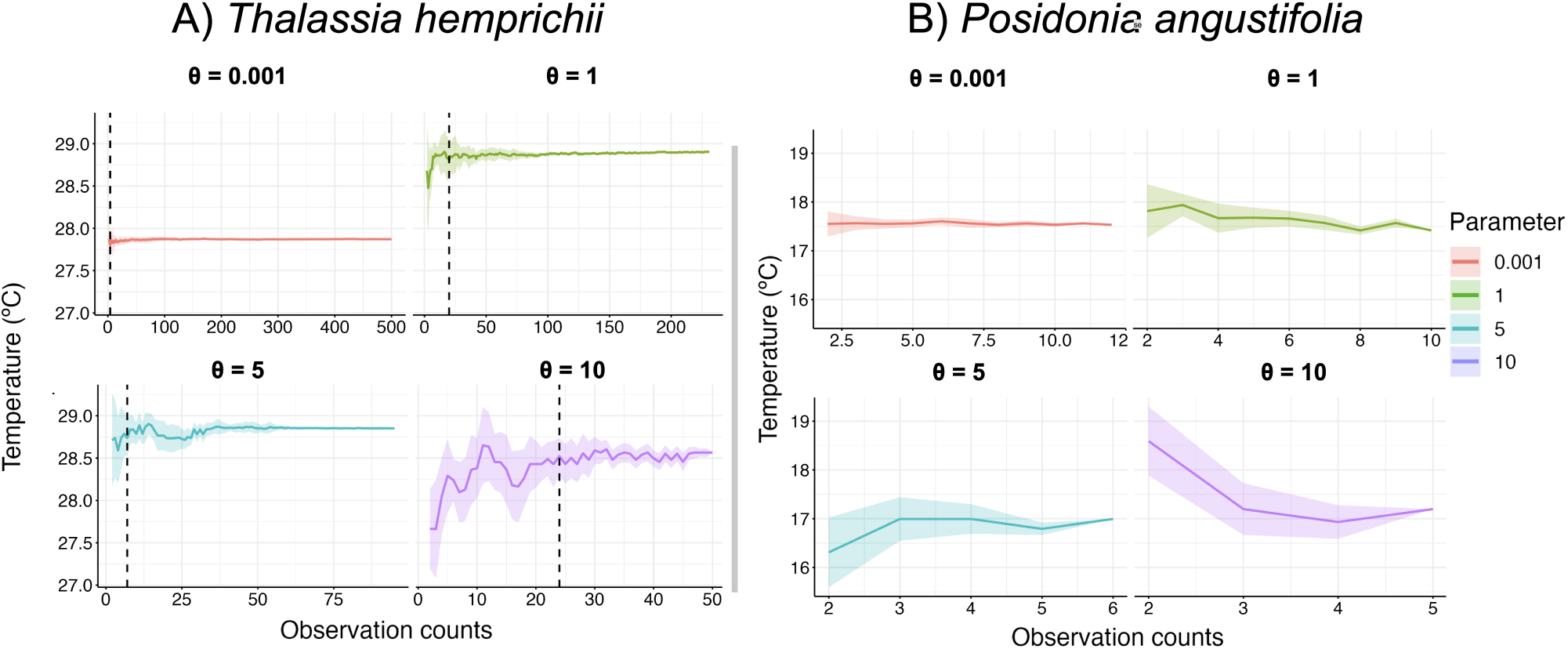
Global gridded map showing seagrass count frequencies. Cells (cell size = 3 °) that are contoured indicate at least a single seagrass presence. Darker colors indicate higher counts. Frequencies where log(10) transformed to normalize the data, making grid cells comparable. Biogeographic realms following Spalding et al., (2007) are plot in different colors under the following acronyms: TAu, Temperate Australasia; CIP, Central Indo-Pacific; SOc, Southern Ocean; TNA, Temperate Northern Atlantic; TAt, Tropical Atlantic; WIP, Western Indo-Pacific; TSAf, Temperate Southern Africa; TNP, Temperate Northern Pacific; TSAm, Temperate South America; EIP, Eastern Indo-Pacific; TEP, Tropical Eastern Pacific; Arc, Arctic

### Application of the new methods

#### BD-TP

The optimal thinning parameter increased progressively with species range size:

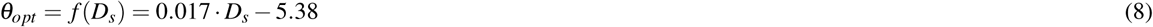

For visual clarity, the relationship was plotted inversely in Fig. 3. Higher thinning parameters were associated with larger range sizes, with no observed loss of niche characterization across the tested gradient. The model predicted that species with a *D_s_* ≤ 308.31 km do not have enough reported coverage to have a stable niche calculation at any thinning parameter. The taxa with the second to fourth largest range sizes (*H. ovalis* group, *H. uninervis* and *T. hemprichii*) are stable at thinning parameters as high as 10°. However, the seagrass species with the largest range, *Z. marina*, differs from the pattern and loses stability at a much lower parameter than predicted (LS-TP = 2.5°, BD-TP = 14.7°). In the lower bound, the inference of the parameter was capped for 26 species, with range sizes 52.90 - 308.30 km, at a minimal thinning parameter of 0.001°. The BD-TP workflow retained 2,306 observations from the original 109,150 records (total retention rate = 0.0211; median species retention rate = 0.49, IQR = 0.02 - 0.95).

**Figure 3.**
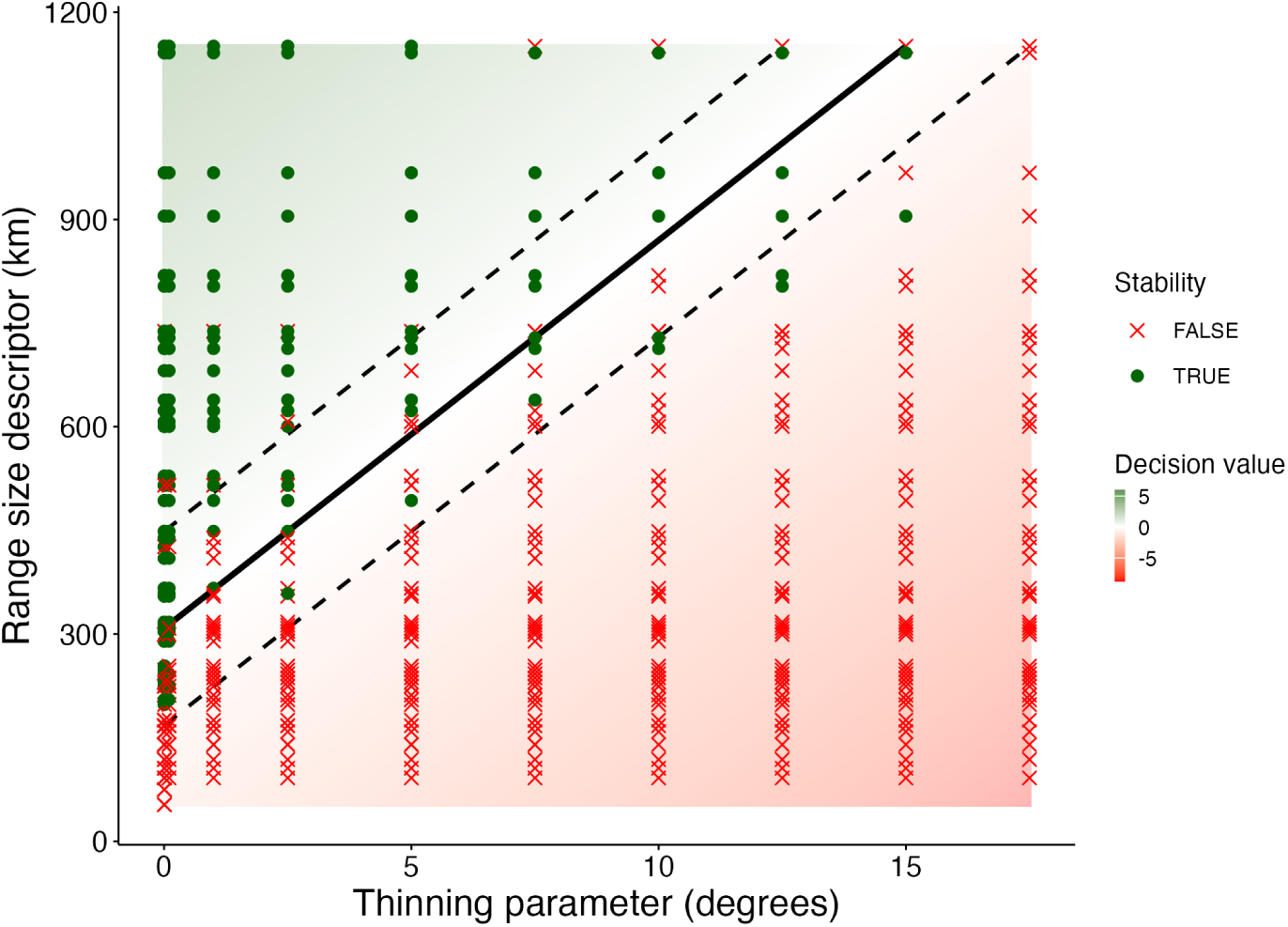
Support Vector Machine (SVM) plot for predicting the optimal spatial thinning parameter for 54 seagrass species. The SVM hyperplane (black solid line) classifies stability based on the predictors *D_s_* (range size descriptor) and *θ* (thinning parameter). Dashed lines indicate margin boundaries. ‘Stability’ designates whether a species-parameter pair has enough information for a stable niche calculation. Thermally stable thinning parameter and species combinations are represented with green dots, whereas thermally unstable combinations are shown in red crosses.

#### LS-TP

The LS-TP method predicted a maximum thinning parameter (*θ_opt_*) of 15^◦^, for the two species with the third largest range size *H. uninervis*. At the lower bound, 15 species with range sizes between 52.90 and 299.12 km were identified to be unstable under all thinning parameters. The LS-TP workflow retained 2,839 observations from the original 109,150 occurrence records (total retention rate = 0.0260; median species retention rate = 0.40; IQR = 0.032 – 0.906).

Both *Z. marina* and *R. maritima*, respectively the species with the largest and the eight largest range sizes, displayed the largest discrepancy between the parameters predicted by the two methods (|*θ_BD_*_−*TP*_ − *θ_LS_*_−*TP*_| *>* 7.3^◦^) and were identified as outliers in comparison to the other deviations with the Tukey 1.5×IQR method, along with *Enhalus acoroides* (Diff = 4.2) and *Halophila spinulosa* (Diff = 4.2). All remaining species showed deviations below 5^◦^, with a median difference of 0.099 ([IQR] = 0 - 1.66) between methods, including species capped to 0.001^◦^.

#### Downstream effects

The four thinning parameters (LS-calculated, BD-calculated, 0.05, and none) resulted in different HSM projections (e.g., Potential suitable area (·10^5^ km^2^) - Original: raw = 2.2, 0.05^◦^ = 3.3, LS = 5.6, BD = 5.7; Biased : raw = 8.9 (·10^4^ km^2^), 0.05^◦^ = 4.5, LS = 4.1, BD = 5.6). The HSI median is shown as a point estimates for each ‘case’ (Fig. **??**). Uncertainty is shown with the mapped standard deviation (available in the Supplementary materials).

**Figure 4.**
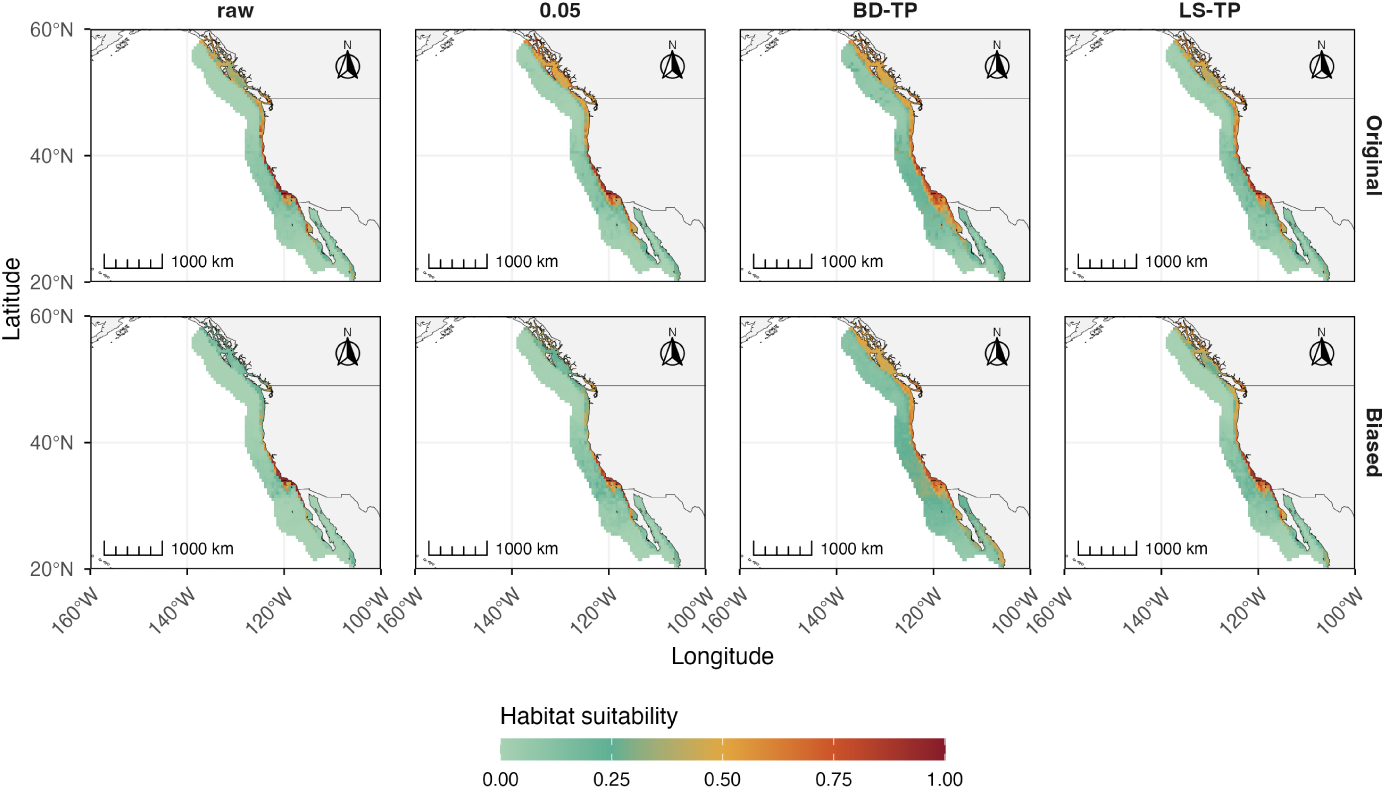
Habitat suitability models for *Phyllospadix torreyi* based on the original (first row) and artificially biased (second row) occurrence datasets. Habitat suitability index (HSI), calculated using BART, is shown as a gradient fill representing the median predicted HSI. For both datasets, models were fitted using four occurrence-data treatments: unthinned data (raw), data thinned at 0.05^◦^, and data thinned using the parameters estimated by BD-TP and LS-TP.

Cross-validated CBIs reported the highest variability among metrics (See See Supplementary Material 8), whilst the CBIs computed with all the datapoints reported higher values (original dataset: raw = 0.95, 0.05° = 0.95, LS-TP = 0.92, BD-TP = 0.93 |biased dataset: raw = 0.98, 0.05° = 0.94, LS-TP = 0.89, BD-TP = 0.6). The thermal curves showed different sensitivity levels according to the data origin and thinning parameter (Fig. **??**). For the datasets derived from the original, a clear bimodality was captured for the raw data, which was progressively less apparent for 0.05^◦^, LS and BD. For the datasets with artificial bias, the raw dataset was overfit and wiggly, whereas the other three curves showed a similar pattern to that of the original datasets.

**Figure 5.**
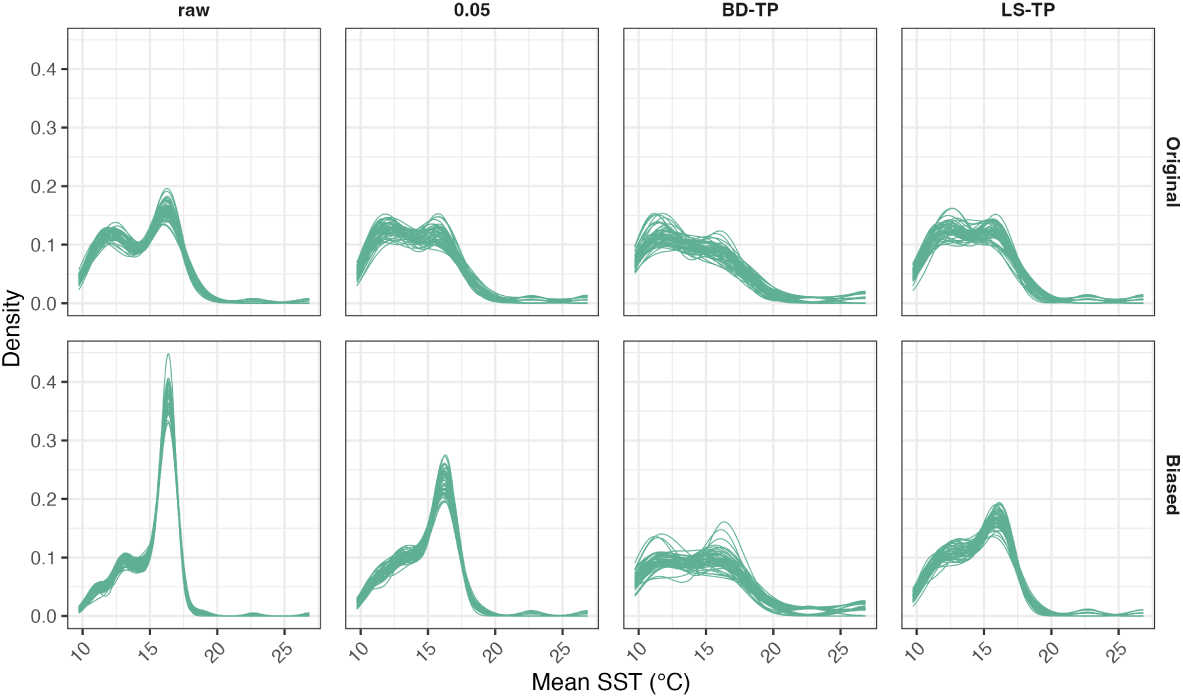
Realised thermal niche curves for *Phyllospadix torreyi* based on the original (first row) and artificially biased (second row) occurrence datasets. Each replicate curves samples 60% of the available data for each thinning parametrization procedure. In both “original” and “biased”, the raw data (raw) and the data thinned with 0.05 and the predicted parameters by BD and LS were inputted into the model.

## DISCUSSION

Sampling bias in opportunistic datasets remains a well-documented issue that can distort ecological inference across spatial scales (Ball-Damerow et al., 2019; Isaac et al., 2014; Oliver et al., 2021; Coron et al., 2018; Erickson and Smith, 2021). By introducing a data-driven framework for thinning parameterization, we replace subjective preprocessing decisions with objective, reproducible criteria and provide a consistent basis for bias correction in ecological analyses.

The framework integrates two complementary approaches that capture different aspects of thinning behaviour. BD-TP links thinning intensity to species range size and provides a scalable rule to adjust spatial filtering across taxa with contrasting distributional properties. LS-TP identifies the maximum thinning level that preserves stable niche descriptors without requiring information on range size. These approaches define a consistent strategy to balance bias reduction and information retention by constraining the optimal thinning parameter to the maximum predicted (BD-TP) or observed (LS-TP) value that precedes niche destabilization. In this work, we defined thermal niche stability as the persistence and consistency of thermal characterization across increasing thinning intensities, up to the point at which model outcomes become dominated by stochastic retention of occurrences rather than by underlying ecological structure.

The two approaches differed in their record retention patterns. LS-TP generally retained a larger proportion of occurrence records, whereas BD-TP applied stronger filtering to species with broad geographic distributions. The largest discrepancies between methods occurred in two widespread species, while most remaining species showed relatively small deviations between approaches. These differences reflect the distinct conceptual foundations of the two methods: LS-TP constrains thinning based solely on species-specific stabilization limits, whereas BD-TP additionally scales thinning intensity based on geographic range size. BD-TP poses itself as a scalable approach when several taxa of similar biogeographic drivers are considered, whilst LS-TP can be used independently for single-species modelling approaches.

### Range size (*D_s_*) as a model input

The range size descriptor (*D_s_*) approximates key aspects of species distribution while accounting for sampling structure. Conceptually, it approximates the extent of occurrence (EOO; i.e. the area encompassed within the outermost geographic limits of a species’ distribution) (Gaston and Fuller, 2009). However, its computational implementation is closer to the area of occupancy (Gaston and Fuller, 2009), as it reflects the spatial coverage of observed occurrences rather than the polygon enclosed by extreme records. This approach was preferred over classical EOO estimators in order to reduce range estimate inflation for coastal distributions. We therefore recommend *D_s_* as a sampling-aware descriptor of range size for coastal taxa.

### Redefining optimality for thinning parametrization

Our results showed that optimal thinning parameters varied systematically across species and that uniform thinning approaches introduced uncontrolled variability in ecological inference. A single parameter can remove critical information for range-restricted taxa while failing to reduce sampling bias in widespread species. These findings aligned with previous work showing that thinning effects differ across species and that no universally optimal parameter exists (Lamboley and Fourcade, 2024). Rather than identifying a single optimal value, our framework defined species-specific criteria that constrain thinning intensity based on data structure and niche stability.

Beyond parameter selection, the framework identified when occurrence data failed to support reliable ecological inference. In our dataset, a 31.48% of the total species failed to reach stabilization across all thinning levels, indicating insufficient sampling coverage (*D_s_* ≤ 308.31 km implying expected instability). Opportunistic datasets often contain uneven and incomplete sampling that can bias ecological analyses if not explicitly addressed (Isaac et al., 2014; Bried and Siepielski, 2017). By identifying these cases, the framework also introduces a diagnostic step that evaluates data quantity adequacy linked to our definition of niche stability. We recommend caution for species deemed as “data deficient” by the proposed methods.

### Usage recommendations

Both methods have been implemented in GeoThinneR v2. We accept both BD-TP and LS-TP as effective tools for managing spatial bias prior to HSM or niche modelling. For single-species, regional and range-restricted studies, we advise using LS-TP first. BD-TP poses a more complete option for multi-species global studies or scalable workflows; if the two disagree by large numbers, further investigation is advised.

### Position within bias-correction paradigms and previously employed parametrization criteria

Most studies select thinning parameters heuristically and rarely compare alternatives explicitly (Nolan et al., 2022; Koldasbayeva et al., 2022; Zárrate Charry et al., 2022). Across the literature, six main parametrization strategies can be identified: (i) thinning to match the environmental raster resolution (Laeseke et al., 2021; Farooq, 2025; Wang et al., 2025; Hu et al., 2021) and (ii) applying a fixed geographic filter, often 10 km (0̃.1^◦^), across all species (Sainz-Villegas et al., 2022; Adelino et al., 2017; Boria et al., 2014). However, these approaches are rarely justified ecologically or in terms of sampling bias reduction (Castellanos et al., 2019). Moreover, the resolution at which sampling bias changes is often bigger than that in both approaches (Zizka et al., 2020). Other strategies include (iii) two-phase filtering combining thinning using the raster-cell and another fixed parameter (Verbruggen et al., 2013; Hu et al., 2021), (iv) accounting for predictor autocorrelation (Suárez-Seoane et al., 2019), (v) species-specific thinning without a mathematical parametrization (Nolan et al., 2022; Koldasbayeva et al., 2022), and (vi) our proposed species-dependent data-driven approaches. Smaller thinning distances are more common in regional studies, whereas coarser filters dominate global analyses (Castle et al., 2022; Manca et al., 2024). Importantly, most comparative studies have focused on continental or smaller extents (Montalvo-Mancheno et al., 2025; Lamboley and Fourcade, 2024; Koldasbayeva et al., 2022; Nolan et al., 2022) and global studies are lacking. In contrast to these existing strategies, our approaches provide a data-driven, taxon-specific framework in which thinning parameters are derived from species-level ecological characteristics rather than than relying on a single, predefined grid size applied uniformly across species.

Attempts to identify an optimal thinning parameter remain limited and theoretical. Castellanos et al. (2019) showed that environmental thinning can reduce model inflation in clustered datasets, although effects varied for range-restricted species, suggesting dependence on environmental heterogeneity and distributional extent. Using virtual species, Lamboley and Fourcade (2024) concluded that stronger thinning consistently reduces clustering at the cost of information loss, and therefore no universal optimum exists. The differing results from Lamboley and Fourcade (2024) and ours are likely caused by differences in how optimality is defined. Ideally, sampling bias should be minimised during field data collection, but this is hardly achievable in practice. Rather than comparing observations to a perfectly sampled distribution, our approach evaluates thinning through a stability metric derived from the structure of the empirical data itself, allowing practical optimization under real-world sampling constraints.

### Pitfalls and challenges

The proposed framework has several limitations related to stabilization, range estimation, and model parametrization. (1) Because stabilization relies on cumulative statistics without an external benchmark, it does not necessarily imply ecological validity. Undersampled endemic species may therefore appear unstable at low thinning parameters despite potentially stabilizing with greater field sampling effort. (2) Comparisons among stabilization curves are also constrained by the fixed cap of 500 sampled observations, which causes unequal x-axis limits across species and thinning intensities when the available record number is lower. (3) Similarly, the range size metric (*D_s_*) is strongly dependent on spatial resolution and sampling coverage, and should therefore be interpreted as a sampling-dependent approximation rather than a true measure of range size. Species may appear range-restricted because of geographic sampling gaps rather than biological rarity. However, this poses no problem to the methods, as this instability is informative in itself, because undersampled and sparsely sampled taxa are expected to become readily unstable under weak thinning. Alternative metrics such as convex hull area were considered and rejected because they are highly sensitive to outliers and coastal geometries. (4) Although the SVM framework reduces arbitrary parametrization, subjective decisions remain regarding the *D_s_*spatial resolution and the tolerance thresholds defining stability. (5) Although most large-ranged species behaved similarly the subcosmopolitan *Z. marina* (present all over the northern hemisphere) and *R. maritima* (present in all continents except Antarctica) represented notable exceptions, likely because strong thinning compresses environmentally distinct regions into single points and because broad-ranged, unevenly sampled taxa require larger thinning values before bias reduction becomes detectable, potentially producing poorly fitted predictions. Consequently, the framework may be more reliable for species with intermediate to big range sizes without an almost cosmopolitan or subcosmopolitan distribution. We recommend to use both methods comparatively and critically. (6) More generally, the framework addresses spatial bias only implicitly and cannot overcome the fundamental problem of missing data in environmentally important regions. (7) The SVM hyperplane should be regarded as descriptive rather than strictly predictive, since both predictors and inferred responses derive from the same occurrence data, causing coupled propagation of uncertainty in stabilization and range estimation. (8) Finally, just as a lower bound was used to constrain negative or null parameter predictions, an upper bound could prevent the formula from extrapolating to ecologically unrealistic thinning resolutions for species with exceptionally large ranges.

## CONCLUSIONS

To our knowledge, this study provides the first diagnostic framework for evaluating data adequacy and thinning sensitivity while formalizing the relationship between thinning intensity, niche stabilization, and range size. Future work could extend the framework beyond thermal niche descriptors and assess stability across multiple environmental dimensions. Incorporating niche breadth and multivariate descriptors may further strengthen ecological interpretation and help determine whether stability along a single environmental axis generalizes to broader niche characterization. Furthermore, applying this framework to other taxonomic groups and ecosystems would help evaluate its generality and identify context-specific refinements, including whether the *D_s_* metric should be adapted for non-coastal distribution geometries. More broadly, future research should examine how thinning interacts with predictor selection and other bias-correction methods, and how these interactions affect estimates of niche stability.

Overall, this study provides a data-driven framework that can serve as a step toward more reproducible thinning parameterization in ecological niche modelling workflows. By replacing purely heuristic parameter selection with explicit stability-based criteria, the framework may improve transparency and reproducibility, while also helping identify species for which available biodiversity records may be insufficient to support stable ecological inference. In doing so, it offers a scalable approach to a previously subjective and under-formalised preprocessing decision.

## ACKNOWLEDGEMENTS

This work was supported by the MARVELS project (PID2025-170011OB-I00), funded by the Spanish Ministry of Science, Innovation and Universities (MICIU) and the Spanish State Research Agency (AEI), and co-funded by the European Union through the European Regional Development Fund (ERDF). Additional support was provided by the Basque Government through the IMasT project (IT2055-26), by the University of the Basque Country (EHU) through the predoctoral fellowship PIF24/222, and the OSCARS-funded project FAME (European Commission’s Horizon Europe Research and Innovation programme) under grant agreement No. 101129751.

## AUTHOR CONTRIBUTIONS

All authors contributed to the investigation, manuscript writing, and manuscript review and editing. Héctor Ibáñez contributed to conceptualization and led data curation, methodology, software development, and validation. Gorka Bidegain contributed to conceptualization, methodology, software development, validation, and supervision. Haitz Saéz de Ocáriz Borde and Jorge Mestre-Tomás contributed to methodology, software development, and validation. Mikel Aingeru Becerro contributed to conceptualization and supervision. Heroen Verbruggen and Alexander Oliver Jueterbock contributed to the investigation, manuscript writing, and manuscript review and editing. All authors reviewed and approved the final version of the manuscript.

## DATA AVAILABILITY

The public datasets used and/or analysed during the current study are already of public domain. We do not own the data that consolidate the final dataset used as an input for our thinning parametrization workflow. The authors have merely acted as aggregators. The code used to download and curate these datasets and perform data-driven parametised thinning and the fine-literature dataset generated *sui-generis* are publicly available on Zenodo (available upon publication) for further reproducibility.

## STATEMENT OF INCLUSION

Our study brings together authors from a different European countries where the tools (HSM, thermal niche) for which the proposed methods have been designed have been widely implemented. All authors were engaged on the research outline, process and publication. Many local and regional studies are cited in the methods and supplementary materials sections.

## SUPPORTING INFORMATION

Additional supporting information is provided for the worked examples:

**Supplementary file S1:**extended methods and methodological results **Supplementary file S2:** Data compiled uniquely for this study and data sources, including (S2.1) the literature supplement data and its sources (csv file), and (S2.2.) the complete list of GBIF and OBIS datasets included in the analyses, including accession details and institutional sources. **Supplementary file S3:** Taxonomic, nomenclature and species selection statement **Supplementary file S4:** Dataset description. **Supplementary file S5:** Tabularised main study outputs. (S5.1) stabilization thresholds across thinning gradients and species. (S5.2) Species-specific thinning parameter outputs and range size descriptor outputs. (S5.3) Number of counts pre- and post-thinning. (S5.4) Bootstrap uncertainty and leave-one-species-out cross-validation for BD-TP. **Supplementary file S6:** Summarised cumulative moving median stabilization trajectories across species and thinning parameters. **Supplementary file S7:** Sensitivity analyses using (S7.1.) alternative range size resolutions, (S7.2) alternative tolerance settings for the stabilization, (S7.3) alternative parameter sets. **Supplementary file S8:**Evaluation metrics for the modeling outputs (thermal niche and HSM) under different thinning approaches and standard deviation maps for the HSM.

## Sup. Mat. - Stop Guessing Thinning Distances: a data-driven framework for spatial thinning parametrization for ecological modelling

- **Supplementary file S1:** extended methods and methodological results
- **Supplementary file S2:** Data compiled uniquely for this study and data sources, including (S2.1) the literature supplement data and its sources (csv file), and (S2.2.) the complete list of GBIF and OBIS datasets included in the analyses, including accession details and institutional sources.
- **Supplementary file S3:** Taxonomic, nomenclature and species selection statement
- **Supplementary file S4:** Dataset description.
- **Supplementary file S5:** Tabularised main study outputs. (S5.1) stabilization thresholds across thinning gradients and species. (S5.2) Species-specific thinning parameter outputs and range size descriptor outputs. (S5.3) Number of counts pre- and post-thinning. (S5.4) Bootstrap uncertainty and leave-one-species-out cross-validation for BD-TP.
- **Supplementary file S6:** Summarised cumulative moving median stabilization trajectories across species and thinning parameters.
- **Supplementary file S7:** Sensitivity analyses using (S7.1.) alternative range size resolutions, (S7.2) alternative tolerance settings for the stabilization, (S7.3) alternative parameter sets.
- **Supplementary file S8:** Evaluation metrics for the modeling outputs (thermal niche and HSM) under different thinning approaches and standard deviation maps for the HSM.

## 1 SUPPLEMENTARY FILE S1: EXTENDED METHODS AND METHODOLOGICAL RESULTS

### 1.1 Data downloads and processing

The seagrass dataset was compiled from four primary sources: (1) large global databases (GBIF, OBIS; search on 03/09/2025 with *rgbif* (Chamberlain et al., 2017) and *robis* (Provoost and Bosch, 2022)), (2) global datasets (UNEP-WCMC, 2020; Assis et al., 2024), (3) regional datasets covering areas and species underrepresented in previous sources (e.g., Sudo and Nakaoka (2020)), and (4) a literature-derived dataset.

#### Global databases

We retrieved occurrence record data by querying with the taxa listed in Table 1 from the Global Biodiversity Information Facility (GBIF) and the Ocean Biodiversity Information System (OBIS) on 03/09/2025. Data was downloaded in two temporal segments (2000–2010 and 2010–2020) for a perfect match with the occurrence data. The query was conducted with the functions ‘*occ_search()*’ and ‘*occurrence()*’ from *rgbif* v3.8.2 Chamberlain et al. (2017) and *robis* v2.11.3 Provoost and Bosch (2022), respectively.

For GBIF data, the following filters were enforced at download: Has coordinate = TRUE, Has geospatial issue = FALSE, Occurrence status = PRESENT, Basis of record ̸= *“FOSSIL_SPECIMEN”, Coordinate uncertainty in metres* ≤ 2000; = NA; ̸= *c(301, 3036, 999, 9999)*. Only species-level occurrences were considered. A doi was generated at each download (2000–2010: [DOI: 10.15468/dl.twmjcn]; 2010–2020: [DOI: 10.15468/dl.xdqkg6]). OBIS data were queried using identical temporal and taxonomic criteria, with post-download filtering via *dplyr* to enforce consistency with the GBIF workflow. Taxa was queried with taxon keys from GBIF, which had been previously extracted via the *rgbif::name_backbone()* function, and from OBIS using the species names.

We restricted records containing explicit uncertainty radii to a maximum uncertainty of 2 km, excluding problematic placeholder values (i.e. 301, 3036, 999,9999) and including NAs. Although a higher uncertainty radius could be used for global studies, a relatively low radius was enforced as a proxy of observation accuracy (i.e., taxonomic identification accuracy). NAs were allowed, as not incorporating them would have led to a significant record drop and other sources do not include this metric.

##### Global datasets

The UNEP-WCMC dataset was developed during the production of Short and Green (2003). Although this source is used extensively in the seagrass modeling literature, we used a very restricted subsample, as half the records do not have assigned year data. Following Short and Green (2003), most records are historic data collected along the XX. century, which we did not deem appropriate for modeling with our temporal extent (2000-2020). Inclusion of the full dataset could have potentially led to spurious association of temperatures different from the conditions at the time of detection and may reflect presence occurrences where local extinctions have taken place. The original dataset was reformatted to DarwinCore. Taxa labelled as *“not reported”* and *“unspecified”* were eliminated.

**Table 1.**
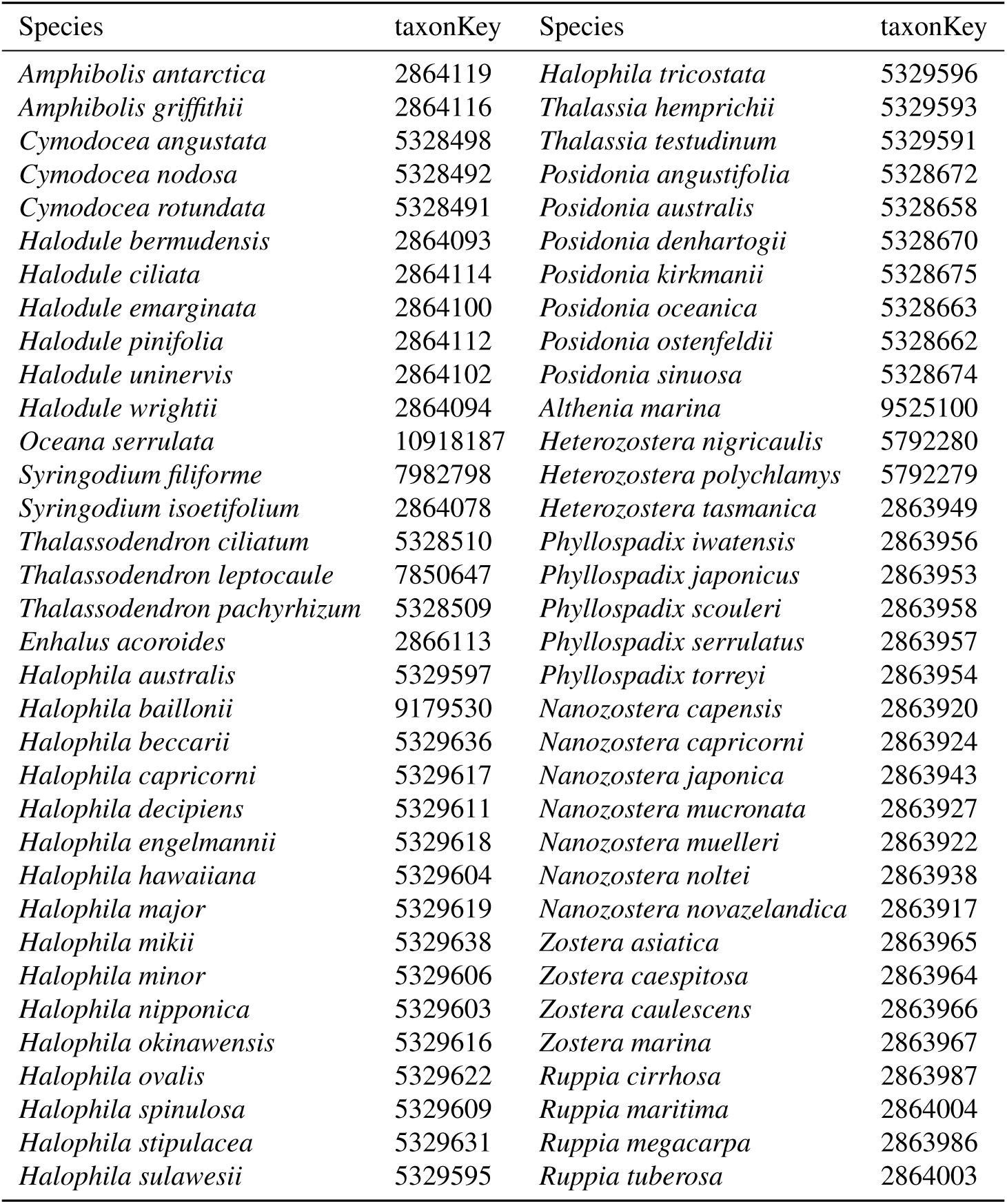
Taxonomic entities queried for the data download from GBIF and OBIS. Taxon keys were used for download from GBIF.

The Assis et al. (2020) dataset (’finalDataBaseSPruned’) was filtered to retain only essential variables: c(dataset Name, basis Of Record, occurrence Status, individual Count, organism Quantity Type, year, month, decimal latitude, decimal longitude, coordinate uncertainty in meters, accepted name, license). We filtered out records with no coordinates or beyond our spatial uncertainty thresholds. Both datasets were separated into decadal subsets (2000-2010 and 2011-2020).

##### Regional datasets

We adapted the dataset in Sudo and Nakaoka (2020) to a DarwinCore through a *dplyr* R pipeline. Both datasets were separated into decadal subsets (2000-2010 and 2011-2020).

##### Supplementation with the literature

The targeted literature search resulted in +2000 occurrence records from over 280 peer-reviewed articles, technical reports, and grey literature items (Supplementary Material II), focusing on data-sparse regions or species. Searches were conducted using Web of Science, Google Scholar, and Google, with combinations of terms “seagrass*”OR”target species” AND “target region”. Local-language sources were consulted where possible. Coordinates were derived from toponyms and projected to WGS 84 (EPSG:4326). A visual uncertainty threshold of 2.5-3 km was enforced, given the lack of numerical uncertainty information in most sources, and the acceptability of this threshold given the raster resolution.

Observation years were extracted directly from publications; if not specified, sampling in year prior to publication was assumed. For multi-year studies, only the first year within the study window was recorded. ‘Year’ was used for later splitting the dataset in two. As not all source materials had the month of sampling, all 2010 records were assigned to the 2000–2010 decade, instead of making the cut in June-July.

This compilation is published under CC-BY in compliance with the EU *sui generis* right, allowing all kinds of reuse, extensions and modifications, depending on the original data permissions.

##### Curation and merge

Unlike workflows that download and merge datasets prior to standardization and cleaning (e.g. Dorey et al. (2023)), we decided to implement the same pipelines to each dataset independently (at query when possible), to reduce data volume and computational load.

License permission was ensured across datasets with the *unique()* function of basic R, identifying the following permits: detected: CC 0, CC 0 (Ca), CC 1.0, CC-BY, CC-BY 3.0, CC-BY 3.0 (Au), CC-BY 4.0, CC-BY-NC, CC-BY-NC 4.0, CC-BY-NC-ND, CC-BY-NC-SA, CC-BY-ND, and CC-BY-SA. The license of non-database sources was ensured manually. Data licensed under no derivatives (CC-BY-ND and CC-BY-NC-ND) and Share-Alike was eliminated due to potential licensing issues. A total of 29 occurrences were eliminated.

The Geodetic System was also checked with ‘*unique()*’ across big datasets and ensured manually otherwise. GBIF downloads are always normalized to WGS84 (EPSG:4326) (Chapman and Wieczorek, 2020). Geodetic consistency was ensured by reprojecting all coordinates to WGS 84 (EPSG:4326).

All datasets (GBIF, OBIS, UNEP-WCMC (2020); Assis et al. (2020); Sudo and Nakaoka (2020); McKenzie and Yoshida (2020), and the literature supplement) were separated into decadal subsets: 2000–2010 (for record dates ≤ 30/06/2010) and 2011–2020 (for record dates ≥ 01/07/2010). Literature-derived records within ± 4–5 years of a decadal boundary were assigned to the nearest decade to retain ecologically relevant but scarce observations. We acknowledge this as a deliberate compromise between retention of key data and precision. Incomplete records (e.g., missing month/day) were defaulted to year-only precision, with 2010 observations included in the 2000–2010 subset.

Prior to merge, all datasets were simplified to key columns to be kept (i.e., decimalLatitude, decimalLongitude, species, family, year and license) or created (i.e., ‘source’ and ‘decade’) following, when possible, DarwinCore naming format. Subsets of the same decade were merged together. Upon merging, deduplication (via ‘*distinct()*’) retained unique combinations of coordinates, species, and decade, eliminating redundant records and collapsing repeated surveys at identical locations into single random-year events.

Taxonomic harmonization was enforced using the species list in Supplementary Material III, resolving synonyms, nomenclatural issues, species complexes, and family names.

A distribution map was generated for each species to identify putative misidentification following expert criteria. Those were manually targeted and eliminated. Automatised cleaning was run with ‘*coordinateCleaner*’ (Zizka et al., 2020). Only 8 records were eliminated in both steps.

##### Temperature data curation and extraction

The temperature was assigned by decade. To avoid a high proportion of non-retrieved temperatures in occurrence points falling outside of the raster (e.g., occurrence points in estuaries, embayments, fjords, or harbours that are not completely covered by the raster), the raster edges were expanded along the coastline. We performed this neighbourhood (focal) gap-filling Fig. 1 on each decadal SST raster by (i) defining a circular moving window of a diameter of 4 cells, (ii) forcing uniform weights inside that window, and (iii) computing a local mean only at cells that are missing (NA).

**Figure 1.**
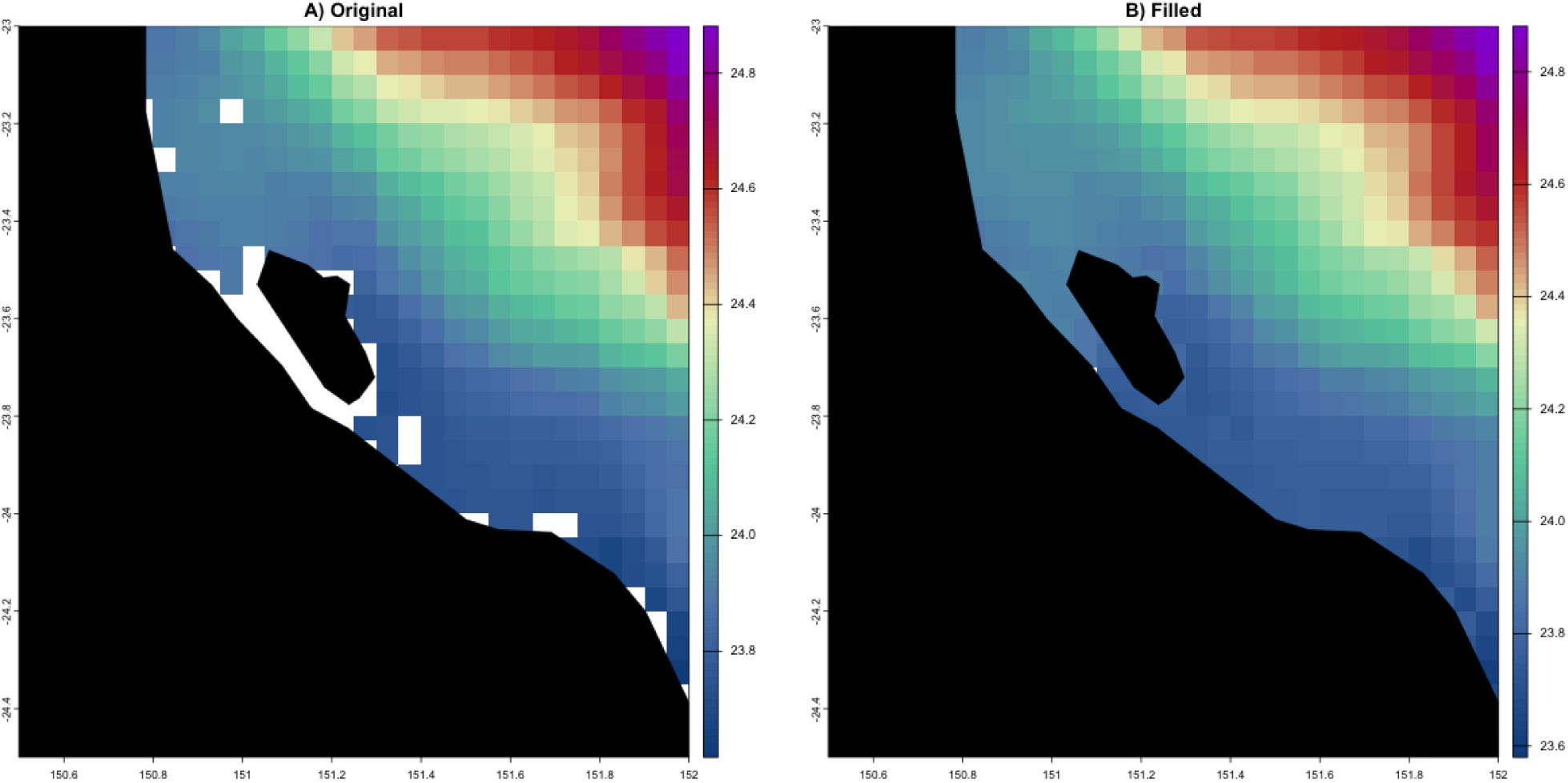
Comparison of the original and filled raster, using the SST mean (2000-2010) from BIO-Oracle v3. The region proximal to Gladstone (Australia) is shown, because of the many *Halophila* and *Nanozostera* observations potentially lost, if this extension is not computed.

Gap filling was performed independently for each decadal SST raster using a focal neighbourhood approach implemented in the Hijmans et al. (2026) package in R. A circular moving window was defined with a diameter of four raster cells (corresponding to 0.2°, given the BIO-ORACLE v3 spatial resolution of 0.05°). The circular neighbourhood was constructed as follows:

~~~
Wcirc <-focalMat(envstack, d = 4 * res(envstack)[1], type = “circle”)
~~~

This step generated a kernel in which all cells whose Euclidean distance from the centre was less than or equal to the specified diameter were included in the neighbourhood. To ensure an equal contribution of all neighbouring cells, the kernel was converted to a uniform (unweighted) window:

~~~
Wcirc[Wcirc > 0] <-1
~~~

Missing raster values were then filled by computing the arithmetic mean of neighbouring cells within the circular window. The operation was applied only to raster cells with missing values (NA), leaving all pre-existing values unchanged and avoiding an artificial smoothing of the original data:

~~~
filledraster <-focal(envstack, w = Wcirc, fun = mean, na.policy = “only”, na.rm = TRUE)
~~~

The ‘na.policy = “only”’ option restricted calculations to focal cells with missing values. ‘*na.rm = TRUE*’ excluded missing neighbours from the computation. If all neighbouring cells within the window were missing, the focal cell remained empty. This approach increased spatial coverage at the raster boundaries (up to ∼2.3 km from the original edge) while minimizing interpolation artifacts.

Occurrence points were converted to a SpatVector in the same coordinate reference system as the BIO-Oracle v3 rasters (EPSG:4326). The local SST at each point was extracted with the *’terra::extract’* function, through bilinear interpolation. The resulting ‘NaN’s were converted into ‘NA’s. Then, NAs (land points and inaccurate locations) were eliminated. Then, both decadal subsets were combined with ‘*bind_rows()*’.

**Table 2.**
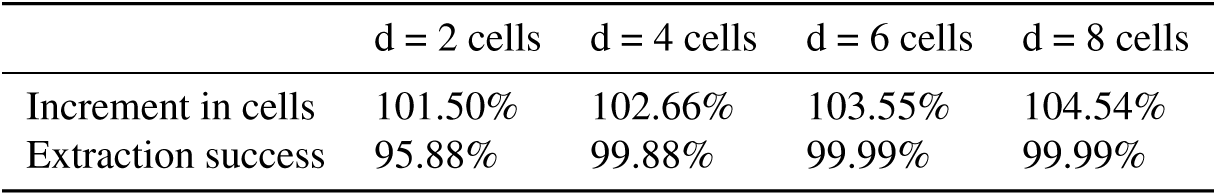
Increment in filled cells (compared to the original number of filled cells) and occurrence points with a successful temperature extraction as focal window size is increased.

A sensitivity check was run to check the impact of different window sizes. The only variation in the results was the increase of occurrence records with a successful temperature extraction. d = 4 was kept to ensure the capture of coastal occurrences whilst filtering out land records. Table 2 shows the increase in filled cells as bigger focal windows are used, along with the increased success in data extraction.

### 1.2 Calculation of the range size

The range of each species *s* was quantified using the range size descriptor (*D_s_*), defined as the diameter of a circular approximation of its range. This metric was derived by clustering all raster cells (resolution = 0.5°) that contained at least one occurrence record on a single surface value and computing the diameter (*D*) of a circle with equivalent area. To avoid the loss of geographic edge information, *D_s_* was calculated exclusively from the unthinned (raw) dataset and subsequently treated as a fixed species-specific constant. Thinning is expected to constrict the range size as different random marginal points are selected in each thinning iteration. In our calculations, the resolution (res = 0.5 °) was selected to find a balance between connecting sparse occurrence records and not overstretching the distribution area by incorporating uninformative areas (i.e. oceanic water and land).

First, occurrence points were converted to a spatial object using the WGS84 coordinate reference system (EPSG:4326). Independently, a global raster grid (0.5° cell size; ∼55.5 km at the Equator) was generated. To compute cell areas accurately, the raster grid was reprojected to the World Cylindrical Equal Area projection (EPSG:6933), and the area of a cell was calculated in square km^2^. However, following steps used the coordinate reference system of the occurrence data (EPSG:4326). For each species, occurrence points were rasterized ‘*rasterize(pts_sp, r, fun = “count”, background = 0)*’ and the total number of occupied cells was calculated ‘*global(D_sp > 0, “sum”, na.rm = TRUE)*’. Occupancy area was estimated by multiplying the number of occupied cells by the area of a single cell in km^2^. To facilitate comparison across species with distributions of different form (fragmented, along heterogeneous coasts…), this two-dimensional area was converted into a one-dimensional metric by assuming a circular geometry and computing the diameter, transforming an area of occupancy metric into an extent of occurrence metric comparable across species. The transformation was run with the following formula:

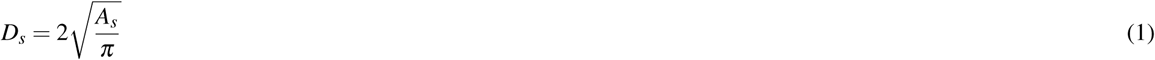

where *D_s_*is the diameter for species *s*, *A_s_* represents the occupied area (km^2^) estimated from the rasterised occurrence cells, and the other elements are derived from the circle area formula *A* = *πr*^2^, which can be simplified into 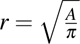 to calculate the radius, which, by definition will have half the value of *D_s_*.

#### Resolution selection

Raster resolution was selected arbitrarily as a compromise between excessive range fragmentation at very fine resolutions and artificial range inflation at coarse resolutions, particularly into non-informative oceanic or terrestrial areas. Six resolutions (0.1°, 0.3°, 0.5°, 1°, 2.5°, and 5°) were evaluated by adapting the distribution-size calculation workflow to iterate across resolutions. For each resolution, species were ranked according to their estimated range size (*D_s,res_*). As a null reference, an additional ranking was generated by randomizing species order using a fixed random seed (’*set.seed(12)*’) for replicability.

The ranked species lists were compared with Kendall’s rank correlations, using the numerical rank position of the species in each resolution according to its predicted range size. Pairwise comparisons between ranked lists were conducted using Kendall’s rank correlation coefficient (*τ*), quantifying the degree of agreement in the relative ordering of species. Correlation values were calculated for all pairs of resolutions and visualized as a correlation matrix (Fig. 2). The correlation analysis showed a high level of consistency across resolutions. Even when comparing the finest resolution (res_0.1) with a resolution 25 times coarser (res_2.5), the correlation remained high (*τ* = 0.84).

**Figure 2.**
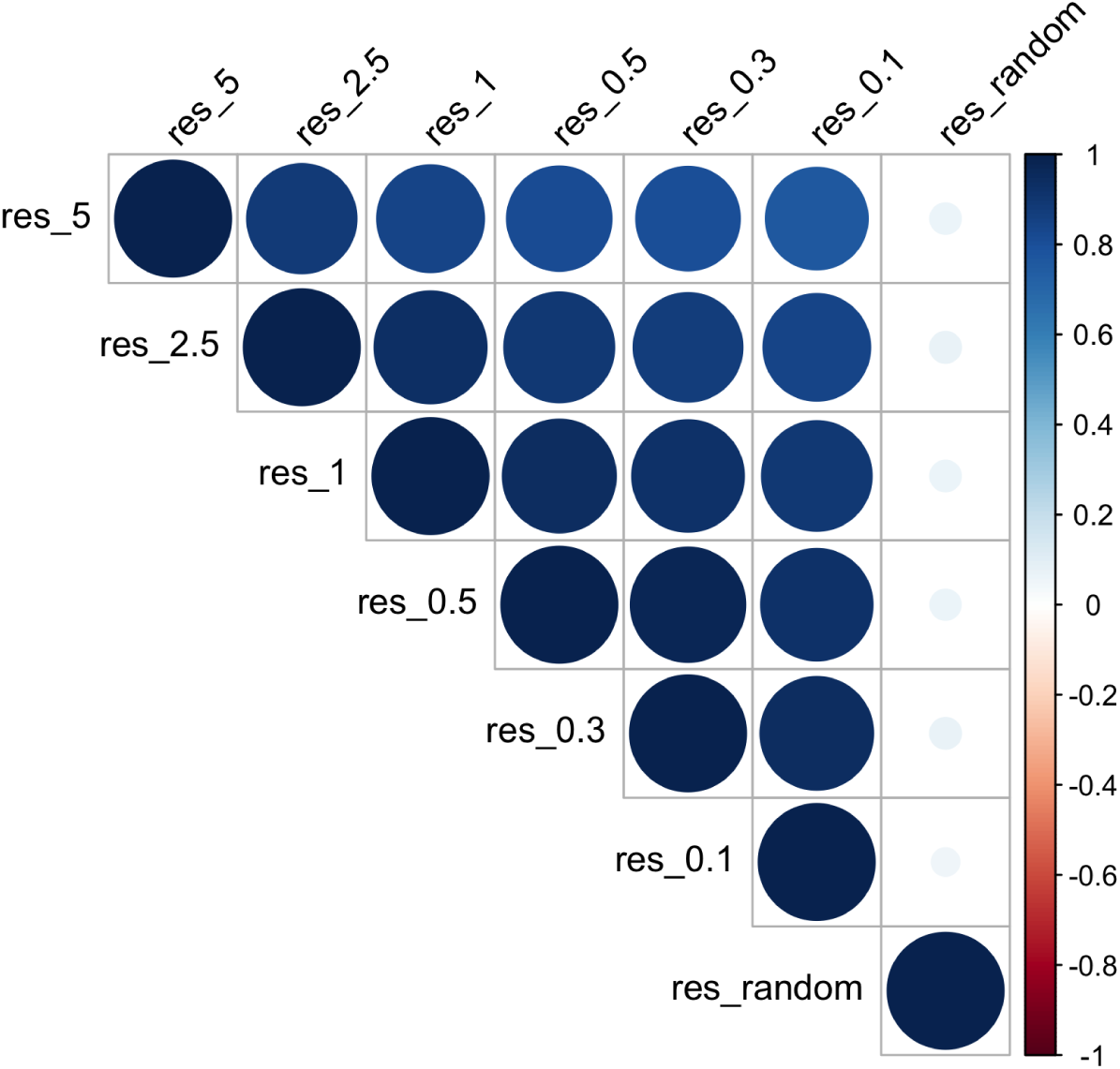
Kendall’s rank correlation matrix comparing species rankings derived from predicted range sizes at each original spatial resolution. Species were ranked according to the resolutions employed (0.1, 0.3, 0.5, 1, 2.5, and 5) and a random species order

### 1.3 Thinned datasets

From the original raw dataset, 13 datasets were independently generated by stratified thinning with Mestre-Tomás (2026). Thinning was iterated over species. Five trials were conducted per iteration, from which only the most conservative (more points retained) was kept. The original dataset was thinned with each item in the following parameter set Θ:

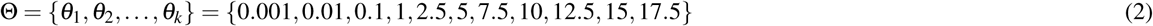

The parameter set was generated by including extremely small and big parameters for a better calibration.

### 1.4 Calculation of niche stabilization

The primary objective of the proposed method is to reduce spatial and sampling bias through data thinning while preserving the suitability of the dataset for niche characterization. Our data and focus have been directed towards the thermal niche, and a widely used temperature-based niche descriptors, the thermal optimum, which is often characterised by the median temperature (Burrows et al., 2020). Although alternative estimators exist, such as the average, the mode or the maximum density of fitted distributions, the median offers a computationally efficient and sufficiently accurate approximation of the niche optimum, whilst being less sensitive to extreme values. Moving statistics offer the opportunity to estimate trends and trajectories, by calculating a descriptor with successive selections of randomly selected data points along a data series (Pierre and Louis, 2012). Although the moving average is more commonly used, we decided to use the moving median for the reasons stated above.

The stabilization procedure includes (i) data input and initialization (including modeling settings), (ii) the formulation of custom functions, (iii) the calculation of replicate moving median trajectories, (iv) the computation of stability metrics, and (v) summarization and visualization. Then, the stability output can be used to calculate both methods.

### Data input and initialization

This stage required four primary inputs: a vector of species names, a list of thinned datasets, an ordered parameter vector used to facet and organize outputs, and a two parameter sets. Moving median settings and rules include (i) a fixed random seed for reproducibility, ‘*set.seed(123)*’; (ii) the maximum subsample size of observations per thinned dataset, ‘*nsamp* `<-` *500*’; (iii) number of replicate subsampling runs, ‘*B* `<-` *10*; and (iv) sampling without replacement, ‘*replacem* `<-` *FALSE*’, to prevent the artificial inflation of temperature frequencies. Tolerance settings and rules include (i) the maximum absolute difference from the final median (in °C), ‘*tol_value* `<-` *0.3*’; (ii) maximum confidence interval width (95% CI ≤ 0.5 °C), ‘*tol_width* `<-` *0.5*’; and (iii) rule that “stability must be maintained for at least 20 consecutive”, n_obs ‘*window_run* `<-` *20*’.

### Custom functions

Two custom functions were created to pull parameter values from the thinned datasets, (i) `stabilize_sp_df`; and determine stability thresholds, (ii) `compute_threshold`.

i. `stabilize_sp_df` computed replicate trajectories of the moving median for each species–parameter combination ((*s, θ*) pairs), using the following function definition:

~~~
stabilize_sp_df <-function(sp, df, B, nsamp, replacem)
~~~

For a given species, the function subsets the corresponding temperature records (annual SST mean), excluding cases ((*s, θ*) pairs) with fewer than three observations to ensure reliable estimation. For each replicate *b* (from a total of *B* replicates), a random sub-sample of size `m = min(nsamp, n)` was drawn without replacement. In this way, if the number of species observations in the thinned dataset was greater than `nsamp` (set previously to 500), `nsamp` observations were sampled. Otherwise, all available records were used. Replacement was disabled, following the original input (`replacem <-FALSE`). Then, *m* row indices were sampled randomly from 1 to the maximum using `sample.int`. For each subset, a cumulative moving median was calcu-lated using the `slide_dbl` function from the `slider` package (Vaughan, 2024). The function calculated the median of the temperature variable while ignoring NA values and using a window that included all previous observations (i.e., a cumulative me-dian): `slide_dbl(x, ∼ median(.x, na.rm = TRUE), .before = Inf)`. The first observation was set to NA because a median was required to contain at least two values: `mov_med = ifelse(n_obs < 2, NA_real_, mov_med)`. To store the results, the function created a new data-frame-like object (a tibble) containing the replicate index, the cumulative number of observations, and the corresponding moving median.

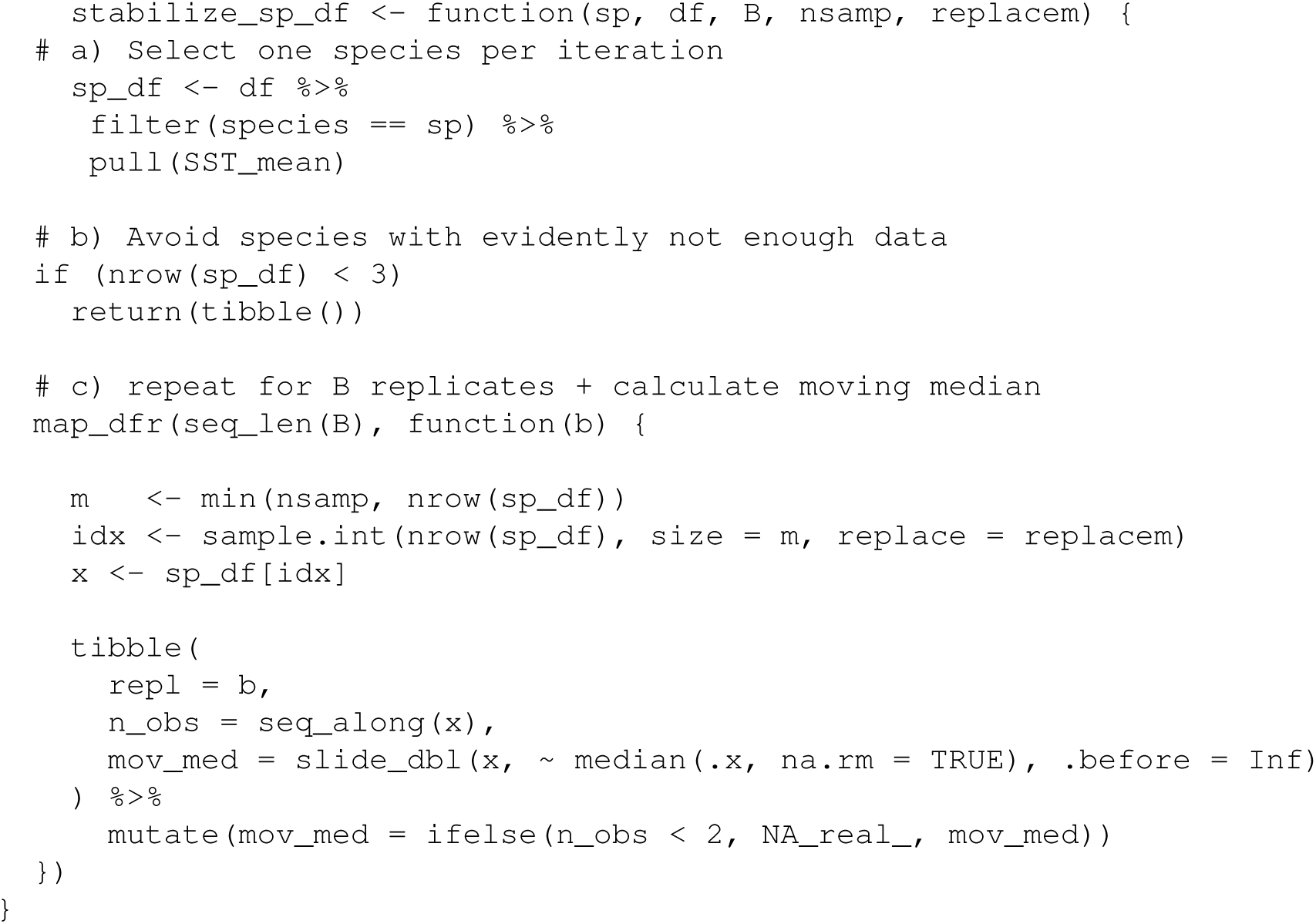

(ii) `compute_threshold` identifies the stabilization threshold (*ST_θ,s_*) using replicate summaries and predefined tolerance criteria. (*s, θ*) pairs with fewer than ten observations were automatically excluded. Otherwise, the “final thermal optimum” was approximated as the mean of the last `tail_n` moving-median values, where `tail_n <-min(20, nrow(summary_df))`. At each step, the following variables were computed: (i) `diff_final`, the absolute difference between the moving median at each step and the tailed final median; (ii) `CI_width`, the difference between the upper and lower bounds of the confidence interval; and (iii) `stable_flag`, a logical stability flag indicating whether the moving median was stable according to both metrics and their respective tolerances. The stability flag was based on two logical expressions. The first, `diff_final <= tol_value`, checked whether the absolute difference between the moving median and the final moving median was less than or equal to `tol_value` (0.3 ^◦^C, as previously specified). The sec-ond, `CI_width <= tol_width`, checked whether the width of the 95% confidence interval was less than or equal to `tol_width` (0.5 ^◦^C). If both conditions were true, the moving median was considered sufficiently close to the final value, as expressed by `stable_flag = (diff_final <= tol_value) & (CI_width <= tol_width)`. Using these summary metrics, *ST_s,θ_* was calculated. To this end, the lengths of consecutive `TRUE` and `FALSE` runs in `stable_flag` were recorded using `rle()`. The final index of each `TRUE` or `FALSE` segment was then obtained using `cumsum()`. The first sequence meeting the minimum-length requirement was defined as the stabilization threshold. This was established by detecting the start index of each `TRUE` segment using `idx_end - run_seq$lengths + 1` and then identifying the `TRUE` segments of sufficient length using `which(run_seq$values & run_seq$lengths >= window_run)`. For each valid case, the function returned the observation count at which stability was first achieved (*ST_s,θ_*); the fraction of observations at which stability was reached (`p_stable`); the tailed final value of the moving median (`final_mov_median`); the difference between the upper and lower confidence-interval bounds at *ST_s,θ_* (`width_at_stable`); and a logical indicator that detemines whether stability was achieved (`stability_reached`).

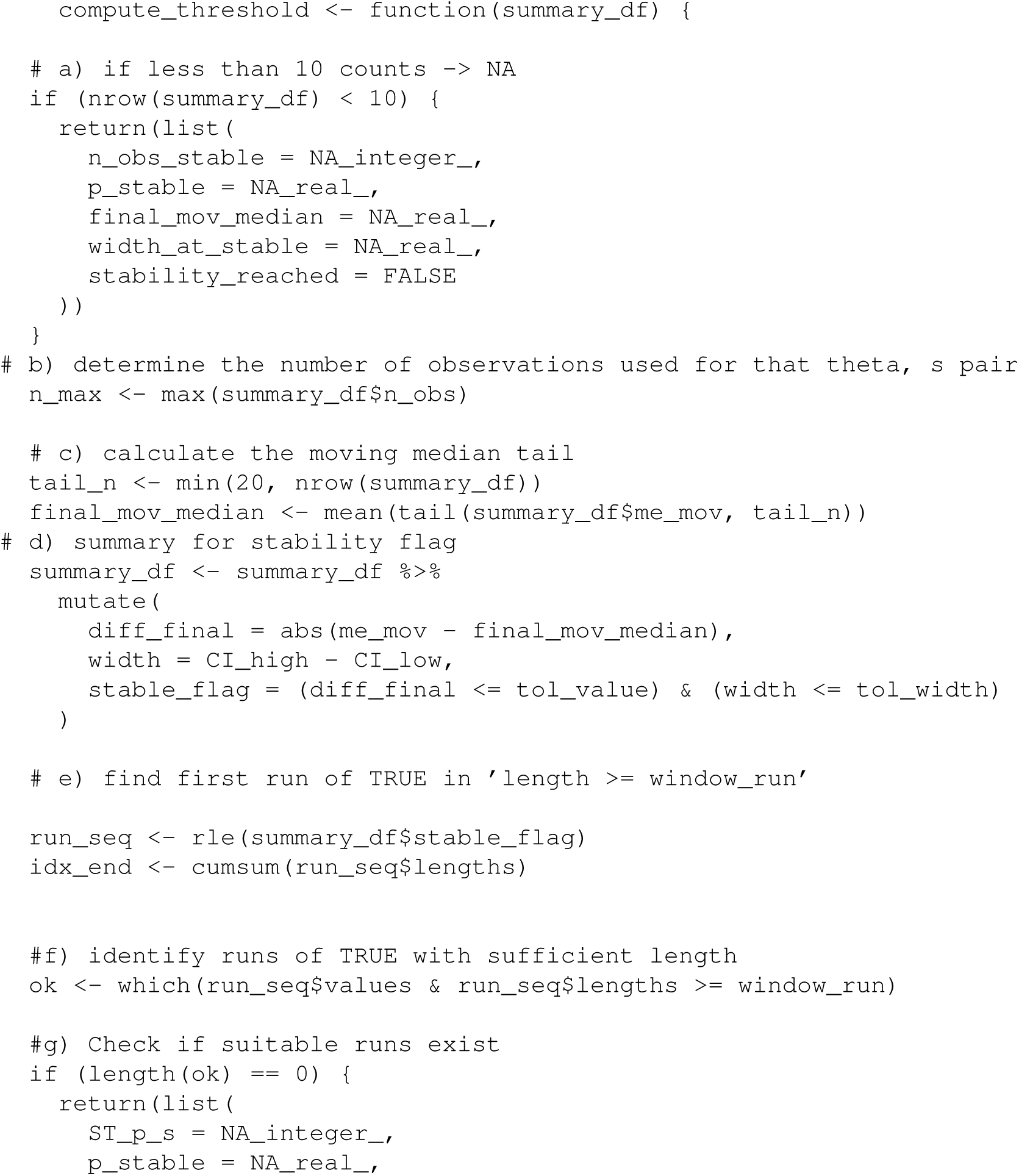
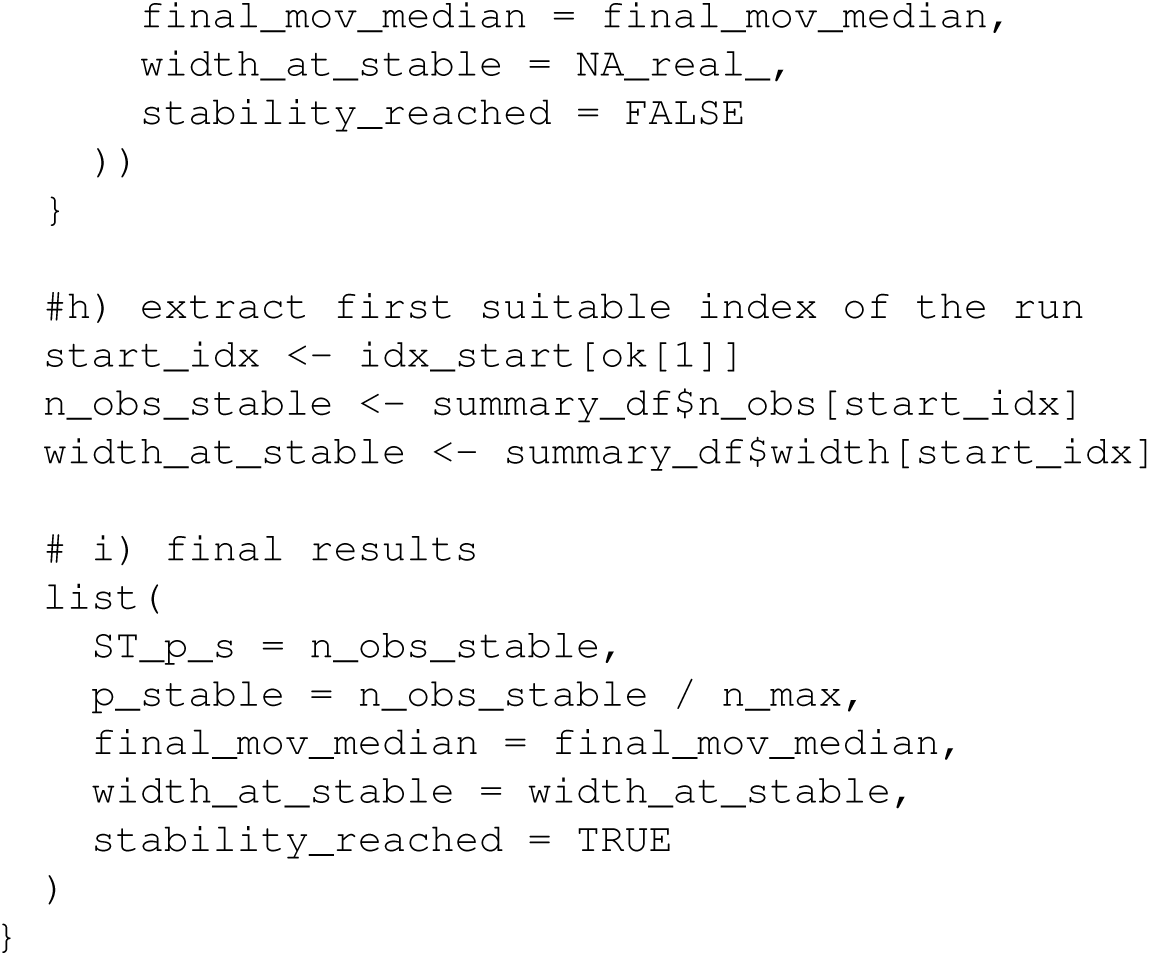

*Stabilization* The stabilization run, where the previous functions were computed, has three sequential parts: (i) replicate computation (replicated moving median calculation), (ii) thresholding, and (iii) replicate aggregation. At the end of the stabilization run, the third input component for SVM, the boolean stability, was generated.

First, moving median trajectories were calculated independently for each species across all thinned datasets (each *s, θ* pair) using nested loops. Lookup vectors were created to later order the outputs, considering the range size (*D_s_*) and raw occurrence counts. To compute and store the results of each (*s, θ*) pair two hierarchical loops were run: an outer loop for the species and an inner loop for the thinned datasets. The inner loop was run for each thinned dataset *θ*, filtering out (*s, θ*) pairs with insufficient data. The thinning parameter value was extracted from the dataset name with the custom function ‘i’ (`parse_param()`). The custom function ‘ii’ (`stabilize_sp_df`) was used to calculate replicate calculations of the moving median for this (*s, θ*) pair. Important metadata was added to the outputs (e.g. species name, thinning parameter). Results of the replicate trajectories of each species *s, θ* pair were compiled in a list.

The stability thresholds (*ST_s,θ_*) were calculated with a similar nested loop. When data counts for each *s, θ* pair were deemed sufficient (*>*3), data was summarised (including the median for all ‘moving median’ replicate trajectories and their 95 % CI bounds). The summaries are used as the input of the (iii) custom function `compute_threshold`. A ‘`stability_results_df`’ data frame is generated including species name, parameter value, maximum number of observations considered, and the output variables of the custom function (see the previous text on the custom function ‘iii’).

Plots were generated for visualization. In the first chunk, plots were computed for each of the *s, θ* pairs, plotting the replicate moving median trajectories against the number of observations considered, grouping the data by replicate and faceting by parameter. Summarised plots were also generated, by generating summaries from replicate data (median of the moving median and 95% confidence interval) and plotting them with the *ST_s,θ_* vertical values.

### 1.5 Sensitivity analysis for different tolerance settings

Default tolerance settings were set at 0.3, 0.5, and 20, for `tol_value`, `tol_width`, and `window_run`, respectively. A sensitivity analysis evaluating 27 combinations of stabilization parameters was performed by counting the number of species that are never stable (regardless of the thinning parameter) for each combination 3. This value was normalised to the total number of species (N = 54) to get the proportion. This proportion was normalised in turn to the value (proportion = 33.27 %) showed that the number of species reaching stability was unaffected by the tolerance around the final moving median (tol_value). In contrast, stability increased significantly with increasing confidence interval tolerance (tol_width) and decreased with increasing stability-window length (window_run). These results suggest that the stabilization procedure is robust to the definition of the target median but moderately sensitive to the stringency of precision and persistence criteria.

**Figure 3.**
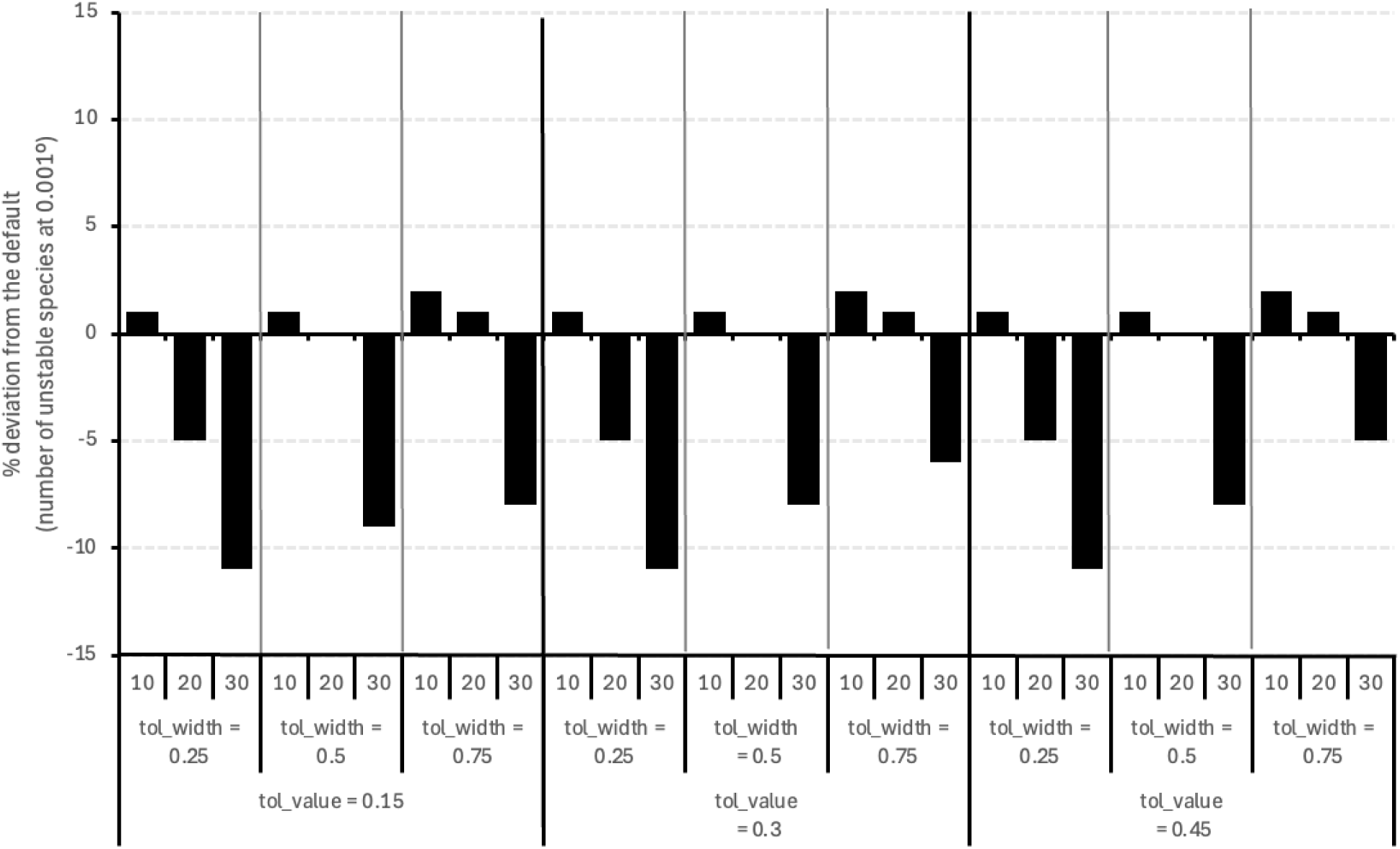
Sensitivity analysis for 27 combinations of the tolerance settings. Combinations were compared by normalising the percentage of species that never reach stabilization and reporting the changes in this number.

### 1.6 Biogeographic-size dependent thinning parametrization (BD-TP)

A data frame with the variables calculated in previous steps and used as the SVM input. Then, a Support Vector Machines algorithm (Vapnik, 1995) is used to extrapolate hyperplane values. These values represent the highest thinning parameter that can be used prior to niche characterization destabilization. The components of a simple linear model were computed analytically to enunciate a linear equation in which *D_s_* is used to predict *θ_opt_* (’*θ* = *m* · *D_s_* −*C*’).

#### Application of Support Vector Machines

A data frame was built with previous outputs: the parameters in numeric format, *D_s_*, and a boolean stability variable. With this input, the SVM model was built with `svm()` from the `e1071` package (Meyer et al., 2024). The range size and the parameter were included as predictors and stability as the class label ‘`svm(stab ∼ diameter_km + parameter_num, …)`’, with a linear kernel. The ‘`probability = TRUE`’ and ‘`scale = TRUE`’ arguments were used to allow the later extraction of probabilities and scaling features internally, respectively.

With the SVM output two prediction grids were generated. The first, created a fine grid of points covering the range of the predictors (*D_s_* and *θ*) to later visualize the decision boundary. With the ‘`predict()`’ function, a categorical value is created to give the predicted class for each grid cell. The second, predicts decision values, that is, how far each point is from the decision boundary.

The SVM was represented as a ggplot graph, using the ‘`scale_fill_gradient2`’ argument to fill the plot, ‘`geom_contour`’ to plot the 0 and the ±1 boundaries, and `’geom_point’` to represent the stability for each *s, θ* pair. The order of the axes (y = *D_s_*, x = *θ*) is plotted inversely to the mathematical formulation of the linear model (*θ* = *f* (*D_s_*); *θ* = *m* · *D_s_* +*C*) for conceptual clarity.

#### Linear model extraction of the hyperplane

We extracted the hyperplane analytically from the SVM model coefficients. The weight vector of the decision function was first reconstructed from the support-vector coefficients and support vectors, and the intercept was obtained from the model parameter (*ρ*). Because the SVM was fitted with standardized predictors, The predictor-specific centering and scaling factors used internally by the SVM were then retrieved and used to transform the hyperplane coefficients back to the original measurement units. In original units, the decision boundary can be written as:

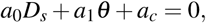

where (*D_s_*) is the range-size descriptor and (*θ*) is the thinning parameter. Rearranging this expression for (*θ*) yielded the linear BD-TP rule (*θ* = *mD_s_* + *c*_0_), with (*m* = −*a*_0_*/a*_1_) and (*c*_0_ = −*a_c_/a*_1_) and with *D_s_*and *θ* as the independent and dependent variables, respectively. The resulting slope and intercept were retained as the analytical estimates of the SVM decision boundary and were used to parameterise the BD-TP rule.

A new data frame with the species list and their *θ_opt_* was created with *θ_opt_*`= pmax(m * diameter_km + c0, 0.001)`. Through this formula, the parameters are predicted regarding the relationship with the range size, but are capped in 0.001° as a minimal parameter. This cap avoids the application of theoretical negative parameters. It also ensures that, extremely close occurrence records are not are considered, even if the predicted parameter is lower than 0.001.

The uncertainty and validity of the method were tested with bootstrapping and Leave-One-Species-Out as specified in Supplementary Material 5.4 and the main script available in the Supplementary materials.

### 1.7 Last stable thinning parametrization (LS-TP)

The data frame generated in the stabilization run (’`stability_results_df`’) is used to find the highest parameter value for which stability is still kept. A custom function was used along `coalesce()` and `cur_data()` to extract the maximum parameter value for the subset of stable parameters. The selection rule was also capped at a minimum *θ* = 0.001.

### 1.8 Method application

The downstream effects of the proposed methods were explored with some of the species with sufficient data for ecological modeling. Among these, a single species, *Phyllospadix torreyi* was selected for simplicity.

Occurrence and environmental data was prepared for GLOSSA and thermal niche plotting. GLOSSA is an R Shiny application implementing Bayesian Additive Regression Trees (BART) for marine species distribution modelling (Mestre-Tomás et al., 2026). A specific data formatting is required for its biological (occurrence data subset) and abiotic (environmental rasters sourced from BIO-Oracle v3) is needed. Occurrences were reformatted by retaining: species name, decimal longitude, decimal latitude, decade, presence/absence indicator, and temporal timestamp. All occurrence records represented presences and were therefore assigned pa = 1. The two decades (2000-2010 and 2010-2020) were assigned the timestamps 1 and 2, respectively. Five environmental variables were considered: bathymetry, temperature, photosynthetically active radiation (PAR), slope, and salinity. We used the decadal average value with a resolution of 0.05 degrees. Raster edges were expanded following the same procedure as in the “Temperature data curation and extraction” subsection. Environmental layers were spatially restricted to the region relevant to *P. torreyi* using the shape of the ecoregions per Spalding et al. (2007) where the species occurs. Environmental rasters were cropped to the polygon extent, masked to retain cells within the selected region, and spatially aggregated using a factor of 10, with the mean used as the aggregation function. The resulting GeoTIFF files were aggregated and exported to a .zip file following the documentation of GLOSSA.

A second *P. torreyi* dataset was generated to examine how the thinning parametrization approaches respond to stronger spatial sampling bias. Artificial sampling bias was introduced around three major cities within the species’ geographical distribution: San Diego, Los Angeles, and San Francisco. A 20 km buffer was generated around these cities using a 3857 crs, allow buffers to be defined in metres. 300 artificial occurrence points were generated randomly in the combined area encompassed by the three buffers. Biased data was prepared for GLOSSA following the same steps used for the original data. For both the original and the biased dataset, three thinning distances and the unthinned occurrence data were tested. Spatial thinning was run with GeoThinneR, conducting five thinning trials, from which only the most conservative one was retained. A fixed random seed of 567 was used to ensure reproducibility. The datasets resulting from thinning were reformatted as well to GLOSSA’s requirements.

The thinning treatments applied to the original *P. torreyi* dataset were: 0.05° thinning, representing thinning at the environmental raster resolution; LS thinning: 0.1°; BD thinning: 0.6°; and raw, representing the original unthinned dataset. BD and LS thinning parametrization was conducted independently for the original and biased datasets. Only BD resulted in different parameters to be used (ori - 0.6°, bias - 1°).

Habitat suitability models were run with GLOSSA. Separate HSM outputs were generated for each combination of dataset type (original or artificially biased) and thinning treatment (raw, 0.05°, LS, or BD). Pseudo-absences were generated randomly within a 0.2^◦^ buffer-restricted area aroudn presences using a 1:1 pseudo-absence-to-presence ratio. Model performance was assessed through 10-fold spatial block cross-validation, with block size determined from the spatial autocorrelation of model residuals, using all evaluation metrics available in GLOSSA.

Median habitat-suitability predictions and their associated standard-deviation rasters were imported back to R, where the final plots were generated. Model-evaluation metrics generated by GLOSSA were also imported and plot in R. Six evaluation statistics were considered: F-score, accuracy (ACC), balanced accuracy (BA), true skill statistic (TSS), continuous Boyce index (CBI), and area under the receiver operating characteristic curve (AUC). The distribution of cross-validation results was visualised using boxplots with individual replicate values superimposed. The evaluation metric value using the entire subset was shown and represented as an asterisk. Results were separated according to the data type (original or artificially biased).

The effect of thinning on the environmental conditions represented by the occurrence data was examined using the mean decadal SST using the *P. torreyi* original and biased data, according to the four tested thinning parameters. Bilinear interpolation was used for raster extraction. The ggplot2 `geom_density` function was used to compute and plot the thermal curves, with its default Gaussian kernel and optimal smoothing bandwidth selected with biased cross-validation (**?**). To evaluate the sensitivity of the estimated thermal niche to the occurrence sample, a repeated resampling was performed 50 times, randomly assigning 60% of occurrence records to the training subset in each replicate. Density curves were plotted separately for the raw, 0.05°, LS, and BD occurrence datasets.

## S3: TAXONOMIC, NOMENCLATURE AND SPECIES SELECTION STATEMENT

This annex justifies (1) the taxonomic limits of the study; (2) the selection of species names, according to the currently most accepted nomenclature and taxonomy; (3) the classification of species in groups/complexes; (4) and ensures result reproducibility and transparency. Synonyms were checked in accordance to Plants of the World Online (POWO; https://powo.science.kew.org/, accessed before Dec/2025), World Flora Online (WFO; https://www.worldfloraonline.org/, accessed before Dec/2025) and Algaebase (https://www.algaebase.org/, accessed before Dec/2025). Taxonomic discrepancies among POWO, WFO, Algaebase, the Global Biodiversity Information Facility (GBIF), and other relevant literature items are discussed here.

We followed the definition of seagrasses presented in Hartog and Kuo (2007), considering marine angiosperms within the order *Alismatales* that live subtidally or intertidally in nearshore systems, having evolved both vegetative and reproductive adaptations to saline conditions. Following the broadest sense of this definition, we implemented a semi-conservative approach which included both seagrasses *sensu stricto* and other phanerogams that are important in the coastal communities they form (seagrasses *sensu lato*). We excluded typically freshwater species that can occur within seagrass ecosystems as seagrass associates or facultative members (Green & Short, 2003), but included *Althenia marina* and the *Ruppia* species whose distribution is most associated to the coastline and which are often encompassed under the umbrella term “seagrass”. However, we only used the occurrences from their marine coastal populations, excluding occurrence records from inland hypersaline lakes.

**Table 1.**
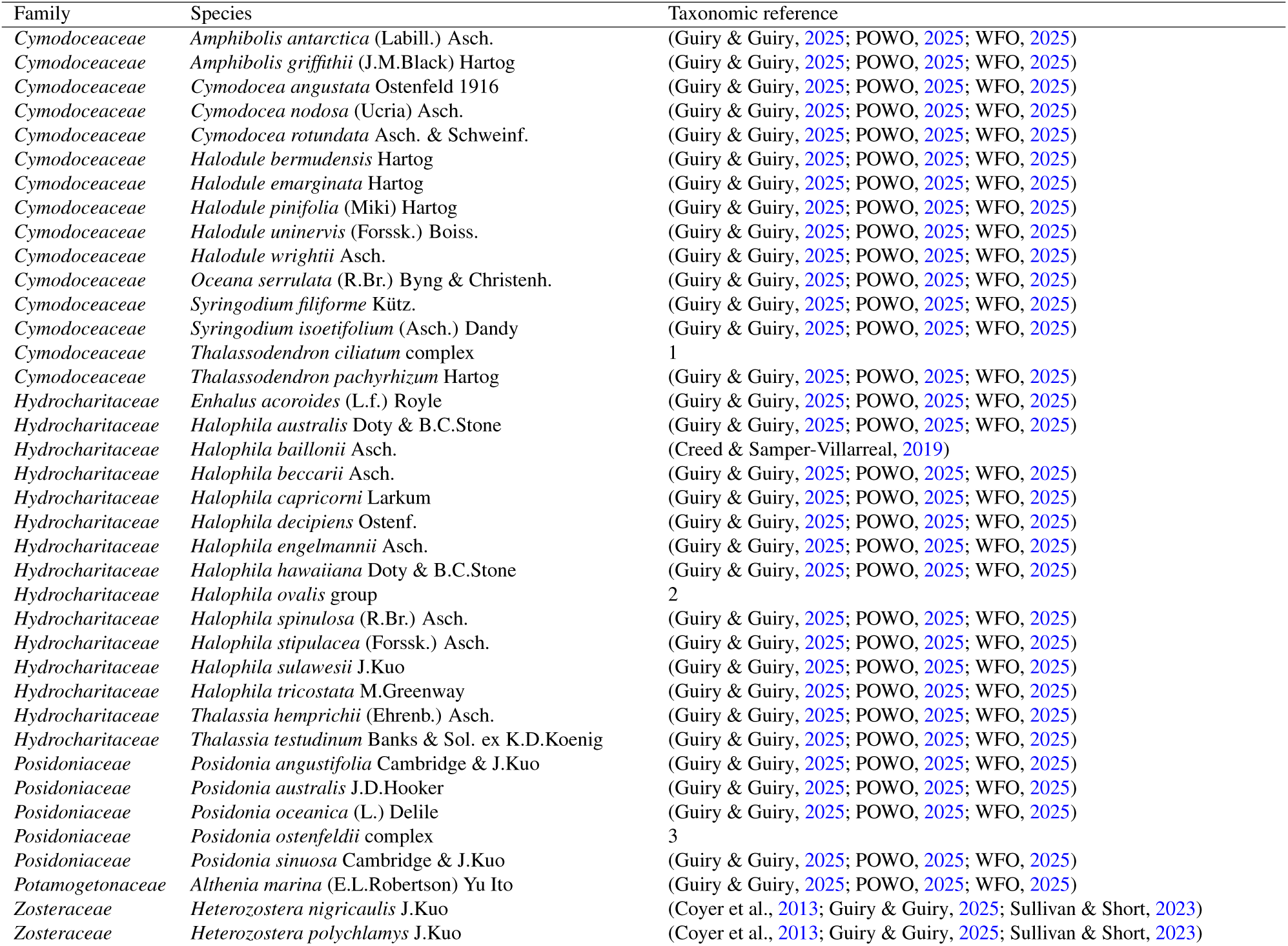
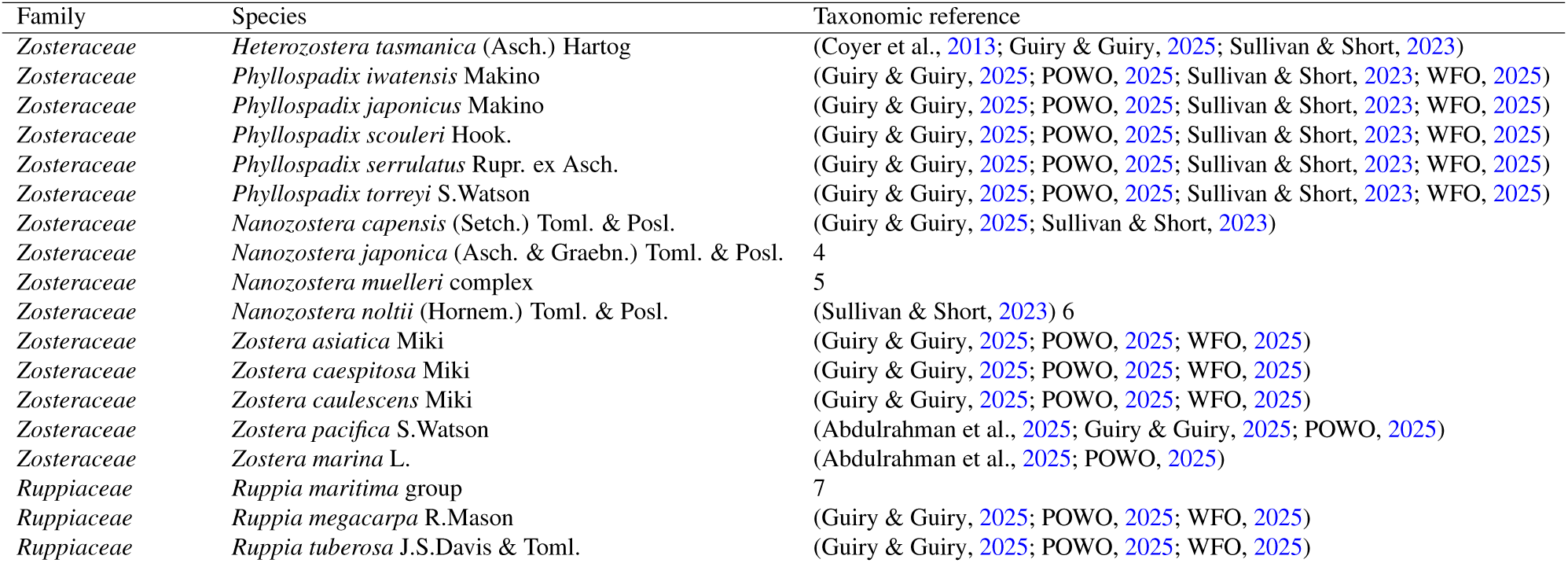
Curated list of seagrass taxa considered, including the taxonomic references that validate their nomenclature and taxonomic status. Comments expand discrepancies across references.

1. The following species are included in the complex: *T. ciliatum* (Forssk.) Hartog and *T. leptocaule* Maria C.Duarte, Bandeira & Romeiras. We grouped them into a complex as sampling efforts prior to the recent description of *T. leptocaule* may have misidentified it for *T. ciliatum*, and its data availability is still very low. However, given the difference in their range sizes, *T. ciliatum* dominates this complex.
2. Our *H. ovalis* group included *H. bullosa* (Setchell) J.Kuo, *H. gaudichaudii* J.Kuo, *H. linearis* Hartog, *H. major* (Zollinger) Miquel, *H. mikii* J.Kuo, *H. minor* (Zoll.) Hartog, *H. nipponica* J.Kuo, *H. okinawensis* J.Kuo, H. ovalis (R.Br.) Hook.f., and *H. ramamurthiana* (K.Ravikumar & R.Ganesan) J.Kuo. We decided to group these species together because of (1) the high taxonomic instability of this section (Kuo, 2020), (2) the difficulties in classifying or identifying species morphologically, (3) the very rapidly changing nomenclature may have led to misidentifications, (4) the taxonomical instability has resulted in errors at classifying these organisms in major biodiversity databases, (5) reports of hybridization within this group (Liu et al., 2020), and (6) our aim towards an analysis that will have the highest classification validity in the near future.
3. *Posidonia ostenfeldii* complex includes *P. ostenfeldii* Hartog, *P. robertsoniae* J.Kuo & Cambridge, *P. denhartogii* J.Kuo & Cambridge, *P. coriacea* Cambridge & J.Kuo and *P. kirkmanii* J.Kuo & Cambridge following Campey et al. (2000) and Hartog and Kuo (2007).
4. The two unnamed cryptic species within *N. japonica* mentioned in Zhang et al. (2025).
5. Following Jacobs et al. (2006) and Sullivan and Short (2023), *N. muelleri* was considered as the currently valid name for *N. muelleri* (Setch.) Toml. & Posl., *N. capricorni* (Setch.) Toml. & Posl., *N. mucronata* (Hartog.) Toml. & Posl., and *N. novazelandica* (Setch.) Toml. & Posl.
6. *Nanozostera noltii* and the subdivisions of *Zostera* sensu lato. We use the *Nanozostera* (and *Heterozostera*) genera under the justification that Coyer et al. (2013) supported genetically the morphological split proposed by Tomlinson and Posluzny (2001), which has been further validated by recent revisions (Sullivan & Short, 2023). The epiphet “*noltii*” is used over “*noltei*” because the species was first described with the *Z. noltii* (Hornemann, 1882) spelling, following the latinised version of E.F. Nolte (Noltius). Thus, the spelling should be kept, per ICN Art. 60.8 and Rec. 60C.1.
7. den Hartog and Triest (2020) concluded that the following three species are separate taxa: *Ruppia maritima* L. *sensu stricto*, *Ruppia cirrhosa* (Petagna) Grande, and *Ruppia spiralis* L. ex Dumort. However, we grouped them within this complex (which was rejected by den Hartog and Triest (2020)) under the justification that many misidentifications could have taken place as a result of the highly variable classification of these *Ruppia* species.

## S4: DATASET DESCRIPTION

A total of 109,150 seagrass records across 54 taxa were compiled in the main dataset used in this study. Fig. 1 shows the raw species counts, after cleaning and deduplicating but prior to any thinning. The Fig. 2 map shows the gridded distribution and record coverage for all species, by overlaying the rasterised log-transformed frequencies on the biogeographical realms proposed in Spalding et al., (2007).

**Figure 1.**
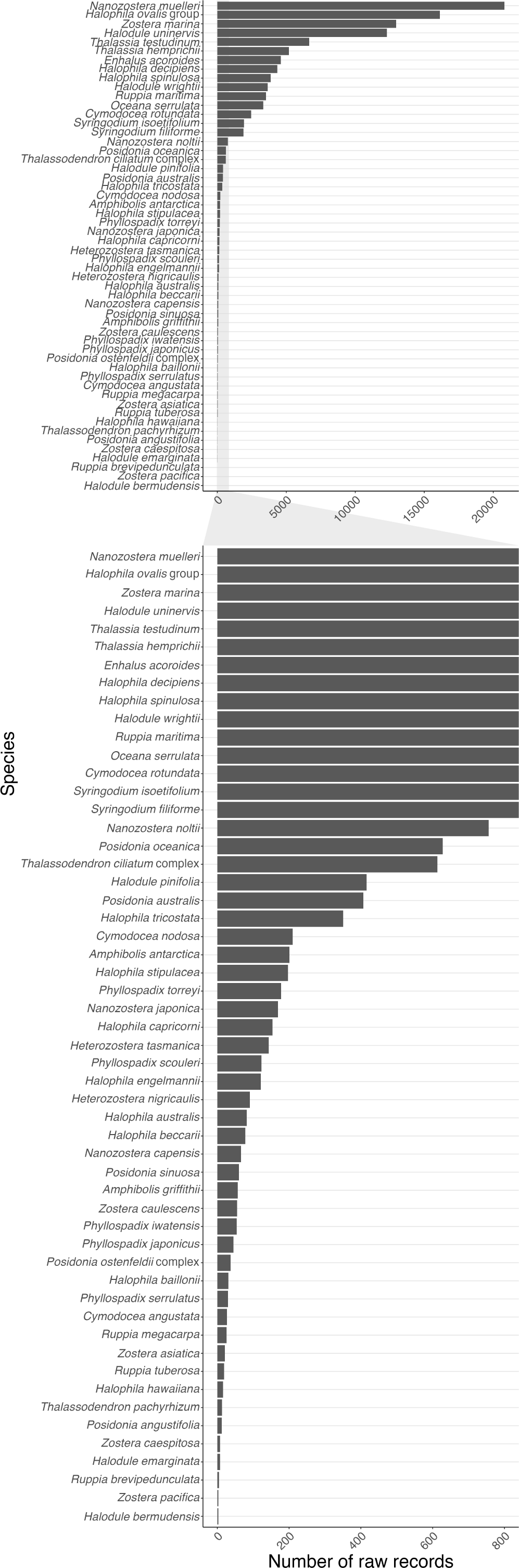
Number of raw records per species. Given the extreme unevenness between record numbers the lower part of the graph expands the x-axis between 0 to 800 records.

**Figure 2.**
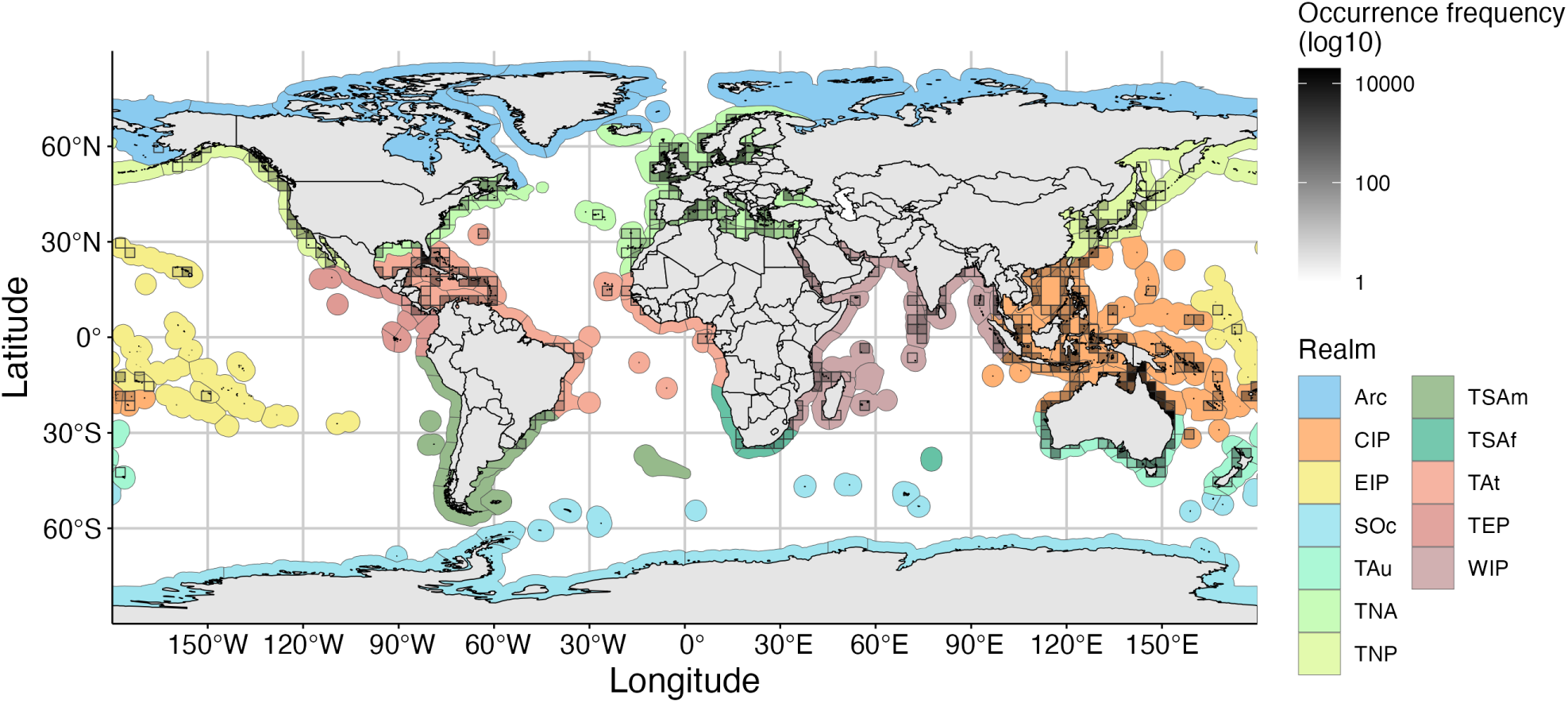
Niche stabilization plots for two seagrasses of different range sizes: (A) *Thalassia hemprichii*, broad ranged; and (B) *Posidonia angustifolia*, small ranged. The vertical dash line represents the stability threshold (ST), if reached.

## S5: SUPPLEMENTARY RESULT TABLES

### S5.1. Stabilization results

The effect of different thinning parameters [0, 17.5] on the thermal stabilization of seagrass species was measured with a stabilization threshold for each species and parameter pair (*ST_s,θ_*). The *ST_s,θ_* was defined as the minimum number of observations required to obtain a stable estimate of the thermal optimum. Table 2 shows the *ST_s,θ_* values for all (*s, θ*) pairs.

**Table 2.**
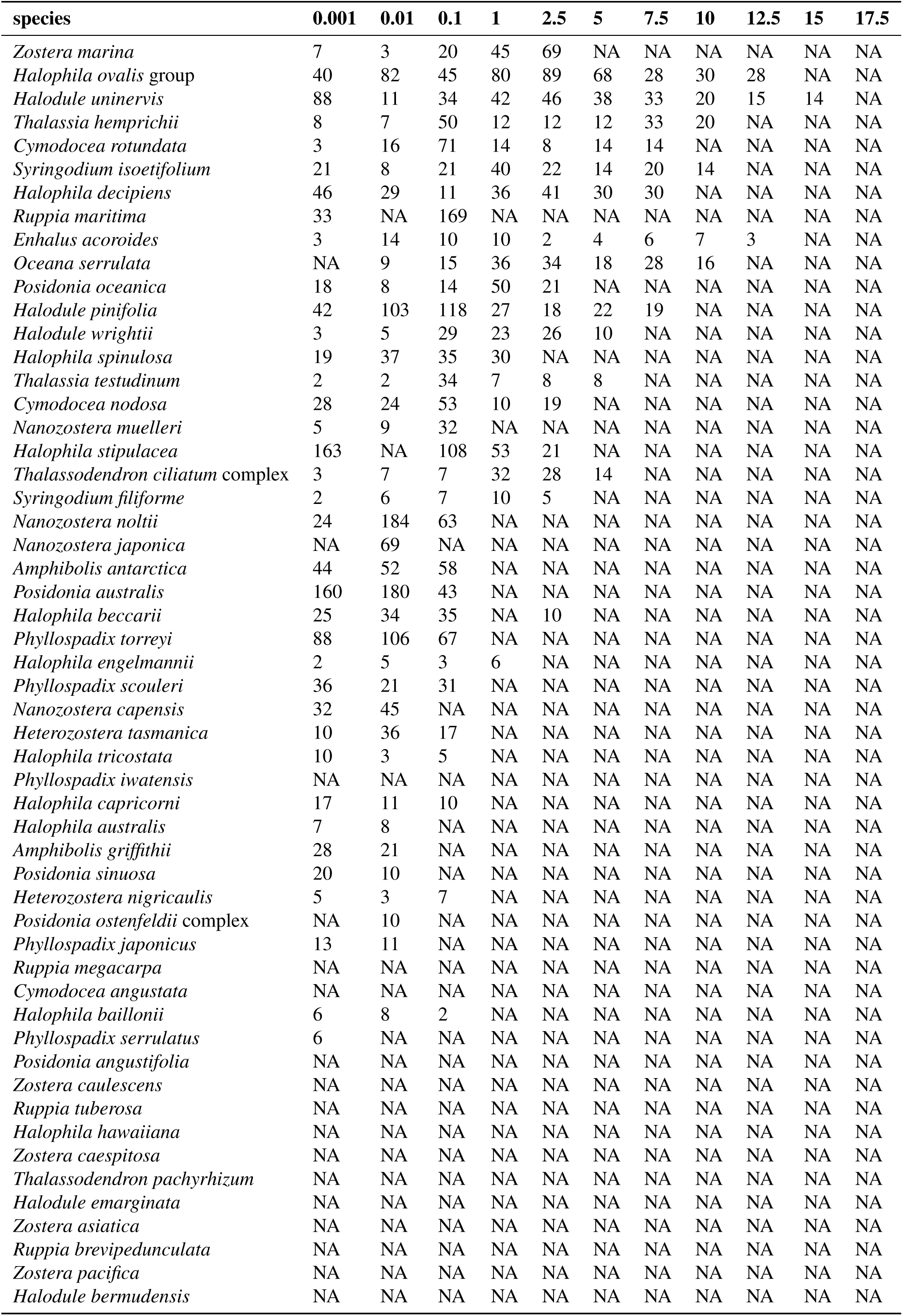
Stabilization threshold (ST) values for each species and parameter pairs. Species are shown as rows and are ordered by decreasing range size. Thinning parameters (measured in latitudinal degrees) are shown as columns. The ST values represent the random observation number at which the thermal curve becomes stable. When stabilization was not reached ST = NA is given.

### S5.2. Main results of our proposed methods

Each species was assigned its range size descriptor (*D_s_*) and thinning parameters predicted by BD-TP and LS-TP in Table 3. The difference between the prediction by both methods was quantified as the absolute difference between both predicted parameters (’*|θ_BD_ −θ_LS_|*’). Predicted parameters of 0.001° are the result of capping at this minimum conceptual parameter. Differences between methods are organised and plot in Fig. 3.

**Table 3.**
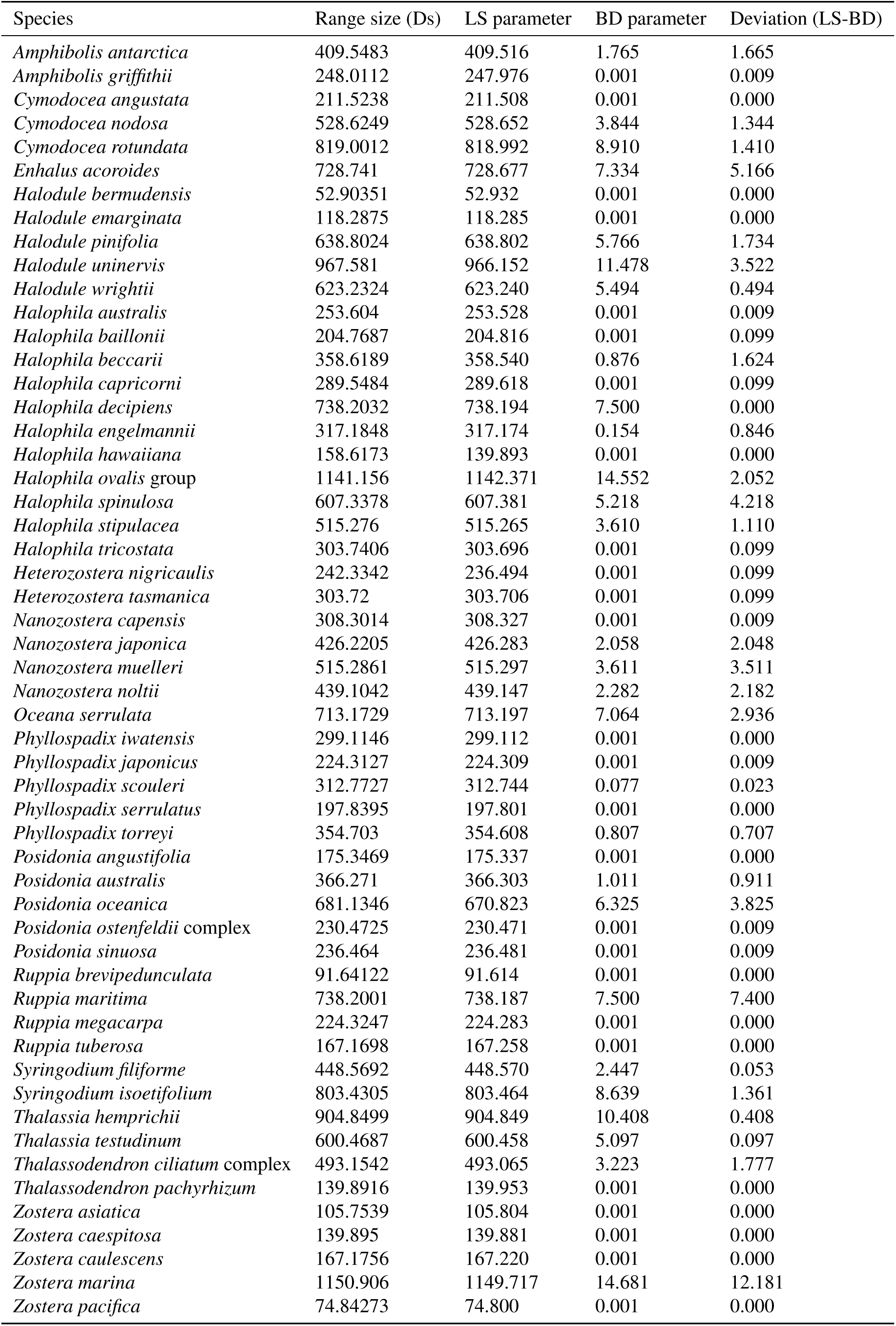
Range size and predicted thinning parameters by seagrass species. Deviation between both methods is shown as the absolute difference between both predicted parameters.

**Figure 3.**
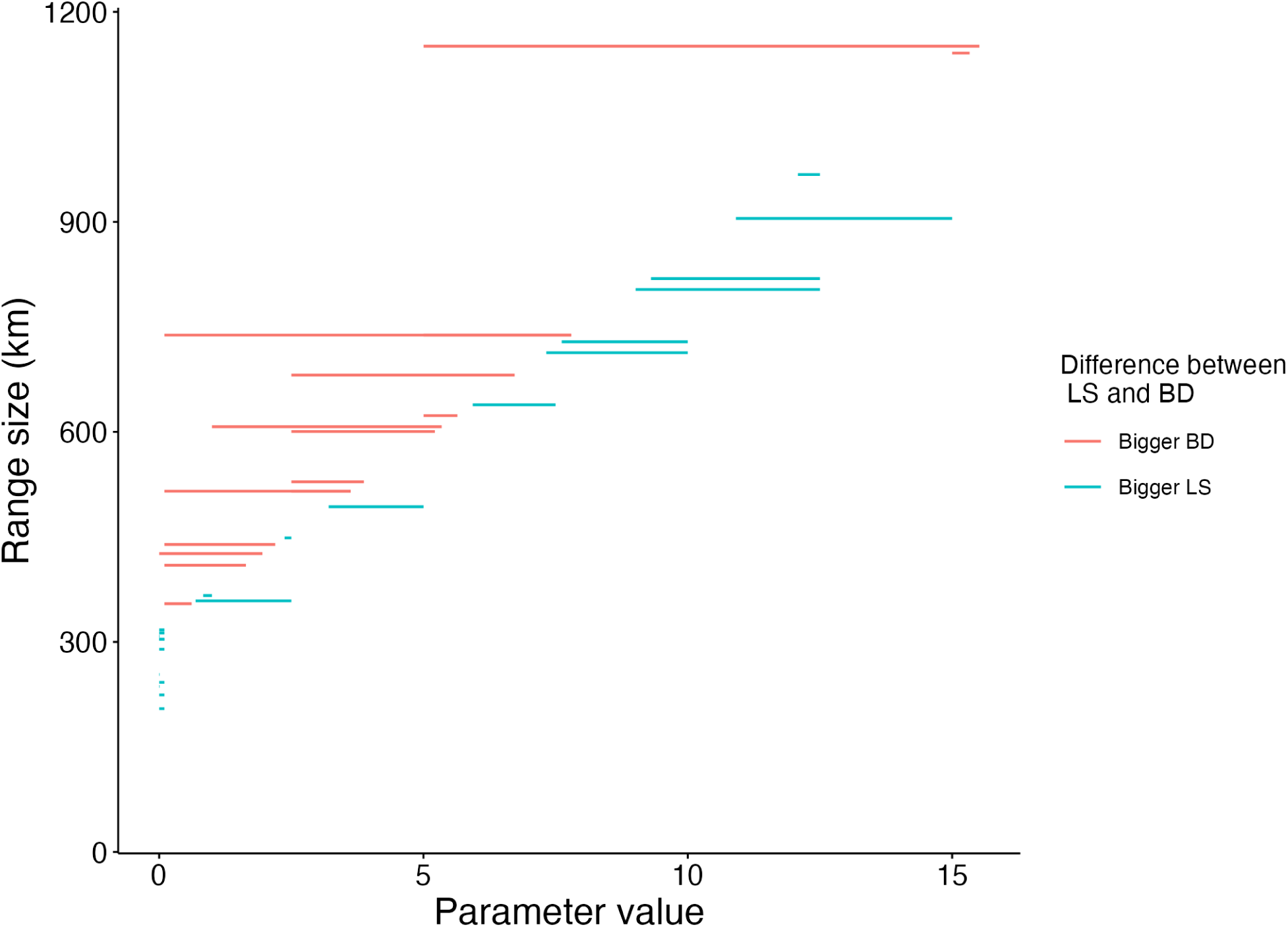
Differences in the parameter predictions by the BD-TP and LS-TP methods. Blue lines identify cases where the LS estimate is bigger than the BD estimate. Red lines identify cases where the BD estimate is bigger than the LS estimate. Barely visible lines show little to no difference between methods.

### S5.3. Number of counts pre- and post-thinning

For each species (ordered by *D_s_*), the original number of counts and the number of points post-thinning, after using BD-TP and LS-TP, is shown in Table 4. Only species deemed with sufficient data were considered.

**Table 4.**
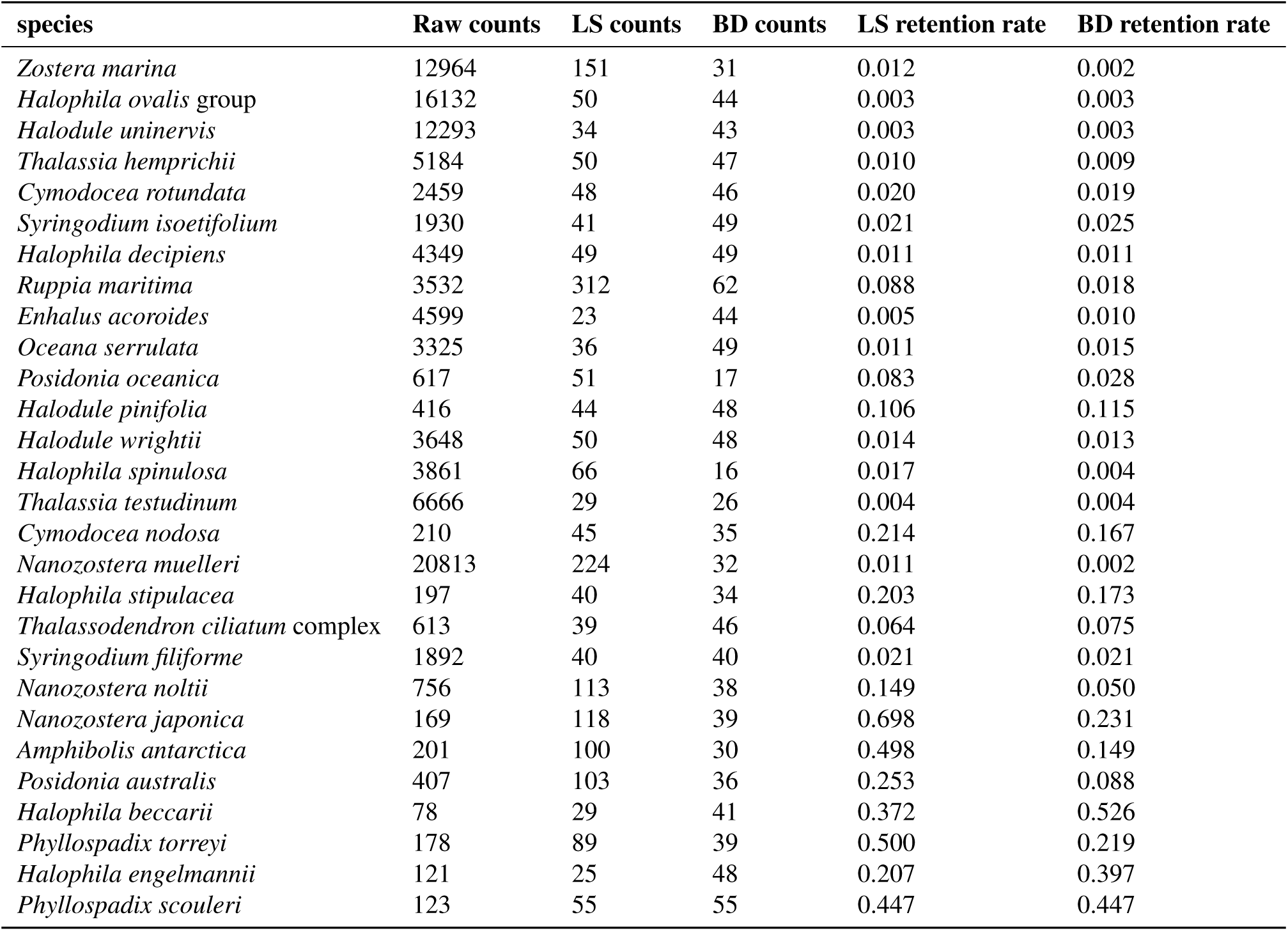
Species occurrence counts before and after the application of thinning with LS-TP and BD-TP parameterization. Only species that reach stability were considered.

**Table 5.**
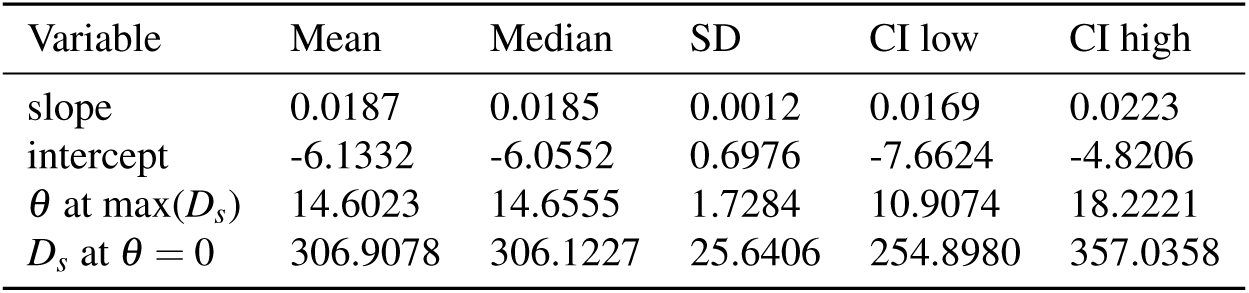
Summary table for cluster bootstrapping (i.e. per species)

**Table 6.**
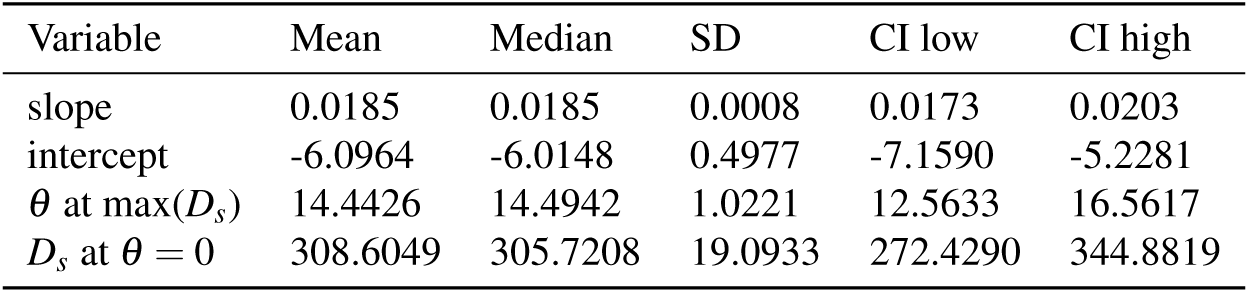
Summary table for row bootstrapping (i.e. per species x parameter)

### S5.4. Bootstrap uncertainty and leave-one-species-out cross-validation for BD-TP

We evaluated bootstrap uncertainty by resampling both at row and species level. The row bootstrap produced estimates of the slope, intercept, and derived BD-TP quantities under the assumption that individual observations are independent. Cluster (i.e. species) bootstrap resampled entire species, preserving dependence among observations of the same species. Cluster bootstrap and row bootstrap results are summarised in Table 5 and Table 6, respectively. The slope and intercept of the BD method formula were resolved analytically. The estimated parameter for the highest observed *D_s_* and *D_s_*at *θ* = 0 were calculated.

The original formula given in the main text

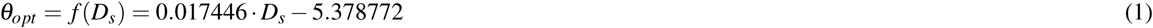

differs slightly from the formula calculated from row bootstrapping (considering the mean between runs)

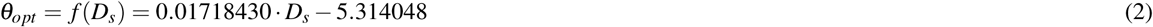

and cluster bootstraping (considering the mean between runs),

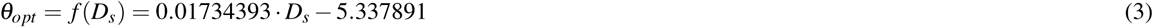

We further evaluated the linear SVM model with a Leave-One-Species-Out (LOSO) cross-validation scheme Table 7. For each iteration, all observations from one species were withheld as the test set, while the model was trained on the remaining species. Performance was summarised with true positives (TP), false positives (FP), true negatives (TN), and false negatives (FN), together with accuracy, precision, recall, and F1 score. Model accuracy typically fell for the species with higher *D_s_*, as they may have a higher influence in the shape of the hyperplane (e.g. *Zostera marina* AUC = 0.55, *Ruppia maritima* AUC = 0.55, or *Enhalus acoroides* AUC = 0.73), although no statistical correlation was found between *D_s_* and any of the evaluation metrics. LOSO results indicate heterogeneous predictive performance among species, with some species showing complete agreement between observed and predicted stability and others showing reduced discrimination, particularly through false-positive or false-negative classifications.

**Table 7.**
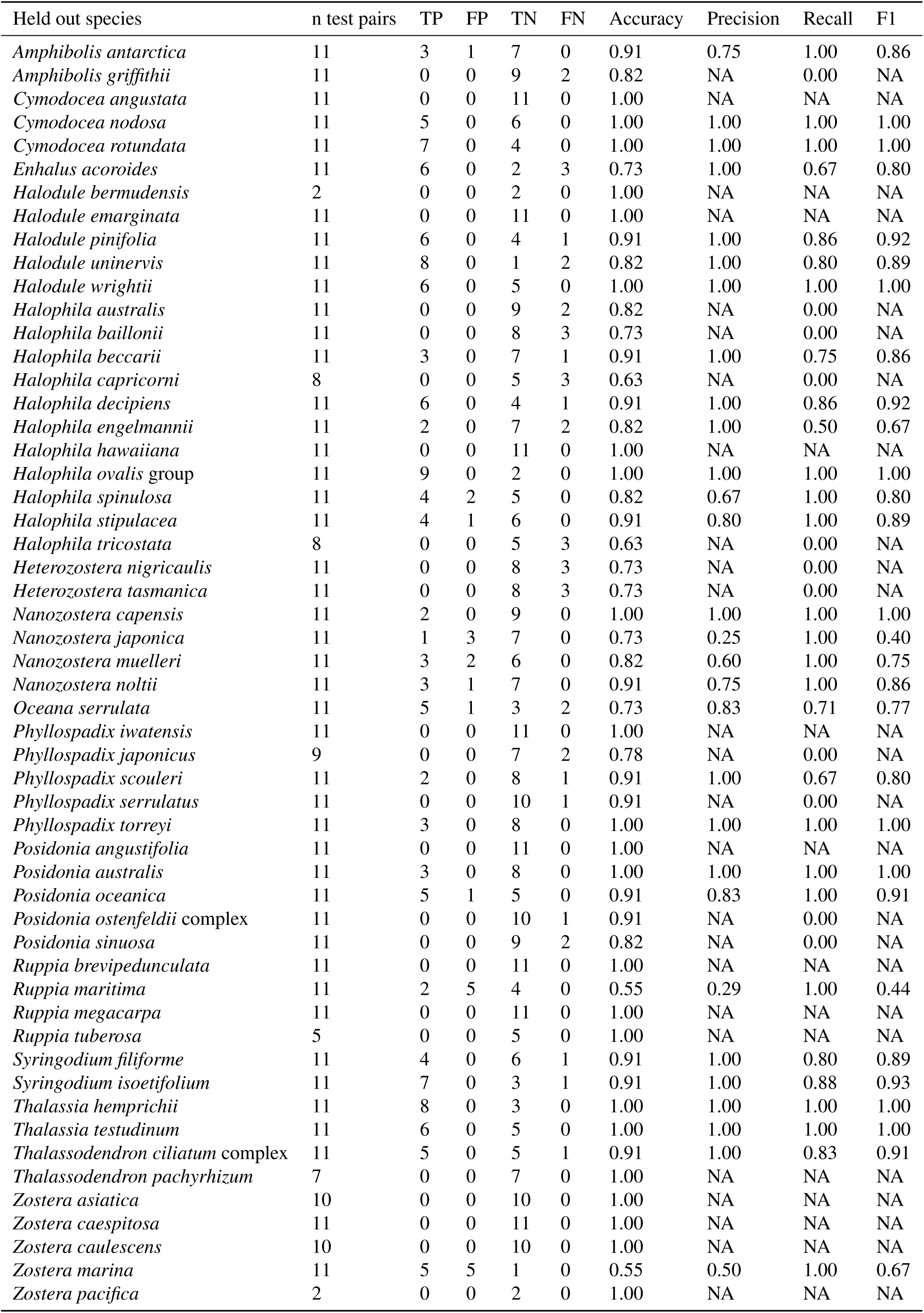
Evaluation metrics from the LOSO analysis. The species that was held out in each run is shown in the left.

## S6: STABILIZATION GRAPHS FOR ALL SPECIES

Stabilization plots of all species for all tested parameters, ordered by decreasing *D_s_*.

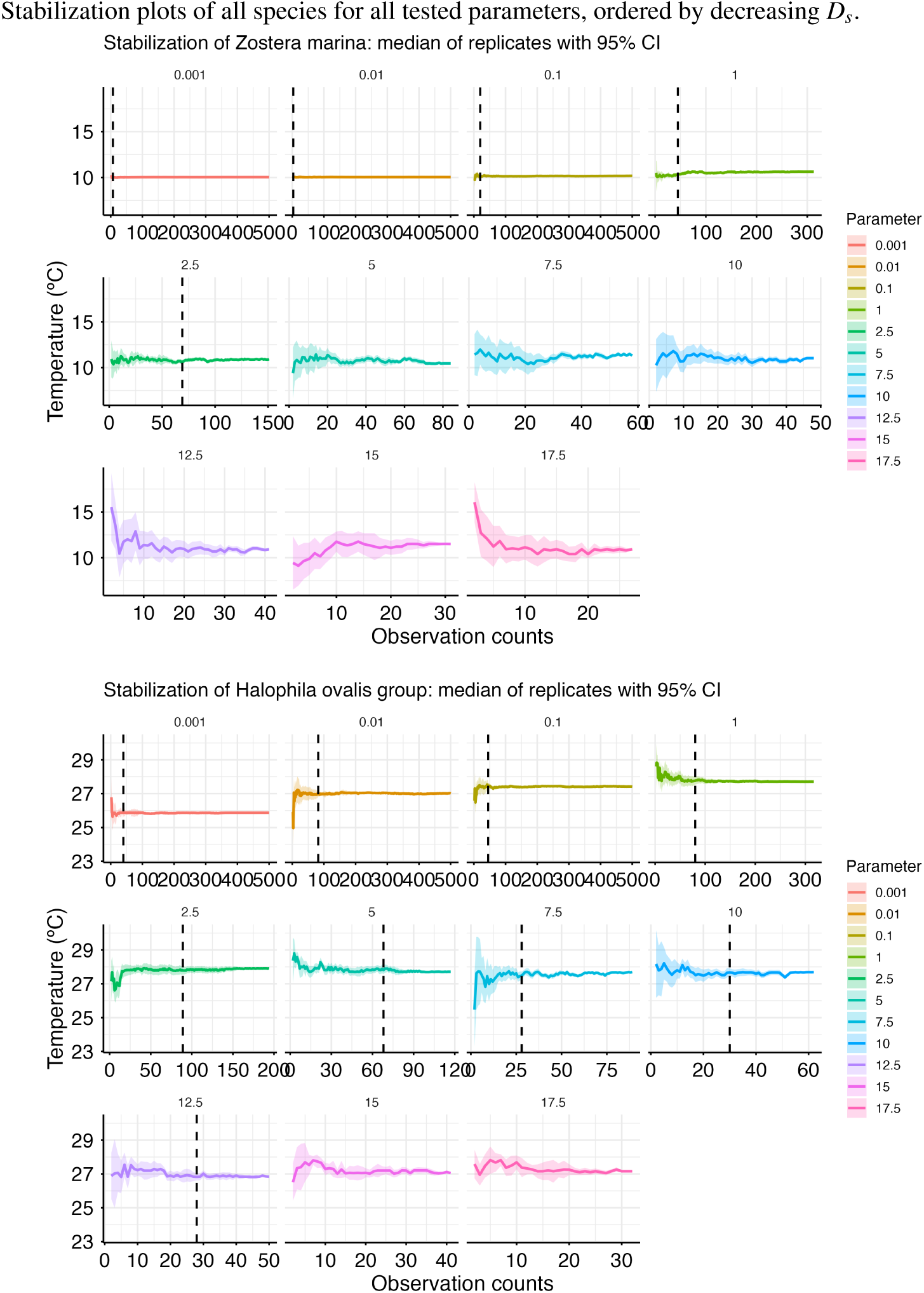
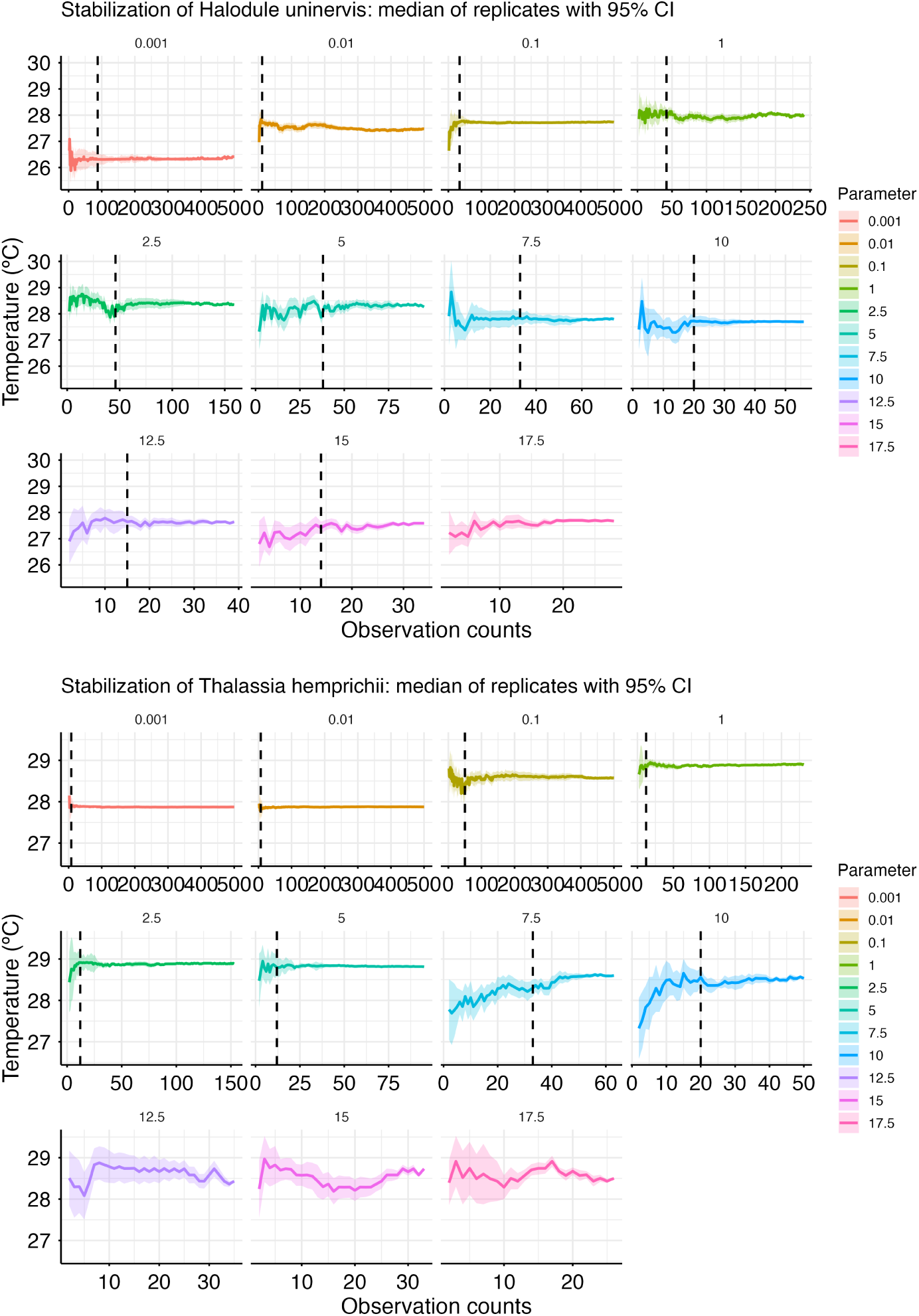
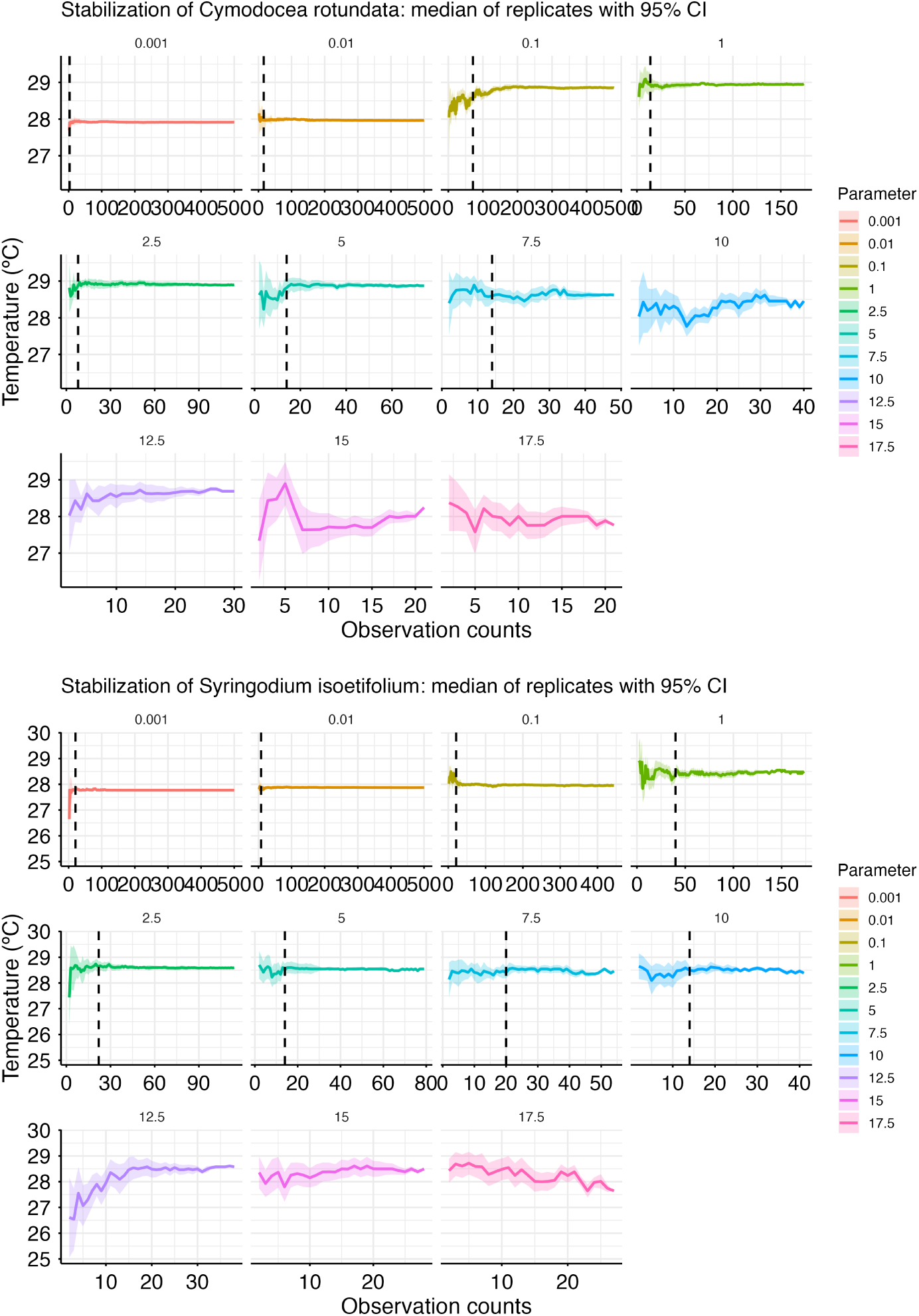
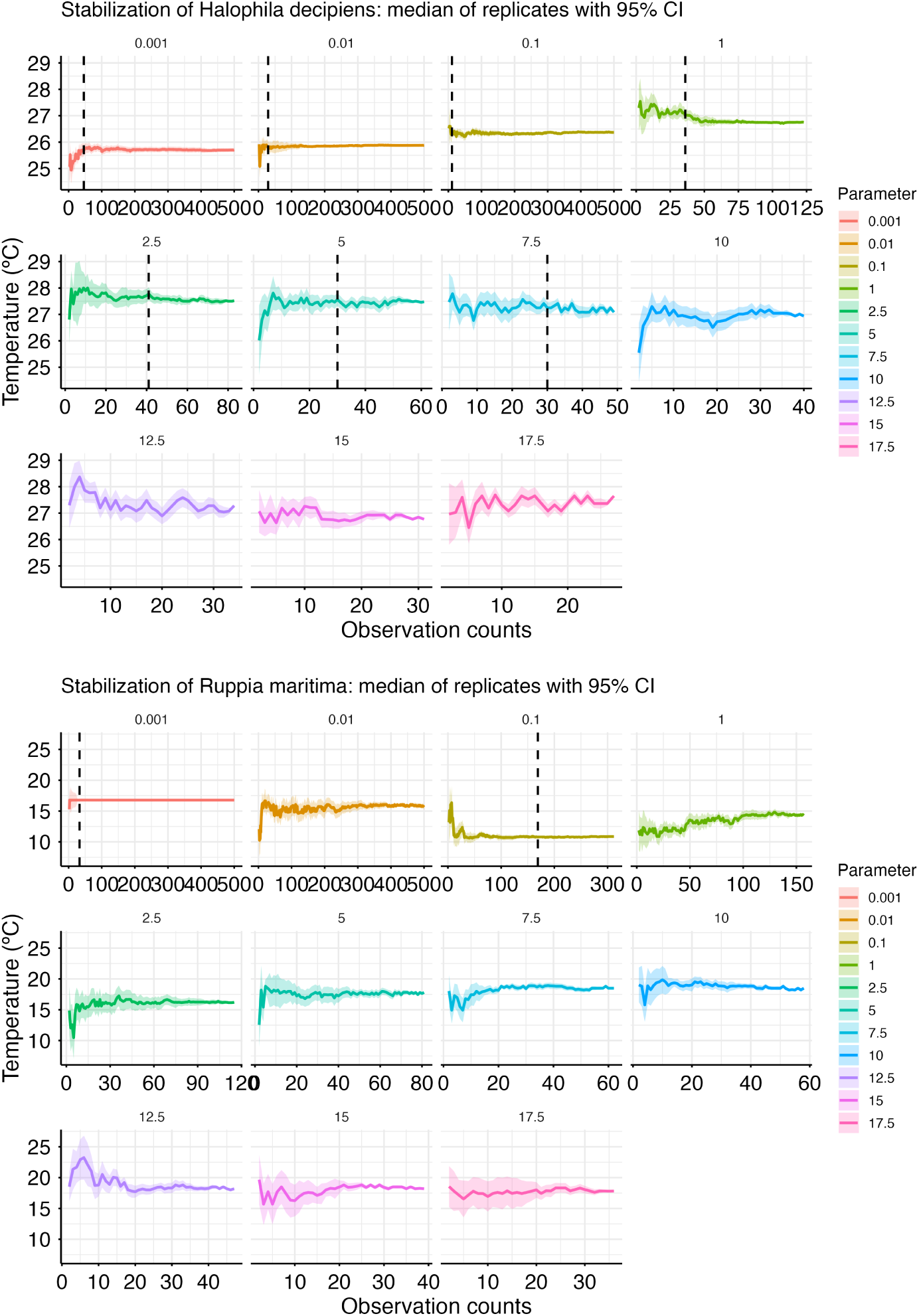
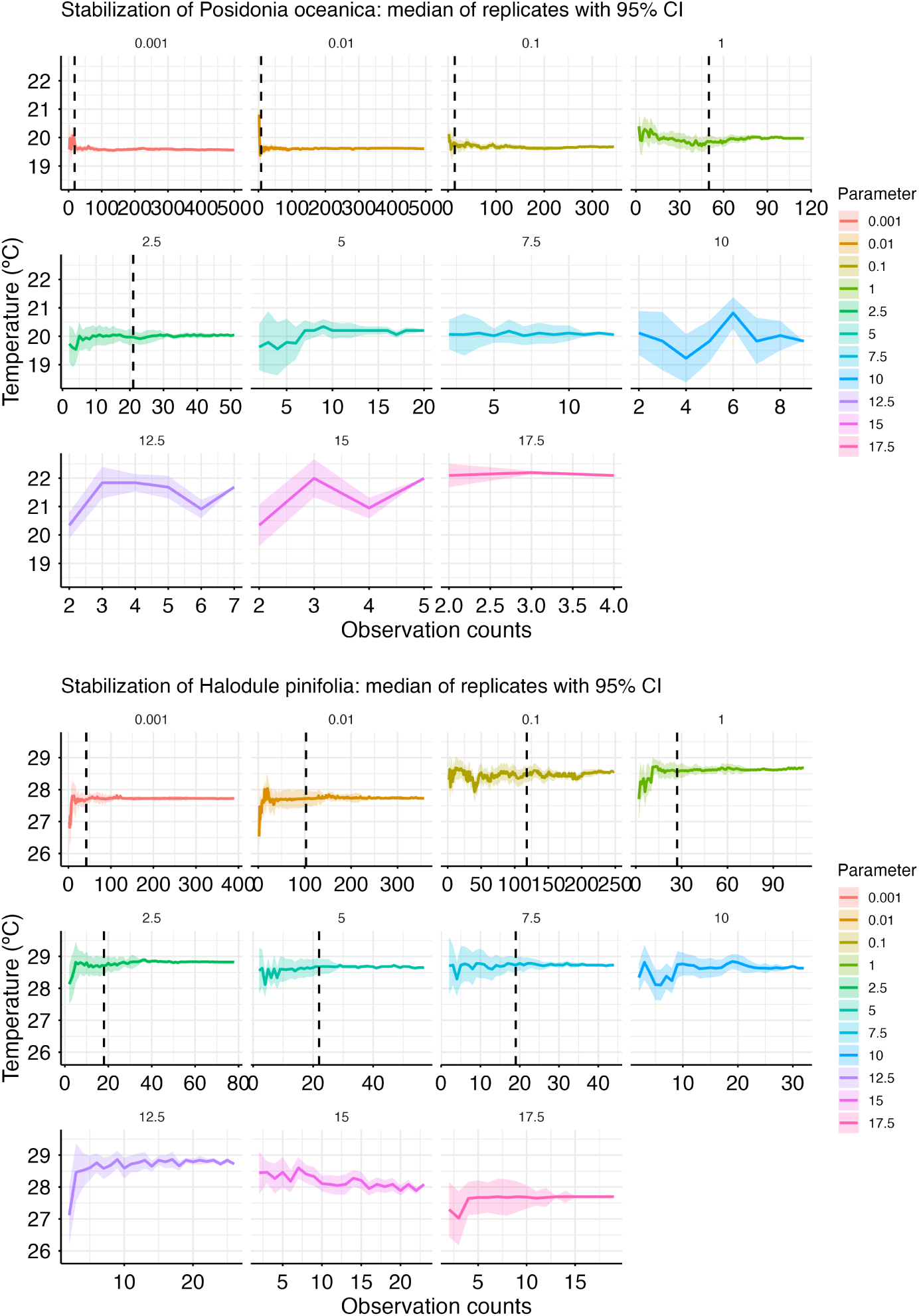
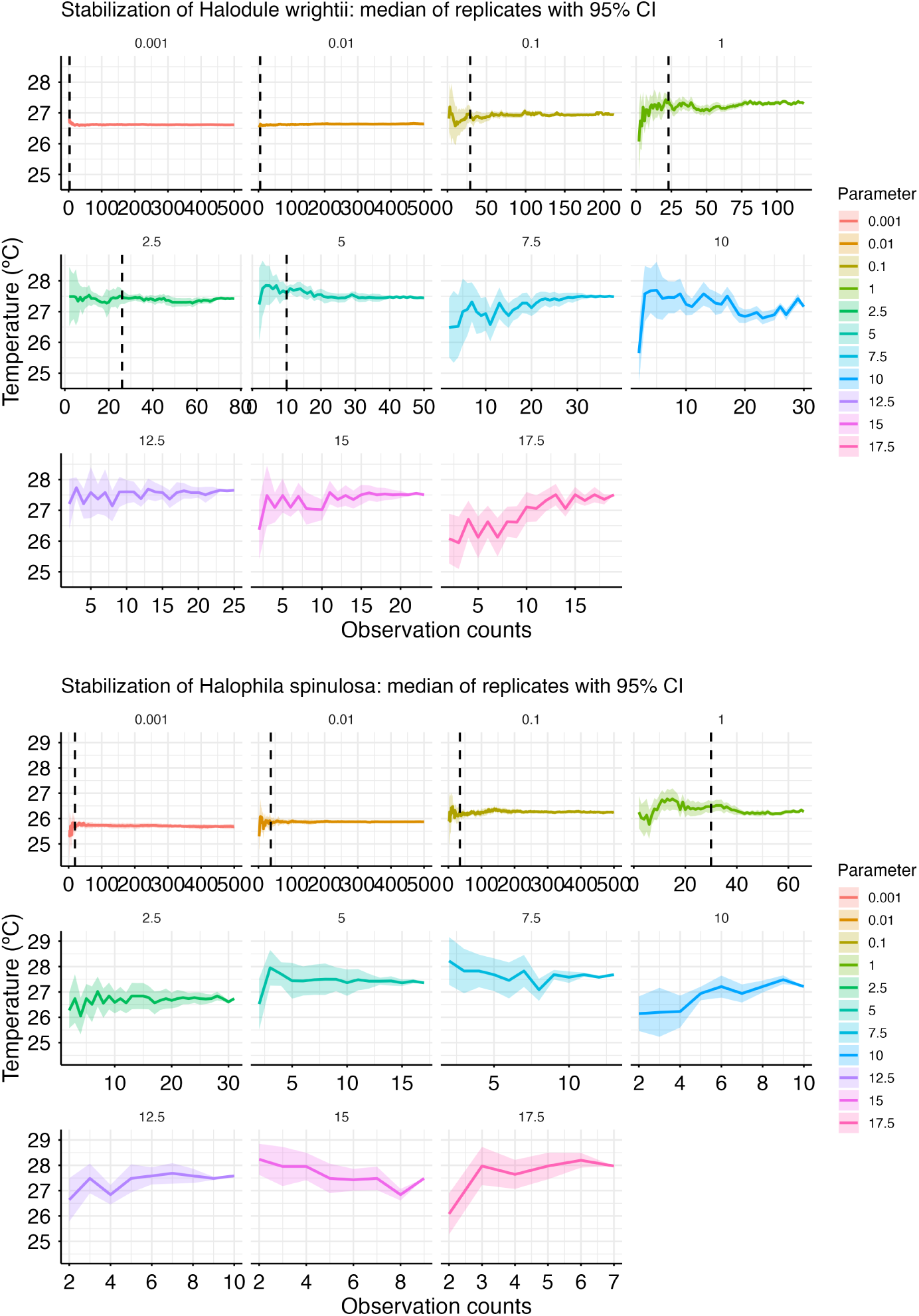
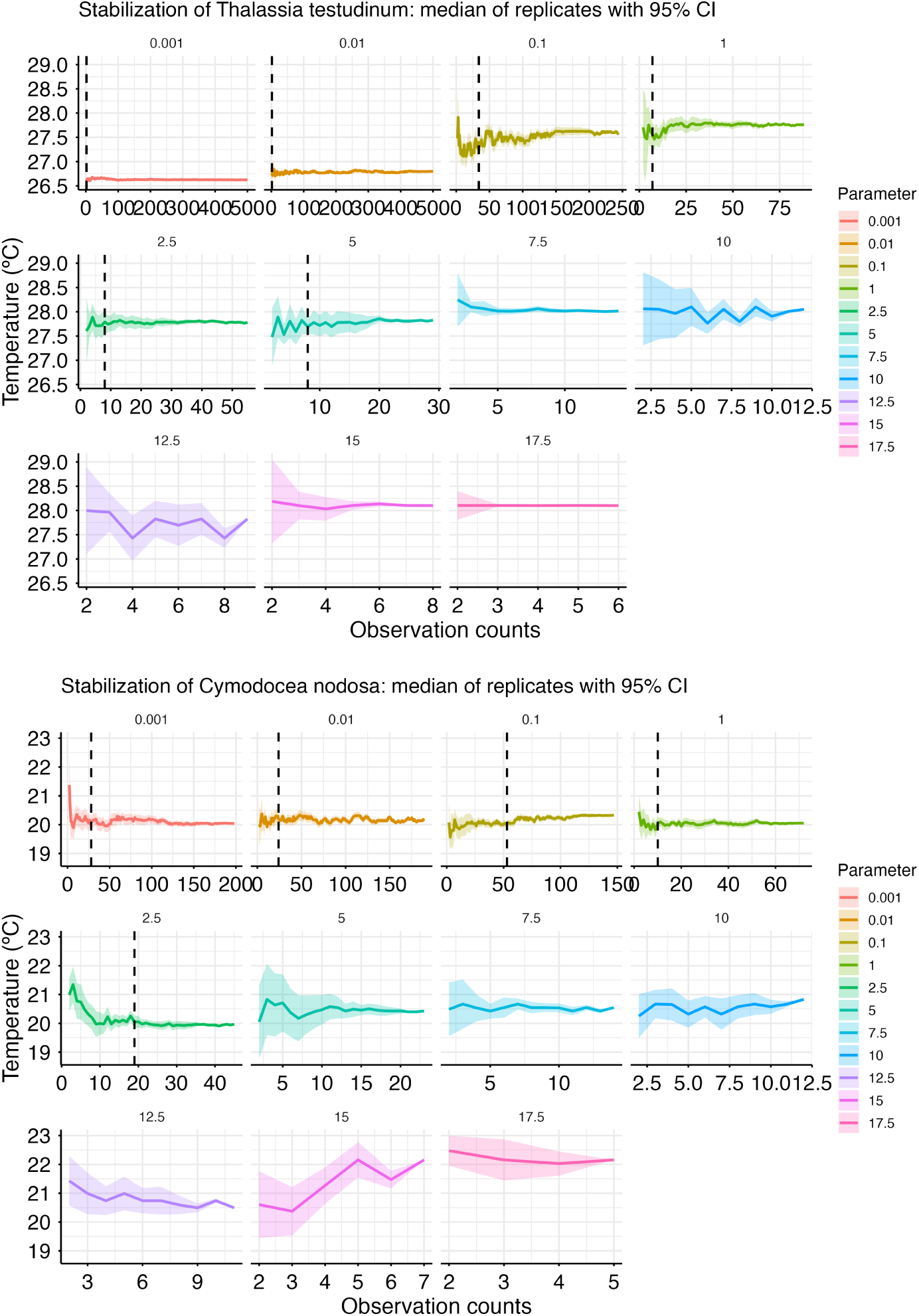
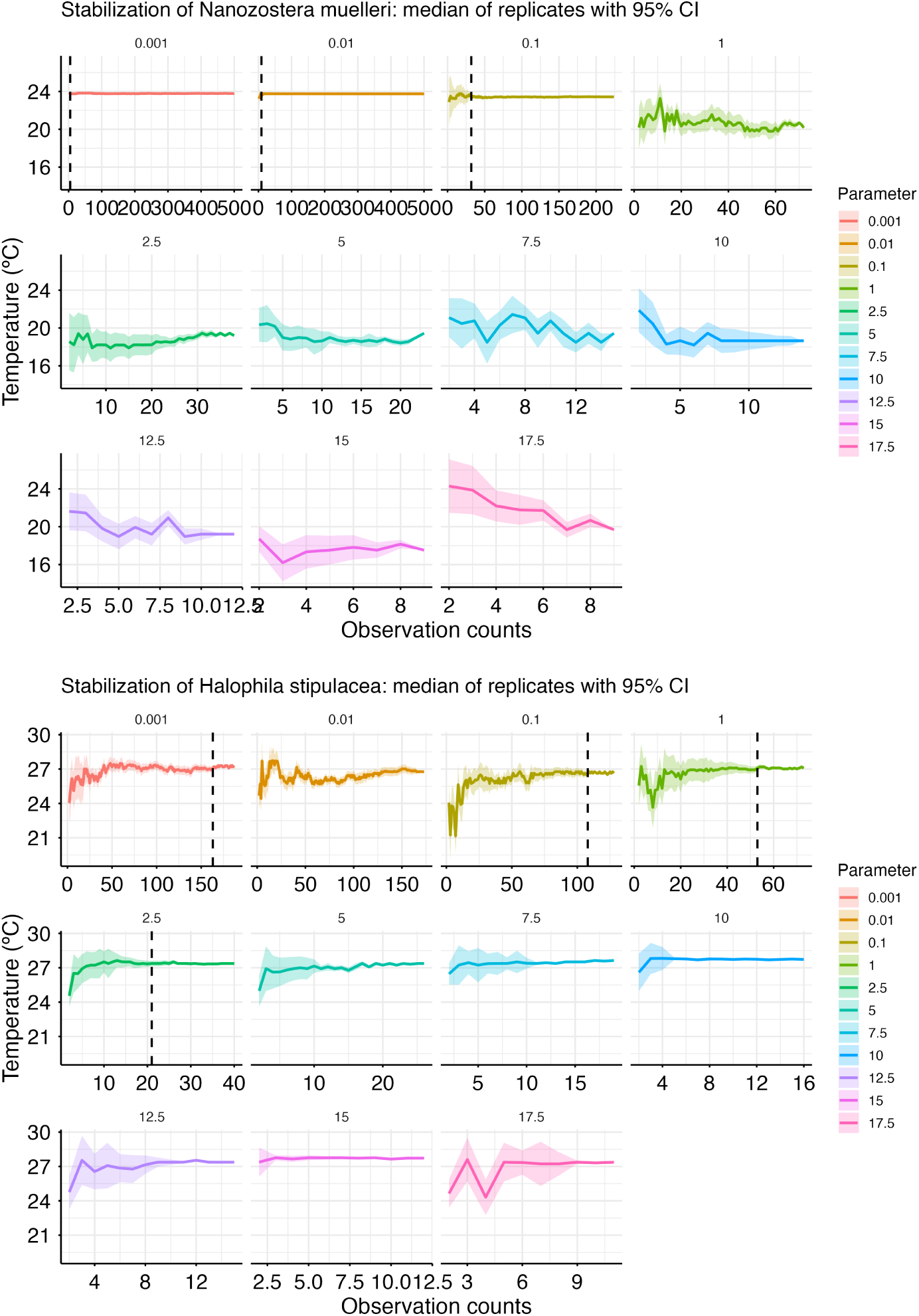
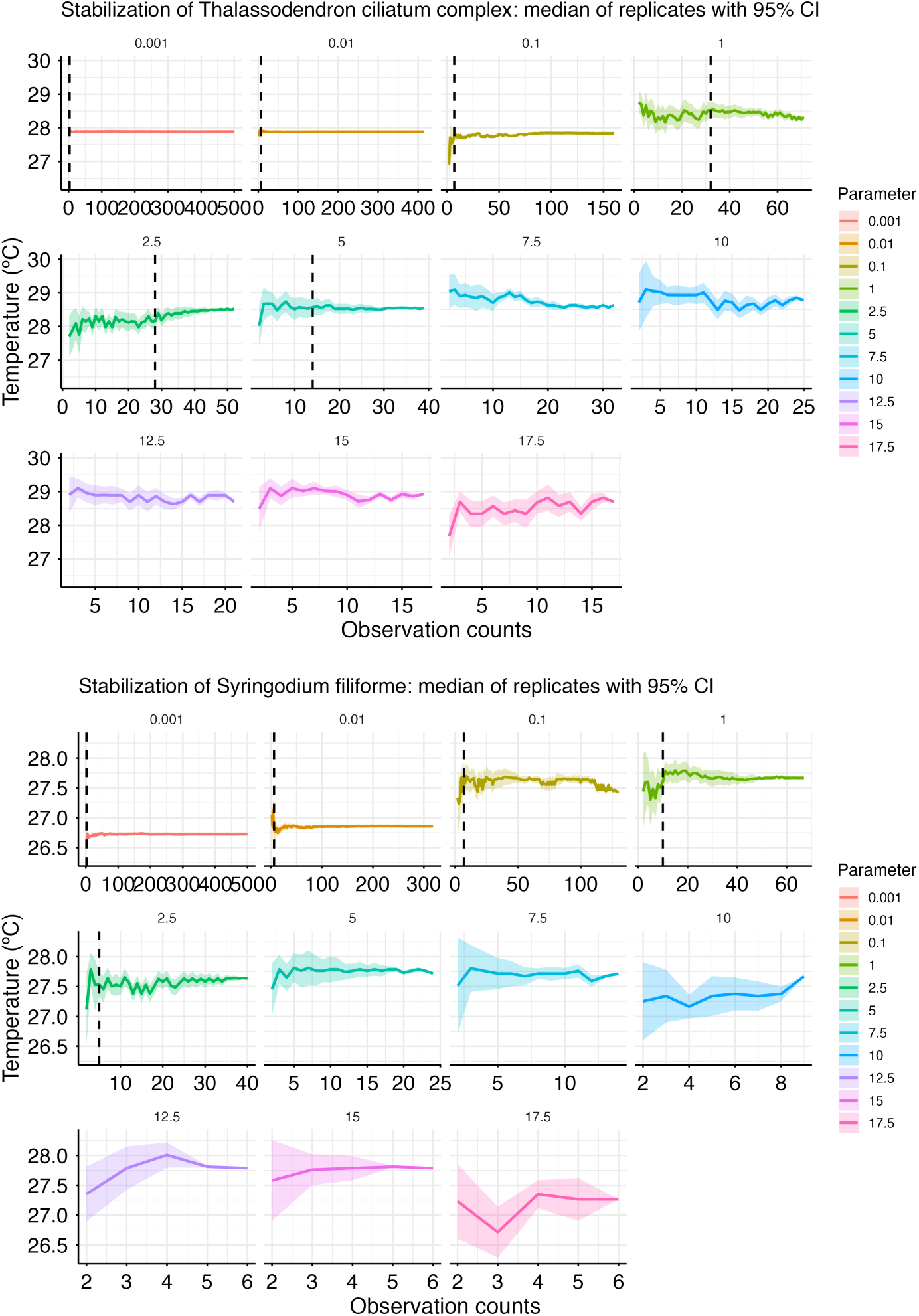
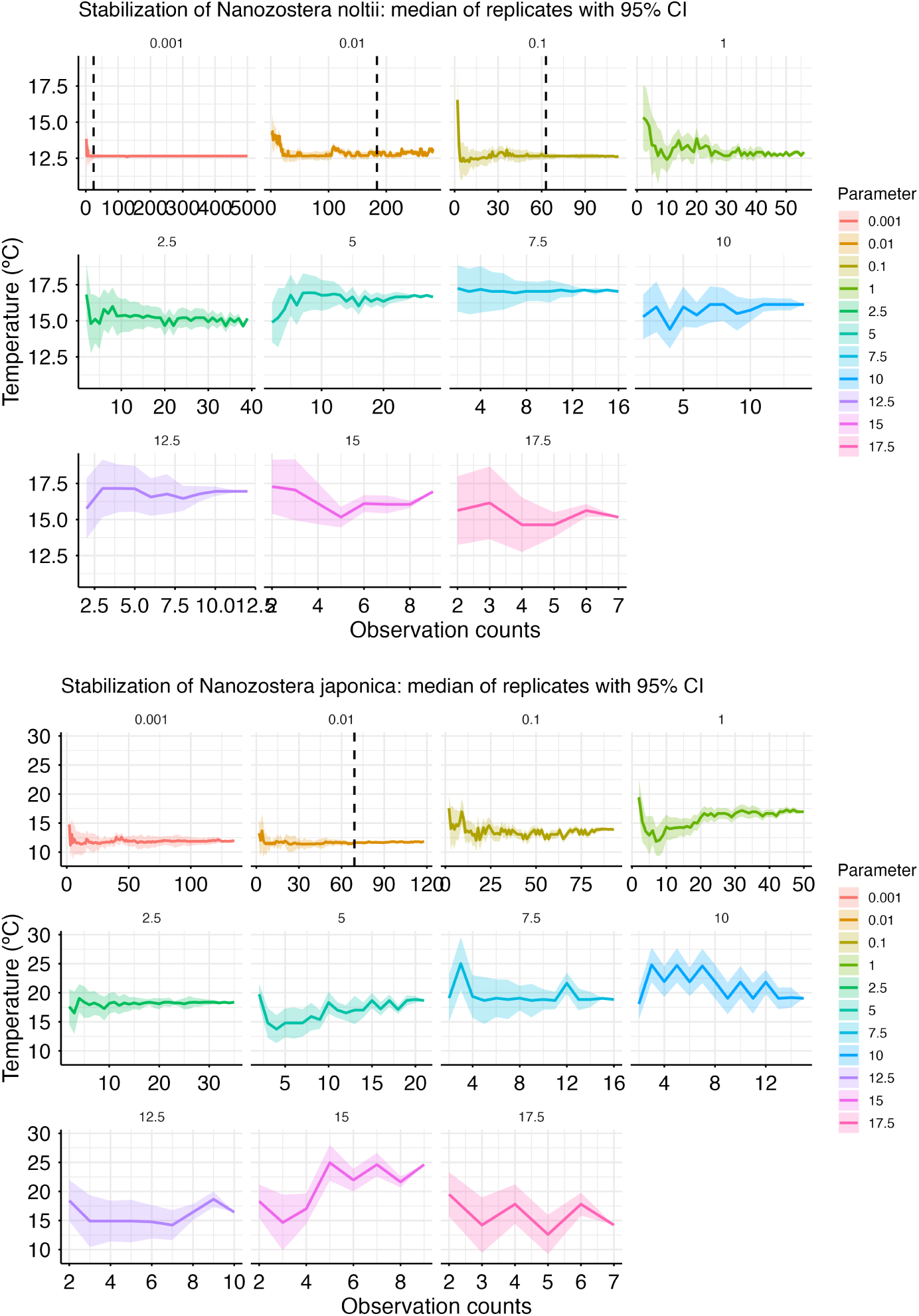
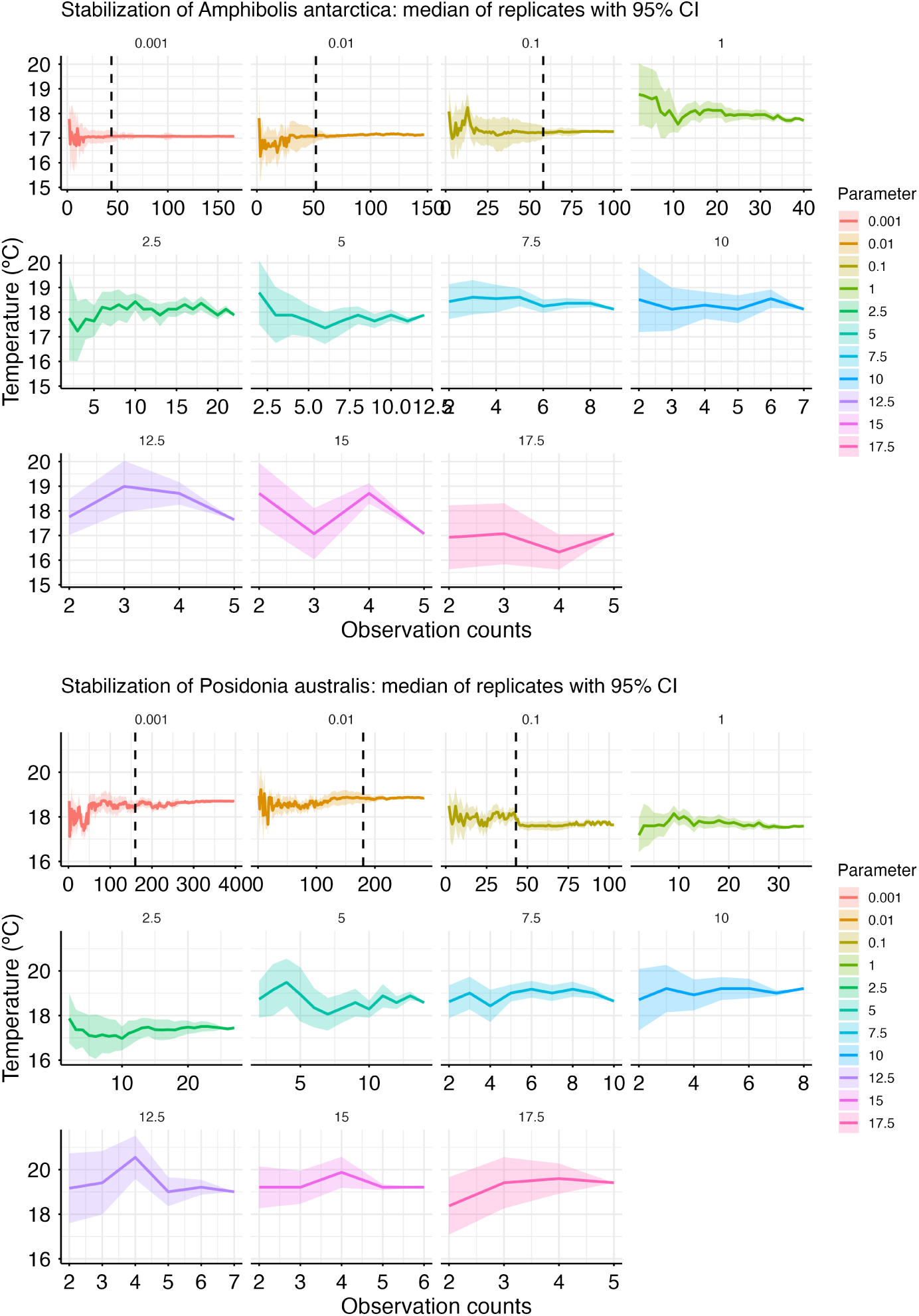
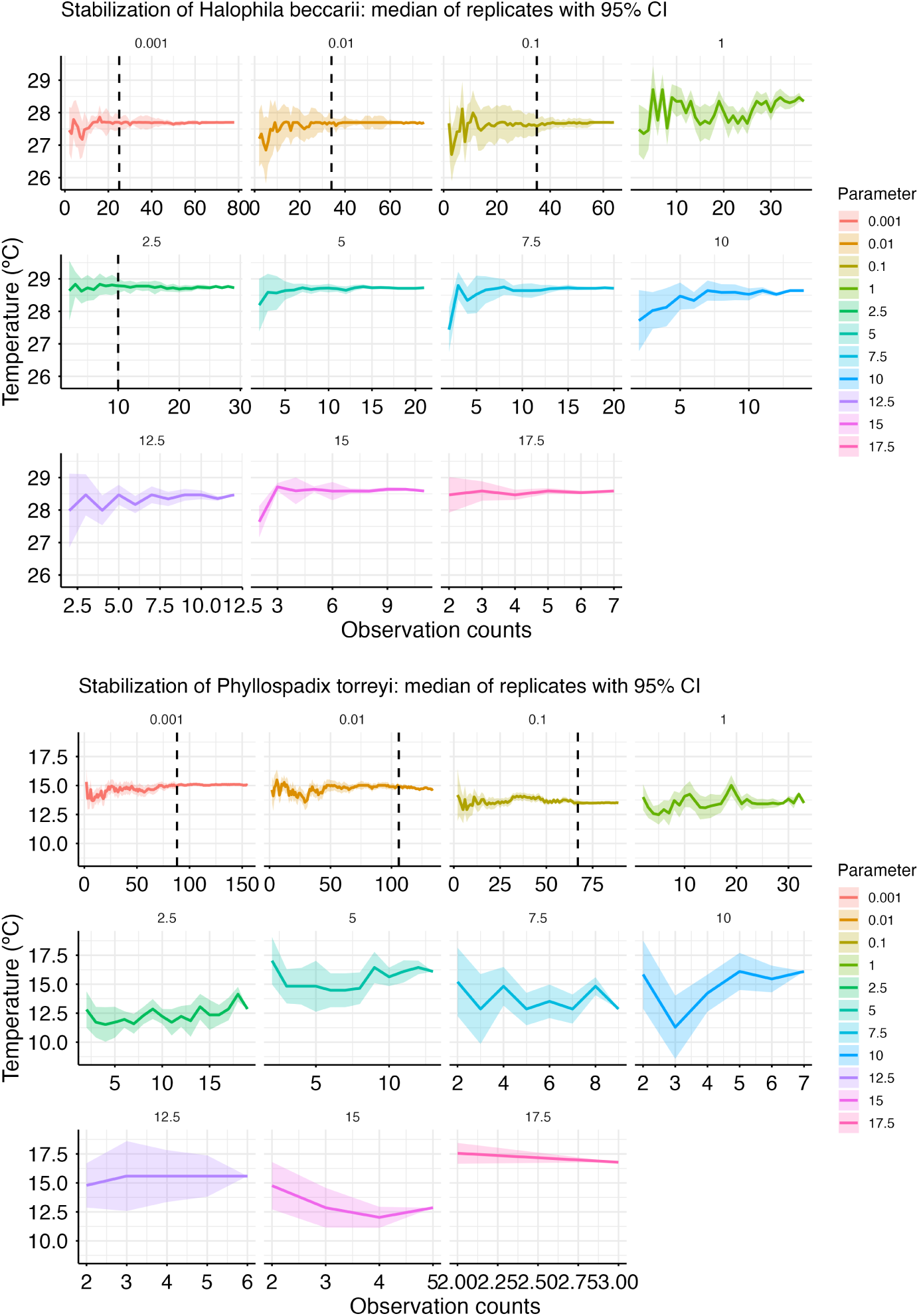
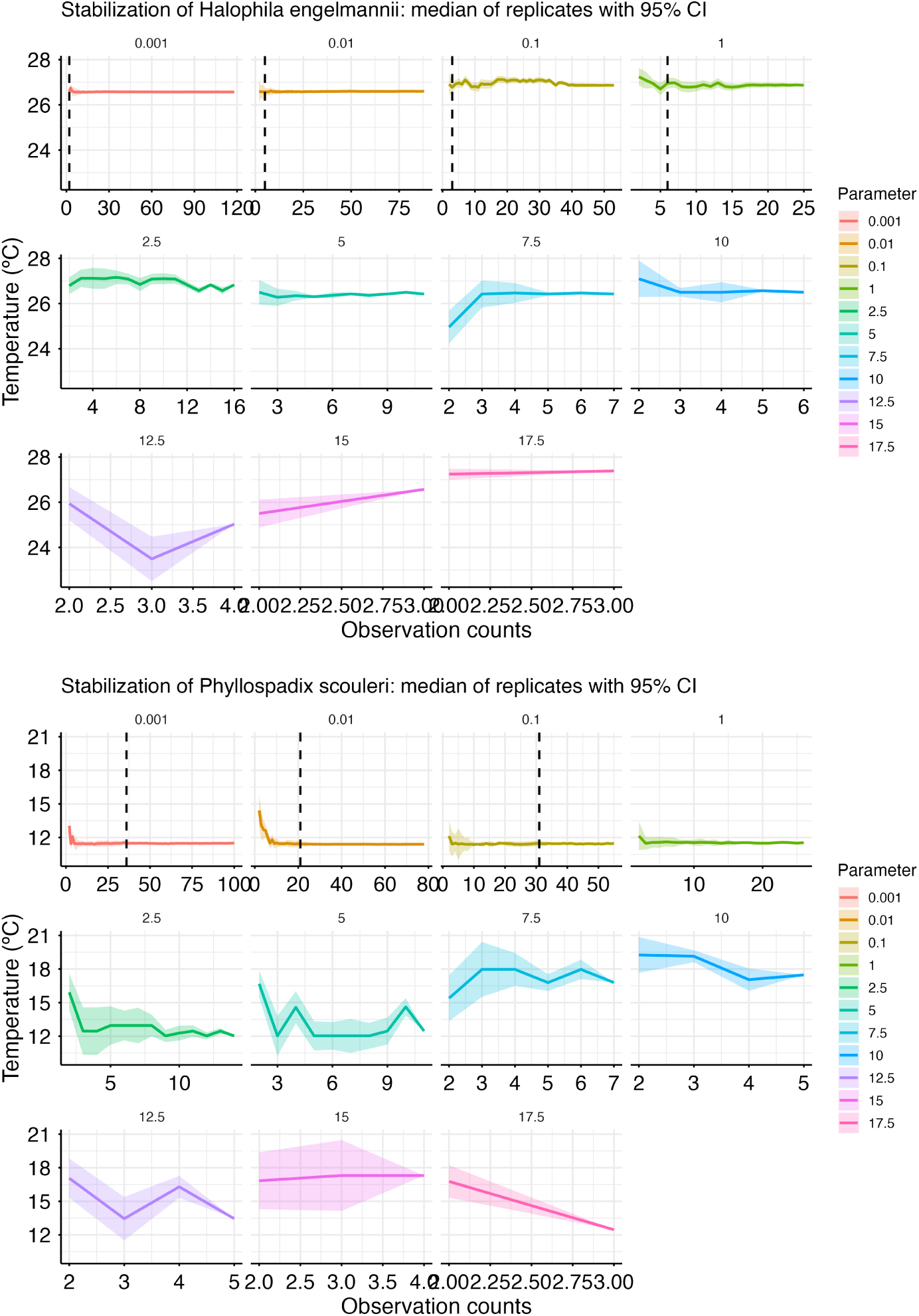
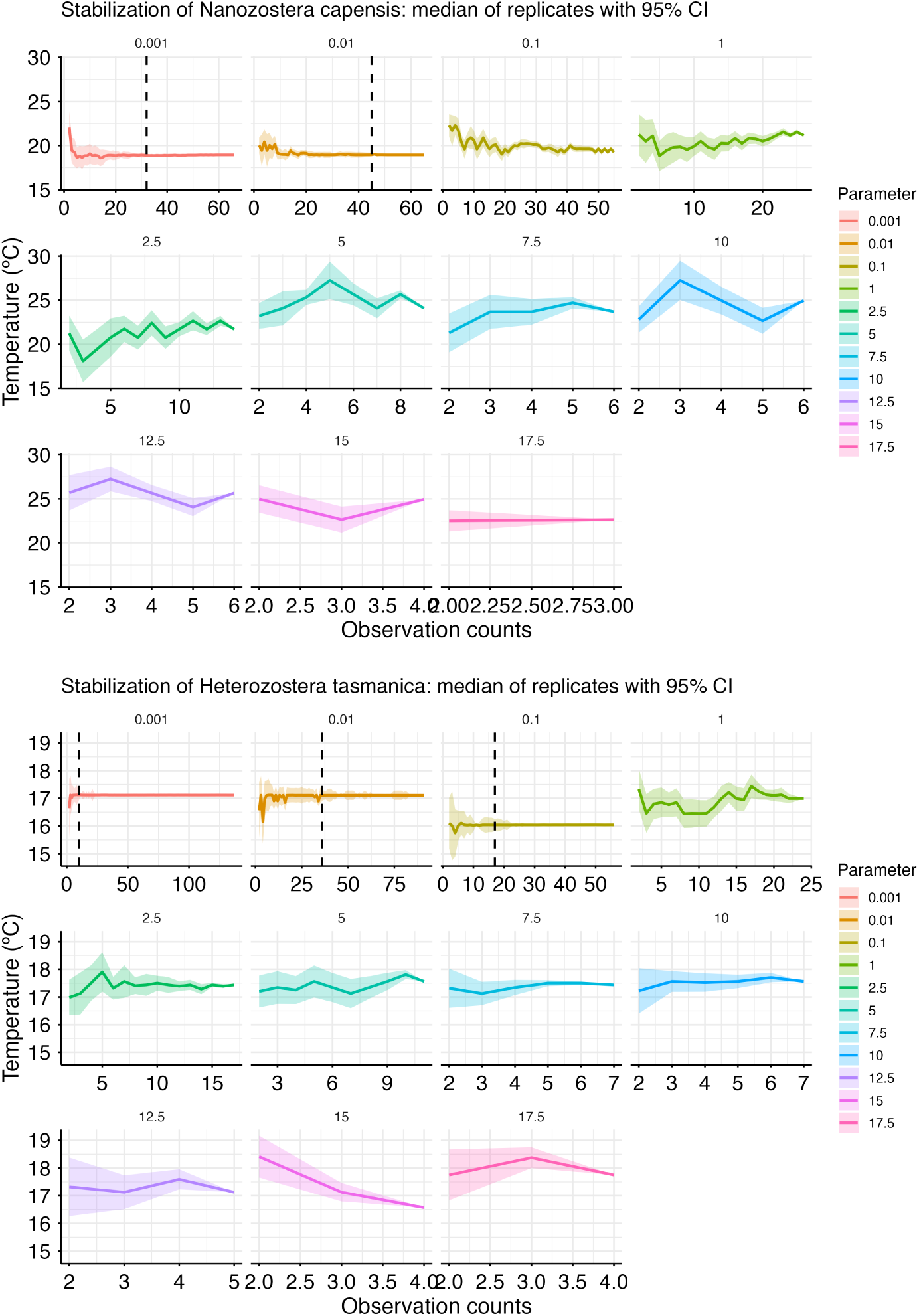
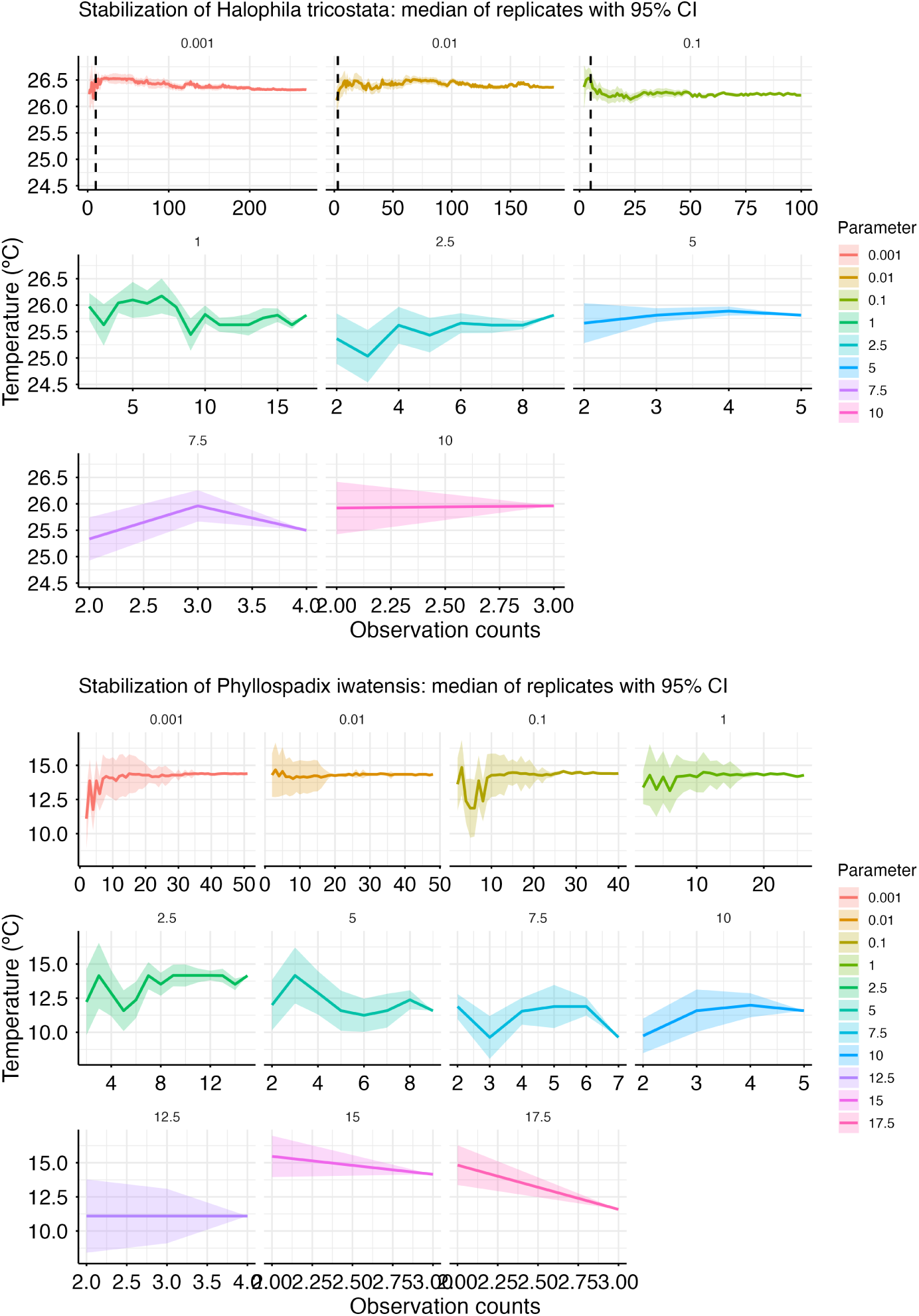
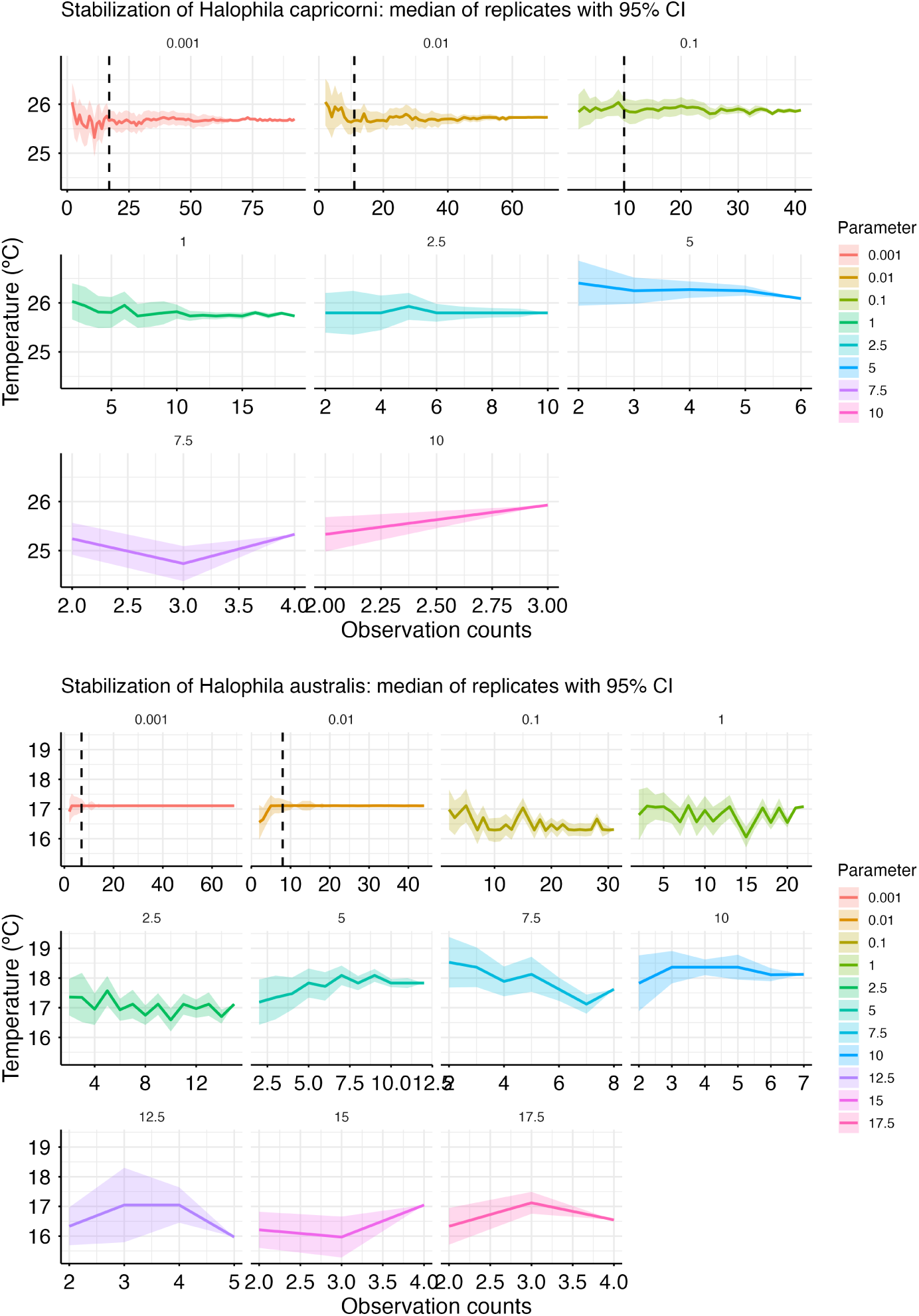
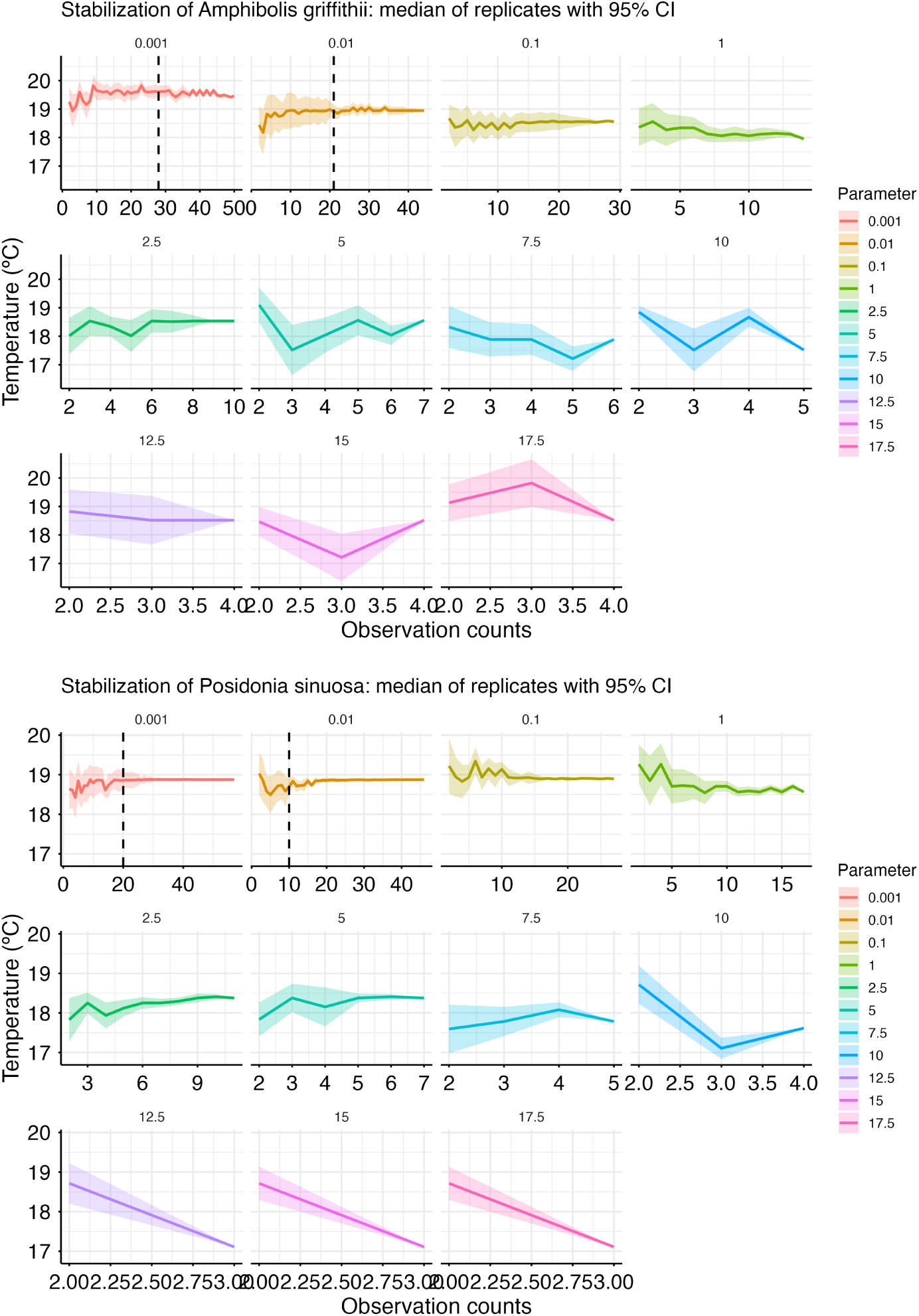
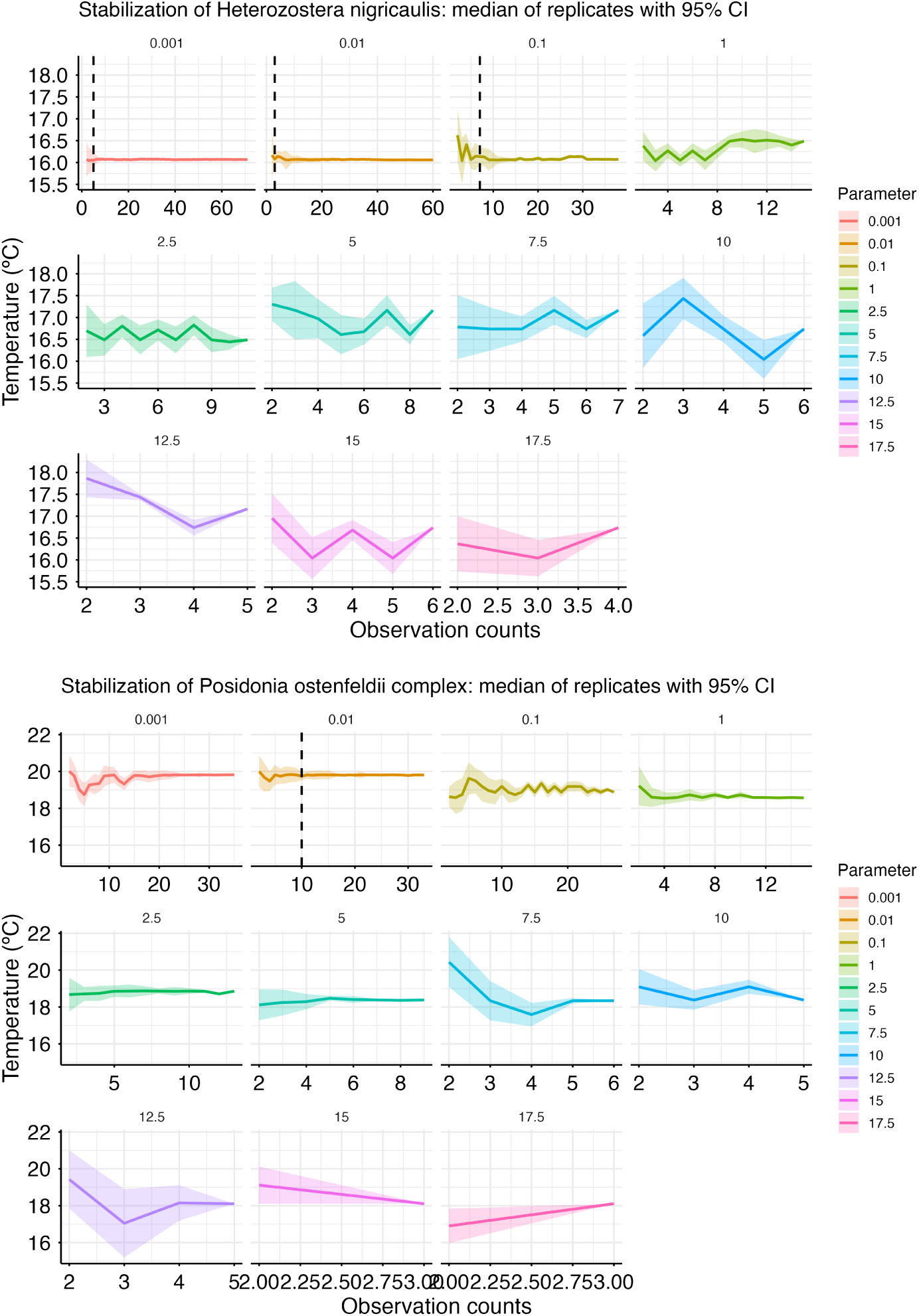
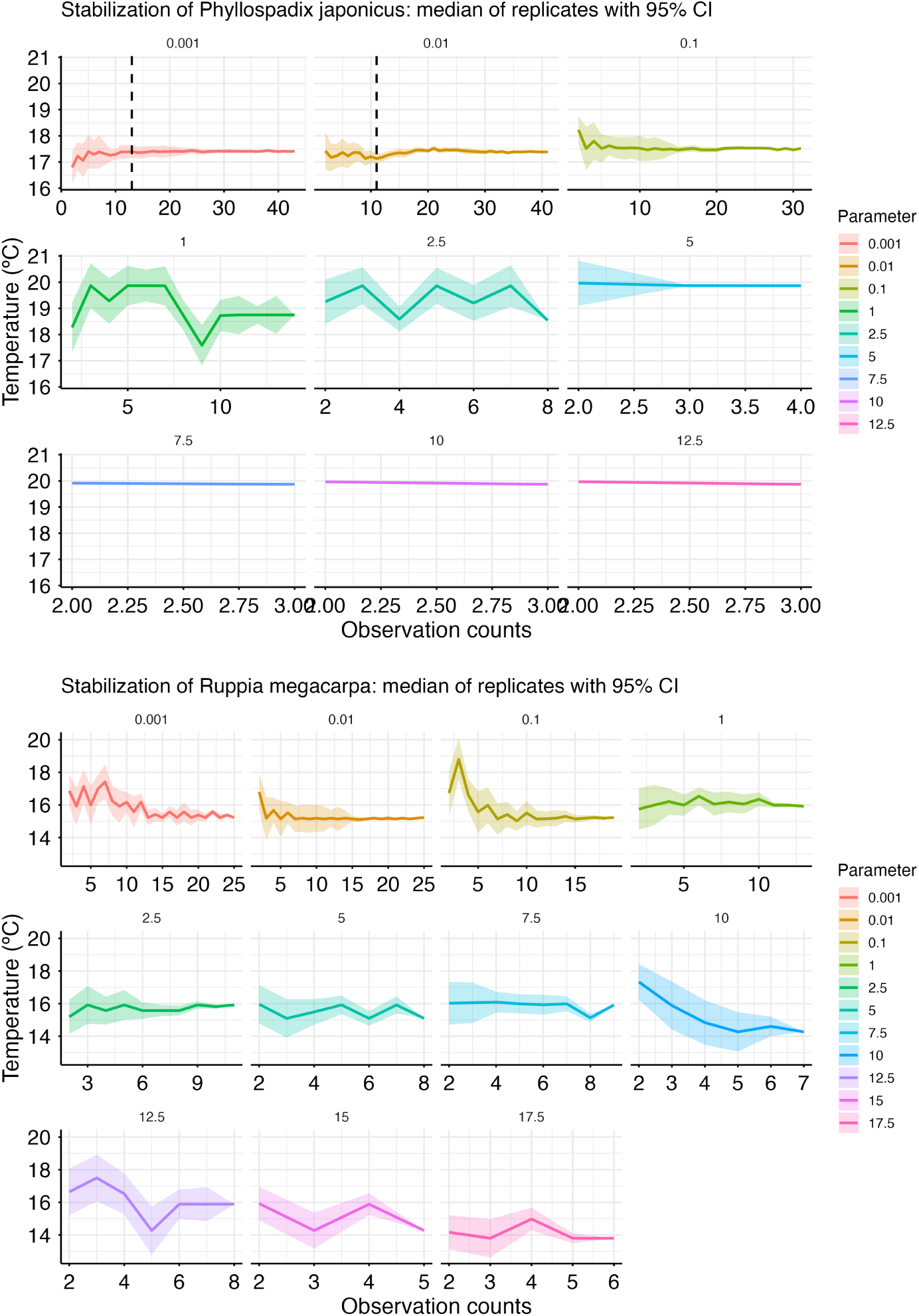
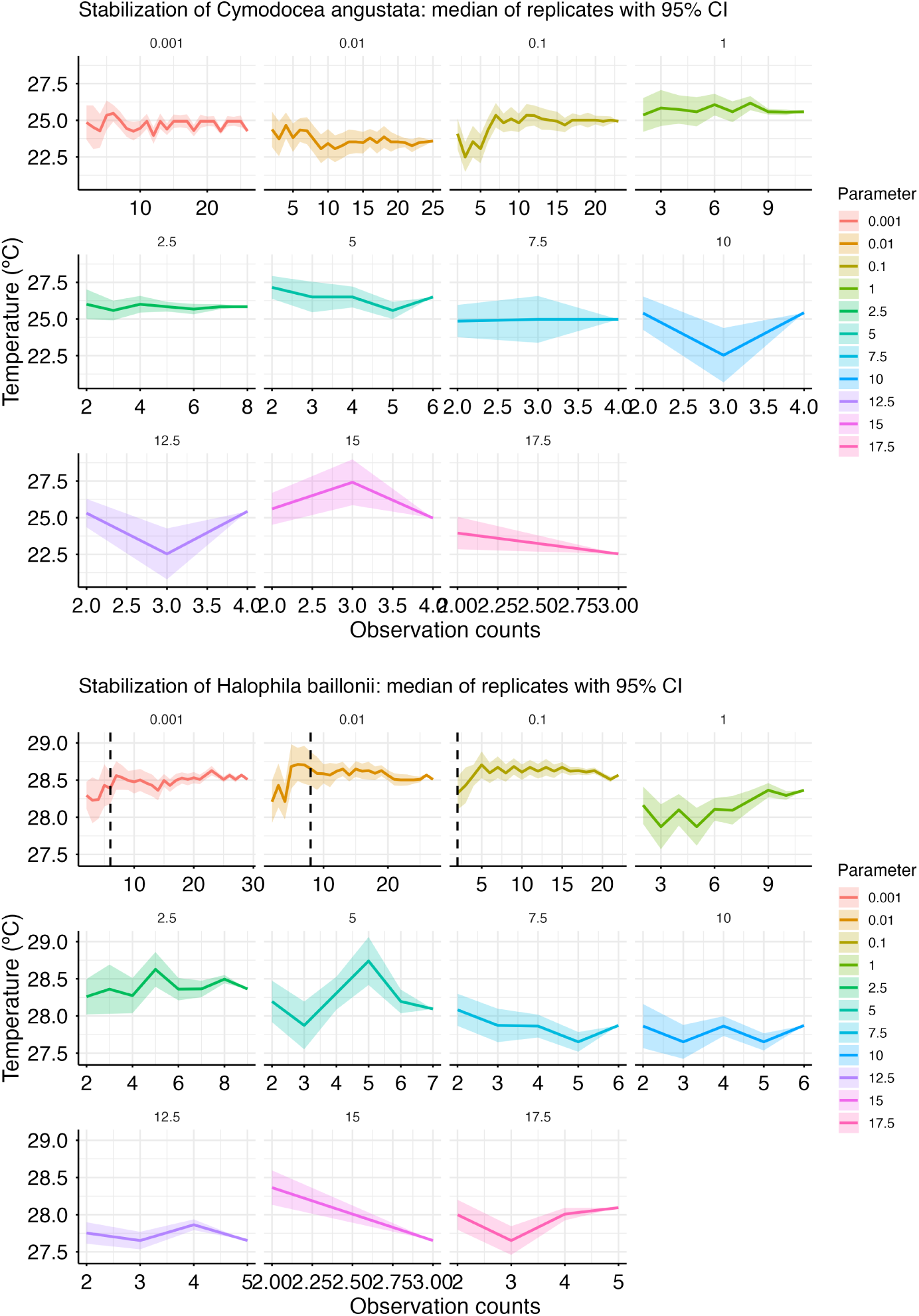
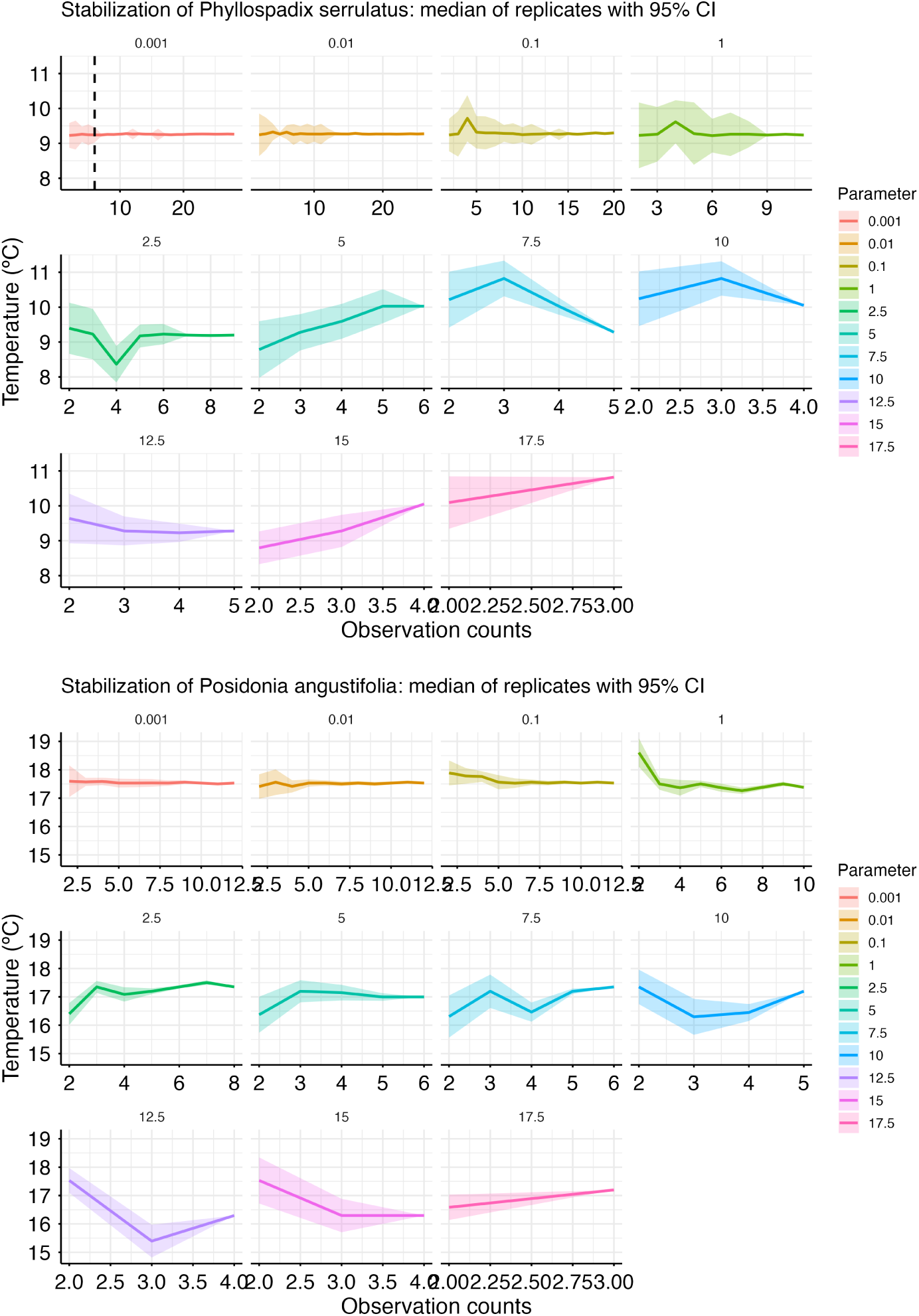
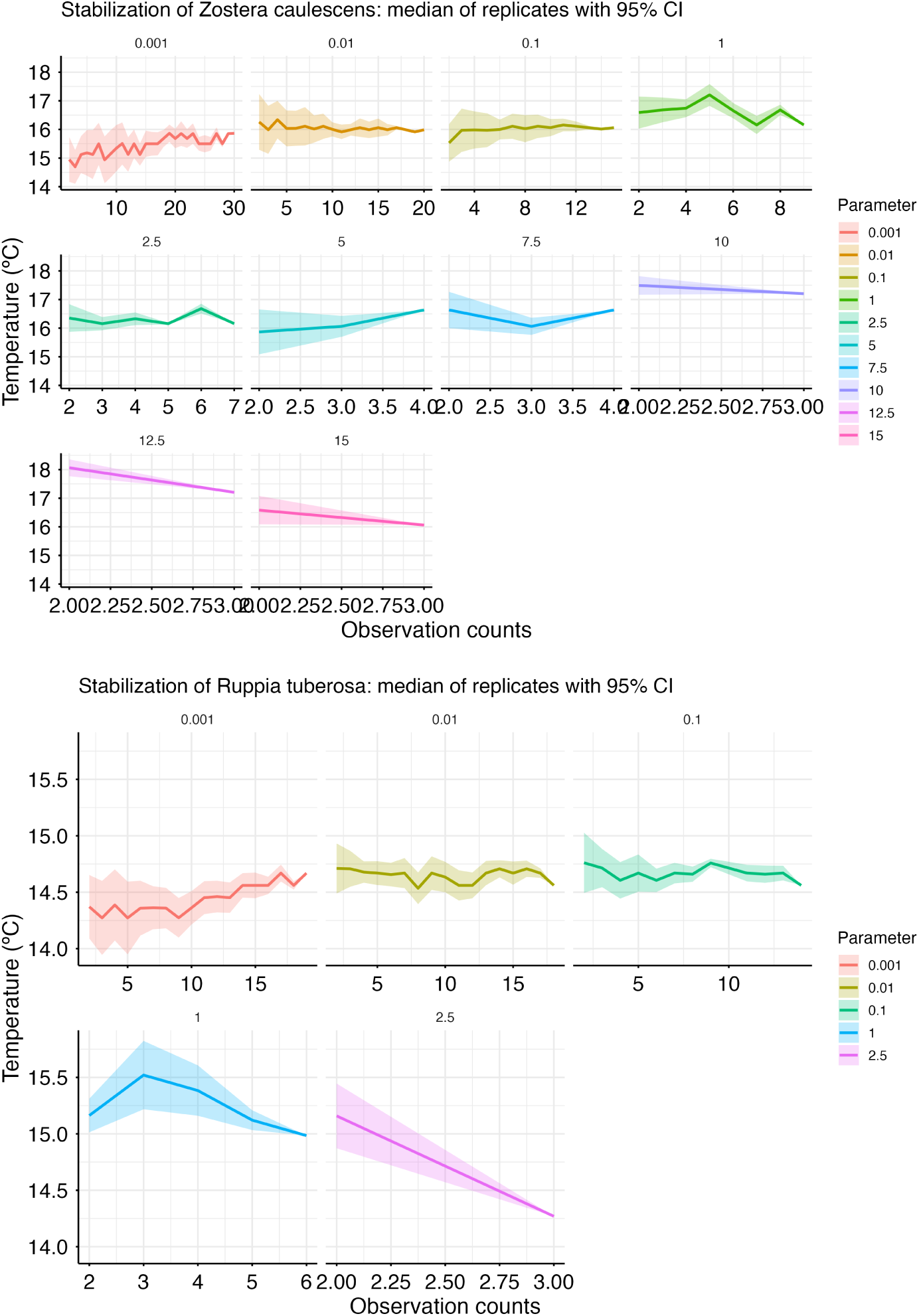
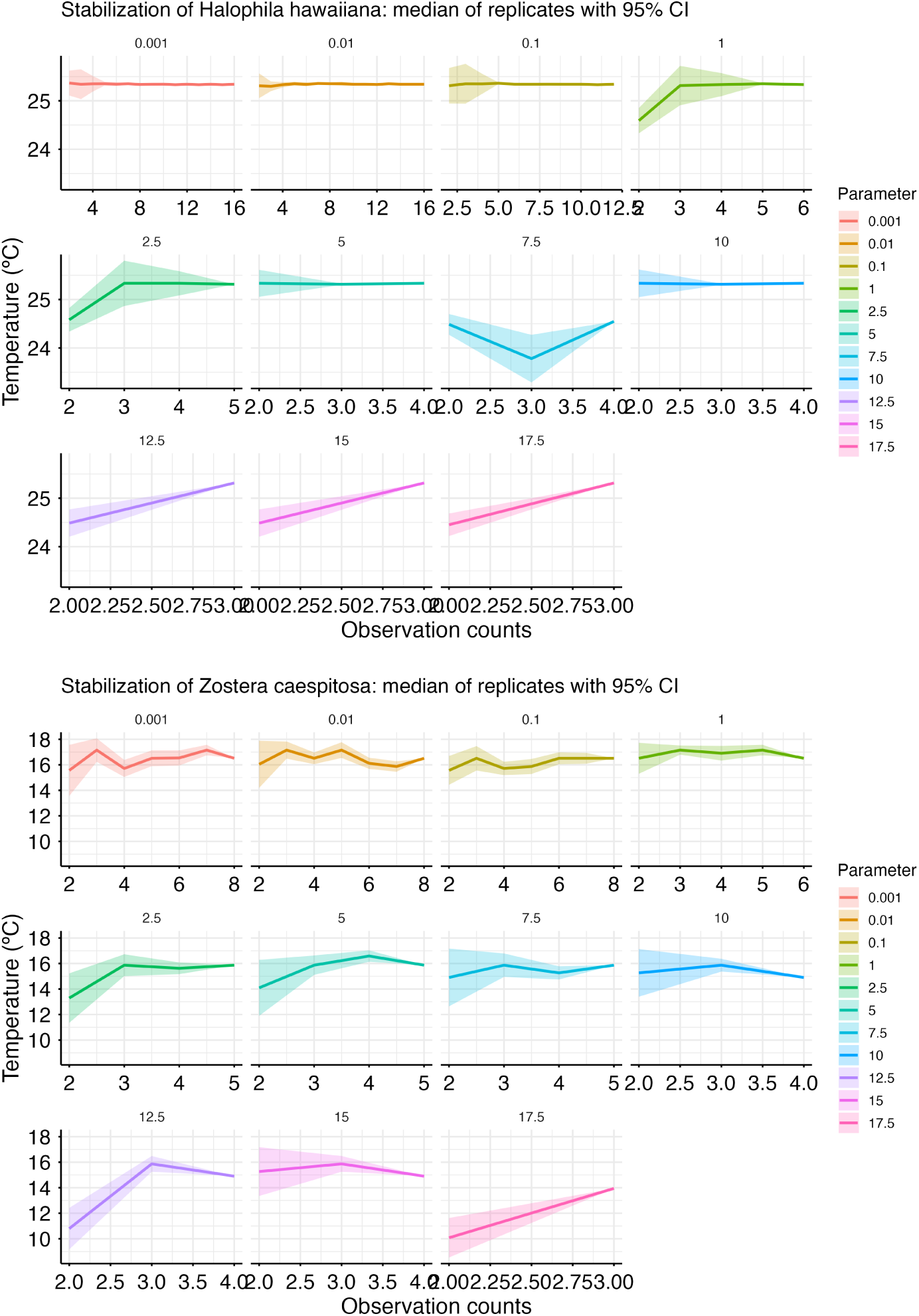
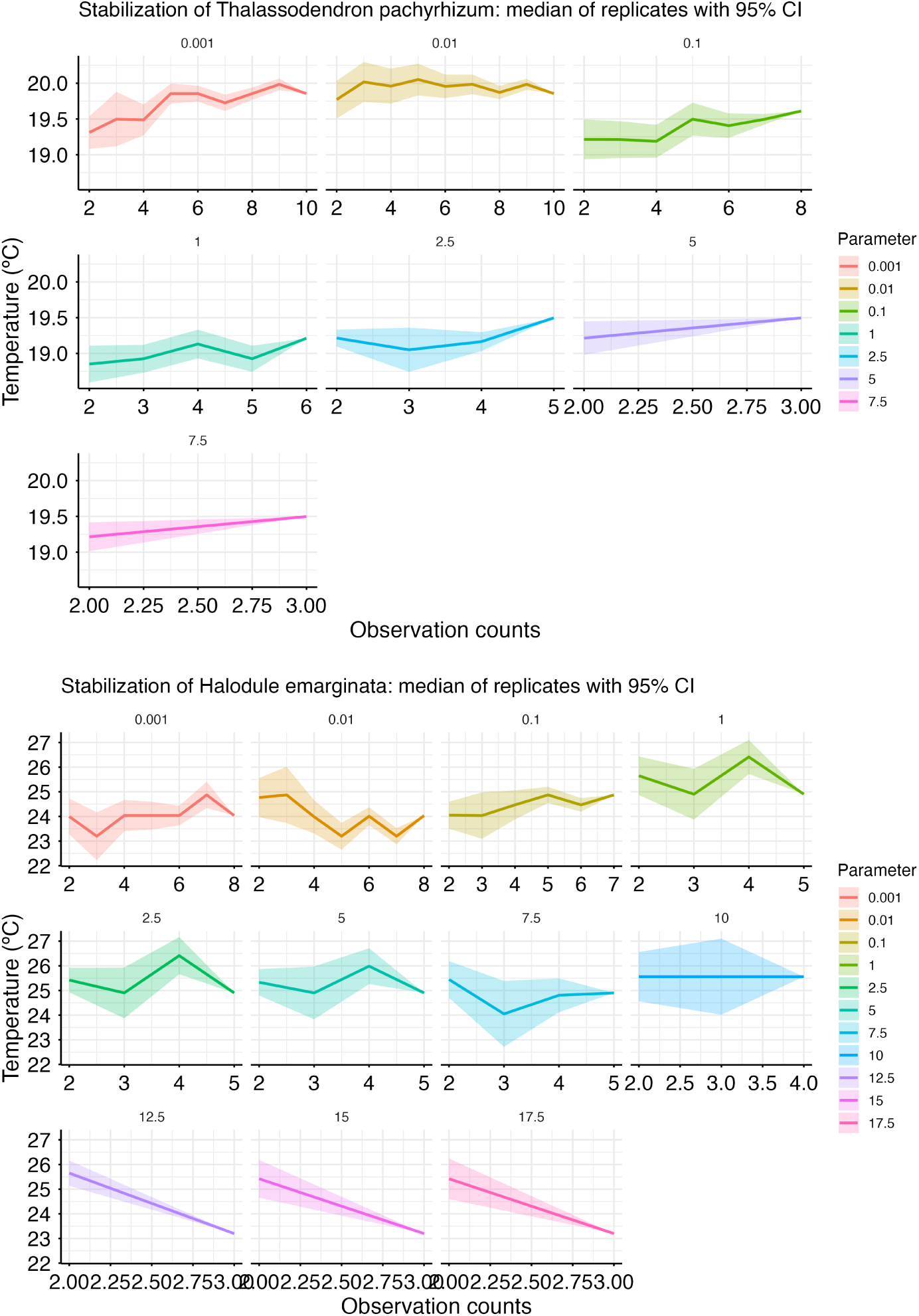
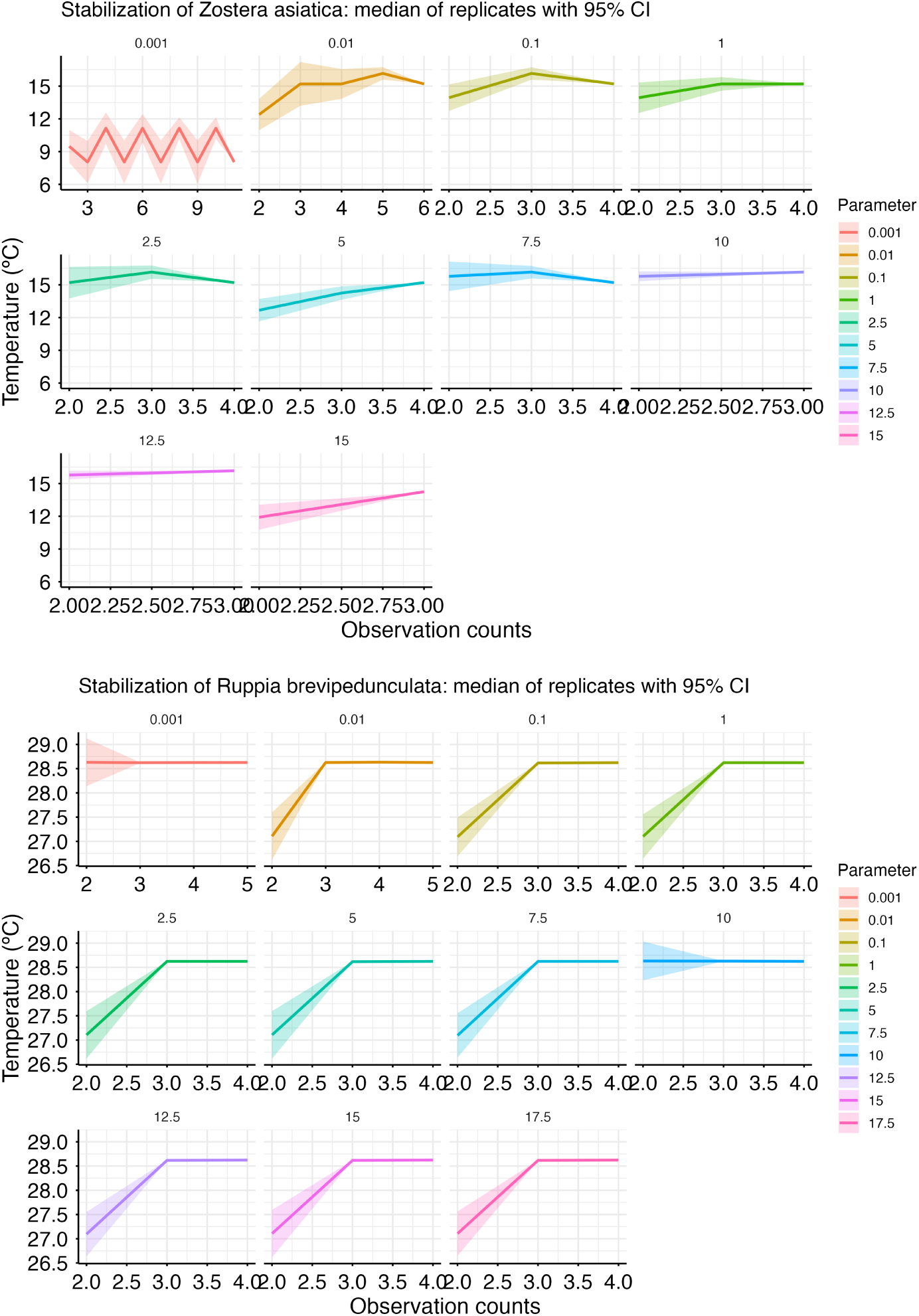
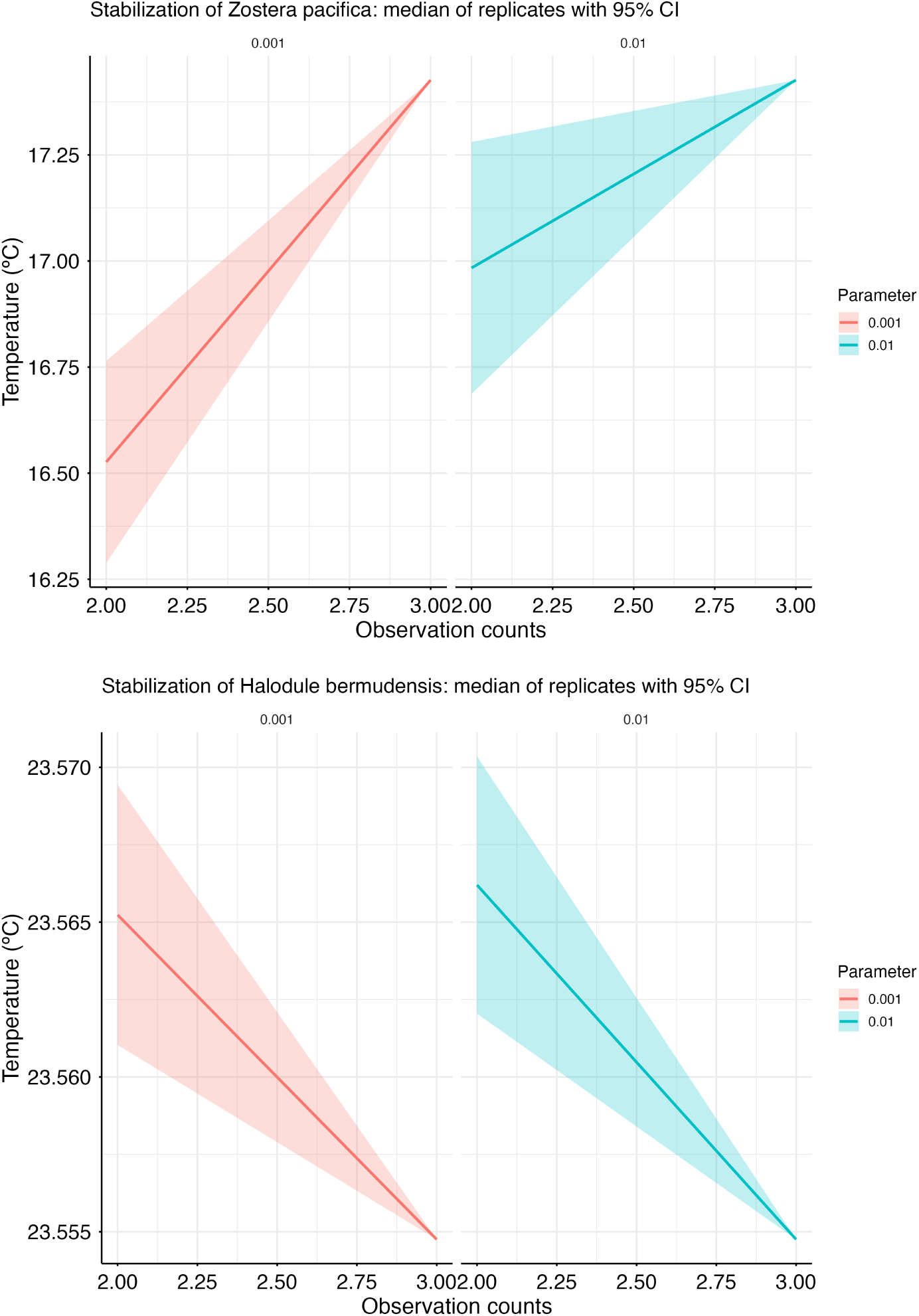

## S7: SENSITIVITY ANALYSES

### S7.1. Alternative range size resolutions

To evaluate the sensitivity of the framework to the spatial resolution used for calculating the range-size descriptor (*D_s_*), analyses were repeated using resolutions 0.25°, 0.50°, and 0.75° (0.5° *±* 50%). Fig. 4 summarises the coefficient of variation (CV) of the results in *D_s_*, and LS-TP and BD-TP parameters across the three resolutions, while Fig. 5 shows the resulting SVM decision surfaces. The metric used as an input (*D_s_*) varies considerably with resolution, yet the quantities of ecological interest (LS and BD parameters) remain largely unchanged.

Range size (Spearman *ρ >* 0.99), LS thinning parameter (Spearman *ρ* = 1), and BD thinning parameter estimates (Spearman *ρ >* 0.98) were highly consistent across spatial resolutions for all pairwise comparisons. As expected, *D_s_* values exhibited moderate variation between resolutions because the metric is directly derived from grid-cell size. LS parameters were identical across all resolutions (CV = 0 for all species), given its independence from *D_s_*. BD-TP predicted parameters showed minimal variation for most species, with only a few outliers displaying higher sensitivity. The main driver that explains the right-skewed CV-data distribution for the BD-TP comparison is the outlier *H. tricostata*, whose *θ_BD_* is capped at resolutions 0.5 and 0.75, but which is calculated as *θ* = 0.42 for the 0.25 resolution.

**Figure 4.**
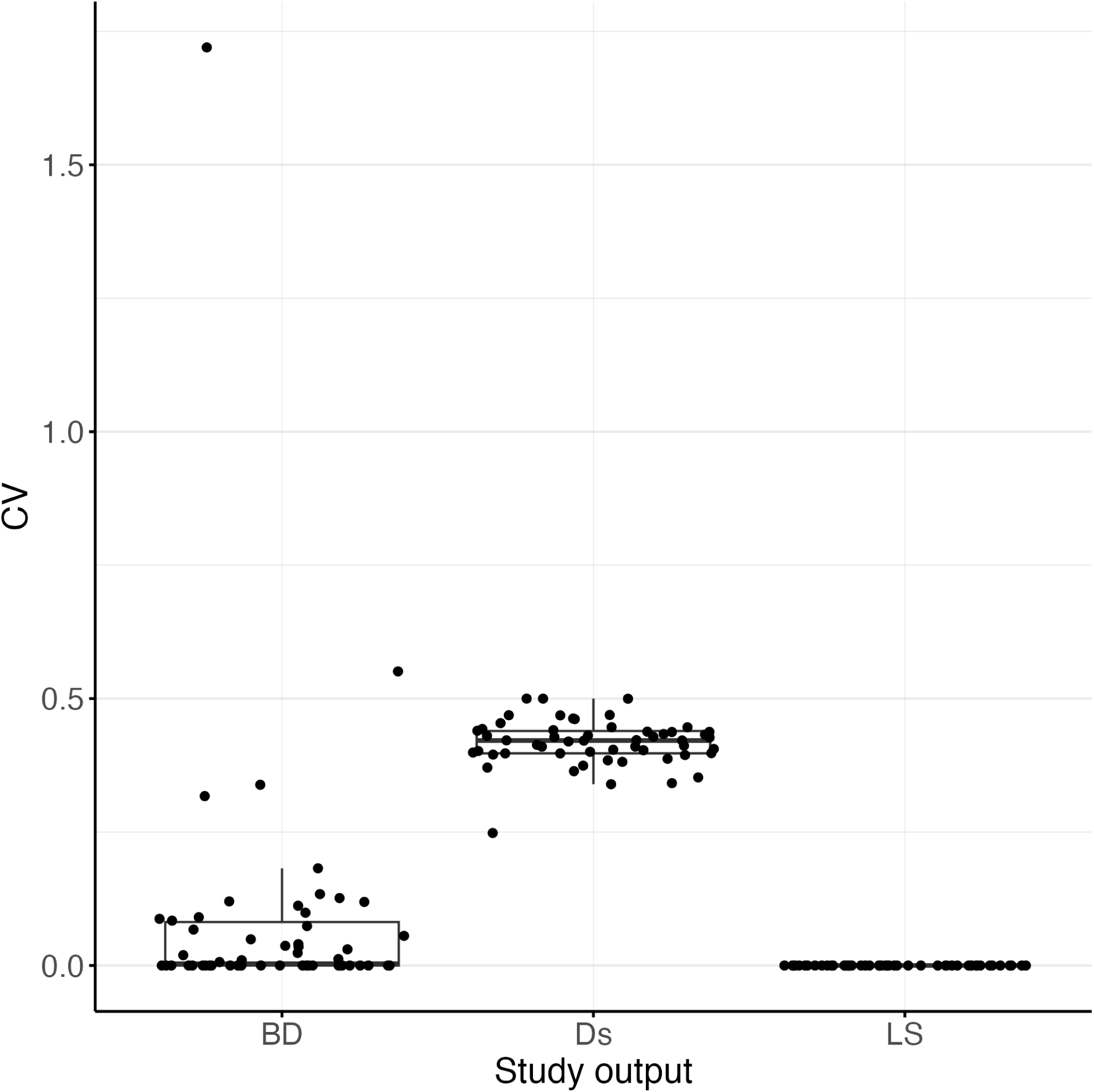
Distribution of the Coefficient of Variation for the study outputs (BD-TP, Ds and LS-TP) for each species (dots).

Comparison of the SVM decision hyperplanes revealed that increasing the resolution used to calculate *D_s_* primarily resulted in a vertical expansion of y-axis (i.e. larger *D_s_* values) and, thus, steeper hyperplanes, while the overall conclusion to be driven from the stability boundary (i.e. the predicted thinning parameter) remained largely unchanged. Hence, the framework is robust to the choice of spatial resolution for *D_s_* computation.

### S7.2. Alternative tolerance settings for the stabilization

Three tolerance settings are key for the stability computations: ‘tol value’, the maximum absolute difference from the final median (default = 0.3); ‘tol width’, the maximum confidence interval width (default = 0.5); and ‘window run’, the number of consecutive observations in which stability is kept (default = 20). A sensitivity analysis was run using deviations of *±*50% from the default. The proportion of species detected as always unstable (for all parameters) was used as a comparison along the 3^3^ possible tolerance setting combinations. Results were summarised in a heatmap (Fig. 6). Only window run affected the results clearly (in 2̃5% from 10 to 30). Tolerance width is the second setting with the highest impact on results although the variation is less marked. Differences due to the tolerance absolute value are negligible.

**Figure 5.**
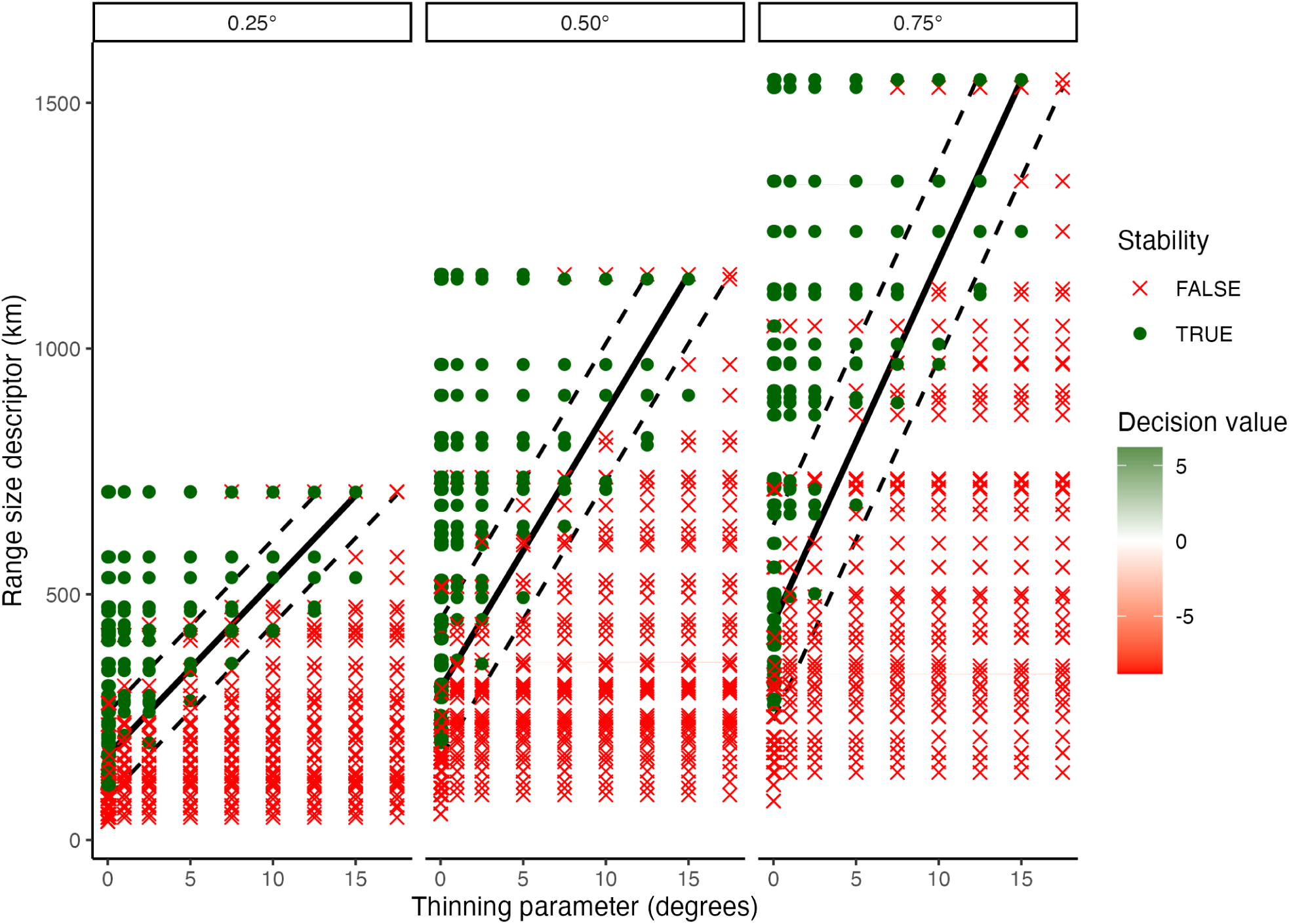
BD-TP and BD-LS results for the three tested resolutions for range size computation, rasterising original records with resolutions of 0.25, 0.5, and 0.75°.

**Figure 6.**
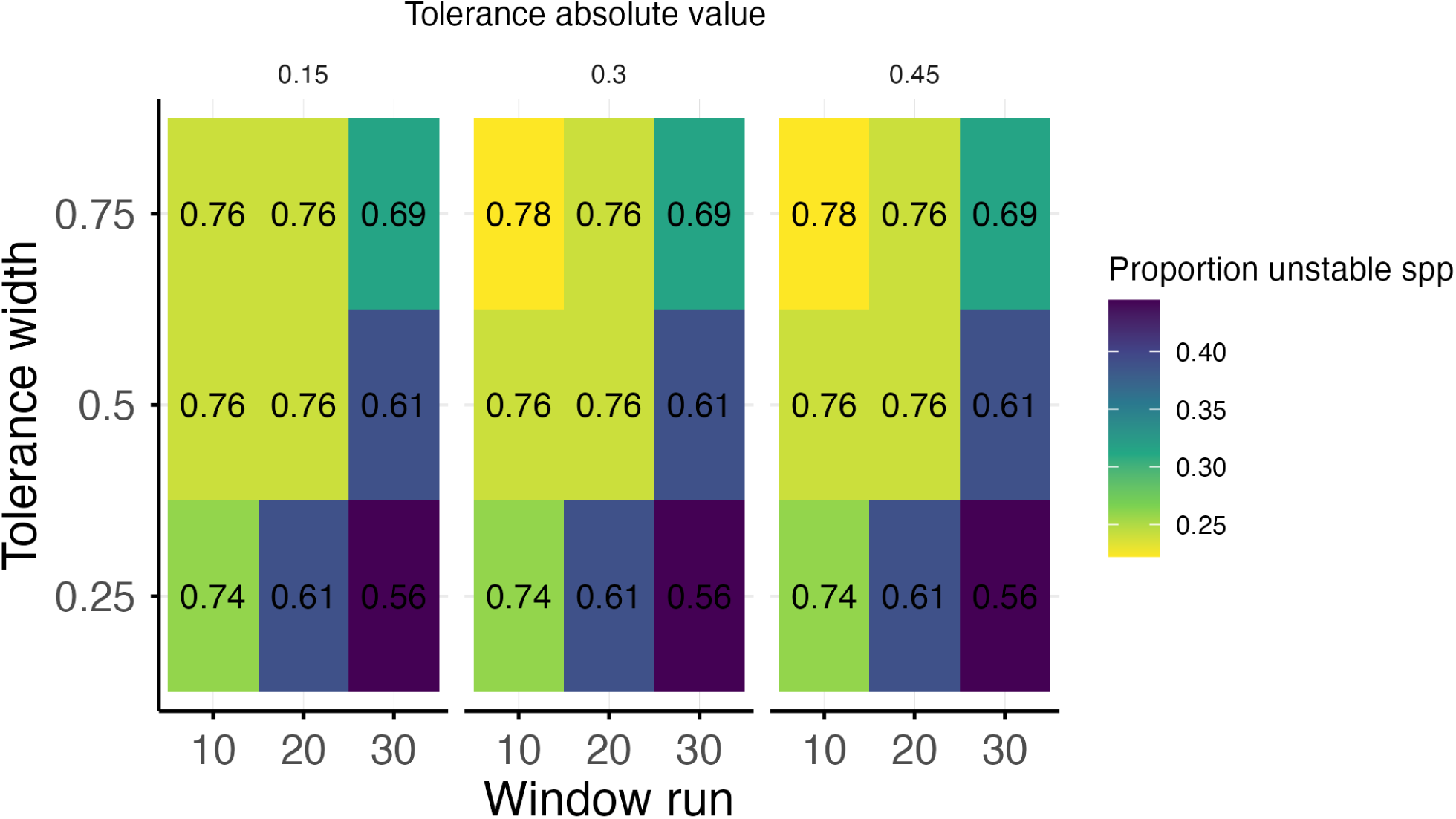
Heatmap showing the sensitivity analysis for tolerance settings. Alternative tolerance settings deviate by *±*50% the default. The proportion of species that are unstable at all thinning parameters (proportion stable spp.) was used for the comparison

### S7.3. Alternative parameter sets

In addition to the default parameter set, we tested different parameter grains, generating parameter sets with the following parameter quantities within the original range (0, 17.5]: 2, 4, 6, 8, 10, 12, 14, 16, 18, 20 (Table 8).

**Table 8.**
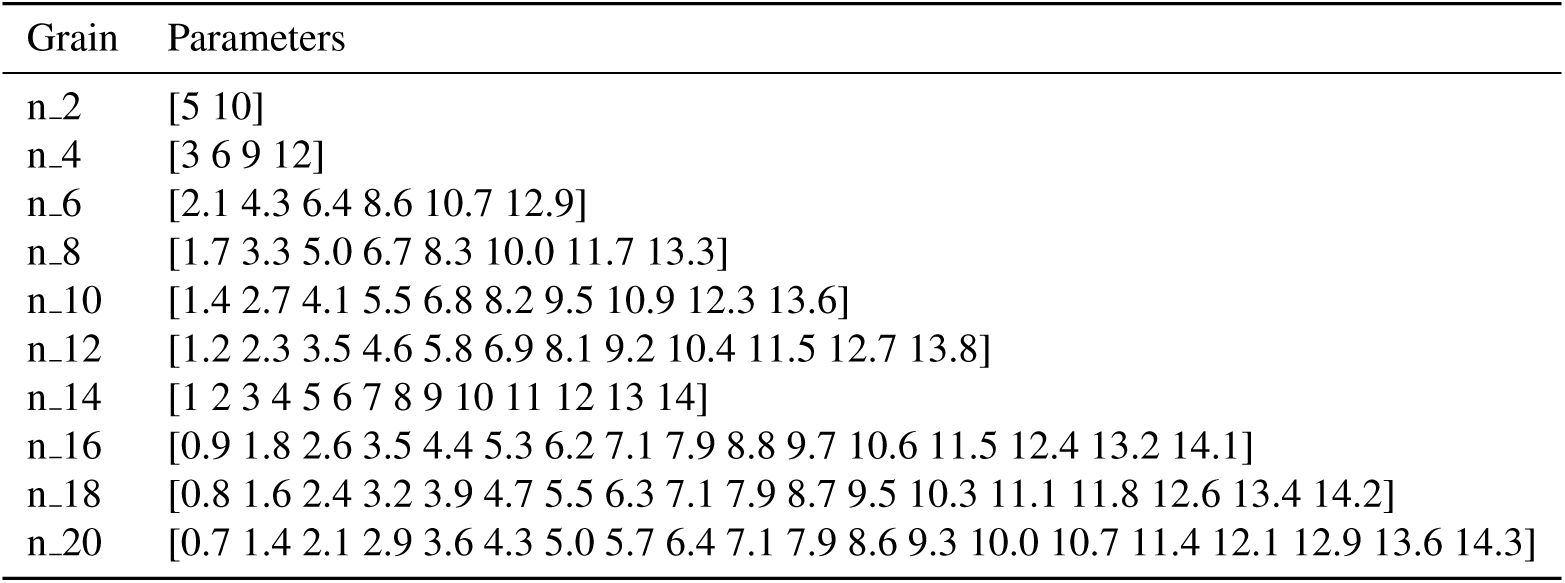
The 10 parameter sets of different grains (2 to 20) used for the sensitivity analysis.

Recalculating the main plot that summarises both BD-TP and LS-TP showed a similar pattern to the one predicted by the default settings, confirming our selection of 10 parameters appropriate as a default (Fig. 7).

**Figure 7.**
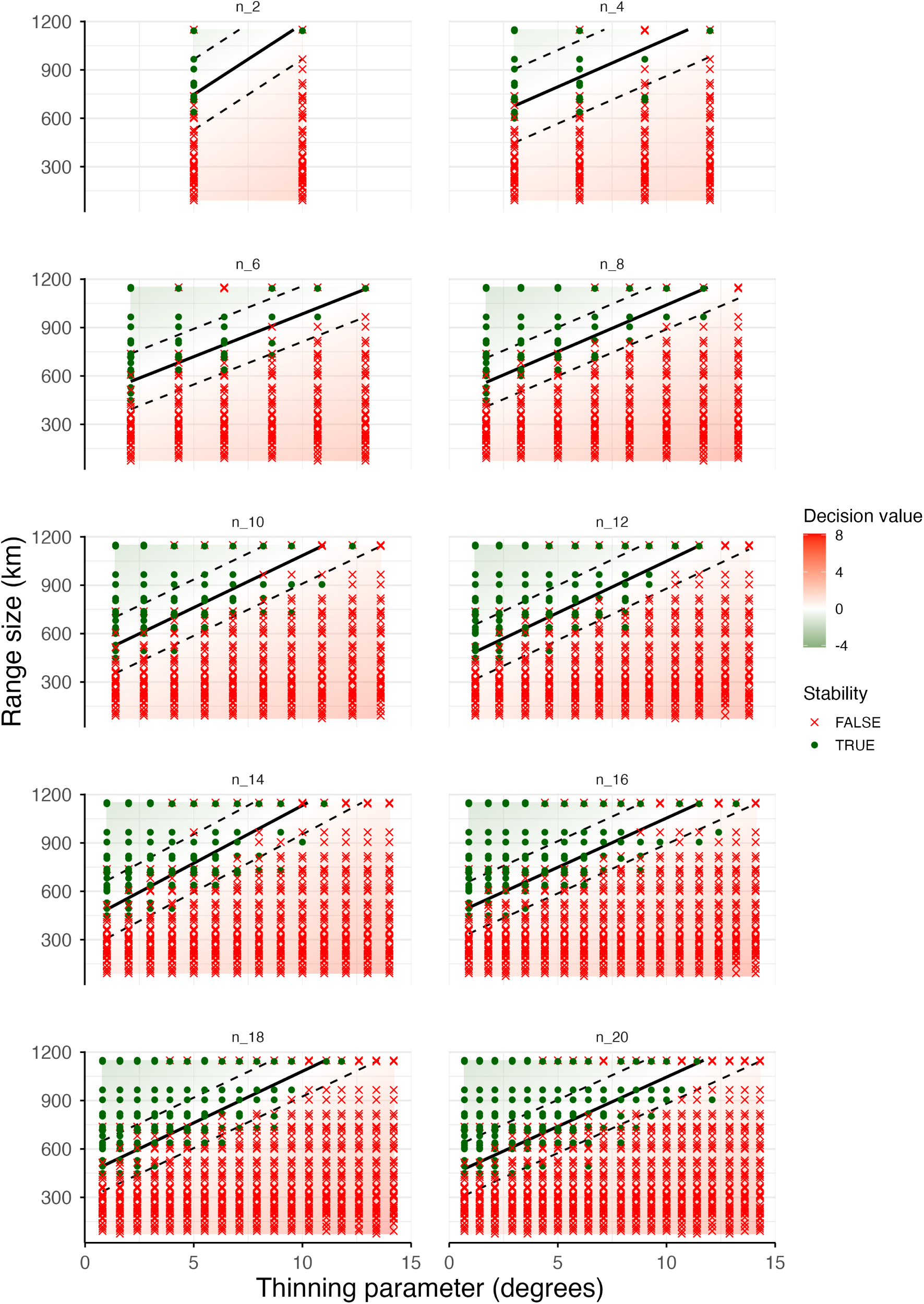
SVM plots showing the BD-TP results for different parameters grains (10 parameter sets). Thermally stable thinning parameter and species combinations are represented with green dots, whereas thermally unstable combinations are shown in red crosses. The black solid line represents the hyperplane predicted by BD-TP. Dashed lines indicate margin boundaries. The LS-TP output can be interpreted as the last stable symbol per species.

## S8: MODELING OUTPUTS

The validity of the proposed methods was assessed using *Phyllospadyx torreyi*, a seagrass species endemic to the North American Pacific coast with a small to intermediate geographic range. This case study provides an expanded comparison of thermal niche estimates and Habitat Suitability Model (HSM) outputs derived from four occurrence datasets: (i) the raw original records, (ii) data thinned to a 0.05° grid (matching the environmental raster resolution as commonly done in the literature), (iii) data thinned using the parameter predicted by LS, and (iv) data thinned using the parameter predicted by BD. Furthermore, models calibrated with the original dataset were compared with those generated from an artificially biased dataset. To simulate sampling bias, 300 additional occurrence records were generated within the combined 20-km buffers surrounding the cities of San Francisco, Los Angeles, and San Diego. For each dataset, the results include: (i) a map of occurrence records, (ii) a Habitat Suitability Index (HSI) map, (iii) cross-validated HSM performance metrics, and (iv) realised thermal niche (RTNs) curves. LS predicted an optimal thinning parameter of 0.1 for both the original and artificially biased datasets, whereas BD predicted parameters of 0.6 and 1.0, respectively.

Habitat suitability models were developed using the R-based online platform GLOSSA (Mestre-Tomás et al., 2026). Five environmental predictors commonly used in seagrass HSMs were selected (Bertelli et al., 2022): slope, sea surface temperature, salinity, bathymetry, and photosynthetically active radiation, all obtained from Bio-ORACLE v3 (Assis et al., 2024). Model calibration employed the habitat suitability workflow implemented in GLOSSA, using cross-validation with ten spatial and temporal folds. Pseudo-absences were generated using the buffer-restricted method with a presence-to-absence ratio of 1:1 and a buffer distance of 0.2 decimal degrees. Models were fitted using the Bayesian Additive Regression Trees (BART) algorithm. Thermal response curves were estimated as density distributions using repeated cross-validation (50 iterations, each using 60% of the occurrence records).

The amount of input occurrence records varied across methods in both the original (raw = 178, 0.05° = 96, LS = 89, BD = 47) and artificially biased (raw = 478, 0.05° = 196, LS = 121, BD = 34) datasets.

Cross-validated performance metrics differed among thinning methods. LS and, particularly, BD generally produced lower evaluation scores of greater variability than the raw and 0.05°-thinned datasets. However, the artificially biased dataset yielded improved performance metrics for the latter two approaches. This pattern suggests that internal cross-validation metrics primarily quantify model fit to the available data rather than ecological realism. Consequently, artificially biased datasets may produce deceptively high performance estimates despite stronger overfitting. Conversely, the lower evaluation scores observed for LS and BD might reflect their more effective reduction of sampling bias, which decreases the number of occurrence records available for model calibration.

**Figure 8.**
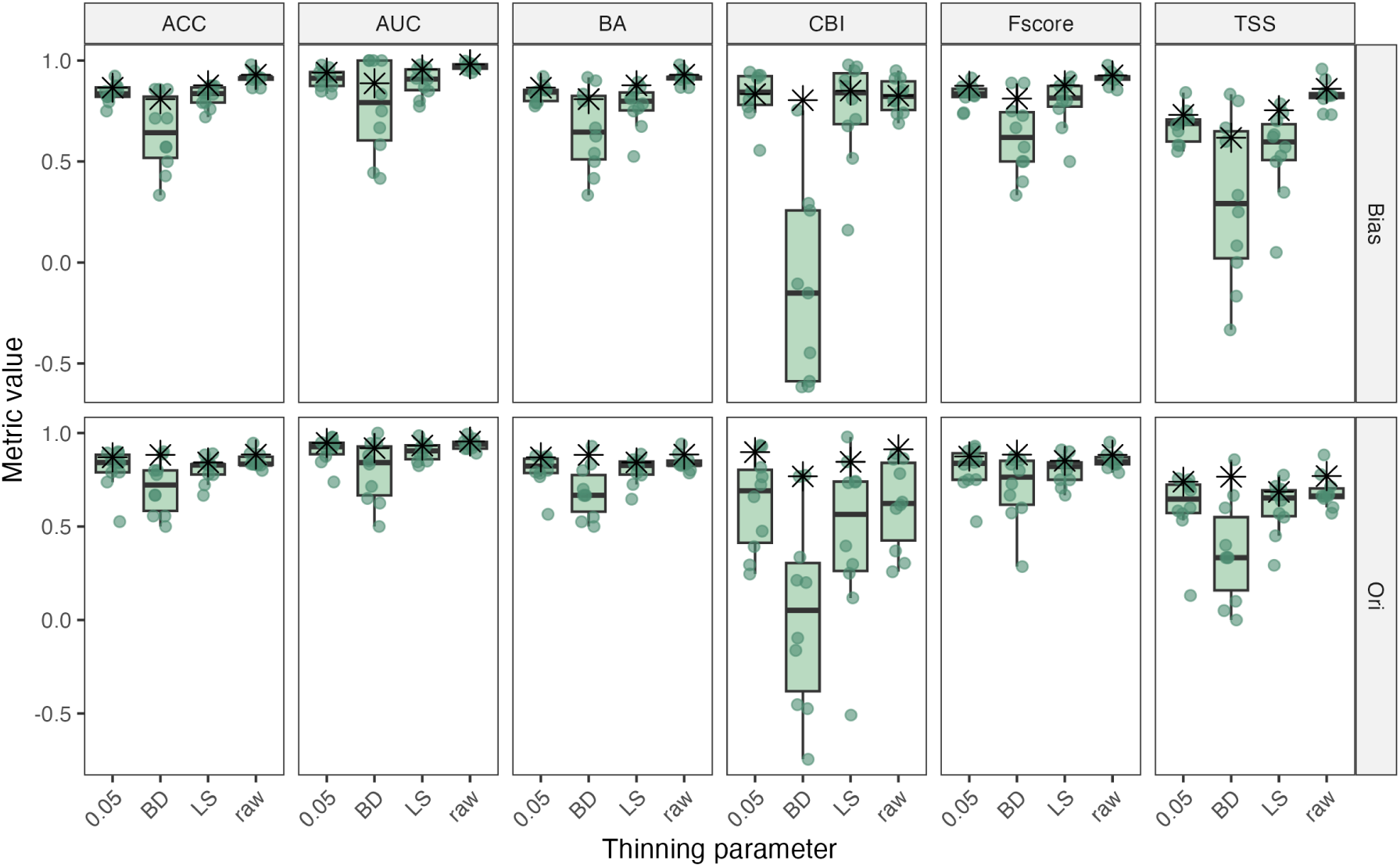
K-fold cross-validated (boxplots) and total value for evaluation metrics. Cross-validation was performed as a subset of the thinned dataset.

The standard deviation of the predicted values across cross-validation folds of the HSM provided a spatial overview of model uncertainty. HSI values were low for all cases.

**Figure 9.**
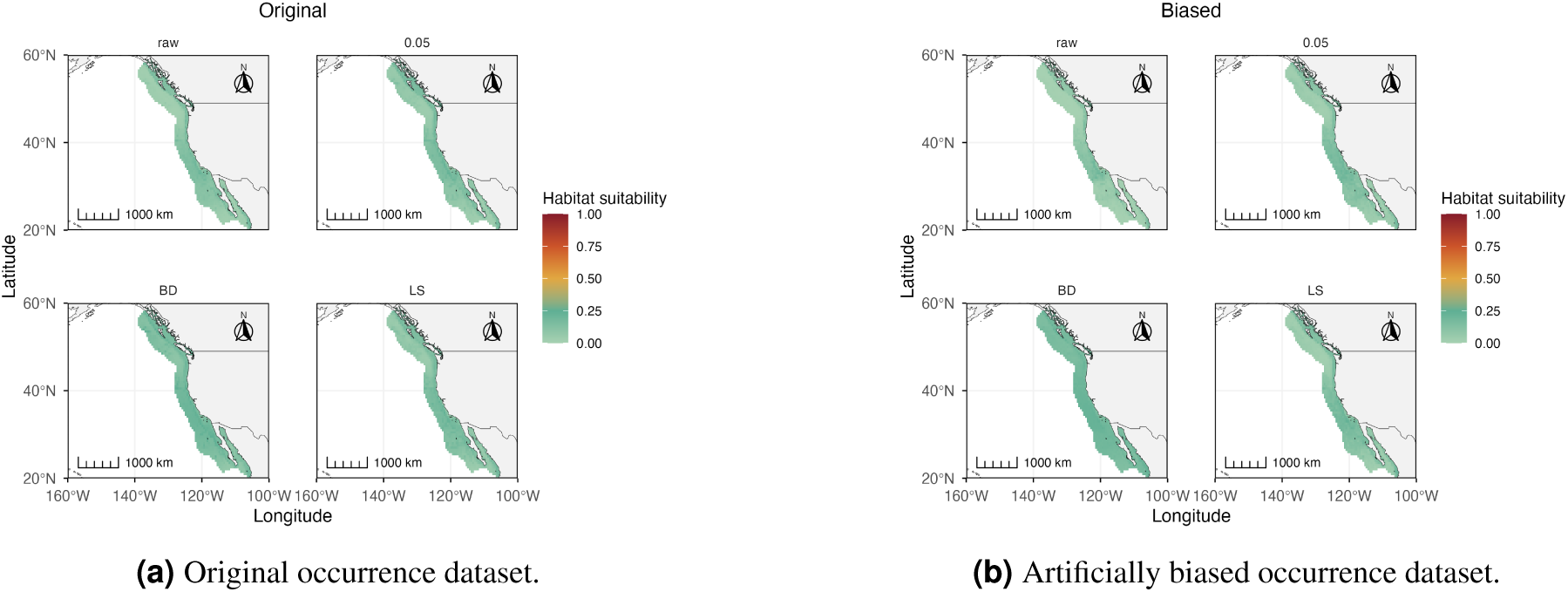
Maps showing the standard deviation of habitat suitability maps for *Phyllospadix torreyi* based on (a) the original and (b) artificially biased occurrence datasets. Habitat suitability indexes were calculated with BART and SD values are shown with a gradient fill. Both in (a) and (b), the raw data (raw) and the data thinned with 0.05 and the predicted parameters by BD and LS were inputted into the model. The predictive distribution is shown as the median.

## REFERENCES

Adelino, J. R. P., Anjos, L. d., and Lima, M. R. (2017). Invasive potential of the pied crow (*corvus albus*) in eastern Brazil: best to eradicate before it spreads. Perspectives in Ecology and Conservation, 15(3):227–233.

Aiello-Lammens, M. E., Boria, R. A., Radosavljevic, A., Vilela, B., and Anderson, R. P. (2015). spThin: an R package for spatial thinning of species occurrence records for use in ecological niche models. Ecography, 38(5):541–545.

Araújo, M. B., Anderson, R. P., Márcia Barbosa, A., Beale, C. M., Dormann, C. F., Early, R., Garcia, R. A., Guisan, A., Maiorano, L., Naimi, B., O’Hara, R. B., Zimmermann, N. E., and Rahbek, C. (2019). Standards for distribution models in biodiversity assessments. Science Advances, 5(1).

Assis, J., Fernández Bejarano, S. J., Salazar, V. W., Schepers, L., Gouvêa, L., Fragkopoulou, E., Leclercq, F., Vanhoorne, B., Tyberghein, L., Serrão, E. A., Verbruggen, H., and De Clerck, O. (2024). Bio-oracle v3.0. pushing marine data layers to the cmip6 earth system models of climate change research. Global Ecology and Biogeography, 33(4).

Ball-Damerow, J. E., Brenskelle, L., Barve, N., Soltis, P. S., Sierwald, P., Bieler, R., LaFrance, R., Ariño, A. H., and Guralnick, R. P. (2019). Research applications of primary biodiversity databases in the digital age. PLOS ONE, 14(9):e0215794.

Bertelli, C. M., Stokes, H. J., Bull, J. C., and Unsworth, R. K. F. (2022). The use of habitat suitability modelling for seagrass: A review. Frontiers in Marine Science, 9.

Boria, R. A., Olson, L. E., Goodman, S. M., and Anderson, R. P. (2014). Spatial filtering to reduce sampling bias can improve the performance of ecological niche models. Ecological Modelling, 275:73–77.

Bried, J. T. and Siepielski, A. M. (2017). Opportunistic data reveal widespread species turnover in Enallagma damselflies at biogeographical scales. Ecography, 41(6):958–970.

Burrows, M. T., Hawkins, S. J., Moore, J. J., Adams, L., Sugden, H., Firth, L., and Mieszkowska, N. (2020). Global-scale species distributions predict temperature-related changes in species composition of rocky shore communities in Britain. Global Change Biology, 26(4):2093–2105.

Castellanos, A. A., Huntley, J. W., Voelker, G., and Lawing, A. M. (2019). Environmental filtering improves ecological niche models across multiple scales. Methods in Ecology and Evolution, 10:481 – 492.

Castle, L., Pinder, J., Lillis, H., and Manca, E. (2022). Using an ensemble modelling approach to predict suitable habitat for *zostera marina* beds, *modiolus modiolus* beds and *sabellaria spinulosa* reefs in UK waters. Technical report, JNCC.

Chamberlain, S., Barve, V., Mcglinn, D., Oldoni, D., Desmet, P., Geffert, L., and Ram, K. (2017). R Python, and Ruby clients for GBIF species occurrence data. PeerJ PrePrints.

Chefaoui, R. M., Duarte, C. M., and Serrão, E. A. (2018). Dramatic loss of seagrass habitat under projected climate change in the Mediterranean Sea. Global Change Biology, 24(10):4919–4928.

Coca de la Iglesia, M., Valcárcel, V., and Medina, N. (2023). A Protocol to Retrieve and Curate Spatial and Climatic Data from Online Biodiversity Databases Using R. BIO-PROTOCOL, 13(20).

Coron, C., Calenge, C., Giraud, C., and Julliard, R. (2018). Bayesian estimation of species relative abundances and habitat preferences using opportunistic data. Environmental and Ecological Statistics, 25(1):71–93.

Cretois, B., Simmonds, E. G., Linnell, J. D. C., van Moorter, B., Rolandsen, C. M., Solberg, E. J., Strand, O., Gundersen, V., Roer, O., and Rød, J. K. (2021). Identifying and correcting spatial bias in opportunistic citizen science data for wild ungulates in Norway. Ecology and Evolution, 11(21):15191–15204.

Dickinson, J. L., Zuckerberg, B., and Bonter, D. N. (2010). Citizen science as an ecological research tool: Challenges and benefits. Annual Review of Ecology, Evolution, and Systematics, 41(1):149–172.

Dorey, J. B., Fischer, E. E., Chesshire, P. R., Nava-Bolaños, A., O’Reilly, R. L., Bossert, S., Collins, S. M., Lichtenberg, E. M., Tucker, E. M., Smith-Pardo, A., Falcon-Brindis, A., Guevara, D. A., Ribeiro, B., de Pedro, D., Pickering, J., Hung, K.-L. J., Parys, K. A., McCabe, L. M., Rogan, M. S., Minckley, R. L., Velazco, S. J. E., Griswold, T., Zarrillo, T. A., Jetz, W., Sica, Y. V., Orr, M. C., Guzman, L. M., Ascher, J. S., Hughes, A. C., and Cobb, N. S. (2023). A globally synthesised and flagged bee occurrence dataset and cleaning workflow. Scientific Data, 10(1).

Duarte, C. M., Dennison, W. C., Orth, R. J. W., and Carruthers, T. J. B. (2008). The charisma of coastal ecosystems: Addressing the imbalance. Estuaries and Coasts, 31(2):233–238.

Erickson, K. D. and Smith, A. B. (2021). Accounting for imperfect detection in data from museums and herbaria when modeling species distributions: combining and contrasting data-level versus model-level bias correction. Ecography, 44(9):1341–1352.

Farooq, S. (2025). Climate-driven spread of giant hogweed *heracleum mantegazzianum* (Sommier & Levier) in Turkey: assessing future invasion risks under CMIP6 climate projections. BMC Plant Biology, 25(1).

Gantchoff, M. G., Conlee, L., and L. Belant, J. (2022). The effectiveness of opportunistic public reports versus professional data to estimate large carnivore distribution. Ecosphere, 13(2).

Gaston, K. J. and Fuller, R. A. (2009). The sizes of species’ geographic ranges. Journal of Applied Ecology, 46(1):1–9.

Hu, Z., Zhang, Q., Zhang, J., Kass, J. M., Mammola, S., Fresia, P., Draisma, S. G. A., Assis, J., Jueterbock, A., Yokota, M., and Zhang, Z. (2021). Intraspecific genetic variation matters when predicting seagrass distribution under climate change. Molecular Ecology, 30(15).

Huang, K., Lai, F., Chen, M., Song, Y., Chen, S., Wubuaysan, Z., and Zhuang, X. (2025). Morphological variation and spatial distribution patterns of *krascheninnikovia compacta* (Losinsk.) Grubov in the Tibetan antelope breeding grounds of the western Kunlun Mountains. Plants, 14(9):1298.

Isaac, N. J. B., van Strien, A. J., August, T. A., de Zeeuw, M. P., and Roy, D. B. (2014). Statistics for citizen science: extracting signals of change from noisy ecological data. Methods in Ecology and Evolution, 5(10):1052–1060.

Jones, M. C. and Cheung, W. W. L. (2014). Multi-model ensemble projections of climate change effects on global marine biodiversity. ICES Journal of Marine Science, 72(3):741–752.

Jueterbock, A., Tyberghein, L., Verbruggen, H., Coyer, J. A., Olsen, J. L., and Hoarau, G. (2013). Climate change impact on seaweed meadow distribution in the north atlantic rocky intertidal. Ecology and Evolution, 3(5):1356–1373.

Koldasbayeva, D., Tregubova, P., Shadrin, D., Gasanov, M., and Pukalchik, M. (2022). Large-scale forecasting of *heracleum sosnowskyi* habitat suitability under the climate change on publicly available data. Scientific Reports, 12(1).

Laeseke, P., Martínez, B., Mansilla, A., and Bischof, K. (2021). Invaders in waiting? non-equilibrium in southern hemisphere seaweed distributions may lead to underestimation of Antarctic invasion potential. Frontiers of Biogeography, 13(4).

Lamboley, Q. and Fourcade, Y. (2024). No optimal spatial filtering distance for mitigating sampling bias in ecological niche models. Journal of Biogeography, 51(9):1783–1794.

Maldonado, C., Molina, C. I., Zizka, A., Persson, C., Taylor, C. M., Albán, J., Chilquillo, E., Rønsted, N., and Antonelli, A. (2015). Estimating species diversity and distribution in the era of big data: to what extent can we trust public databases? Global Ecology and Biogeography, 24(8):973–984.

Manca, F., Benedetti-Cecchi, L., Bradshaw, C. J. A., Cabeza, M., Gustafsson, C., Norkko, A. M., Roslin, T. V., Thomas, D. N., White, L., and Strona, G. (2024). Projected loss of brown macroalgae and seagrasses with global environmental change. Nature Communications, 15(1).

Mestre-Tomás, J. (2026). GeoThinneR: An R Package for Efficient Spatial Thinning of Species Occurrences and Point Data. The R Journal, 18(1):299–314.

Mestre-Tomás, J., Fuster-Alonso, A., Bellido, J. M., and Coll, M. (2026). GLOSSA: A user-friendly R Shiny application for Bayesian machine learning analysis of marine species distribution. Methods in Ecology and Evolution, 17(2):505–517.

Meyer, D., Dimitriadou, E., Hornik, K., Weingessel, A., and Leisch, F. (2024). e1071: Misc Functions of the Department of Statistics, Probability Theory Group (Formerly: E1071), TU Wien. R package version 1.7-16.

Montalvo-Mancheno, C. S., Buettel, J. C., Ondei, S., and Brook, B. W. (2025). A reproducible, data-driven approach to mapping species distributions using presence-only data and biogeographic templates. Ecology and Evolution, 15(10).

Moore, N. A., Morales-Castilla, I., Hargreaves, A. L., Olalla-Tárraga, M. A., Villalobos, F., Calosi, P., Clusella-Trullas, S., Rubalcaba, J. G., Algar, A. C., Martínez, B., Rodríguez, L., Gravel, S., Bennett, J. M., Vega, G. C., Rahbek, C., Araújo, M. B., Bernhardt, J. R., and Sunday, J. M. (2023). Temperate species underfill their tropical thermal potentials on land. Nature Ecology and Evolution, 7(12):1993–2003.

Moudrý, V., Bazzichetto, M., Remelgado, R., Devillers, R., Lenoir, J., Mateo, R. G., Lembrechts, J. J., Sillero, N., Lecours, V., Cord, A. F., Barták, V., Balej, P., Rocchini, D., Torresani, M., Arenas-Castro, S., Man, M., Prajzlerová, D., Gdulová, K., Prošek, J., Marchetto, E., Zarzo-Arias, A., Gábor, L., Leroy, F., Martini, M., Malavasi, M., Cazzolla Gatti, R., Wild, J., and Šímová, P. (2024). Optimising occurrence data in species distribution models: sample size, positional uncertainty, and sampling bias matter. Ecography, 2024(12).

Nolan, V., Gilbert, F., Reed, T., and Reader, T. (2022). Distribution models calibrated with independent field data predict two million ancient and veteran trees in England. Ecological Applications, 32(8).

Oliver, R. Y., Meyer, C., Ranipeta, A., Winner, K., and Jetz, W. (2021). Global and national trends, gaps, and opportunities in documenting and monitoring species distributions. PLOS Biology, 19(8):e3001336.

Provoost, P. and Bosch, S. (2022). robis: Ocean Biodiversity Information System (OBIS) Client. R package version 2.11.3.

R Core Team (2025). R: A Language and Environment for Statistical Computing. R Foundation for Statistical Computing, Vienna, Austria.

Reddy, S. and Dávalos, L. M. (2003). Geographical sampling bias and its implications for conservation priorities in Africa. Journal of Biogeography, 30(11):1719–1727.

Sainz-Villegas, S., de la Hoz, C. F., Juanes, J. A., and Puente, A. (2022). Predicting non-native seaweeds global distributions: The importance of tuning individual algorithms in ensembles to obtain biologically meaningful results. Frontiers in Marine Science, 9.

Scott, D. W. and Terrell, G. R. (1987). Biased and unbiased cross-validation in density estimation. Journal of the American Statistical Association, 82(400):1131–1146.

Sudo, K. and Nakaoka, M. (2020). Fine-scale distribution of tropical seagrass beds in Southeast Asia. Ecological Research, 35(6):994–1000.

Suárez-Seoane, S., Jiménez-Alfaro, B., and Obeso, J. R. (2019). Habitat-partitioning improves regional distribution models in multi-habitat species: a case study with the European bilberry. Biodiversity and Conservation, 29(3):987–1008.

Tulloch, A. I. T. and Szabo, J. K. (2012). A behavioural ecology approach to understand volunteer surveying for citizen science datasets. Emu - Austral Ornithology, 112(4):313–325.

UNEP-WCMC, S. F. (2020). Global distribution of seagrasses (version 7.0). seventh update to the data layer used in Green and Short (2003). UN Environment World Conservation Monitoring Centre: Cambridge, UK. Available from http://data.unep-wcmc.org/datasets/7. Accessed on 15 June 2025.

Van Eupen, C., Maes, D., Herremans, M., Swinnen, K. R., Somers, B., and Luca, S. (2021). The impact of data quality filtering of opportunistic citizen science data on species distribution model performance. Ecological Modelling, 444:109453.

Vapnik, V. N. (1995). *The Nature of Statistical Learning Theory*. Springer New York.

Vaughan, D. (2024). slider: Sliding Window Functions. R package version 0.3.2.

Verbruggen, H., Tyberghein, L., Belton, G. S., Mineur, F., Jueterbock, A., Hoarau, G., Gurgel, C. F. D., and De Clerck, O. (2013). Improving transferability of introduced species’ distribution models: New tools to forecast the spread of a highly invasive seaweed. PLoS ONE, 8(6):e68337.

Wang, P., Zhang, L., Li, J., Yang, C., Xie, G., and Wang, W. (2025). Using ecological modeling to study the response of distribution dynamics of *Paraglenea fortunei* (Coleoptera: Cerambycidae) to human activities and climate change to in Northeast Asia. Ecology and Evolution, 15(7).

Wickham, H. (2016). *ggplot2: Elegant Graphics for Data Analysis*. Springer-Verlag New York.

Wickham, H., Averick, M., Bryan, J., Chang, W., McGowan, L. D., François, R., Grolemund, G., Hayes, A., Henry, L., Hester, J., Kuhn, M., Pedersen, T. L., Miller, E., Bache, S. M., Müller, K., Ooms, J., Robinson, D., Seidel, D. P., Spinu, V., Takahashi, K., Vaughan, D., Wilke, C., Woo, K., and Yutani, H. (2019). Welcome to the tidyverse. Journal of Open Source Software, 4(43):1686.

Xu, Q., Wang, X., Yi, J., and Wang, Y. (2024). Bias correction in species distribution models based on geographic and environmental characteristics. Ecological Informatics, 81:102604.

Yates, K. L., Bouchet, P. J., Caley, M. J., Mengersen, K., Randin, C. F., Parnell, S., Fielding, A. H., Bamford, A. J., Ban, S., Barbosa, A. M., Dormann, C. F., Elith, J., Embling, C. B., Ervin, G. N., Fisher, R., Gould, S., Graf, R. F., Gregr, E. J., Halpin, P. N., Heikkinen, R. K., Heinanen, S., Jones, A. R., Krishnakumar, P. K., Lauria, V., Lozano-Montes, H., Mannocci, L., Mellin, C., Mesgaran, M. B., Moreno-Amat, E., Mormede, S., Novaczek, E., Oppel, S., Ortuño Crespo, G., Peterson, A. T., Rapacciuolo, G., Roberts, J. J., Ross, R. E., Scales, K. L., Schoeman, D., Snelgrove, P., Sundblad, G., Thuiller, W., Torres, L. G., Verbruggen, H., Wang, L., Wenger, S., Whittingham, M. J., Zharikov, Y., Zurell, D., and Sequeira, A. M. (2018). Outstanding challenges in the transferability of ecological models. Trends in Ecology and Evolution, 33(10):790–802.

Zizka, A., Antonelli, A., and Silvestro, D. (2020). sampbias, a method for quantifying geographic sampling biases in species distribution data. Ecography, 44(1):25–32.

Zizka, A., Silvestro, D., Andermann, T., Azevedo, J., Duarte Ritter, C., Edler, D., Farooq, H., Herdean, A., Ariza, M., Scharn, R., Svantesson, S., Wengström, N., Zizka, V., and Antonelli, A. (2019). Coordinatecleaner: Standardized cleaning of occurrence records from biological collection databases. Methods in Ecology and Evolution, 10(5):744–751.

Zárrate Charry, D. A., González-Maya, J. F., Arias-Alzate, A., Jiménez-Alvarado, J. S., Reyes Arias, J. D., Armenteras, D., and Betts, M. G. (2022). Connectivity conservation at the crossroads: protected areas versus payments for ecosystem services in conserving connectivity for Colombian carnivores. Royal Society Open Science, 9(1).

## REFERENCES

Assis, J., Fragkopoulou, E., Frade, D., Neiva, J., Oliveira, A., Abecasis, D., Faugeron, S., and Serrão, E. A. (2020). A fine-tuned global distribution dataset of marine forests. Scientific Data, 7(1).

Chapman, A. and Wieczorek, J. (2020). Georeferencing Best Practices.

Hijmans, R. J., Brown, A., and Barbosa, M. (2026). *terra: Spatial Data Analysis*. R package version 1.8-95.

McKenzie, L. J. and Yoshida, R. L. (2020). Over a decade monitoring fiji’s seagrass condition demonstrates resilience to anthropogenic pressures and extreme climate events. Marine Pollution Bulletin, 160:111636.

Pierre, L. and Louis, L. (2012). Numerical ecology. Elsevier, Amsterdam, 3rd edition.

Short, F. T. and Green, E. P. (2003). World Atlas of Seagrasses. University of California Press.

Spalding, M. D., Fox, H. E., Allen, G. R., Davidson, N., Ferdaña, Z. A., Finlayson, M., Halpern, B. S., Jorge, M. A., Lombana, A., Lourie, S. A., Martin, K. D., McManus, E., Molnar, J., Recchia, C. A., and Robertson, J. (2007). Marine ecoregions of the world: A bioregionalization of coastal and shelf areas. BioScience, 57(7):573–583.

Vapnik, V. N. (1995). *The Nature of Statistical Learning Theory*. Springer New York.

Vaughan, D. (2024). slider: Sliding Window Functions. R package version 0.3.2.

## REFERENCES

Abdulrahman, S., Aguirre, A., Arriaga, E., Avina, A. C., Badajos, A., Badillo, Y. A., Bañuelos, A. M., Pulido, J., Bucio Valdovinos, B.-s., Champaco, T., Chavez, R., Chavez-Felix, G., Cruz, M., Distancia, M., Dzubay, K., Espinoza, A., Estrada, J., Fernandez, D., Gutierrez-Villanueva, A., … Valdez, H. (2025). Complete chloroplast genome of the marine eelgrass *Zostera pacifica* (*Zosteraceae*, *Plantae*) from Monterey, California (L. Pritchard, Ed.). Microbiology Resource Announcements, 14(2). 10.1128/mra.01189-24

Assis, J., Fernández Bejarano, S. J., Salazar, V. W., Schepers, L., Gouvêa, L., Fragkopoulou, E., Leclercq, F., Vanhoorne, B., Tyberghein, L., Serrão, E. A., Verbruggen, H., & De Clerck, O. (2024). Bio-oracle v3.0. pushing marine data layers to the cmip6 earth system models of climate change research. Global Ecology and Biogeography, 33(4). 10.1111/geb.13813

Bertelli, C. M., Stokes, H. J., Bull, J. C., & Unsworth, R. K. F. (2022). The use of habitat suitability modelling for seagrass: A review. Frontiers in Marine Science, 9. 10.3389/fmars.2022.997831

Campey, M., Waycott, M., & Kendrick, G. (2000). Re-evaluating species boundaries among members of the *Posidonia ostenfeldii* species complex (*Posidoniaceae*) – morphological and genetic variation. Aquatic Botany, 66(1), 41–56. 10.1016/s0304-3770(99)00015-7

Coyer, J. A., Hoarau, G., Kuo, J., Tronholm, A., Veldsink, J., & Olsen, J. L. (2013). Phylogeny and temporal divergence of the seagrass family *Zosteraceae* using one nuclear and three chloroplast loci. Systematics and Biodiversity, 11(3), 271–284. 10.1080/14772000.2013.821187

Creed, J. C., & Samper-Villarreal, J. (2019). Clarification of the nomenclature of the seagrass *Halophila baillonii* Ascherson. Aquatic Botany, 154, 42–44. 10.1016/j.aquabot.2019.01.002

den Hartog, C., & Triest, L. (2020). A profound view and discourse on the typification and status of three confused taxa: *Ruppia maritima*, *R. spiralis* and *R. cirrhosa*. Botanica Marina, 63(3), 229–239. 10.1515/bot-2019-0045

Green, E., & Short, F. (2003). World Atlas of Seagrasses. Prepared by the UNEP World Conservation Monitoring Center. University of California Press.

Guiry, M., & Guiry, G. (2025). Algaebase. World-wide electronic publication, National University of Ireland, Galway [Accessed: 04 June 2025]. https://www.algaebase.org/

Hartog, C. d., & Kuo, J. (2007). Taxonomy and biogeography of seagrasses. In A. W. Larkum, R. J. Orth, & C. M. Duarte (Eds.), Seagrasses: Biology, ecology and conservation. Springer. 10.1007/978-1-4020-2983-7 1

Jacobs, S., Les, D., & Moody, M. (2006). New combinations in Australasian *Zostera* (*Zosteraceae*). Telopea, 11, 127–128.

Kuo, J. (2020). Taxonomy of the genus *Halophila* Thouars (*Hydocharitaceae*): A review. Plants, 9(12), 1732. 10.3390/plants9121732

Liu, S. Y. V., Kumara, T. P., & Hsu, C.-H. (2020). Genetic identification and hybridization in the seagrass genus *Halophila* (*Hydrocharitaceae*) in Sri Lankan waters. PeerJ, 8, e10027. 10.7717/peerj.10027

Mestre-Tomás, J., Fuster-Alonso, A., Bellido, J. M., & Coll, M. (2026). GLOSSA: A user-friendly R Shiny application for Bayesian machine learning analysis of marine species distribution. Methods in Ecology and Evolution, 17(2), 505–517. 10.1111/2041-210x.70167

POWO. (2025). Plants of the World Online [Facilitated by the Royal Botanic Gardens, Kew. Published on the Internet; http://www.plantsoftheworldonline.org/. Accessed: March 13, 2026]. http://www.plantsoftheworldonline.org/

Sullivan, B. K., & Short, F. T. (2023). Taxonomic revisions in *Zosteraceae* (*Zostera*, *Nanozostera*, Heterozostera and Phyl-lospadix). Aquatic Botany, 187, 103636. 10.1016/j.aquabot.2023.103636

Tomlinson, P. B., & Posluzny, U. (2001). Generic limits in the seagrass family *Zosteraceae*. TAXON, 50(2), 429–437. 10.2307/1223890

WFO. (2025). World Flora Online Consortium [Published on the Internet; http://www.worldfloraonline.org/. Accessed: March 13, 2026]. http://www.worldfloraonline.org/

Zhang, X., Yu, L., Li, Y.-L., Suonan, Z., Komatsu, T., Qiu, G., Xu, S., Yue, S., Xu, M., Wang, F., Zhang, Y., Lee, K.-S., Liu, J.-X., Zhou, Y., & Reusch, T. B. H. (2025). Uncovering the *Nanozostera japonica* species complex suggests cryptic speciation and underestimated seagrass diversity. New Phytologist, 247(5), 2086–2097. 10.1111/nph.70355

