## Supplementary materials 1-8 for "Stop guessing thinning distances: a data-driven framework for spatial thinning parametrization for ecological modelling"

DRAFT

**Table 1.** Taxonomic entities queried for the data download from GBIF and OBIS. Taxon keys were used for download from GBIF.

| Species | taxonKey | Species | taxonKey |
|---|---|---|---|
| *Amphibolis antarctica* | 2864119 | *Halophila tricostata* | 5329596 |
| *Amphibolis griffithii* | 2864116 | *Thalassia hemprichii* | 5329593 |
| *Cymodocea angustata* | 5328498 | *Thalassia testudinum* | 5329591 |
| *Cymodocea nodosa* | 5328492 | *Posidonia angustifolia* | 5328672 |
| *Cymodocea rotundata* | 5328491 | *Posidonia australis* | 5328658 |
| *Halodule bermudensis* | 2864093 | *Posidonia denhartogii* | 5328670 |
| *Halodule ciliata* | 2864114 | *Posidonia kirkmanii* | 5328675 |
| *Halodule emarginata* | 2864100 | *Posidonia oceanica* | 5328663 |
| *Halodule pinifolia* | 2864112 | *Posidonia ostenfeldii* | 5328662 |
| *Halodule uninervis* | 2864102 | *Posidonia sinuosa* | 5328674 |
| *Halodule wrightii* | 2864094 | *Althenia marina* | 9525100 |
| *Oceana serrulata* | 10918187 | *Heterozostera nigricaulis* | 5792280 |
| *Syringodium filiforme* | 7982798 | *Heterozostera polychlamys* | 5792279 |
| *Syringodium isoetifolium* | 2864078 | *Heterozostera tasmanica* | 2863949 |
| *Thalassodendron ciliatum* | 5328510 | *Phyllospadix iwatensis* | 2863956 |
| *Thalassodendron leptocaule* | 7850647 | *Phyllospadix japonicus* | 2863953 |
| *Thalassodendron pachyrhizum* | 5328509 | *Phyllospadix scouleri* | 2863958 |
| *Enhalus acoroides* | 2866113 | *Phyllospadix serrulatus* | 2863957 |
| *Halophila australis* | 5329597 | *Phyllospadix torreyi* | 2863954 |
| *Halophila baillonii* | 9179530 | *Nanozostera capensis* | 2863920 |
| *Halophila beccarii* | 5329636 | *Nanozostera capricorni* | 2863924 |
| *Halophila capricorni* | 5329617 | *Nanozostera japonica* | 2863943 |
| *Halophila decipiens* | 5329611 | *Nanozostera mucronata* | 2863927 |
| *Halophila engelmannii* | 5329618 | *Nanozostera muelleri* | 2863922 |
| *Halophila hawaiiana* | 5329604 | *Nanozostera noltei* | 2863938 |
| *Halophila major* | 5329619 | *Nanozostera novazelandica* | 2863917 |
| *Halophila mikii* | 5329638 | *Zostera asiatica* | 2863965 |
| *Halophila minor* | 5329606 | *Zostera caespitosa* | 2863964 |
| *Halophila nipponica* | 5329603 | *Zostera caulescens* | 2863966 |
| *Halophila okinawensis* | 5329616 | *Zostera marina* | 2863967 |
| *Halophila ovalis* | 5329622 | *Ruppia cirrhosa* | 2863987 |
| *Halophila spinulosa* | 5329609 | *Ruppia maritima* | 2864004 |
| *Halophila stipulacea* | 5329631 | *Ruppia megacarpa* | 2863986 |
| *Halophila sulawesii* | 5329595 | *Ruppia tuberosa* | 2864003 |

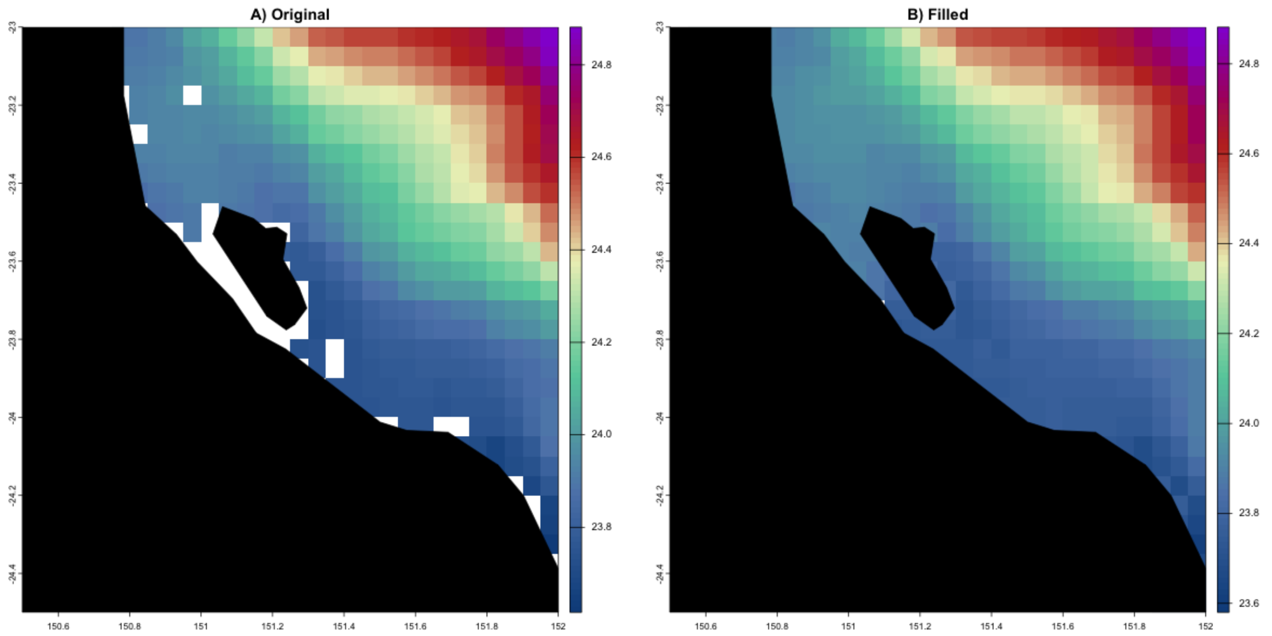


**Figure 1.** Comparison of the original and filled raster, using the SST mean (2000-2010) from BIO-Oracle v3. The region proximal to Gladstone (Australia) is shown, because of the many *Halophila* and *Nanozostera* observations potentially lost, if this extension is not computed.

Gap filling was performed independently for each decadal SST raster using a focal neighbourhood approach implemented in the Hijmans et al. (2026) package in R. A circular moving window was defined with a diameter of four raster cells (corresponding to 0.2°, given the BIO-ORACLE v3 spatial resolution of 0.05°). The circular neighbourhood was constructed as follows:

```
Wcirc <- focalMat(envstack, d = 4 * res(envstack)[1], type = "circle")
```

This step generated a kernel in which all cells whose Euclidean distance from the centre was less than or equal to the specified diameter were included in the neighbourhood. To ensure an equal contribution of all neighbouring cells, the kernel was converted to a uniform (unweighted) window:

```
Wcirc[Wcirc > 0] <- 1
```

Missing raster values were then filled by computing the arithmetic mean of neighbouring cells within the circular window. The operation was applied only to raster cells with missing values (NA), leaving all pre-existing values unchanged and avoiding an artificial smoothing of the original data:

```
filledraster <- focal(envstack, w = Wcirc, fun = mean, na.policy = "only", na.rm = TRUE)
```

The ’na.policy = "only"’ option restricted calculations to focal cells with missing values. ’*na.rm = TRUE*’ excluded missing neighbours from the computation. If all neighbouring cells within the window were missing, the focal cell remained empty. This approach increased spatial coverage at the raster boundaries (up to ~2.3 km from the original edge) while minimizing interpolation artifacts.

Occurrence points were converted to a SpatVector in the same coordinate reference system as the BIO-Oracle v3 rasters (EPSG:4326). The local SST at each point was extracted with the *’terra::extract’* function, through bilinear interpolation. The

**Table 2.** Increment in filled cells (compared to the original number of filled cells) and occurrence points with a successful temperature extraction as focal window size is increased

| | d = 2 cells | d = 4 cells | d = 6 cells | d = 8 cells |
|---|---|---|---|---|
| Increment in cells | 101.50% | 102.66% | 103.55% | 104.54% |
| Extraction success | 95.88% | 99.88% | 99.99% | 99.99% |

$$D_s = 2\sqrt{\frac{A_s}{\pi}} \tag{1}$$

where $D_s$ is the diameter for species $s$, $A_s$ represents the occupied area (km$^2$) estimated from the rasterised occurrence cells, and the other elements are derived from the circle area formula $A = \pi r^2$, which can be simplified into $r = \sqrt{\frac{A}{\pi}}$ to calculate the radius, which, by definition will have half the value of $D_s$.

*Resolution selection*

Raster resolution was selected arbitrarily as a compromise between excessive range fragmentation at very fine resolutions and artificial range inflation at coarse resolutions, particularly into non-informative oceanic or terrestrial areas. Six resolutions (0.1°, 0.3°, 0.5°, 1°, 2.5°, and 5°) were evaluated by adapting the distribution-size calculation workflow to iterate across resolutions. For each resolution, species were ranked according to their estimated range size ($D_{s,res}$). As a null reference, an additional ranking was generated by randomizing species order using a fixed random seed ('*set.seed(12)*') for replicability.

The ranked species lists were compared with Kendall's rank correlations, using the numerical rank position of the species in each resolution according to its predicted range size. Pairwise comparisons between ranked lists were conducted using Kendall's rank correlation coefficient ($\tau$), quantifying the degree of agreement in the relative ordering of species. Correlation values were calculated for all pairs of resolutions and visualized as a correlation matrix (Fig. 2). The correlation analysis showed a high level of consistency across resolutions. Even when comparing the finest resolution (res_0.1) with a resolution 25 times coarser (res_2.5), the correlation remained high ($\tau = 0.84$).

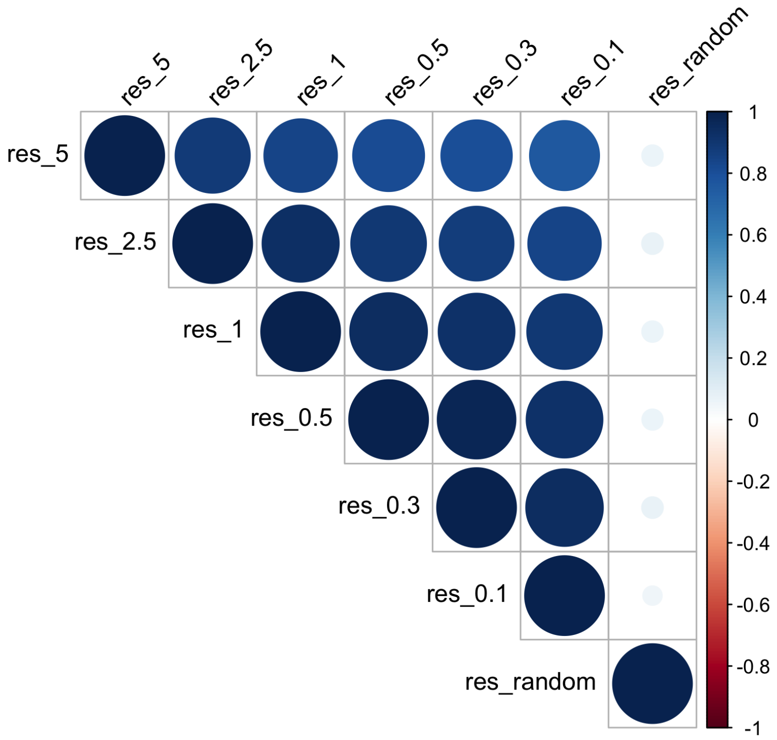


**Figure 2.** Kendall's rank correlation matrix comparing species rankings derived from predicted range sizes at each original spatial resolution. Species were ranked according to the resolutions employed (0.1, 0.3, 0.5, 1, 2.5, and 5) and a random species order

##### 1.3 Thinned datasets

From the original raw dataset, 13 datasets were independently generated by stratified thinning with Mestre-Tomás (2026). Thinning was iterated over species. Five trials were conducted per iteration, from which only the most conservative (more points retained) was kept. The original dataset was thinned with each item in the following parameter set $\Theta$:

$$\Theta = \{\theta_1, \theta_2, \ldots, \theta_k\} = \{0.001, 0.01, 0.1, 1, 2.5, 5, 7.5, 10, 12.5, 15, 17.5\} \quad (2)$$

The parameter set was generated by including extremely small and big parameters for a better calibration.

##### 1.4 Calculation of niche stabilization

```
    stabilize_sp_df <- function(sp, df, B, nsamp, replacem) {
  # a) Select one species per iteration
    sp_df <- df %>%
     filter(species == sp) %>%
     pull(SST_mean)

  # b) Avoid species with evidently not enough data
  if (nrow(sp_df) < 3)
    return(tibble())

  # c) repeat for B replicates + calculate moving median
  map_dfr(seq_len(B), function(b) {

    m    <- min(nsamp, nrow(sp_df))
    idx <- sample.int(nrow(sp_df), size = m, replace = replacem)
    x <- sp_df[idx]

    tibble(
      repl = b,
      n_obs = seq_along(x),
      mov_med = slide_dbl(x, ~ median(.x, na.rm = TRUE), .before = Inf)
    ) %>%
      mutate(mov_med = ifelse(n_obs < 2, NA_real_, mov_med))
  })
}
```

```
    compute_threshold <- function(summary_df) {

  # a) if less than 10 counts -> NA
  if (nrow(summary_df) < 10) {
    return(list(
      n_obs_stable = NA_integer_,
      p_stable = NA_real_,
      final_mov_median = NA_real_,
      width_at_stable = NA_real_,
      stability_reached = FALSE
    ))
  }
# b) determine the number of observations used for that theta, s pair
  n_max <- max(summary_df$n_obs)

  # c) calculate the moving median tail
  tail_n <- min(20, nrow(summary_df))
  final_mov_median <- mean(tail(summary_df$me_mov, tail_n))
# d) summary for stability flag
  summary_df <- summary_df %>%
    mutate(
      diff_final = abs(me_mov - final_mov_median),
      width = CI_high - CI_low,
      stable_flag = (diff_final <= tol_value) & (width <= tol_width)
    )

  # e) find first run of TRUE in 'length >= window_run'

  run_seq <- rle(summary_df$stable_flag)
  idx_end <- cumsum(run_seq$lengths)


  #f) identify runs of TRUE with sufficient length
  ok <- which(run_seq$values & run_seq$lengths >= window_run)

  #g) Check if suitable runs exist
  if (length(ok) == 0) {
    return(list(
      ST_p_s = NA_integer_,
      p_stable = NA_real_,
```

DRAFT

```
      final_mov_median = final_mov_median,
      width_at_stable = NA_real_,
      stability_reached = FALSE
    ))
  }

  #h) extract first suitable index of the run
  start_idx <- idx_start[ok[1]]
  n_obs_stable <- summary_df$n_obs[start_idx]
  width_at_stable <- summary_df$width[start_idx]

  # i) final results
  list(
    ST_p_s = n_obs_stable,
    p_stable = n_obs_stable / n_max,
    final_mov_median = final_mov_median,
    width_at_stable = width_at_stable,
    stability_reached = TRUE
  )
}
```

*Stabilization* The stabilization run, where the previous functions were computed, has three sequential parts: (i) replicate computation (replicated moving median calculation), (ii) thresholding, and (iii) replicate aggregation. At the end of the stabilization run, the third input component for SVM, the boolean stability, was generated.

First, moving median trajectories were calculated independently for each species across all thinned datasets (each $s, \theta$ pair) using nested loops. Lookup vectors were created to later order the outputs, considering the range size ($D_s$) and raw occurrence counts. To compute and store the results of each $(s, \theta)$ pair two hierarchical loops were run: an outer loop for the species and an inner loop for the thinned datasets. The inner loop was run for each thinned dataset $\theta$, filtering out $(s, \theta)$ pairs with insufficient data. The thinning parameter value was extracted from the dataset name with the custom function 'i' (`parse_param()`). The custom function 'ii' (`stabilize_sp_df`) was used to calculate replicate calculations of the moving median for this $(s, \theta)$ pair. Important metadata was added to the outputs (e.g. species name, thinning parameter). Results of the replicate trajectories of each species $s, \theta$ pair were compiled in a list.

DRAFT

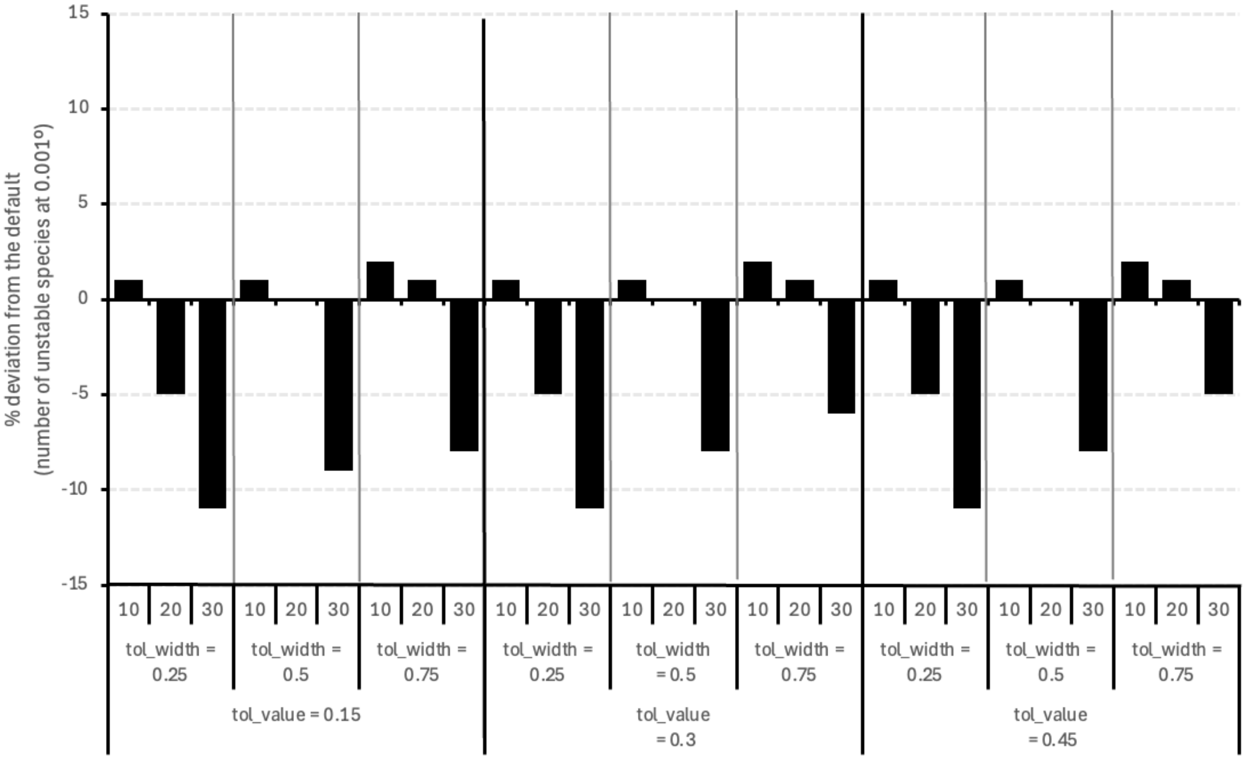


**Figure 3.** Sensitivity analysis for 27 combinations of the tolerance settings. Combinations were compared by normalising the percentage of species that never reach stabilization and reporting the changes in this number.

##### 1.6 Biogeographic-size dependent thinning parametrization (BD-TP)

A data frame with the variables calculated in previous steps and used as the SVM input. Then, a Support Vector Machines algorithm (Vapnik, 1995) is used to extrapolate hyperplane values. These values represent the highest thinning parameter that can be used prior to niche characterization destabilization. The components of a simple linear model were computed analytically to enunciate a linear equation in which $D_s$ is used to predict $\theta_{opt}$ ('$\theta = m \cdot D_s - C$').

**Application of Support Vector Machines**

A data frame was built with previous outputs: the parameters in numeric format, $D_s$, and a boolean stability variable. With this input, the SVM model was built with `svm()` from the `e1071` package (Meyer et al., 2024). The range size and the parameter were included as predictors and stability as the class label '`svm(stab ~ diameter_km + parameter_num, ...)`', with a linear kernel. The '`probability = TRUE`' and '`scale = TRUE`' arguments were used to allow the later extraction of probabilities and scaling features internally, respectively.

With the SVM output two prediction grids were generated. The first, created a fine grid of points covering the range of the predictors ($D_s$ and $\theta$) to later visualize the decision boundary. With the '`predict()`' function, a categorical value is created to give the predicted class for each grid cell. The second, predicts decision values, that is, how far each point is from the decision boundary.

The SVM was represented as a ggplot graph, using the '`scale_fill_gradient2`' argument to fill the plot, '`geom_contour`' to plot the 0 and the $\pm 1$ boundaries, and '`geom_point`' to represent the stability for each $s, \theta$ pair. The order of the axes (y = $D_s$, x = $\theta$) is plotted inversely to the mathematical formulation of the linear model ($\theta = f(D_s)$; $\theta = m \cdot D_s + C$) for conceptual clarity.

**Linear model extraction of the hyperplane**

We extracted the hyperplane analytically from the SVM model coefficients. The weight vector of the decision function was first reconstructed from the support-vector coefficients and support vectors, and the intercept was obtained from the model parameter ($\rho$). Because the SVM was fitted with standardized predictors, The predictor-specific centering and scaling factors used internally by the SVM were then retrieved and used to transform the hyperplane coefficients back to the original measurement units. In original units, the decision boundary can be written as:

$$a_0 D_s + a_1 \theta + a_c = 0,$$

DRAFT

where ($D_s$) is the range-size descriptor and ($\theta$) is the thinning parameter. Rearranging this expression for ($\theta$) yielded the linear BD-TP rule ($\theta = mD_s + c_0$), with ($m = -a_0/a_1$) and ($c_0 = -a_c/a_1$) and with $D_s$ and $\theta$ as the independent and dependent variables, respectively. The resulting slope and intercept were retained as the analytical estimates of the SVM decision boundary and were used to parameterise the BD-TP rule.

A new data frame with the species list and their $\theta_{opt}$ was created with $\theta_{opt}$ = `pmax(m * diameter_km +c0, 0.001)`. Through this formula, the parameters are predicted regarding the relationship with the range size, but are capped in 0.001º as a minimal parameter. This cap avoids the application of theoretical negative parameters. It also ensures that, extremely close occurrence records are not are considered, even if the predicted parameter is lower than 0.001.

DRAFT

**Table 1.** Curated list of seagrass taxa considered, including the taxonomic references that validate their nomenclature and taxonomic status. Comments expand discrepancies across references

| Family | Species | Taxonomic reference |
|---|---|---|
| *Cymodoceaceae* | *Amphibolis antarctica* (Labill.) Asch. | (Guiry & Guiry, 2025; POWO, 2025; WFO, 2025) |
| *Cymodoceaceae* | *Amphibolis griffithii* (J.M.Black) Hartog | (Guiry & Guiry, 2025; POWO, 2025; WFO, 2025) |
| *Cymodoceaceae* | *Cymodocea angustata* Ostenfeld 1916 | (Guiry & Guiry, 2025; POWO, 2025; WFO, 2025) |
| *Cymodoceaceae* | *Cymodocea nodosa* (Ucria) Asch. | (Guiry & Guiry, 2025; POWO, 2025; WFO, 2025) |
| *Cymodoceaceae* | *Cymodocea rotundata* Asch. & Schweinf. | (Guiry & Guiry, 2025; POWO, 2025; WFO, 2025) |
| *Cymodoceaceae* | *Halodule bermudensis* Hartog | (Guiry & Guiry, 2025; POWO, 2025; WFO, 2025) |
| *Cymodoceaceae* | *Halodule emarginata* Hartog | (Guiry & Guiry, 2025; POWO, 2025; WFO, 2025) |
| *Cymodoceaceae* | *Halodule pinifolia* (Miki) Hartog | (Guiry & Guiry, 2025; POWO, 2025; WFO, 2025) |
| *Cymodoceaceae* | *Halodule uninervis* (Forssk.) Boiss. | (Guiry & Guiry, 2025; POWO, 2025; WFO, 2025) |
| *Cymodoceaceae* | *Halodule wrightii* Asch. | (Guiry & Guiry, 2025; POWO, 2025; WFO, 2025) |
| *Cymodoceaceae* | *Oceana serrulata* (R.Br.) Byng & Christenh. | (Guiry & Guiry, 2025; POWO, 2025; WFO, 2025) |
| *Cymodoceaceae* | *Syringodium filiforme* Kütz. | (Guiry & Guiry, 2025; POWO, 2025; WFO, 2025) |
| *Cymodoceaceae* | *Syringodium isoetifolium* (Asch.) Dandy | (Guiry & Guiry, 2025; POWO, 2025; WFO, 2025) |
| *Cymodoceaceae* | *Thalassodendron ciliatum* complex | 1 |
| *Cymodoceaceae* | *Thalassodendron pachyrhizum* Hartog | (Guiry & Guiry, 2025; POWO, 2025; WFO, 2025) |
| *Hydrocharitaceae* | *Enhalus acoroides* (L.f.) Royle | (Guiry & Guiry, 2025; POWO, 2025; WFO, 2025) |
| *Hydrocharitaceae* | *Halophila australis* Doty & B.C.Stone | (Guiry & Guiry, 2025; POWO, 2025; WFO, 2025) |
| *Hydrocharitaceae* | *Halophila baillonii* Asch. | (Creed & Samper-Villarreal, 2019) |
| *Hydrocharitaceae* | *Halophila beccarii* Asch. | (Guiry & Guiry, 2025; POWO, 2025; WFO, 2025) |
| *Hydrocharitaceae* | *Halophila capricorni* Larkum | (Guiry & Guiry, 2025; POWO, 2025; WFO, 2025) |
| *Hydrocharitaceae* | *Halophila decipiens* Ostenf. | (Guiry & Guiry, 2025; POWO, 2025; WFO, 2025) |
| *Hydrocharitaceae* | *Halophila engelmannii* Asch. | (Guiry & Guiry, 2025; POWO, 2025; WFO, 2025) |
| *Hydrocharitaceae* | *Halophila hawaiiana* Doty & B.C.Stone | (Guiry & Guiry, 2025; POWO, 2025; WFO, 2025) |
| *Hydrocharitaceae* | *Halophila ovalis* group | 2 |
| *Hydrocharitaceae* | *Halophila spinulosa* (R.Br.) Asch. | (Guiry & Guiry, 2025; POWO, 2025; WFO, 2025) |
| *Hydrocharitaceae* | *Halophila stipulacea* (Forssk.) Asch. | (Guiry & Guiry, 2025; POWO, 2025; WFO, 2025) |
| *Hydrocharitaceae* | *Halophila sulawesii* J.Kuo | (Guiry & Guiry, 2025; POWO, 2025; WFO, 2025) |
| *Hydrocharitaceae* | *Halophila tricostata* M.Greenway | (Guiry & Guiry, 2025; POWO, 2025; WFO, 2025) |
| *Hydrocharitaceae* | *Thalassia hemprichii* (Ehrenb.) Asch. | (Guiry & Guiry, 2025; POWO, 2025; WFO, 2025) |
| *Hydrocharitaceae* | *Thalassia testudinum* Banks & Sol. ex K.D.Koenig | (Guiry & Guiry, 2025; POWO, 2025; WFO, 2025) |
| *Posidoniaceae* | *Posidonia angustifolia* Cambridge & J.Kuo | (Guiry & Guiry, 2025; POWO, 2025; WFO, 2025) |
| *Posidoniaceae* | *Posidonia australis* J.D.Hooker | (Guiry & Guiry, 2025; POWO, 2025; WFO, 2025) |
| *Posidoniaceae* | *Posidonia oceanica* (L.) Delile | (Guiry & Guiry, 2025; POWO, 2025; WFO, 2025) |
| *Posidoniaceae* | *Posidonia ostenfeldii* complex | 3 |
| *Posidoniaceae* | *Posidonia sinuosa* Cambridge & J.Kuo | (Guiry & Guiry, 2025; POWO, 2025; WFO, 2025) |
| *Potamogetonaceae* | *Althenia marina* (E.L.Robertson) Yu Ito | (Guiry & Guiry, 2025; POWO, 2025; WFO, 2025) |
| *Zosteraceae* | *Heterozostera nigricaulis* J.Kuo | (Coyer et al., 2013; Guiry & Guiry, 2025; Sullivan & Short, 2023) |
| *Zosteraceae* | *Heterozostera polychlamys* J.Kuo | (Coyer et al., 2013; Guiry & Guiry, 2025; Sullivan & Short, 2023) |

DRAFT

**Table 1.** Curated list of seagrass taxa considered, including the taxonomic references that validate their nomenclature and taxonomic status. Comments expand discrepancies across references

| Family | Species | Taxonomic reference |
|---|---|---|
| *Zosteraceae* | *Heterozostera tasmanica* (Asch.) Hartog | (Coyer et al., 2013; Guiry & Guiry, 2025; Sullivan & Short, 2023) |
| *Zosteraceae* | *Phyllospadix iwatensis* Makino | (Guiry & Guiry, 2025; POWO, 2025; Sullivan & Short, 2023; WFO, 2025) |
| *Zosteraceae* | *Phyllospadix japonicus* Makino | (Guiry & Guiry, 2025; POWO, 2025; Sullivan & Short, 2023; WFO, 2025) |
| *Zosteraceae* | *Phyllospadix scouleri* Hook. | (Guiry & Guiry, 2025; POWO, 2025; Sullivan & Short, 2023; WFO, 2025) |
| *Zosteraceae* | *Phyllospadix serrulatus* Rupr. ex Asch. | (Guiry & Guiry, 2025; POWO, 2025; Sullivan & Short, 2023; WFO, 2025) |
| *Zosteraceae* | *Phyllospadix torreyi* S.Watson | (Guiry & Guiry, 2025; POWO, 2025; Sullivan & Short, 2023; WFO, 2025) |
| *Zosteraceae* | *Nanozostera capensis* (Setch.) Toml. & Posl. | (Guiry & Guiry, 2025; Sullivan & Short, 2023) |
| *Zosteraceae* | *Nanozostera japonica* (Asch. & Graebn.) Toml. & Posl. | 4 |
| *Zosteraceae* | *Nanozostera muelleri* complex | 5 |
| *Zosteraceae* | *Nanozostera noltii* (Hornem.) Toml. & Posl. | (Sullivan & Short, 2023) 6 |
| *Zosteraceae* | *Zostera asiatica* Miki | (Guiry & Guiry, 2025; POWO, 2025; WFO, 2025) |
| *Zosteraceae* | *Zostera caespitosa* Miki | (Guiry & Guiry, 2025; POWO, 2025; WFO, 2025) |
| *Zosteraceae* | *Zostera caulescens* Miki | (Guiry & Guiry, 2025; POWO, 2025; WFO, 2025) |
| *Zosteraceae* | *Zostera pacifica* S.Watson | (Abdulrahman et al., 2025; Guiry & Guiry, 2025; POWO, 2025) |
| *Zosteraceae* | *Zostera marina* L. | (Abdulrahman et al., 2025; POWO, 2025) |
| *Ruppiaceae* | *Ruppia maritima* group | 7 |
| *Ruppiaceae* | *Ruppia megacarpa* R.Mason | (Guiry & Guiry, 2025; POWO, 2025; WFO, 2025) |
| *Ruppiaceae* | *Ruppia tuberosa* J.S.Davis & Toml. | (Guiry & Guiry, 2025; POWO, 2025; WFO, 2025) |

Bertelli, C. M., Stokes, H. J., Bull, J. C., & Unsworth, R. K. F. (2022). The use of habitat suitability modelling for seagrass: A review. *Frontiers in Marine Science*, *9*. https://doi.org/10.3389/fmars.2022.997831

Campey, M., Waycott, M., & Kendrick, G. (2000). Re-evaluating species boundaries among members of the *Posidonia ostenfeldii* species complex (*Posidoniaceae*) – morphological and genetic variation. *Aquatic Botany*, *66*(1), 41–56. https://doi.org/10.1016/s0304-3770(99)00015-7

Coyer, J. A., Hoarau, G., Kuo, J., Tronholm, A., Veldsink, J., & Olsen, J. L. (2013). Phylogeny and temporal divergence of the seagrass family *Zosteraceae* using one nuclear and three chloroplast loci. *Systematics and Biodiversity*, *11*(3), 271–284. https://doi.org/10.1080/14772000.2013.821187

Creed, J. C., & Samper-Villarreal, J. (2019). Clarification of the nomenclature of the seagrass *Halophila baillonii* Ascherson. *Aquatic Botany*, *154*, 42–44. https://doi.org/10.1016/j.aquabot.2019.01.002

den Hartog, C., & Triest, L. (2020). A profound view and discourse on the typification and status of three confused taxa: *Ruppia maritima*, *R. spiralis* and *R. cirrhosa*. *Botanica Marina*, *63*(3), 229–239. https://doi.org/10.1515/bot-2019-0045


DRAFT

Green, E., & Short, F. (2003). *World Atlas of Seagrasses. Prepared by the UNEP World Conservation Monitoring Center*. University of California Press.

Guiry, M., & Guiry, G. (2025). Algaebase. World-wide electronic publication, National University of Ireland, Galway [Accessed: 04 June 2025]. https://www.algaebase.org/

Hartog, C. d., & Kuo, J. (2007). Taxonomy and biogeography of seagrasses. In A. W. Larkum, R. J. Orth, & C. M. Duarte (Eds.), *Seagrasses: Biology, ecology and conservation*. Springer. https://doi.org/10.1007/978-1-4020-2983-7_1

Jacobs, S., Les, D., & Moody, M. (2006). New combinations in Australasian *Zostera* (*Zosteraceae*). *Telopea*, *11*, 127–128.

Kuo, J. (2020). Taxonomy of the genus *Halophila* Thouars (*Hydocharitaceae*): A review. *Plants*, *9*(12), 1732. https://doi.org/10.3390/plants9121732

Liu, S. Y. V., Kumara, T. P., & Hsu, C.-H. (2020). Genetic identification and hybridization in the seagrass genus *Halophila* (*Hydrocharitaceae*) in Sri Lankan waters. *PeerJ*, *8*, e10027. https://doi.org/10.7717/peerj.10027

Mestre-Tomás, J., Fuster-Alonso, A., Bellido, J. M., & Coll, M. (2026). GLOSSA: A user-friendly R Shiny application for Bayesian machine learning analysis of marine species distribution. *Methods in Ecology and Evolution*, *17*(2), 505–517. https://doi.org/10.1111/2041-210x.70167

POWO. (2025). Plants of the World Online [Facilitated by the Royal Botanic Gardens, Kew. Published on the Internet; http://www.plantsoftheworldonline.org/. Accessed: March 13, 2026]. http://www.plantsoftheworldonline.org/

Sullivan, B. K., & Short, F. T. (2023). Taxonomic revisions in *Zosteraceae* (*Zostera*, *Nanozostera*, *Heterozostera* and *Phyllospadix*). *Aquatic Botany*, *187*, 103636. https://doi.org/10.1016/j.aquabot.2023.103636

Tomlinson, P. B., & Posluzny, U. (2001). Generic limits in the seagrass family *Zosteraceae*. *TAXON*, *50*(2), 429–437. https://doi.org/10.2307/1223890

WFO. (2025). World Flora Online Consortium [Published on the Internet; http://www.worldfloraonline.org/. Accessed: March 13, 2026]. http://www.worldfloraonline.org/

Zhang, X., Yu, L., Li, Y.-L., Suonan, Z., Komatsu, T., Qiu, G., Xu, S., Yue, S., Xu, M., Wang, F., Zhang, Y., Lee, K.-S., Liu, J.-X., Zhou, Y., & Reusch, T. B. H. (2025). Uncovering the *Nanozostera japonica* species complex suggests cryptic speciation and underestimated seagrass diversity. *New Phytologist*, *247*(5), 2086–2097. https://doi.org/10.1111/nph.70355

DRAFT

#### S4: DATASET DESCRIPTION

A total of 109,150 seagrass records across 54 taxa were compiled in the main dataset used in this study. Fig. 1 shows the raw species counts, after cleaning and deduplicating but prior to any thinning. The Fig. 2 map shows the gridded distribution and record coverage for all species, by overlaying the rasterised log-transformed frequencies on the biogeographical realms proposed in Spalding et al., (2007).

DRAFT

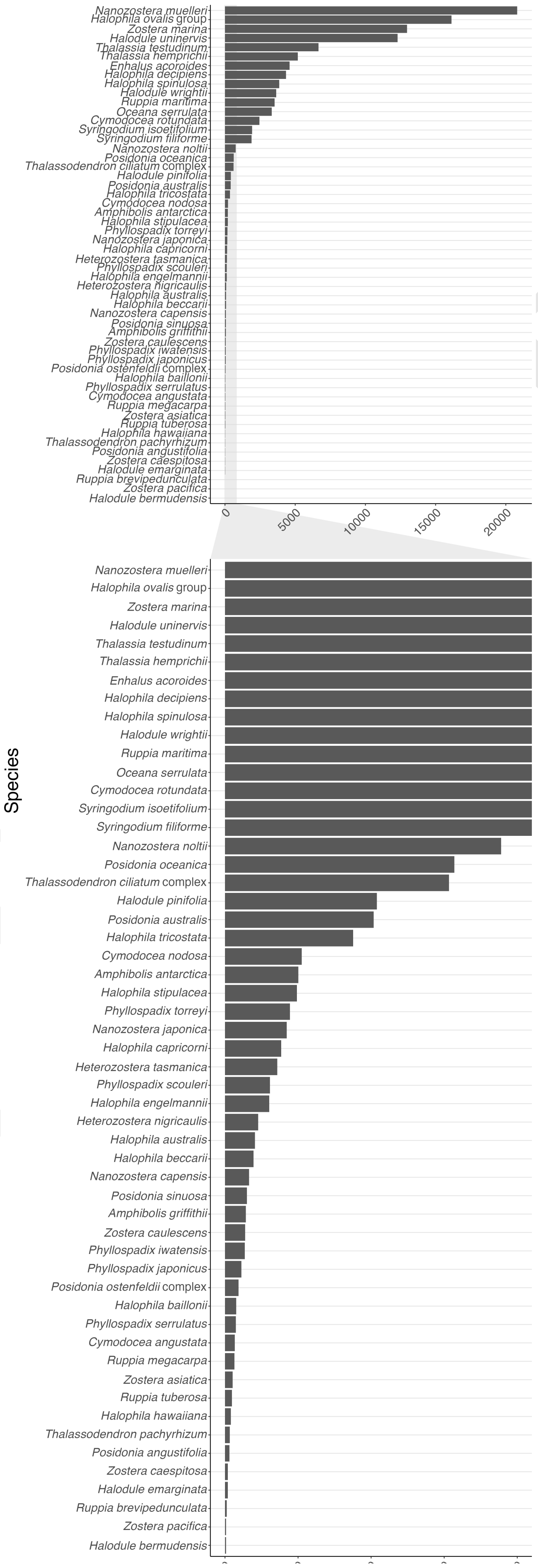
Nanozostera muelleri
Halophila ovalis group
Zostera marina
Halodule uninervis
Thalassia testudinum
Thalassia hemprichii
Enhalus acoroides
Halophila decipiens
Halophila spinulosa
Halodule wrightii
Ruppia maritima
Oceana serrulata
Cymodocea rotundata
Syringodium isoetifolium
Syringodium filiforme
Nanozostera noltii
Posidonia oceanica
Thalassodendron ciliatum complex
Halodule pinifolia
Posidonia australis
Halophila tricostata
Cymodocea nodosa
Amphibolis antarctica
Halophila stipulacea
Phyllospadix torreyi
Nanozostera japonica
Halophila capricorni
Heterozostera tasmanica
Phyllospadix scouleri
Halophila engelmannii
Heterozostera nigricaulis
Halophila australis
Halophila beccarii
Nanozostera capensis
Posidonia sinuosa
Amphibolis griffithii
Zostera caulescens
Phyllospadix iwatensis
Phyllospadix japonicus
Posidonia ostenfeldii complex
Halophila baillonii
Phyllospadix serrulatus
Cymodocea angustata
Ruppia megacarpa
Zostera asiatica
Ruppia tuberosa
Halophila hawaiiana
Thalassodendron pachyrhizum
Posidonia angustifolia
Zostera caespitosa
Halodule emarginata
Ruppia brevipedunculata
Zostera pacifica
Halodule bermudensis
0
5000
10000
15000
20000
Species

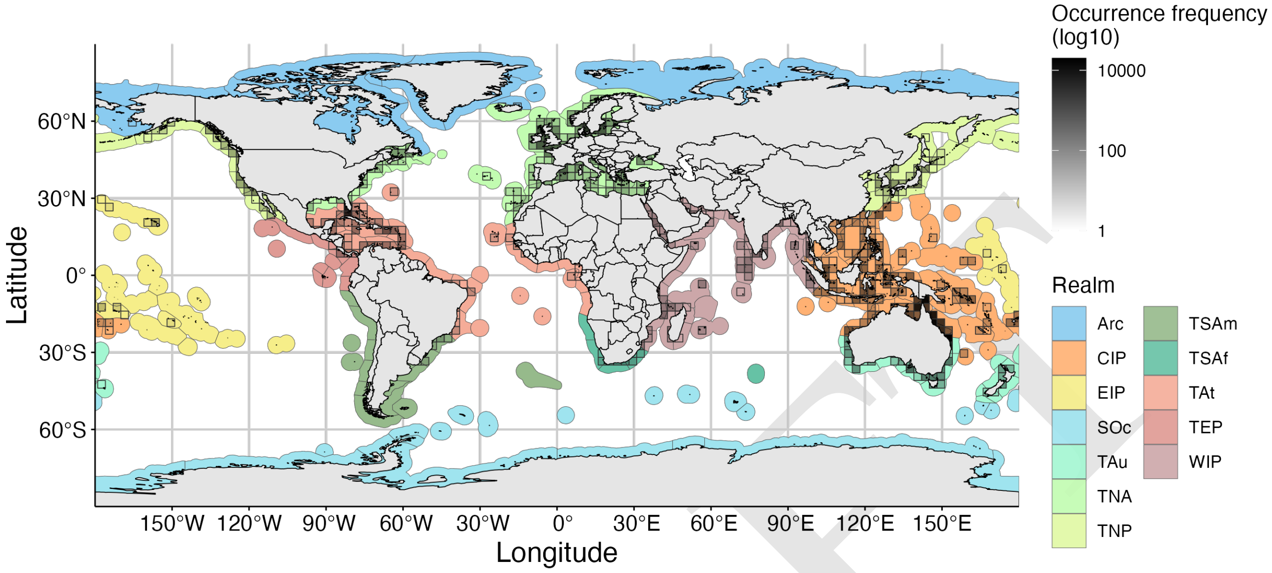


**Figure 2.** Global gridded map showing seagrass count frequencies. Cells (cell size = 3 °) that are contoured indicate at least a single seagrass presence. Darker colors indicate higher counts. Frequencies where log(10) transformed to normalize the data, making grid cells comparable. Biogeographic realms following Spalding et al., (2007) are plot in different colors under the following acronyms: TAu, Temperate Australasia; CIP, Central Indo-Pacific; SOc, Southern Ocean; TNA, Temperate Northern Atlantic; TAt, Tropical Atlantic; WIP, Western Indo-Pacific; TSAf, Temperate Southern Africa; TNP, Temperate Northern Pacific; TSAm, Temperate South America; EIP, Eastern Indo-Pacific; TEP, Tropical Eastern Pacific; Arc, Arctic

DRAFT

### S5: SUPPLEMENTARY RESULT TABLES

#### S5.1. Stabilization results

The effect of different thinning parameters [0, 17.5] on the thermal stabilization of seagrass species was measured with a stabilization threshold for each species and parameter pair ($ST_{s,\theta}$). The $ST_{s,\theta}$ was defined as the minimum number of observations required to obtain a stable estimate of the thermal optimum. Table 2 shows the $ST_{s,\theta}$ values for all $(s,\theta)$ pairs.

DRAFT

**Table 2.** Stabilization threshold (ST) values for each species and parameter pairs. Species are shown as rows and are ordered by decreasing range size. Thinning parameters (measured in latitudinal degrees) are shown as columns. The ST values represent the random observation number at which the thermal curve becomes stable. When stabilization was not reached ST = NA is given.

| species | 0.001 | 0.01 | 0.1 | 1 | 2.5 | 5 | 7.5 | 10 | 12.5 | 15 | 17.5 |
|---|---|---|---|---|---|---|---|---|---|---|---|
| *Zostera marina* | 7 | 3 | 20 | 45 | 69 | NA | NA | NA | NA | NA | NA |
| *Halophila ovalis* group | 40 | 82 | 45 | 80 | 89 | 68 | 28 | 30 | 28 | NA | NA |
| *Halodule uninervis* | 88 | 11 | 34 | 42 | 46 | 38 | 33 | 20 | 15 | 14 | NA |
| *Thalassia hemprichii* | 8 | 7 | 50 | 12 | 12 | 12 | 33 | 20 | NA | NA | NA |
| *Cymodocea rotundata* | 3 | 16 | 71 | 14 | 8 | 14 | 14 | NA | NA | NA | NA |
| *Syringodium isoetifolium* | 21 | 8 | 21 | 40 | 22 | 14 | 20 | 14 | NA | NA | NA |
| *Halophila decipiens* | 46 | 29 | 11 | 36 | 41 | 30 | 30 | NA | NA | NA | NA |
| *Ruppia maritima* | 33 | NA | 169 | NA | NA | NA | NA | NA | NA | NA | NA |
| *Enhalus acoroides* | 3 | 14 | 10 | 10 | 2 | 4 | 6 | 7 | 3 | NA | NA |
| *Oceana serrulata* | NA | 9 | 15 | 36 | 34 | 18 | 28 | 16 | NA | NA | NA |
| *Posidonia oceanica* | 18 | 8 | 14 | 50 | 21 | NA | NA | NA | NA | NA | NA |
| *Halodule pinifolia* | 42 | 103 | 118 | 27 | 18 | 22 | 19 | NA | NA | NA | NA |
| *Halodule wrightii* | 3 | 5 | 29 | 23 | 26 | 10 | NA | NA | NA | NA | NA |
| *Halophila spinulosa* | 19 | 37 | 35 | 30 | NA | NA | NA | NA | NA | NA | NA |
| *Thalassia testudinum* | 2 | 2 | 34 | 7 | 8 | 8 | NA | NA | NA | NA | NA |
| *Cymodocea nodosa* | 28 | 24 | 53 | 10 | 19 | NA | NA | NA | NA | NA | NA |
| *Nanozostera muelleri* | 5 | 9 | 32 | NA | NA | NA | NA | NA | NA | NA | NA |
| *Halophila stipulacea* | 163 | NA | 108 | 53 | 21 | NA | NA | NA | NA | NA | NA |
| *Thalassodendron ciliatum* complex | 3 | 7 | 7 | 32 | 28 | 14 | NA | NA | NA | NA | NA |
| *Syringodium filiforme* | 2 | 6 | 7 | 10 | 5 | NA | NA | NA | NA | NA | NA |
| *Nanozostera noltii* | 24 | 184 | 63 | NA | NA | NA | NA | NA | NA | NA | NA |
| *Nanozostera japonica* | NA | 69 | NA | NA | NA | NA | NA | NA | NA | NA | NA |
| *Amphibolis antarctica* | 44 | 52 | 58 | NA | NA | NA | NA | NA | NA | NA | NA |
| *Posidonia australis* | 160 | 180 | 43 | NA | NA | NA | NA | NA | NA | NA | NA |
| *Halophila beccarii* | 25 | 34 | 35 | NA | 10 | NA | NA | NA | NA | NA | NA |
| *Phyllospadix torreyi* | 88 | 106 | 67 | NA | NA | NA | NA | NA | NA | NA | NA |
| *Halophila engelmannii* | 2 | 5 | 3 | 6 | NA | NA | NA | NA | NA | NA | NA |
| *Phyllospadix scouleri* | 36 | 21 | 31 | NA | NA | NA | NA | NA | NA | NA | NA |
| *Nanozostera capensis* | 32 | 45 | NA | NA | NA | NA | NA | NA | NA | NA | NA |
| *Heterozostera tasmanica* | 10 | 36 | 17 | NA | NA | NA | NA | NA | NA | NA | NA |
| *Halophila tricostata* | 10 | 3 | 5 | NA | NA | NA | NA | NA | NA | NA | NA |
| *Phyllospadix iwatensis* | NA | NA | NA | NA | NA | NA | NA | NA | NA | NA | NA |
| *Halophila capricorni* | 17 | 11 | 10 | NA | NA | NA | NA | NA | NA | NA | NA |
| *Halophila australis* | 7 | 8 | NA | NA | NA | NA | NA | NA | NA | NA | NA |
| *Amphibolis griffithii* | 28 | 21 | NA | NA | NA | NA | NA | NA | NA | NA | NA |
| *Posidonia sinuosa* | 20 | 10 | NA | NA | NA | NA | NA | NA | NA | NA | NA |
| *Heterozostera nigricaulis* | 5 | 3 | 7 | NA | NA | NA | NA | NA | NA | NA | NA |
| *Posidonia ostenfeldii* complex | NA | 10 | NA | NA | NA | NA | NA | NA | NA | NA | NA |
| *Phyllospadix japonicus* | 13 | 11 | NA | NA | NA | NA | NA | NA | NA | NA | NA |
| *Ruppia megacarpa* | NA | NA | NA | NA | NA | NA | NA | NA | NA | NA | NA |
| *Cymodocea angustata* | NA | NA | NA | NA | NA | NA | NA | NA | NA | NA | NA |
| *Halophila baillonii* | 6 | 8 | 2 | NA | NA | NA | NA | NA | NA | NA | NA |
| *Phyllospadix serrulatus* | 6 | NA | NA | NA | NA | NA | NA | NA | NA | NA | NA |
| *Posidonia angustifolia* | NA | NA | NA | NA | NA | NA | NA | NA | NA | NA | NA |
| *Zostera caulescens* | NA | NA | NA | NA | NA | NA | NA | NA | NA | NA | NA |
| *Ruppia tuberosa* | NA | NA | NA | NA | NA | NA | NA | NA | NA | NA | NA |
| *Halophila hawaiiana* | NA | NA | NA | NA | NA | NA | NA | NA | NA | NA | NA |
| *Zostera caespitosa* | NA | NA | NA | NA | NA | NA | NA | NA | NA | NA | NA |
| *Thalassodendron pachyrhizum* | NA | NA | NA | NA | NA | NA | NA | NA | NA | NA | NA |
| *Halodule emarginata* | NA | NA | NA | NA | NA | NA | NA | NA | NA | NA | NA |
| *Zostera asiatica* | NA | NA | NA | NA | NA | NA | NA | NA | NA | NA | NA |
| *Ruppia brevipedunculata* | NA | NA | NA | NA | NA | NA | NA | NA | NA | NA | NA |
| *Zostera pacifica* | NA | NA | NA | NA | NA | NA | NA | NA | NA | NA | NA |
| *Halodule bermudensis* | NA | NA | NA | NA | NA | NA | NA | NA | NA | NA | NA |

##### S5.2. Main results of our proposed methods

Each species was assigned its range size descriptor ($D_s$) and thinning parameters predicted by BD-TP and LS-TP in Table 3. The difference between the prediction by both methods was quantified as the absolute difference between both predicted parameters ('$|\theta_{BD} - \theta_{LS}|$'). Predicted parameters of 0.001º are the result of capping at this minimum conceptual parameter. Differences between methods are organised and plot in Fig. 3.

DRAFT

**Table 3.** Range size and predicted thinning parameters by seagrass species. Deviation between both methods is shown as the absolute difference between both predicted parameters

| Species | Range size (Ds) | LS parameter | BD parameter | Deviation (LS-BD) |
|---|---|---|---|---|
| *Amphibolis antarctica* | 409.5483 | 409.516 | 1.765 | 1.665 |
| *Amphibolis griffithii* | 248.0112 | 247.976 | 0.001 | 0.009 |
| *Cymodocea angustata* | 211.5238 | 211.508 | 0.001 | 0.000 |
| *Cymodocea nodosa* | 528.6249 | 528.652 | 3.844 | 1.344 |
| *Cymodocea rotundata* | 819.0012 | 818.992 | 8.910 | 1.410 |
| *Enhalus acoroides* | 728.741 | 728.677 | 7.334 | 5.166 |
| *Halodule bermudensis* | 52.90351 | 52.932 | 0.001 | 0.000 |
| *Halodule emarginata* | 118.2875 | 118.285 | 0.001 | 0.000 |
| *Halodule pinifolia* | 638.8024 | 638.802 | 5.766 | 1.734 |
| *Halodule uninervis* | 967.581 | 966.152 | 11.478 | 3.522 |
| *Halodule wrightii* | 623.2324 | 623.240 | 5.494 | 0.494 |
| *Halophila australis* | 253.604 | 253.528 | 0.001 | 0.009 |
| *Halophila baillonii* | 204.7687 | 204.816 | 0.001 | 0.099 |
| *Halophila beccarii* | 358.6189 | 358.540 | 0.876 | 1.624 |
| *Halophila capricorni* | 289.5484 | 289.618 | 0.001 | 0.099 |
| *Halophila decipiens* | 738.2032 | 738.194 | 7.500 | 0.000 |
| *Halophila engelmannii* | 317.1848 | 317.174 | 0.154 | 0.846 |
| *Halophila hawaiiana* | 158.6173 | 139.893 | 0.001 | 0.000 |
| *Halophila ovalis* group | 1141.156 | 1142.371 | 14.552 | 2.052 |
| *Halophila spinulosa* | 607.3378 | 607.381 | 5.218 | 4.218 |
| *Halophila stipulacea* | 515.276 | 515.265 | 3.610 | 1.110 |
| *Halophila tricostata* | 303.7406 | 303.696 | 0.001 | 0.099 |
| *Heterozostera nigricaulis* | 242.3342 | 236.494 | 0.001 | 0.099 |
| *Heterozostera tasmanica* | 303.72 | 303.706 | 0.001 | 0.099 |
| *Nanozostera capensis* | 308.3014 | 308.327 | 0.001 | 0.009 |
| *Nanozostera japonica* | 426.2205 | 426.283 | 2.058 | 2.048 |
| *Nanozostera muelleri* | 515.2861 | 515.297 | 3.611 | 3.511 |
| *Nanozostera noltii* | 439.1042 | 439.147 | 2.282 | 2.182 |
| *Oceana serrulata* | 713.1729 | 713.197 | 7.064 | 2.936 |
| *Phyllospadix iwatensis* | 299.1146 | 299.112 | 0.001 | 0.000 |
| *Phyllospadix japonicus* | 224.3127 | 224.309 | 0.001 | 0.009 |
| *Phyllospadix scouleri* | 312.7727 | 312.744 | 0.077 | 0.023 |
| *Phyllospadix serrulatus* | 197.8395 | 197.801 | 0.001 | 0.000 |
| *Phyllospadix torreyi* | 354.703 | 354.608 | 0.807 | 0.707 |
| *Posidonia angustifolia* | 175.3469 | 175.337 | 0.001 | 0.000 |
| *Posidonia australis* | 366.271 | 366.303 | 1.011 | 0.911 |
| *Posidonia oceanica* | 681.1346 | 670.823 | 6.325 | 3.825 |
| *Posidonia ostenfeldii* complex | 230.4725 | 230.471 | 0.001 | 0.009 |
| *Posidonia sinuosa* | 236.464 | 236.481 | 0.001 | 0.009 |
| *Ruppia brevipedunculata* | 91.64122 | 91.614 | 0.001 | 0.000 |
| *Ruppia maritima* | 738.2001 | 738.187 | 7.500 | 7.400 |
| *Ruppia megacarpa* | 224.3247 | 224.283 | 0.001 | 0.000 |
| *Ruppia tuberosa* | 167.1698 | 167.258 | 0.001 | 0.000 |
| *Syringodium filiforme* | 448.5692 | 448.570 | 2.447 | 0.053 |
| *Syringodium isoetifolium* | 803.4305 | 803.464 | 8.639 | 1.361 |
| *Thalassia hemprichii* | 904.8499 | 904.849 | 10.408 | 0.408 |
| *Thalassia testudinum* | 600.4687 | 600.458 | 5.097 | 0.097 |
| *Thalassodendron ciliatum* complex | 493.1542 | 493.065 | 3.223 | 1.777 |
| *Thalassodendron pachyrhizum* | 139.8916 | 139.953 | 0.001 | 0.000 |
| *Zostera asiatica* | 105.7539 | 105.804 | 0.001 | 0.000 |
| *Zostera caespitosa* | 139.895 | 139.881 | 0.001 | 0.000 |
| *Zostera caulescens* | 167.1756 | 167.220 | 0.001 | 0.000 |
| *Zostera marina* | 1150.906 | 1149.717 | 14.681 | 12.181 |
| *Zostera pacifica* | 74.84273 | 74.800 | 0.001 | 0.000 |



DRAFT

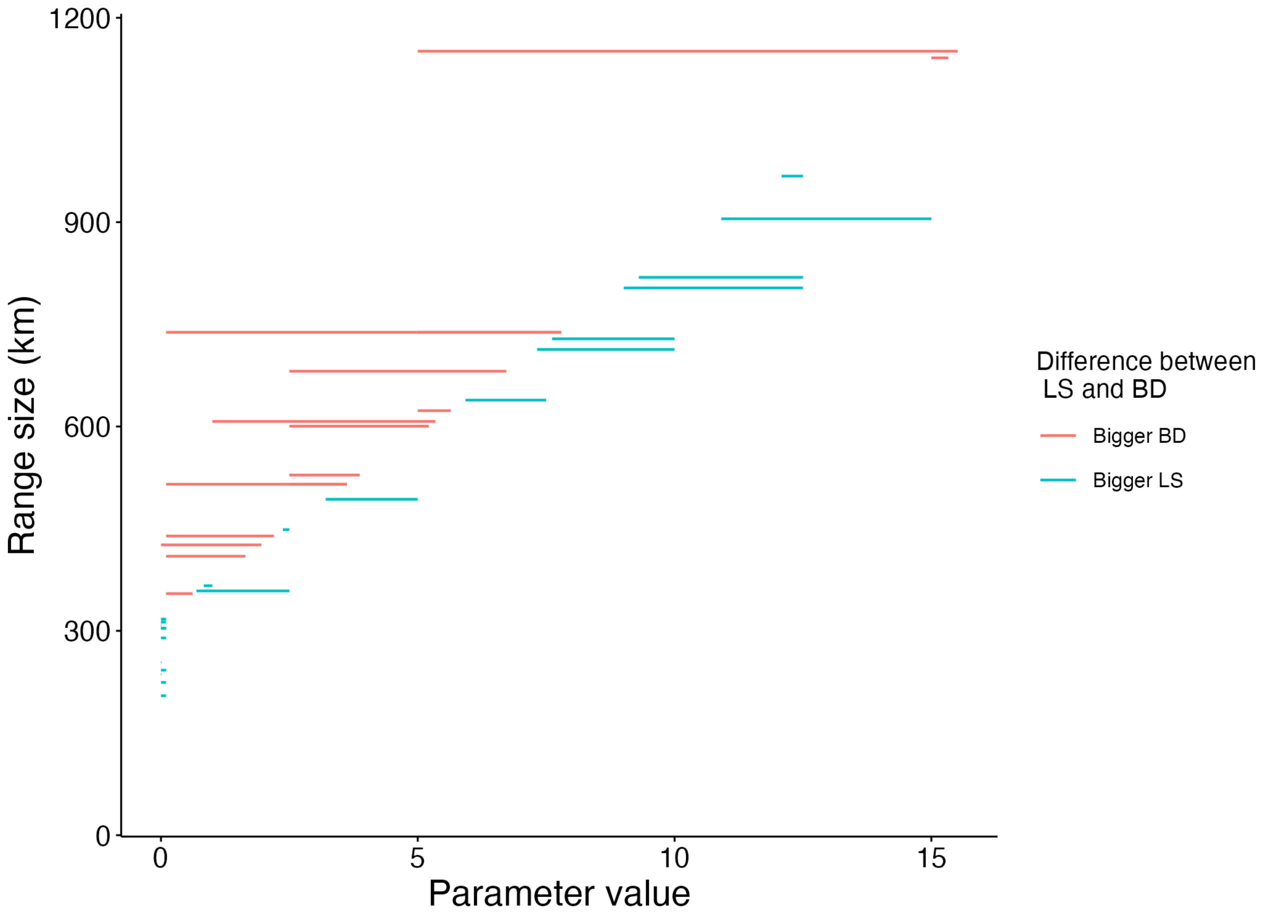


**Figure 3.** Differences in the parameter predictions by the BD-TP and LS-TP methods. Blue lines identify cases where the LS estimate is bigger than the BD estimate. Red lines identify cases where the BD estimate is bigger than the LS estimate. Barely visible lines show little to no difference between methods.

##### S5.3. Number of counts pre- and post- thinning

For each species (ordered by $D_s$), the original number of counts and the number of points post-thinning, after using BD-TP and LS-TP, is shown in Table 4. Only species deemed with sufficient data were considered.

DRAFT

**Table 4.** Species occurrence counts before and after the application of thinning with LS-TP and BD-TP parameterization. Only species that reach stability were considered.

| species | Raw counts | LS counts | BD counts | LS retention rate | BD retention rate |
|---|---|---|---|---|---|
| *Zostera marina* | 12964 | 151 | 31 | 0.012 | 0.002 |
| *Halophila ovalis* group | 16132 | 50 | 44 | 0.003 | 0.003 |
| *Halodule uninervis* | 12293 | 34 | 43 | 0.003 | 0.003 |
| *Thalassia hemprichii* | 5184 | 50 | 47 | 0.010 | 0.009 |
| *Cymodocea rotundata* | 2459 | 48 | 46 | 0.020 | 0.019 |
| *Syringodium isoetifolium* | 1930 | 41 | 49 | 0.021 | 0.025 |
| *Halophila decipiens* | 4349 | 49 | 49 | 0.011 | 0.011 |
| *Ruppia maritima* | 3532 | 312 | 62 | 0.088 | 0.018 |
| *Enhalus acoroides* | 4599 | 23 | 44 | 0.005 | 0.010 |
| *Oceana serrulata* | 3325 | 36 | 49 | 0.011 | 0.015 |
| *Posidonia oceanica* | 617 | 51 | 17 | 0.083 | 0.028 |
| *Halodule pinifolia* | 416 | 44 | 48 | 0.106 | 0.115 |
| *Halodule wrightii* | 3648 | 50 | 48 | 0.014 | 0.013 |
| *Halophila spinulosa* | 3861 | 66 | 16 | 0.017 | 0.004 |
| *Thalassia testudinum* | 6666 | 29 | 26 | 0.004 | 0.004 |
| *Cymodocea nodosa* | 210 | 45 | 35 | 0.214 | 0.167 |
| *Nanozostera muelleri* | 20813 | 224 | 32 | 0.011 | 0.002 |
| *Halophila stipulacea* | 197 | 40 | 34 | 0.203 | 0.173 |
| *Thalassodendron ciliatum* complex | 613 | 39 | 46 | 0.064 | 0.075 |
| *Syringodium filiforme* | 1892 | 40 | 40 | 0.021 | 0.021 |
| *Nanozostera noltii* | 756 | 113 | 38 | 0.149 | 0.050 |
| *Nanozostera japonica* | 169 | 118 | 39 | 0.698 | 0.231 |
| *Amphibolis antarctica* | 201 | 100 | 30 | 0.498 | 0.149 |
| *Posidonia australis* | 407 | 103 | 36 | 0.253 | 0.088 |
| *Halophila beccarii* | 78 | 29 | 41 | 0.372 | 0.526 |
| *Phyllospadix torreyi* | 178 | 89 | 39 | 0.500 | 0.219 |
| *Halophila engelmannii* | 121 | 25 | 48 | 0.207 | 0.397 |
| *Phyllospadix scouleri* | 123 | 55 | 55 | 0.447 | 0.447 |

DRAFT

**Table 5.** Summary table for cluster bootstrapping (i.e. per species)

| Variable | Mean | Median | SD | CI low | CI high |
|---|---|---|---|---|---|
| slope | 0.0187 | 0.0185 | 0.0012 | 0.0169 | 0.0223 |
| intercept | -6.1332 | -6.0552 | 0.6976 | -7.6624 | -4.8206 |
| $\theta$ at max($D_s$) | 14.6023 | 14.6555 | 1.7284 | 10.9074 | 18.2221 |
| $D_s$ at $\theta = 0$ | 306.9078 | 306.1227 | 25.6406 | 254.8980 | 357.0358 |

**Table 6.** Summary table for row bootstrapping (i.e. per species x parameter)

| Variable | Mean | Median | SD | CI low | CI high |
|---|---|---|---|---|---|
| slope | 0.0185 | 0.0185 | 0.0008 | 0.0173 | 0.0203 |
| intercept | -6.0964 | -6.0148 | 0.4977 | -7.1590 | -5.2281 |
| $\theta$ at max($D_s$) | 14.4426 | 14.4942 | 1.0221 | 12.5633 | 16.5617 |
| $D_s$ at $\theta = 0$ | 308.6049 | 305.7208 | 19.0933 | 272.4290 | 344.8819 |

##### S5.4. Bootstrap uncertainty and leave-one-species-out cross-validation for BD-TP

We evaluated bootstrap uncertainty by resampling both at row and species level. The row bootstrap produced estimates of the slope, intercept, and derived BD-TP quantities under the assumption that individual observations are independent. Cluster (i.e. species) bootstrap resampled entire species, preserving dependence among observations of the same species. Cluster bootstrap and row bootstrap results are summarised in Table 5 and Table 6, respectively. The slope and intercept of the BD method formula were resolved analytically. The estimated parameter for the highest observed $D_s$ and $D_s$ at $\theta = 0$ were calculated.

The original formula given in the main text

$$\theta_{opt} = f(D_s) = 0.017446 \cdot D_s - 5.378772 \tag{1}$$

differs slightly from the formula calculated from row bootstrapping (considering the mean between runs)

$$\theta_{opt} = f(D_s) = 0.01718430 \cdot D_s - 5.314048 \tag{2}$$

and cluster bootstraping (considering the mean between runs),

$$\theta_{opt} = f(D_s) = 0.01734393 \cdot D_s - 5.337891 \tag{3}$$

We further evaluated the linear SVM model with a Leave-One-Species-Out (LOSO) cross-validation scheme Table 7. For each iteration, all observations from one species were withheld as the test set, while the model was trained on the remaining species. Performance was summarised with true positives (TP), false positives (FP), true negatives (TN), and false negatives (FN), together with accuracy, precision, recall, and F1 score. Model accuracy typically fell for the species with higher $D_s$, as they may have a higher influence in the shape of the hyperplane (e.g. *Zostera marina* AUC = 0.55, *Ruppia maritima* AUC = 0.55, or *Enhalus acoroides* AUC = 0.73), although no statistical correlation was found between $D_s$ and any of the evaluation metrics. LOSO results indicate heterogeneous predictive performance among species, with some species showing complete agreement between observed and predicted stability and others showing reduced discrimination, particularly through false-positive or false-negative classifications.

DRAFT

**Table 7.** Evaluation metrics from the LOSO analysis. The species that was held out in each run is shown in the left.

| Held out species | n test pairs | TP | FP | TN | FN | Accuracy | Precision | Recall | F1 |
|---|---|---|---|---|---|---|---|---|---|
| *Amphibolis antarctica* | 11 | 3 | 1 | 7 | 0 | 0.91 | 0.75 | 1.00 | 0.86 |
| *Amphibolis griffithii* | 11 | 0 | 0 | 9 | 2 | 0.82 | NA | 0.00 | NA |
| *Cymodocea angustata* | 11 | 0 | 0 | 11 | 0 | 1.00 | NA | NA | NA |
| *Cymodocea nodosa* | 11 | 5 | 0 | 6 | 0 | 1.00 | 1.00 | 1.00 | 1.00 |
| *Cymodocea rotundata* | 11 | 7 | 0 | 4 | 0 | 1.00 | 1.00 | 1.00 | 1.00 |
| *Enhalus acoroides* | 11 | 6 | 0 | 2 | 3 | 0.73 | 1.00 | 0.67 | 0.80 |
| *Halodule bermudensis* | 2 | 0 | 0 | 2 | 0 | 1.00 | NA | NA | NA |
| *Halodule emarginata* | 11 | 0 | 0 | 11 | 0 | 1.00 | NA | NA | NA |
| *Halodule pinifolia* | 11 | 6 | 0 | 4 | 1 | 0.91 | 1.00 | 0.86 | 0.92 |
| *Halodule uninervis* | 11 | 8 | 0 | 1 | 2 | 0.82 | 1.00 | 0.80 | 0.89 |
| *Halodule wrightii* | 11 | 6 | 0 | 5 | 0 | 1.00 | 1.00 | 1.00 | 1.00 |
| *Halophila australis* | 11 | 0 | 0 | 9 | 2 | 0.82 | NA | 0.00 | NA |
| *Halophila baillonii* | 11 | 0 | 0 | 8 | 3 | 0.73 | NA | 0.00 | NA |
| *Halophila beccarii* | 11 | 3 | 0 | 7 | 1 | 0.91 | 1.00 | 0.75 | 0.86 |
| *Halophila capricorni* | 8 | 0 | 0 | 5 | 3 | 0.63 | NA | 0.00 | NA |
| *Halophila decipiens* | 11 | 6 | 0 | 4 | 1 | 0.91 | 1.00 | 0.86 | 0.92 |
| *Halophila engelmannii* | 11 | 2 | 0 | 7 | 2 | 0.82 | 1.00 | 0.50 | 0.67 |
| *Halophila hawaiiana* | 11 | 0 | 0 | 11 | 0 | 1.00 | NA | NA | NA |
| *Halophila ovalis* group | 11 | 9 | 0 | 2 | 0 | 1.00 | 1.00 | 1.00 | 1.00 |
| *Halophila spinulosa* | 11 | 4 | 2 | 5 | 0 | 0.82 | 0.67 | 1.00 | 0.80 |
| *Halophila stipulacea* | 11 | 4 | 1 | 6 | 0 | 0.91 | 0.80 | 1.00 | 0.89 |
| *Halophila tricostata* | 8 | 0 | 0 | 5 | 3 | 0.63 | NA | 0.00 | NA |
| *Heterozostera nigricaulis* | 11 | 0 | 0 | 8 | 3 | 0.73 | NA | 0.00 | NA |
| *Heterozostera tasmanica* | 11 | 0 | 0 | 8 | 3 | 0.73 | NA | 0.00 | NA |
| *Nanozostera capensis* | 11 | 2 | 0 | 9 | 0 | 1.00 | 1.00 | 1.00 | 1.00 |
| *Nanozostera japonica* | 11 | 1 | 3 | 7 | 0 | 0.73 | 0.25 | 1.00 | 0.40 |
| *Nanozostera muelleri* | 11 | 3 | 2 | 6 | 0 | 0.82 | 0.60 | 1.00 | 0.75 |
| *Nanozostera noltii* | 11 | 3 | 1 | 7 | 0 | 0.91 | 0.75 | 1.00 | 0.86 |
| *Oceana serrulata* | 11 | 5 | 1 | 3 | 2 | 0.73 | 0.83 | 0.71 | 0.77 |
| *Phyllospadix iwatensis* | 11 | 0 | 0 | 11 | 0 | 1.00 | NA | NA | NA |
| *Phyllospadix japonicus* | 9 | 0 | 0 | 7 | 2 | 0.78 | NA | 0.00 | NA |
| *Phyllospadix scouleri* | 11 | 2 | 0 | 8 | 1 | 0.91 | 1.00 | 0.67 | 0.80 |
| *Phyllospadix serrulatus* | 11 | 0 | 0 | 10 | 1 | 0.91 | NA | 0.00 | NA |
| *Phyllospadix torreyi* | 11 | 3 | 0 | 8 | 0 | 1.00 | 1.00 | 1.00 | 1.00 |
| *Posidonia angustifolia* | 11 | 0 | 0 | 11 | 0 | 1.00 | NA | NA | NA |
| *Posidonia australis* | 11 | 3 | 0 | 8 | 0 | 1.00 | 1.00 | 1.00 | 1.00 |
| *Posidonia oceanica* | 11 | 5 | 1 | 5 | 0 | 0.91 | 0.83 | 1.00 | 0.91 |
| *Posidonia ostenfeldii* complex | 11 | 0 | 0 | 10 | 1 | 0.91 | NA | 0.00 | NA |
| *Posidonia sinuosa* | 11 | 0 | 0 | 9 | 2 | 0.82 | NA | 0.00 | NA |
| *Ruppia brevipedunculata* | 11 | 0 | 0 | 11 | 0 | 1.00 | NA | NA | NA |
| *Ruppia maritima* | 11 | 2 | 5 | 4 | 0 | 0.55 | 0.29 | 1.00 | 0.44 |
| *Ruppia megacarpa* | 11 | 0 | 0 | 11 | 0 | 1.00 | NA | NA | NA |
| *Ruppia tuberosa* | 5 | 0 | 0 | 5 | 0 | 1.00 | NA | NA | NA |
| *Syringodium filiforme* | 11 | 4 | 0 | 6 | 1 | 0.91 | 1.00 | 0.80 | 0.89 |
| *Syringodium isoetifolium* | 11 | 7 | 0 | 3 | 1 | 0.91 | 1.00 | 0.88 | 0.93 |
| *Thalassia hemprichii* | 11 | 8 | 0 | 3 | 0 | 1.00 | 1.00 | 1.00 | 1.00 |
| *Thalassia testudinum* | 11 | 6 | 0 | 5 | 0 | 1.00 | 1.00 | 1.00 | 1.00 |
| *Thalassodendron ciliatum* complex | 11 | 5 | 0 | 5 | 1 | 0.91 | 1.00 | 0.83 | 0.91 |
| *Thalassodendron pachyrhizum* | 7 | 0 | 0 | 7 | 0 | 1.00 | NA | NA | NA |
| *Zostera asiatica* | 10 | 0 | 0 | 10 | 0 | 1.00 | NA | NA | NA |
| *Zostera caespitosa* | 11 | 0 | 0 | 11 | 0 | 1.00 | NA | NA | NA |
| *Zostera caulescens* | 10 | 0 | 0 | 10 | 0 | 1.00 | NA | NA | NA |
| *Zostera marina* | 11 | 5 | 5 | 1 | 0 | 0.55 | 0.50 | 1.00 | 0.67 |
| *Zostera pacifica* | 2 | 0 | 0 | 2 | 0 | 1.00 | NA | NA | NA |

DRAFT

#### S6: STABILIZATION GRAPHS FOR ALL SPECIES

Stabilization plots of all species for all tested parameters, ordered by decreasing $D_s$.

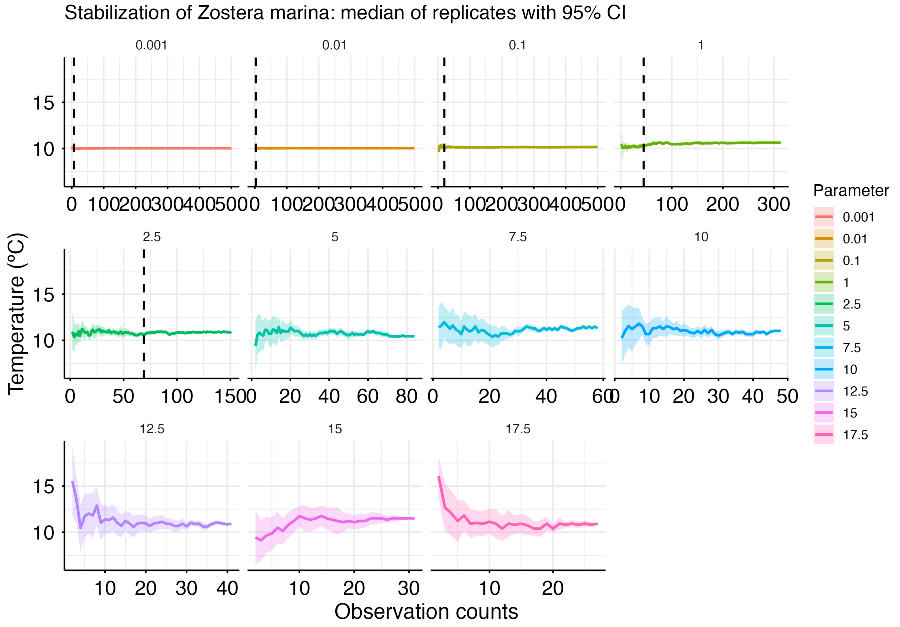


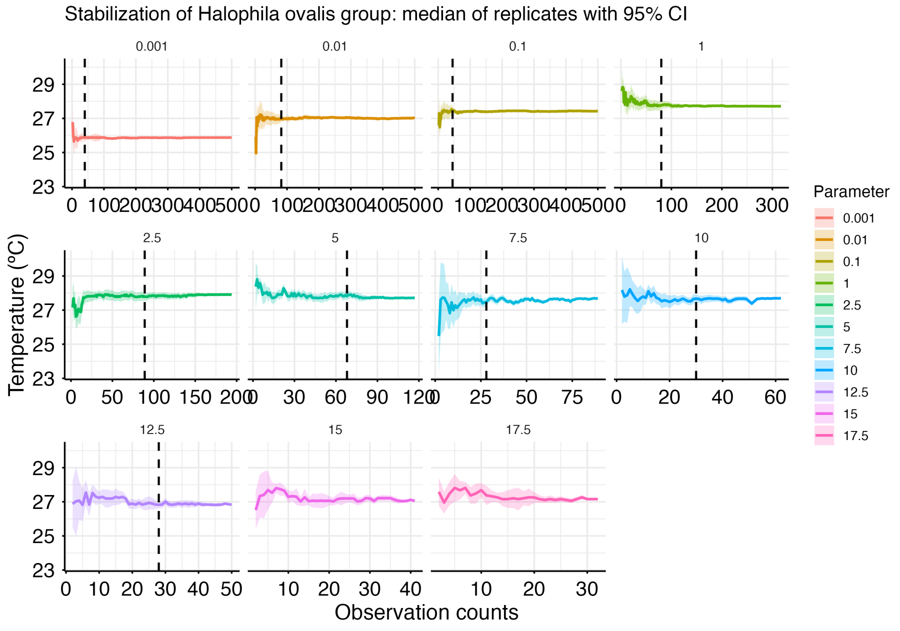

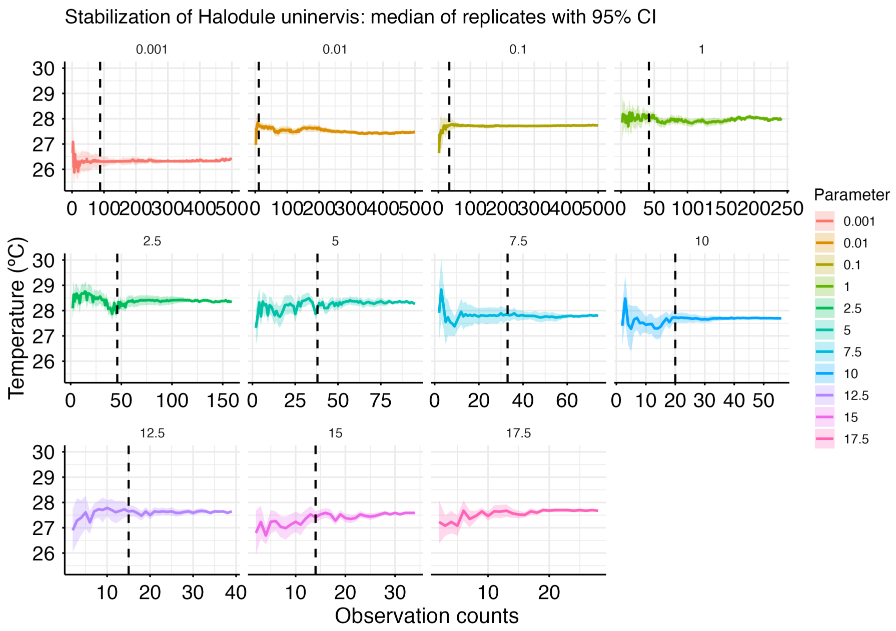
Stabilization of Halodule uninervis: median of replicates with 95% CI
Temperature (°C)
Observation counts
Parameter
0.001
0.01
0.1
1
2.5
5
7.5
10
12.5
15
17.5

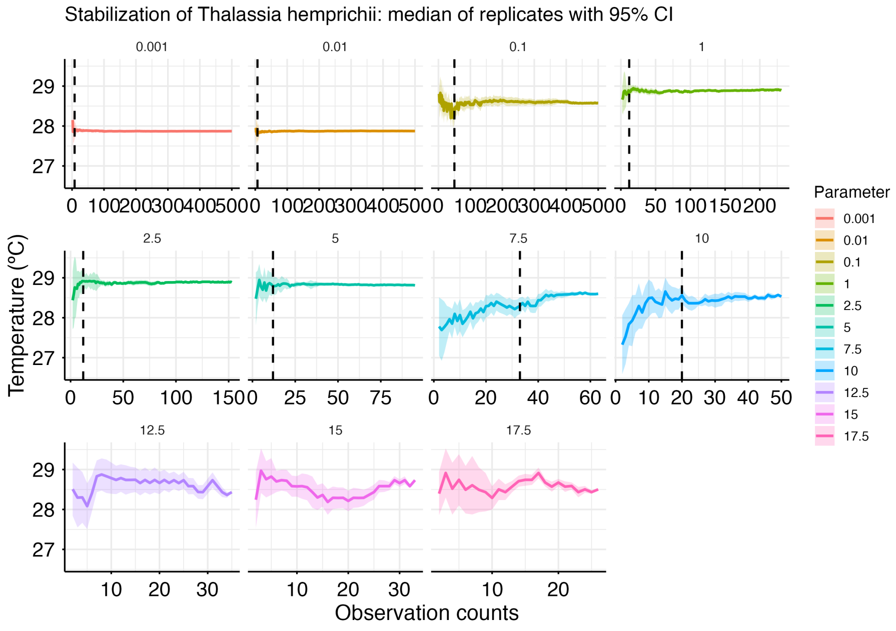
Stabilization of Thalassia hemprichii: median of replicates with 95% CI
Temperature (°C)
Observation counts
Parameter
0.001
0.01
0.1
1
2.5
5
7.5
10
12.5
15
17.5

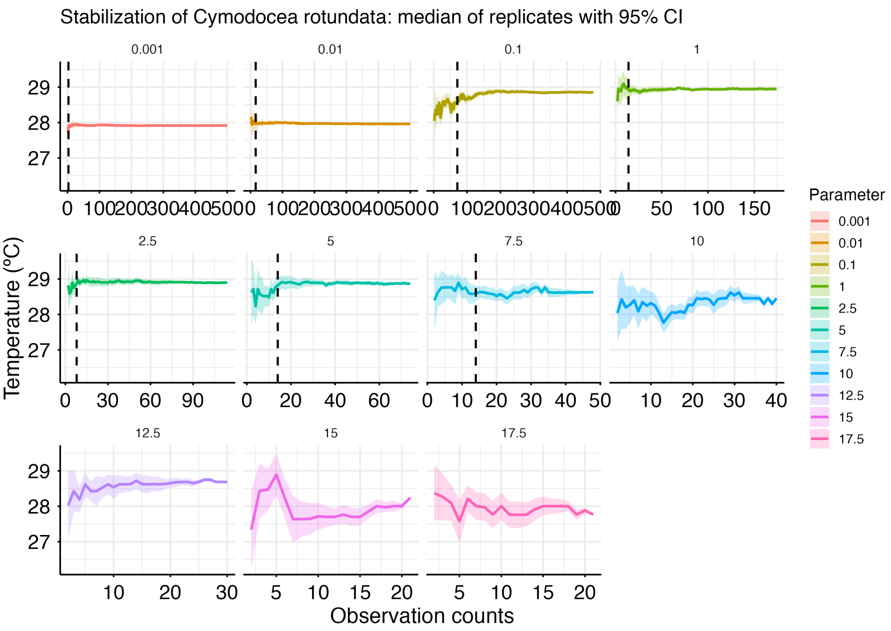
Stabilization of Cymodocea rotundata: median of replicates with 95% CI
0.001
0.01
0.1
1
2.5
5
7.5
10
12.5
15
17.5
Temperature (°C)
Observation counts
Parameter
0.001
0.01
0.1
1
2.5
5
7.5
10
12.5
15
17.5

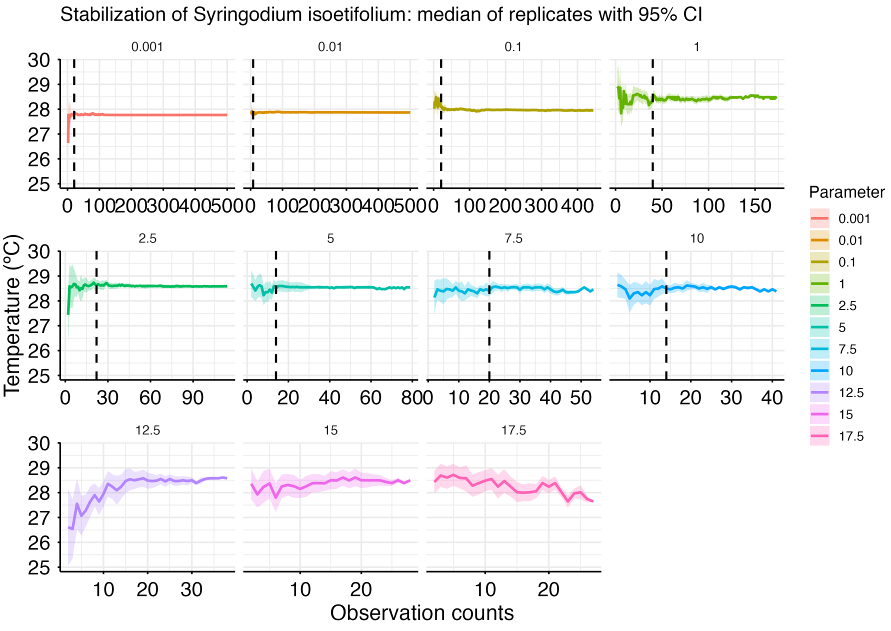
Stabilization of Syringodium isoetifolium: median of replicates with 95% CI
0.001
0.01
0.1
1
2.5
5
7.5
10
12.5
15
17.5
Temperature (°C)
Observation counts
Parameter
0.001
0.01
0.1
1
2.5
5
7.5
10
12.5
15
17.5

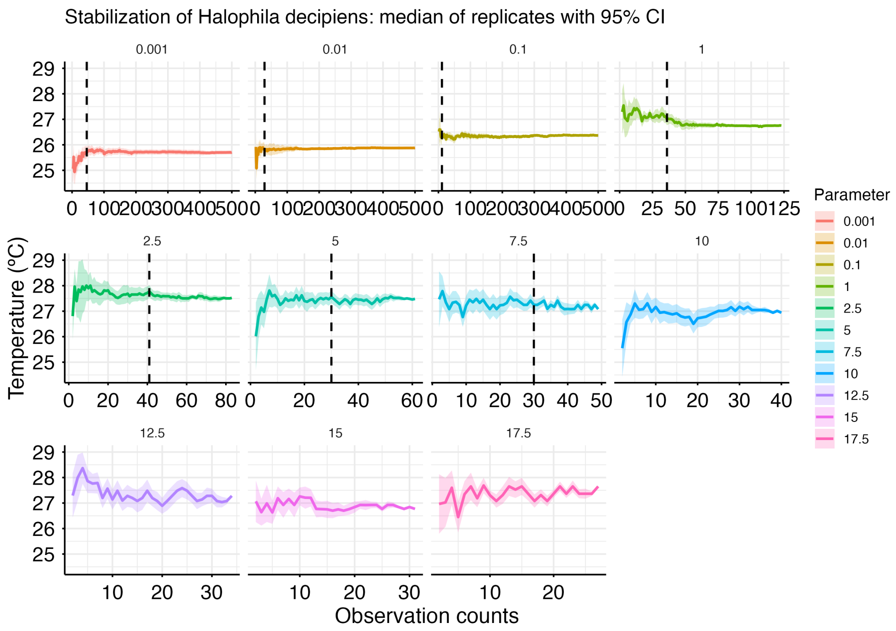
Stabilization of Halophila decipiens: median of replicates with 95% CI
0.001
0.01
0.1
1
2.5
5
7.5
10
12.5
15
17.5
Temperature (°C)
Observation counts
Parameter
0.001
0.01
0.1
1
2.5
5
7.5
10
12.5
15
17.5

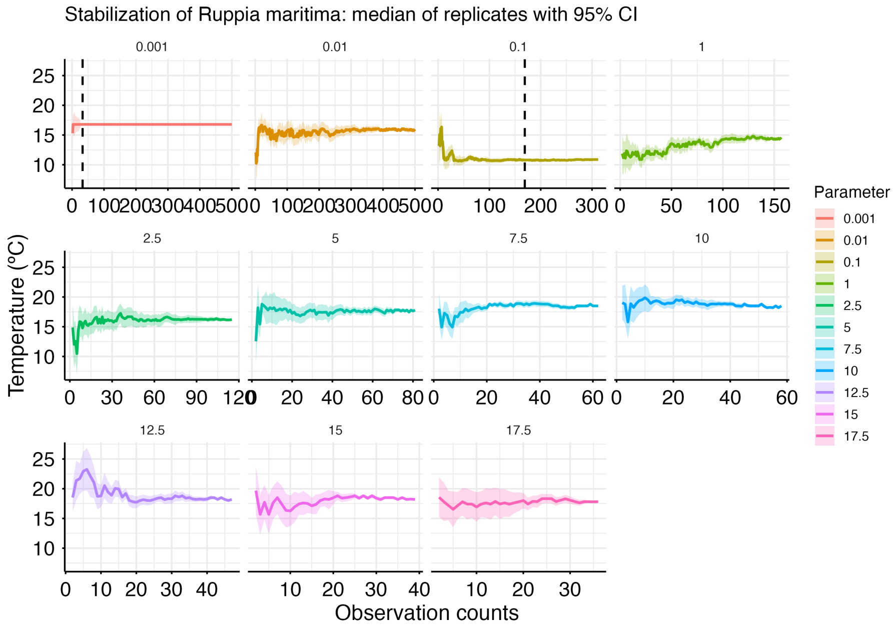
Stabilization of Ruppia maritima: median of replicates with 95% CI
0.001
0.01
0.1
1
2.5
5
7.5
10
12.5
15
17.5
Temperature (°C)
Observation counts
Parameter
0.001
0.01
0.1
1
2.5
5
7.5
10
12.5
15
17.5

Stabilization of Enhalus acoroides: median of replicates with 95% CI

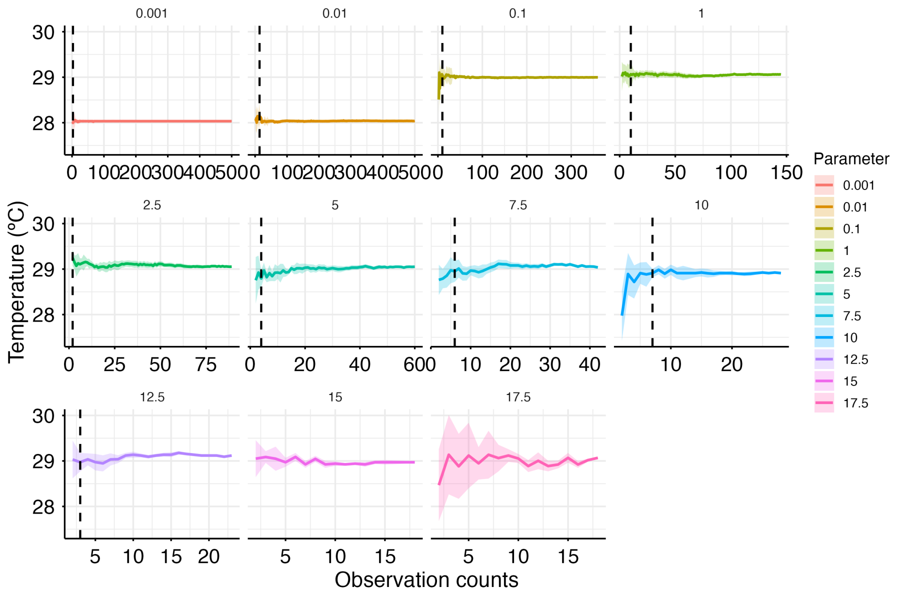


Stabilization of Oceana serrulata: median of replicates with 95% CI

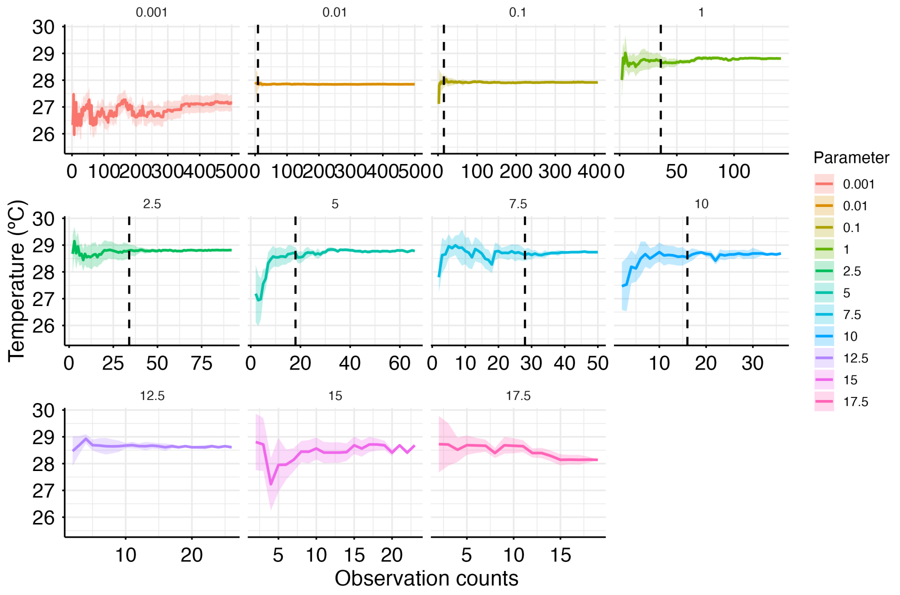

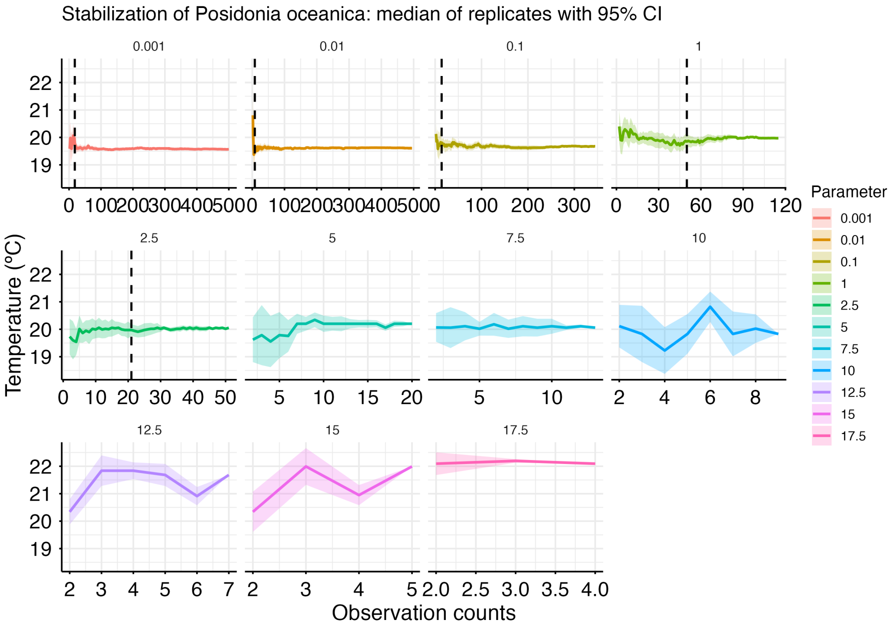
Stabilization of Posidonia oceanica: median of replicates with 95% CI
0.001
0.01
0.1
1
2.5
5
7.5
10
12.5
15
17.5
Temperature (°C)
Observation counts
Parameter
0.001
0.01
0.1
1
2.5
5
7.5
10
12.5
15
17.5

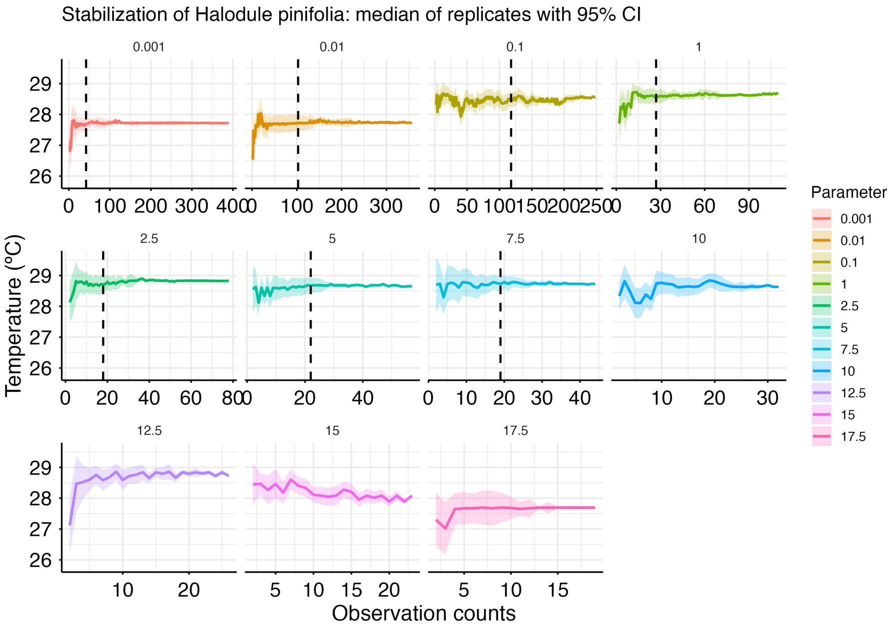
Stabilization of Halodule pinifolia: median of replicates with 95% CI
0.001
0.01
0.1
1
2.5
5
7.5
10
12.5
15
17.5
Temperature (°C)
Observation counts
Parameter
0.001
0.01
0.1
1
2.5
5
7.5
10
12.5
15
17.5

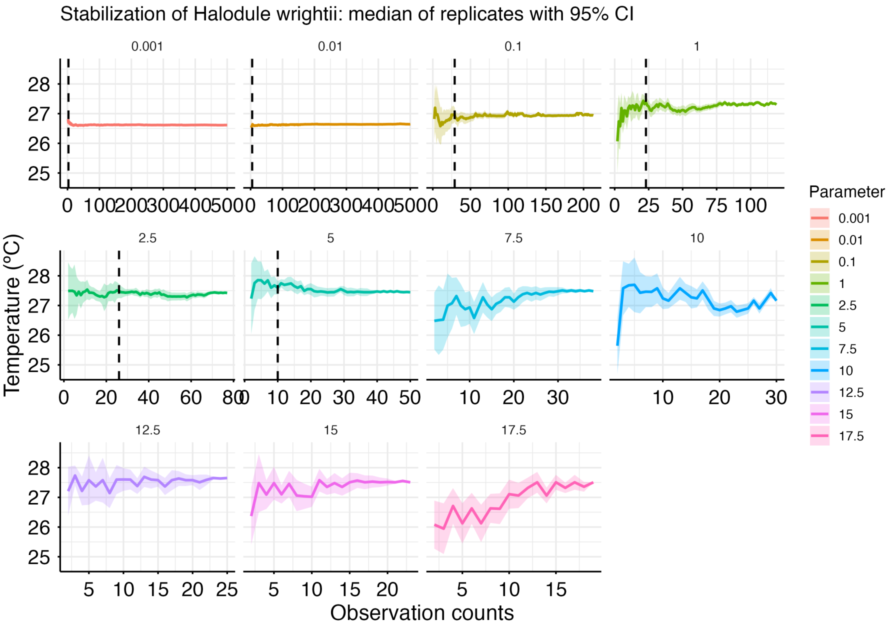
Stabilization of Halodule wrightii: median of replicates with 95% CI
0.001
0.01
0.1
1
2.5
5
7.5
10
12.5
15
17.5
Temperature (°C)
Observation counts
Parameter
0.001
0.01
0.1
1
2.5
5
7.5
10
12.5
15
17.5

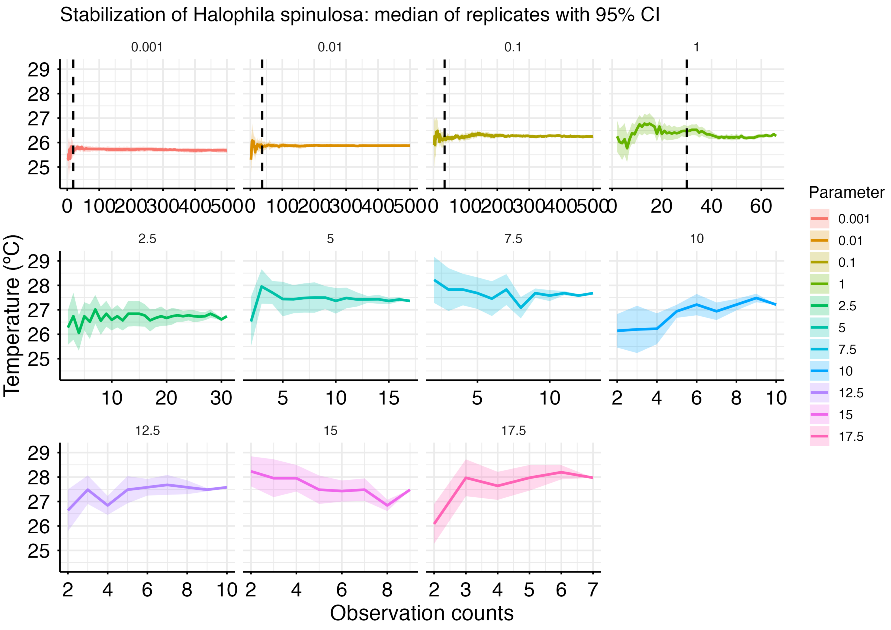
Stabilization of Halophila spinulosa: median of replicates with 95% CI
0.001
0.01
0.1
1
2.5
5
7.5
10
12.5
15
17.5
Temperature (°C)
Observation counts
Parameter
0.001
0.01
0.1
1
2.5
5
7.5
10
12.5
15
17.5

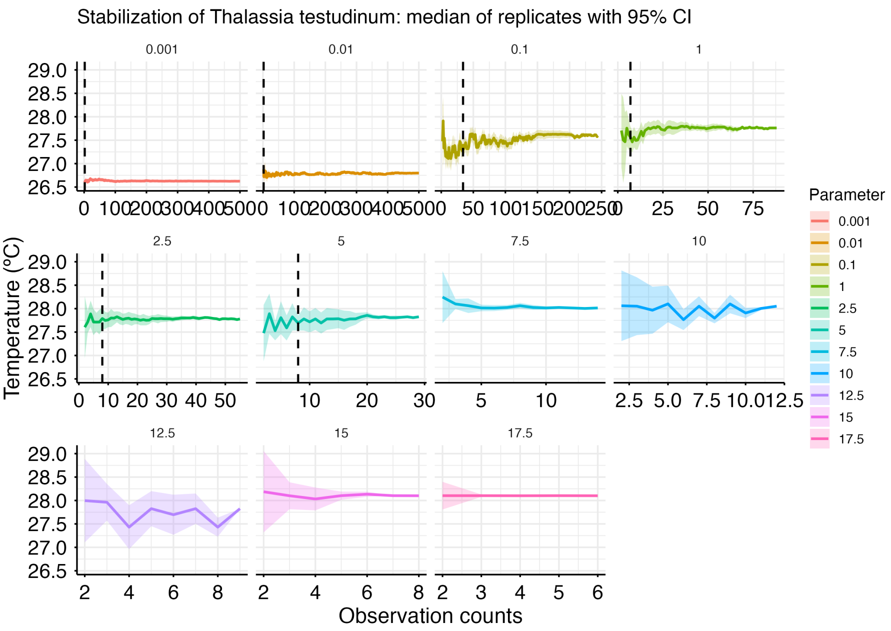
Stabilization of Thalassia testudinum: median of replicates with 95% CI
0.001
0.01
0.1
1
2.5
5
7.5
10
12.5
15
17.5
Temperature (°C)
Observation counts
Parameter
0.001
0.01
0.1
1
2.5
5
7.5
10
12.5
15
17.5

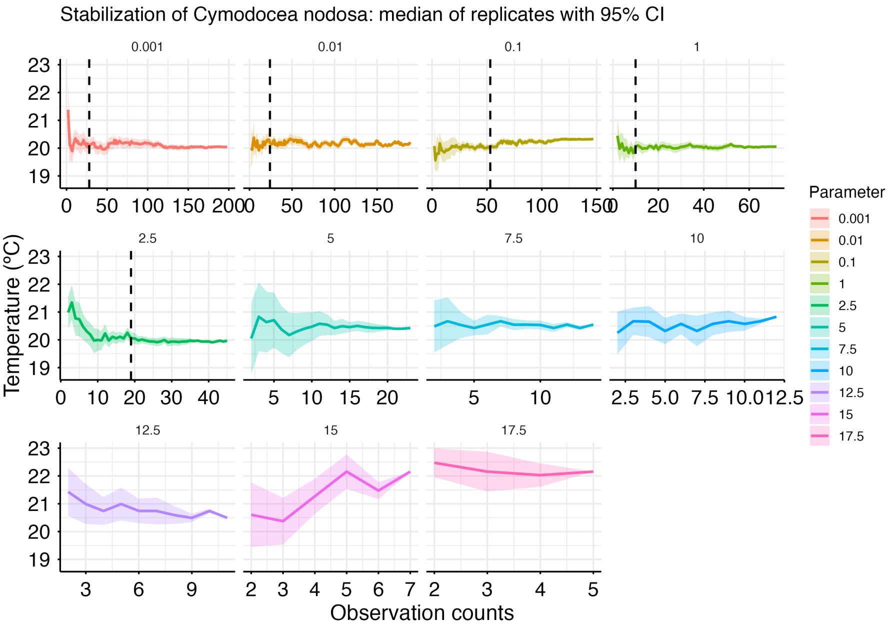
Stabilization of Cymodocea nodosa: median of replicates with 95% CI
0.001
0.01
0.1
1
2.5
5
7.5
10
12.5
15
17.5
Temperature (°C)
Observation counts
Parameter
0.001
0.01
0.1
1
2.5
5
7.5
10
12.5
15
17.5

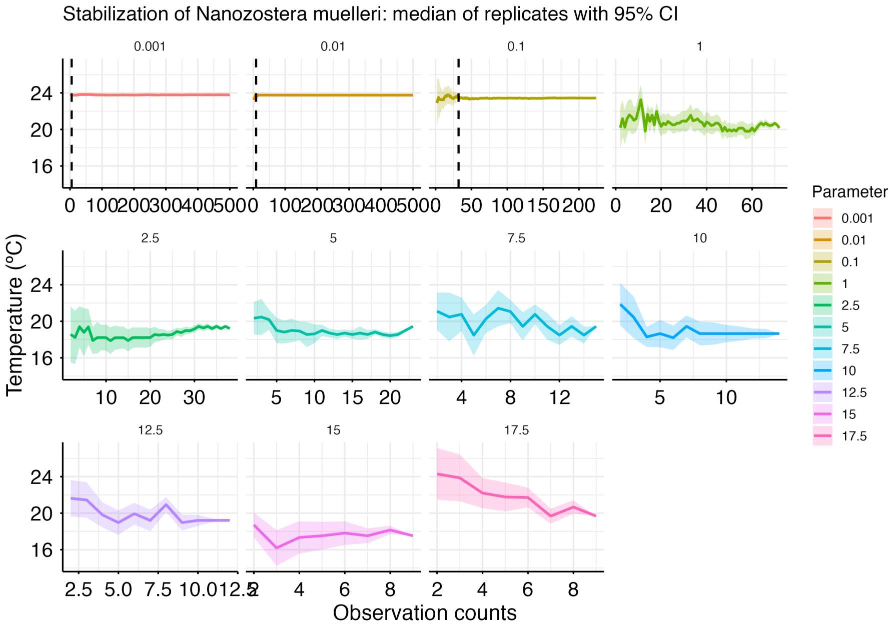
Stabilization of Nanozostera muelleri: median of replicates with 95% CI
Temperature (°C)
Observation counts
Parameter
0.001
0.01
0.1
1
2.5
5
7.5
10
12.5
15
17.5

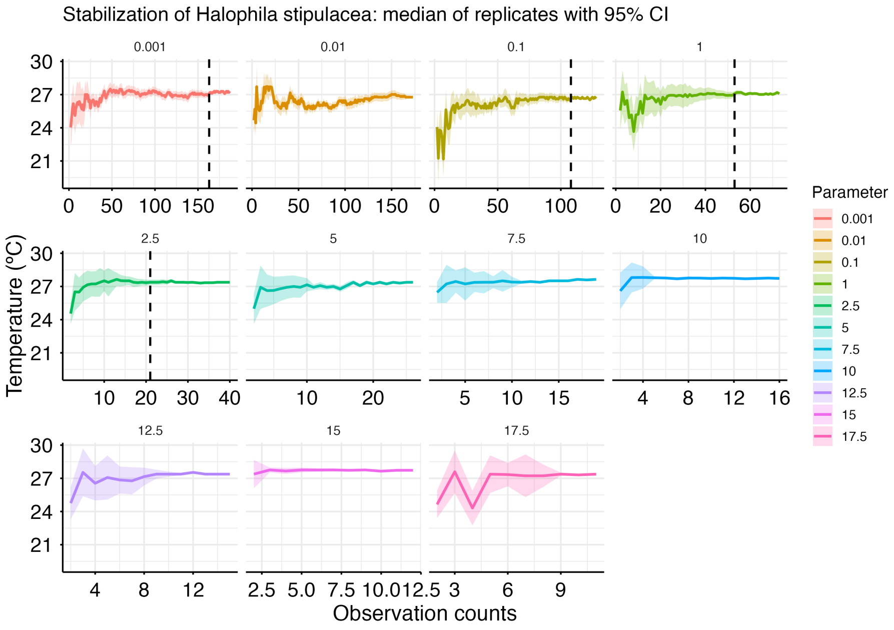
Stabilization of Halophila stipulacea: median of replicates with 95% CI
Temperature (°C)
Observation counts
Parameter
0.001
0.01
0.1
1
2.5
5
7.5
10
12.5
15
17.5

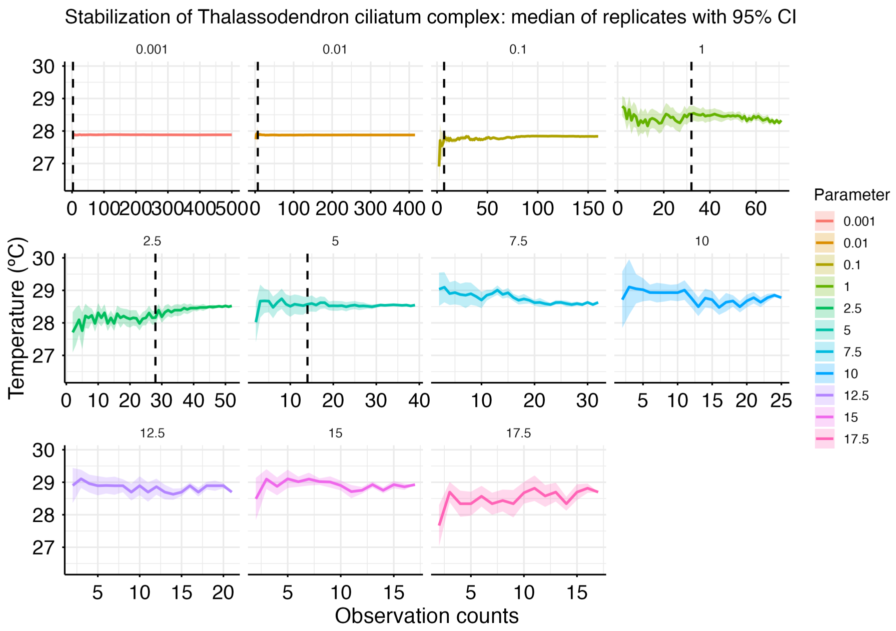

Stabilization of Thalassodendron ciliatum complex: median of replicates with 95% CI
Temperature (°C)
Observation counts
Parameter
0.001
0.01
0.1
1
2.5
5
7.5
10
12.5
15
17.5


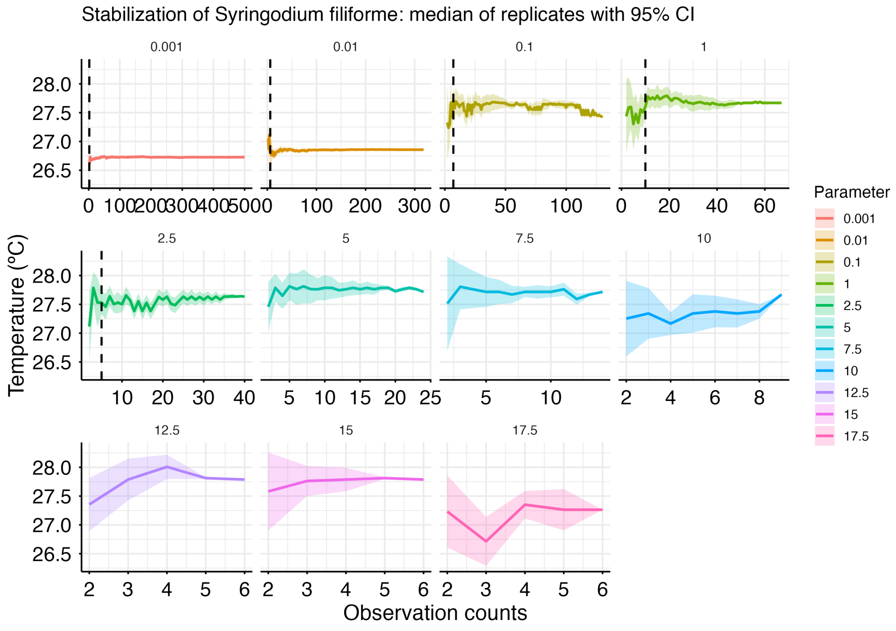

Stabilization of Syringodium filiforme: median of replicates with 95% CI
Temperature (°C)
Observation counts
Parameter
0.001
0.01
0.1
1
2.5
5
7.5
10
12.5
15
17.5

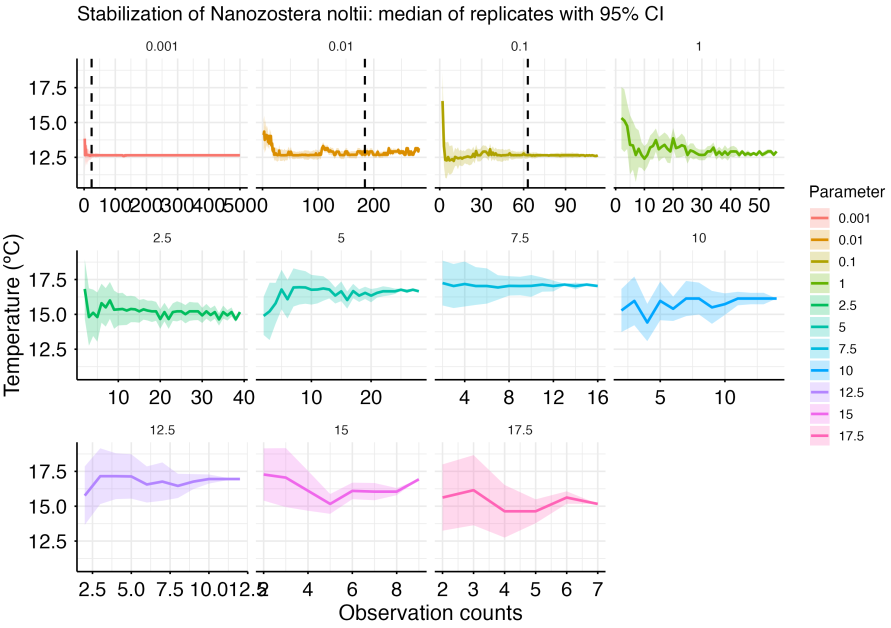
Stabilization of Nanozostera noltii: median of replicates with 95% CI
0.001
0.01
0.1
1
2.5
5
7.5
10
12.5
15
17.5
Temperature (°C)
Observation counts
Parameter

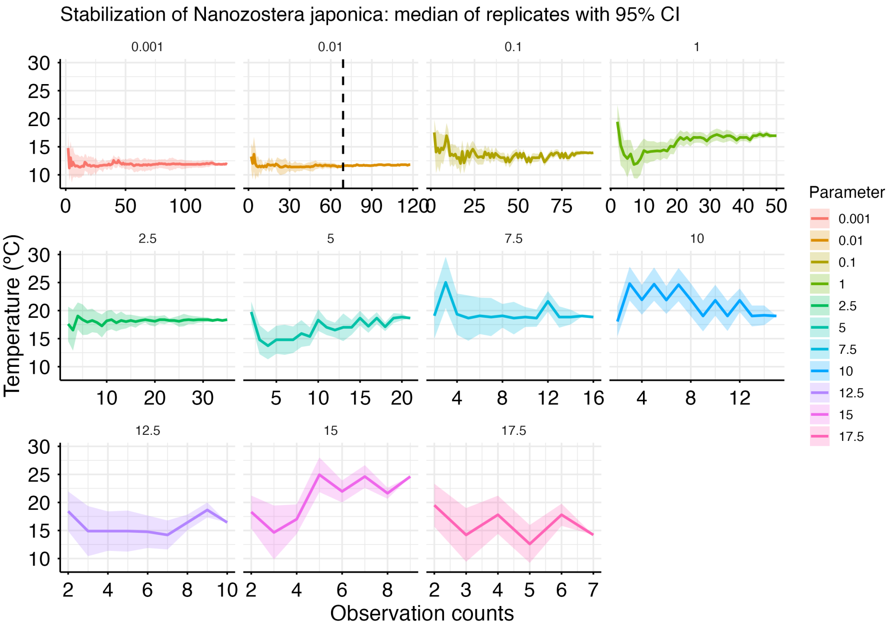
Stabilization of Nanozostera japonica: median of replicates with 95% CI
0.001
0.01
0.1
1
2.5
5
7.5
10
12.5
15
17.5
Temperature (°C)
Observation counts
Parameter

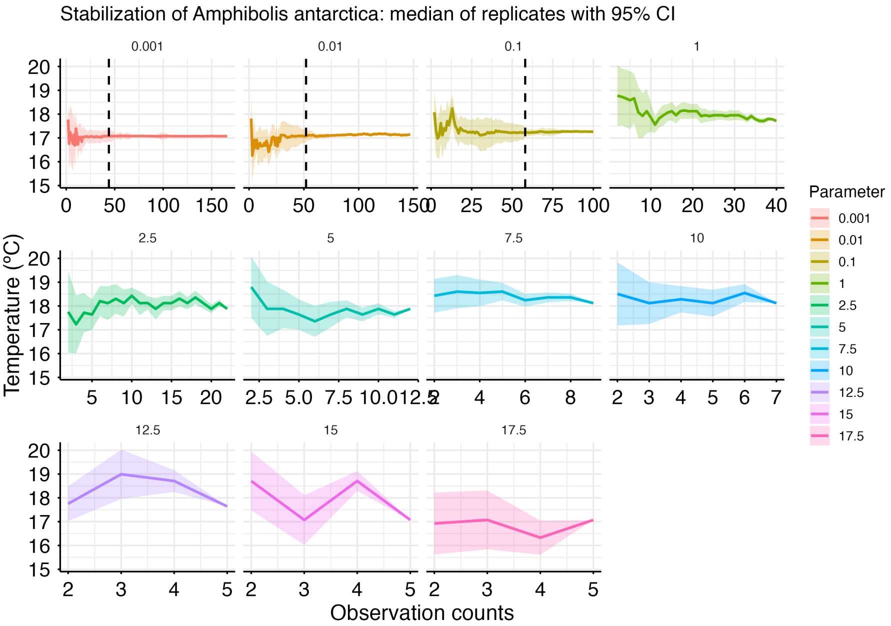
Stabilization of Amphibolis antarctica: median of replicates with 95% CI
0.001
0.01
0.1
1
2.5
5
7.5
10
12.5
15
17.5
Temperature (°C)
Observation counts
Parameter
0.001
0.01
0.1
1
2.5
5
7.5
10
12.5
15
17.5

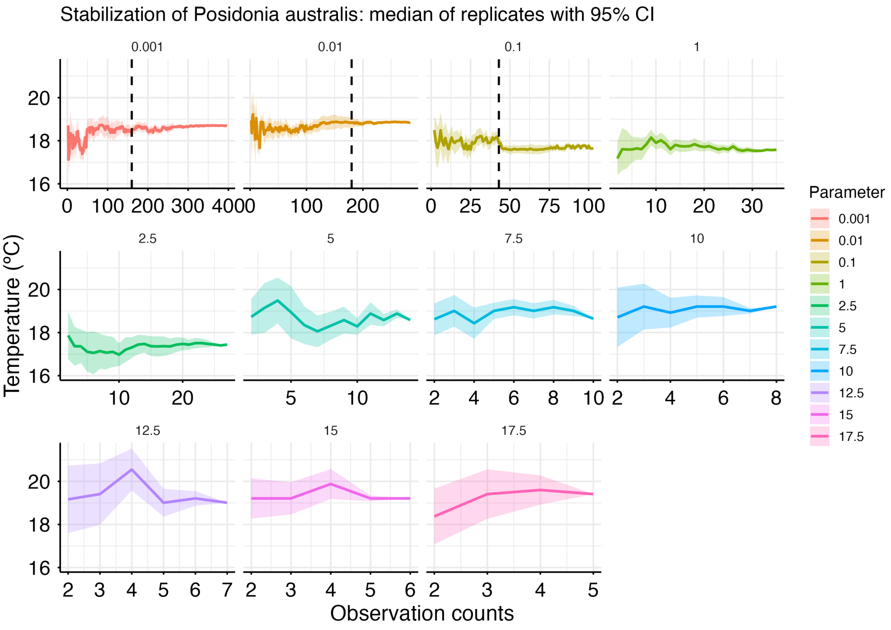
Stabilization of Posidonia australis: median of replicates with 95% CI
0.001
0.01
0.1
1
2.5
5
7.5
10
12.5
15
17.5
Temperature (°C)
Observation counts
Parameter
0.001
0.01
0.1
1
2.5
5
7.5
10
12.5
15
17.5

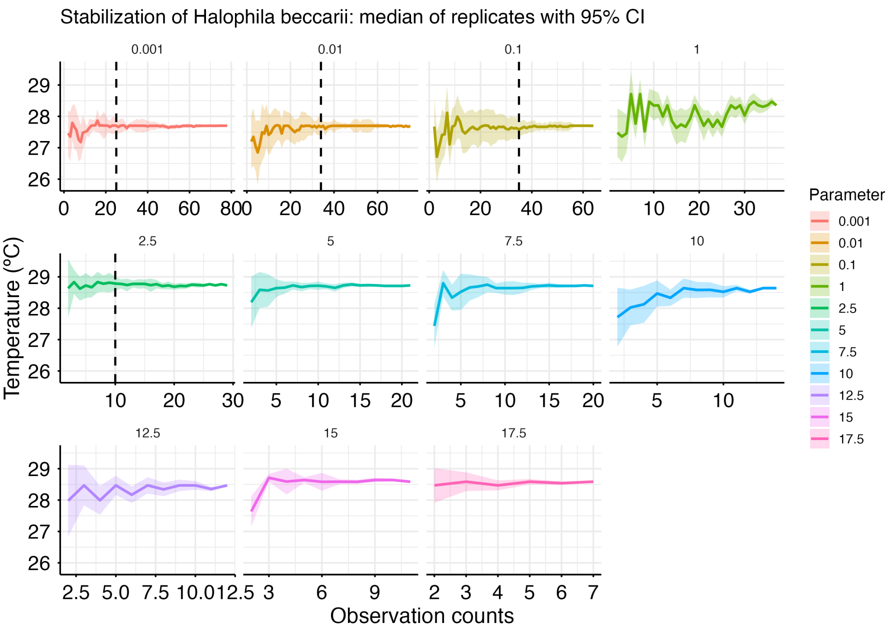
Stabilization of Halophila beccarii: median of replicates with 95% CI
0.001
0.01
0.1
1
2.5
5
7.5
10
12.5
15
17.5
Temperature (°C)
Observation counts
Parameter
0.001
0.01
0.1
1
2.5
5
7.5
10
12.5
15
17.5

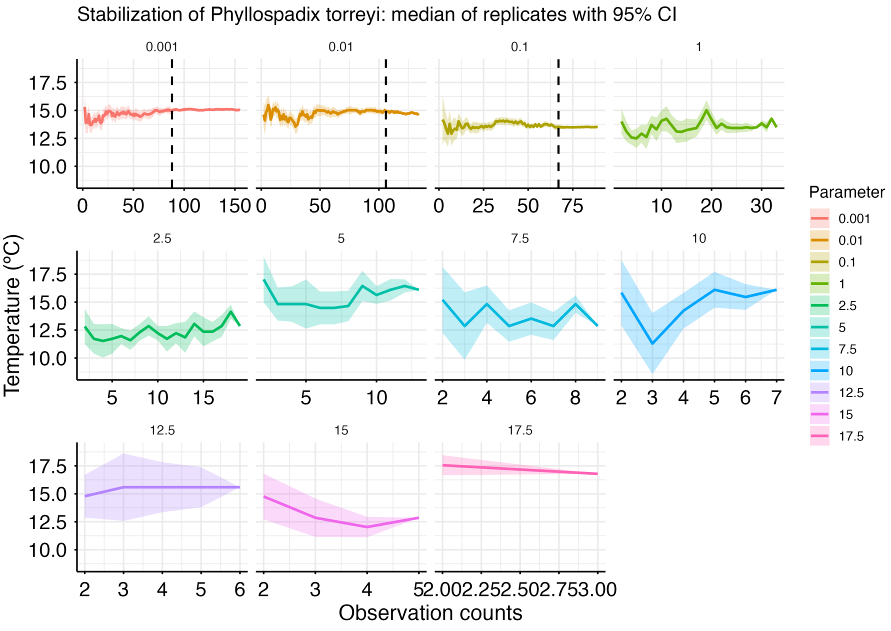
Stabilization of Phyllospadix torreyi: median of replicates with 95% CI
0.001
0.01
0.1
1
2.5
5
7.5
10
12.5
15
17.5
Temperature (°C)
Observation counts
Parameter
0.001
0.01
0.1
1
2.5
5
7.5
10
12.5
15
17.5

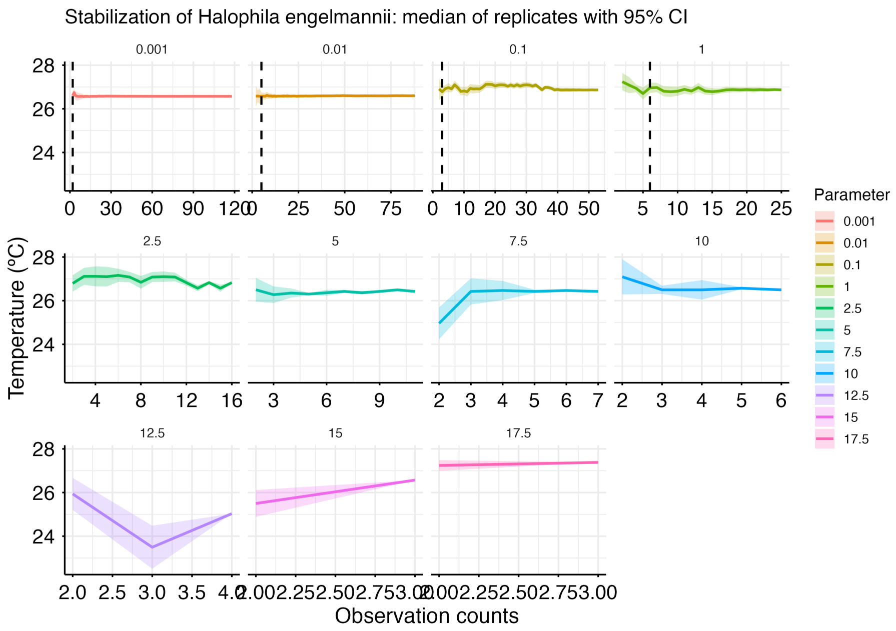
Stabilization of Halophila engelmannii: median of replicates with 95% CI
0.001
0.01
0.1
1
2.5
5
7.5
10
12.5
15
17.5
Temperature (°C)
Observation counts
Parameter
0.001
0.01
0.1
1
2.5
5
7.5
10
12.5
15
17.5

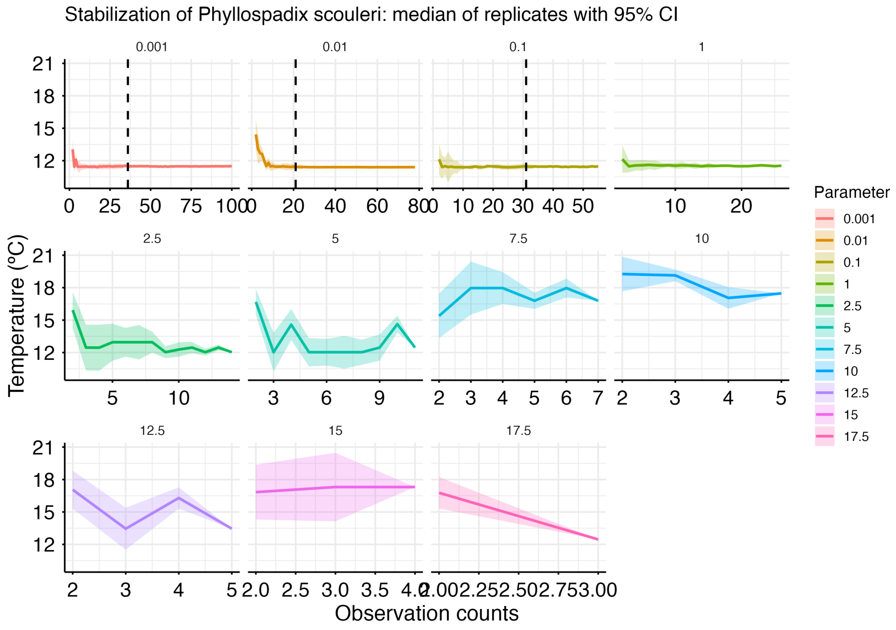
Stabilization of Phyllospadix scouleri: median of replicates with 95% CI
0.001
0.01
0.1
1
2.5
5
7.5
10
12.5
15
17.5
Temperature (°C)
Observation counts
Parameter
0.001
0.01
0.1
1
2.5
5
7.5
10
12.5
15
17.5

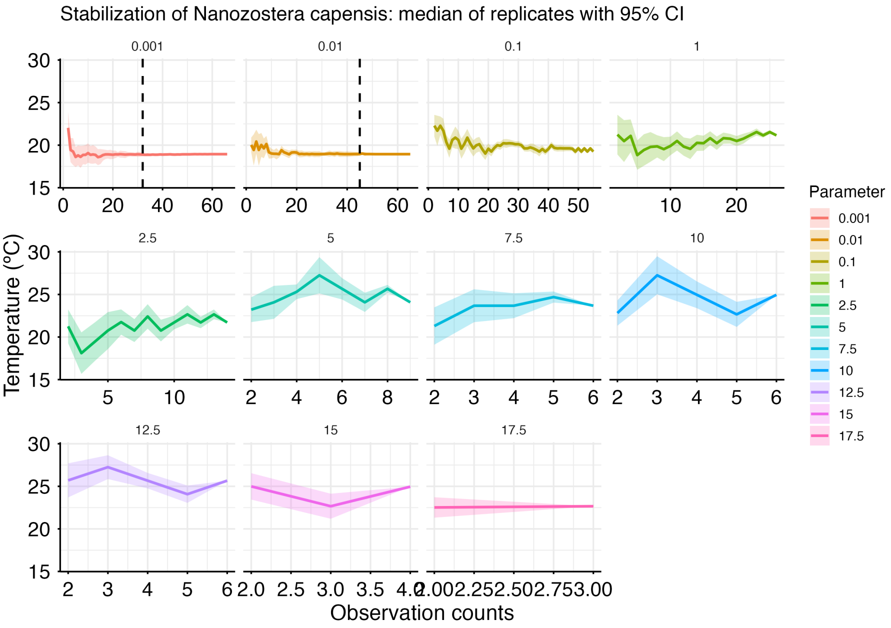
Stabilization of Nanozostera capensis: median of replicates with 95% CI
0.001
0.01
0.1
1
2.5
5
7.5
10
12.5
15
17.5
Temperature (°C)
Observation counts
Parameter

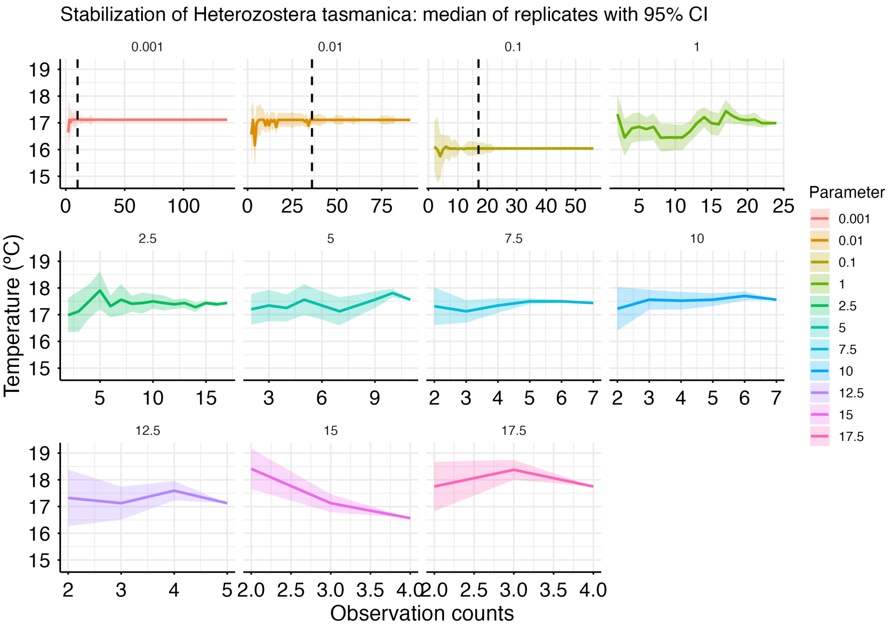
Stabilization of Heterozostera tasmanica: median of replicates with 95% CI
0.001
0.01
0.1
1
2.5
5
7.5
10
12.5
15
17.5
Temperature (°C)
Observation counts
Parameter

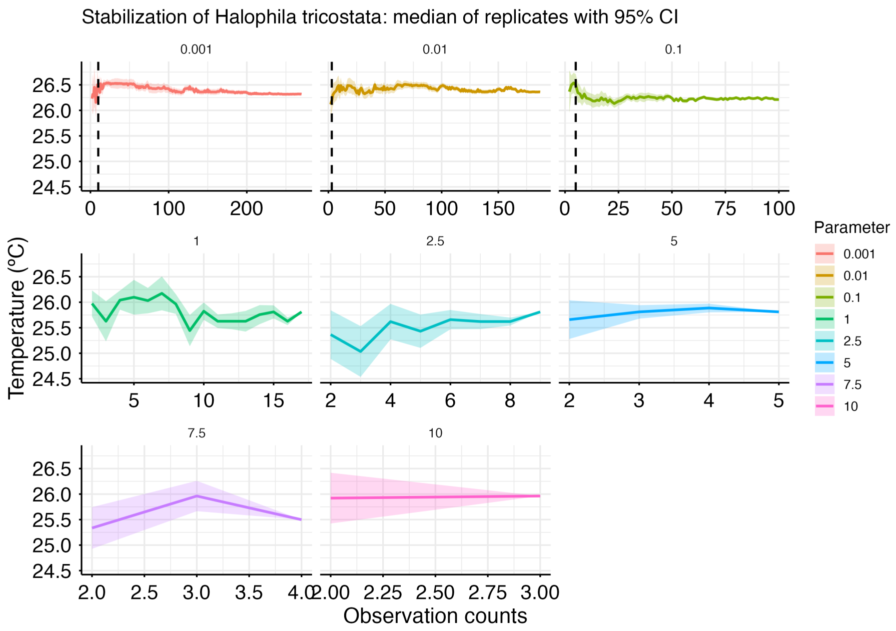
Stabilization of Halophila tricostata: median of replicates with 95% CI
0.001
0.01
0.1
1
2.5
5
7.5
10
Temperature (°C)
Observation counts
Parameter
0.001
0.01
0.1
1
2.5
5
7.5
10

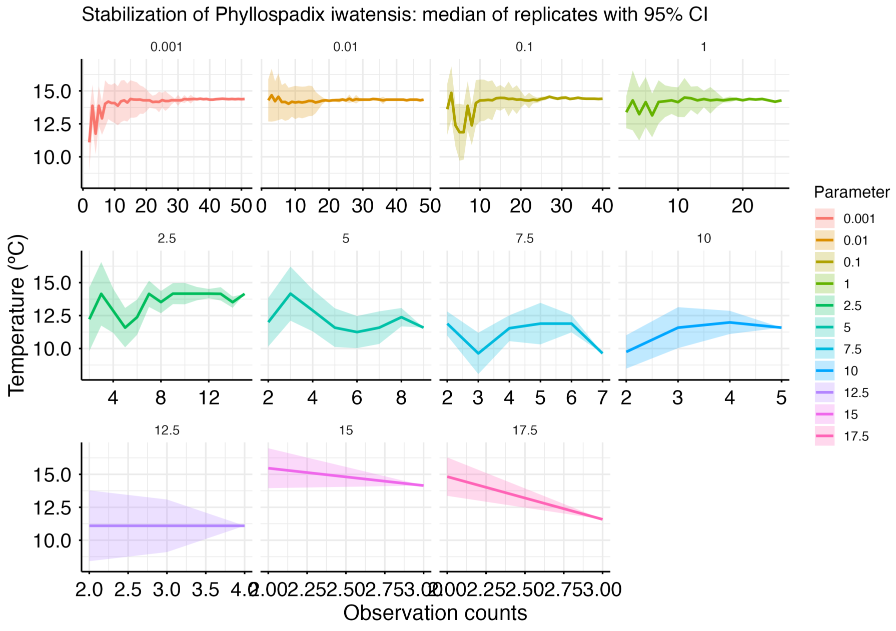
Stabilization of Phyllospadix iwatensis: median of replicates with 95% CI
0.001
0.01
0.1
1
2.5
5
7.5
10
12.5
15
17.5
Temperature (°C)
Observation counts
Parameter
0.001
0.01
0.1
1
2.5
5
7.5
10
12.5
15
17.5

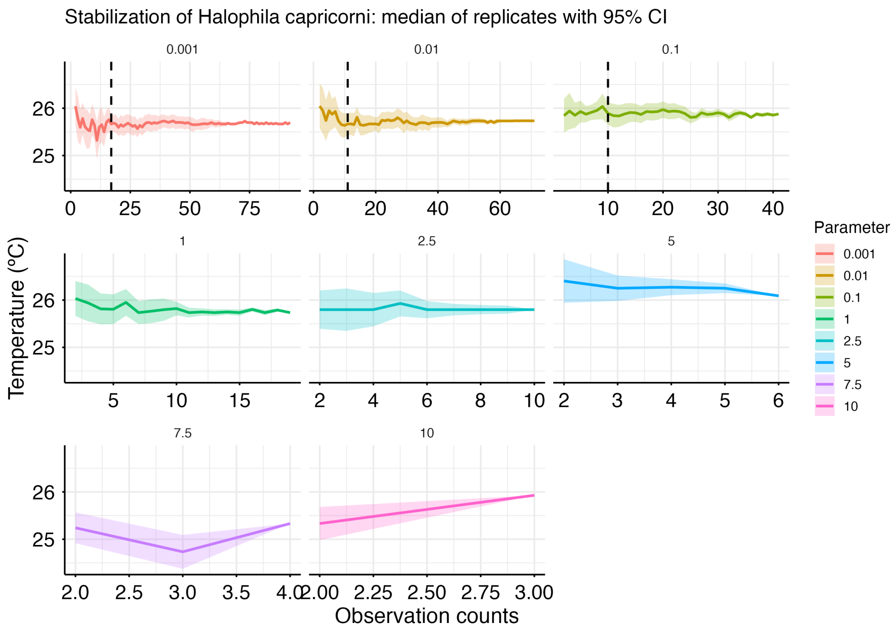
Stabilization of Halophila capricorni: median of replicates with 95% CI
0.001
0.01
0.1
1
2.5
5
7.5
10
Temperature (°C)
Observation counts
Parameter
0.001
0.01
0.1
1
2.5
5
7.5
10

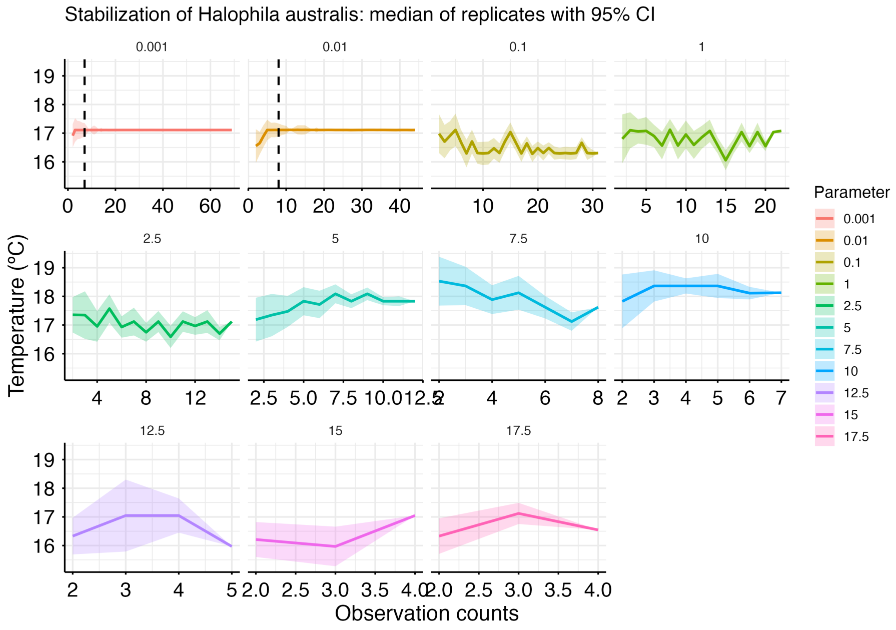
Stabilization of Halophila australis: median of replicates with 95% CI
0.001
0.01
0.1
1
2.5
5
7.5
10
12.5
15
17.5
Temperature (°C)
Observation counts
Parameter
0.001
0.01
0.1
1
2.5
5
7.5
10
12.5
15
17.5

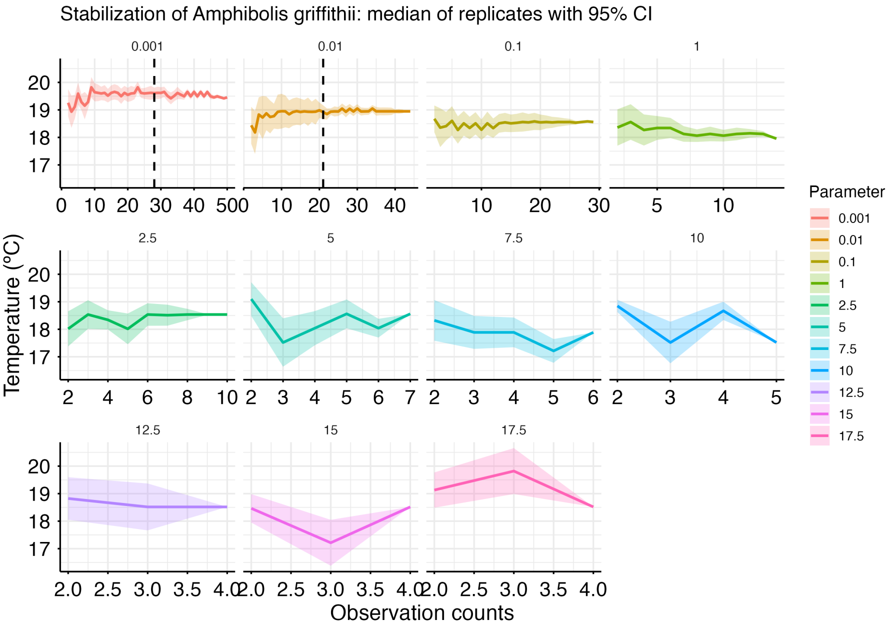
Stabilization of Amphibolis griffithii: median of replicates with 95% CI
Temperature (°C)
Observation counts
Parameter
0.001
0.01
0.1
1
2.5
5
7.5
10
12.5
15
17.5

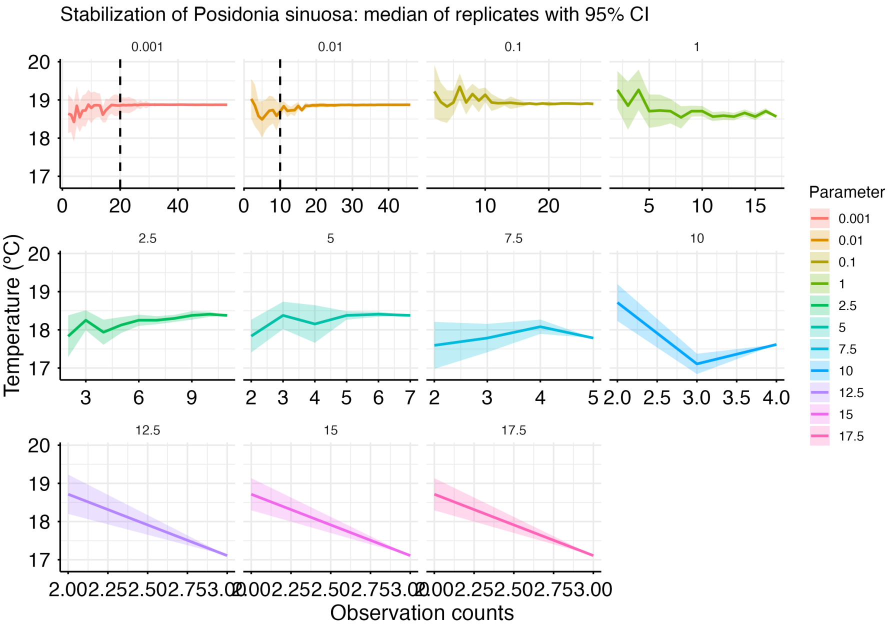
Stabilization of Posidonia sinuosa: median of replicates with 95% CI
Temperature (°C)
Observation counts
Parameter
0.001
0.01
0.1
1
2.5
5
7.5
10
12.5
15
17.5

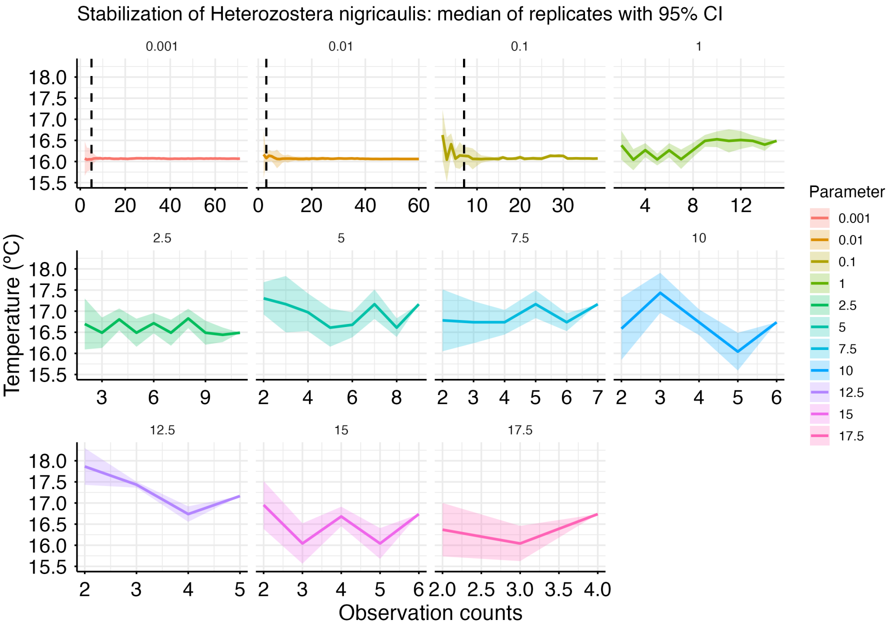
Stabilization of Heterozostera nigricaulis: median of replicates with 95% CI
0.001
0.01
0.1
1
2.5
5
7.5
10
12.5
15
17.5
Temperature (°C)
Observation counts
Parameter
0.001
0.01
0.1
1
2.5
5
7.5
10
12.5
15
17.5

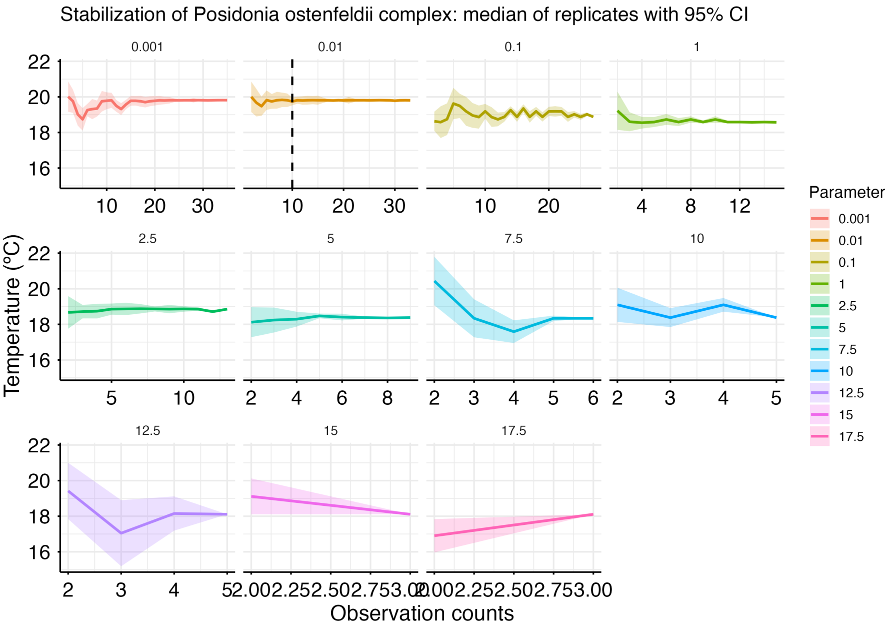
Stabilization of Posidonia ostenfeldii complex: median of replicates with 95% CI
0.001
0.01
0.1
1
2.5
5
7.5
10
12.5
15
17.5
Temperature (°C)
Observation counts
Parameter
0.001
0.01
0.1
1
2.5
5
7.5
10
12.5
15
17.5

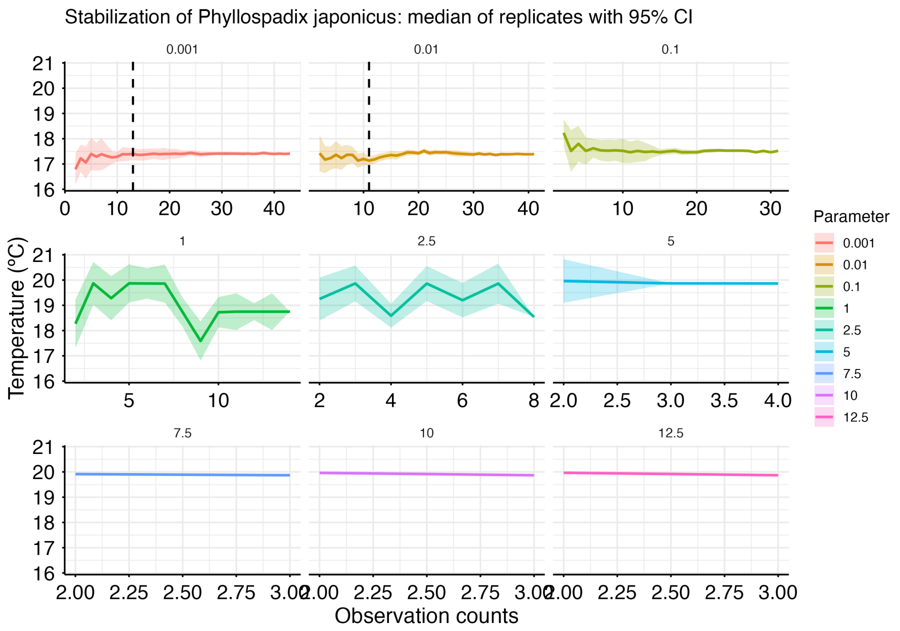
Stabilization of Phyllospadix japonicus: median of replicates with 95% CI
Temperature (ºC)
Observation counts
Parameter
0.001
0.01
0.1
1
2.5
5
7.5
10
12.5

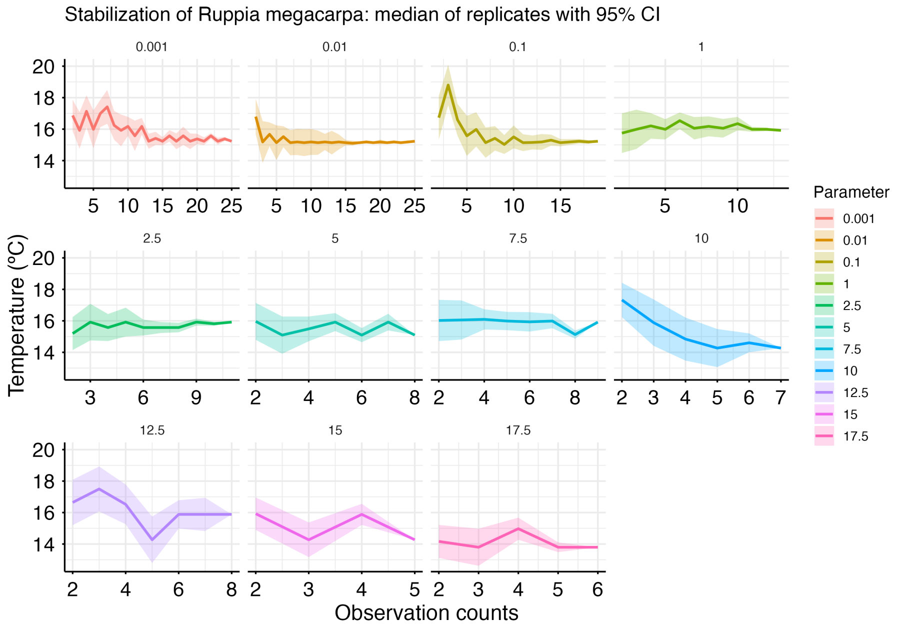
Stabilization of Ruppia megacarpa: median of replicates with 95% CI
Temperature (ºC)
Observation counts
Parameter
0.001
0.01
0.1
1
2.5
5
7.5
10
12.5
15
17.5

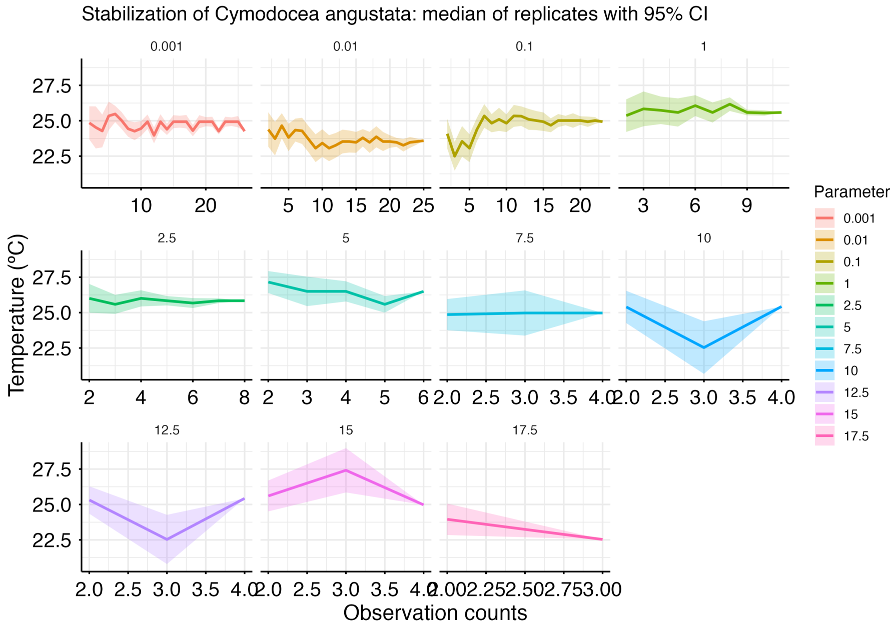
Stabilization of Cymodocea angustata: median of replicates with 95% CI
Temperature (°C)
Observation counts
Parameter
0.001
0.01
0.1
1
2.5
5
7.5
10
12.5
15
17.5

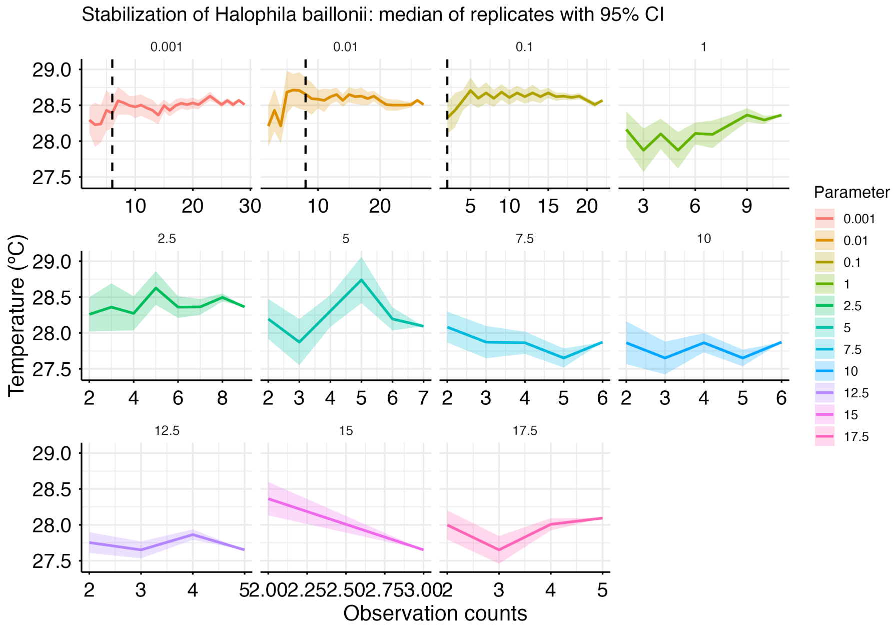
Stabilization of Halophila baillonii: median of replicates with 95% CI
Temperature (°C)
Observation counts
Parameter
0.001
0.01
0.1
1
2.5
5
7.5
10
12.5
15
17.5

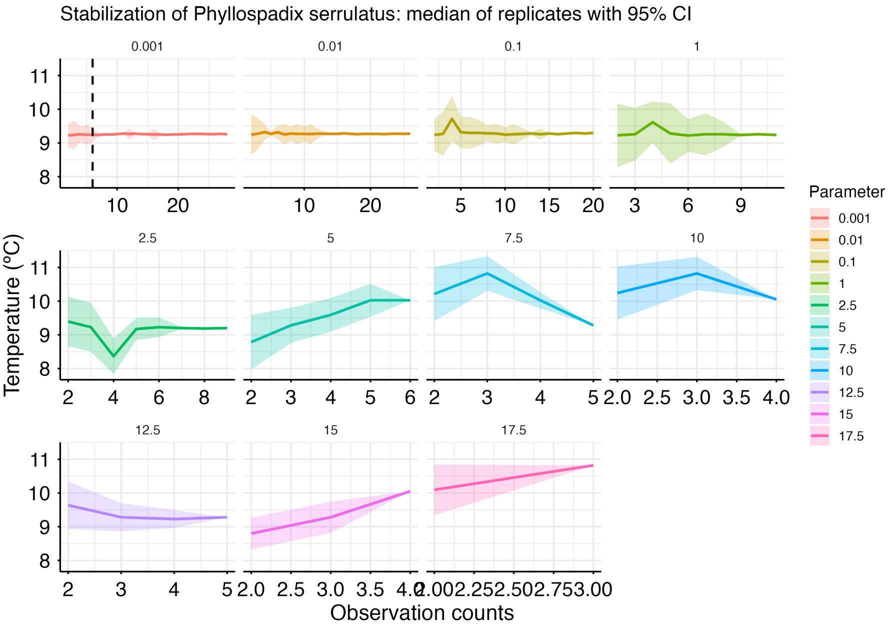
Stabilization of Phyllospadix serrulatus: median of replicates with 95% CI
0.001
0.01
0.1
1
2.5
5
7.5
10
12.5
15
17.5
Temperature (°C)
Observation counts
Parameter
0.001
0.01
0.1
1
2.5
5
7.5
10
12.5
15
17.5

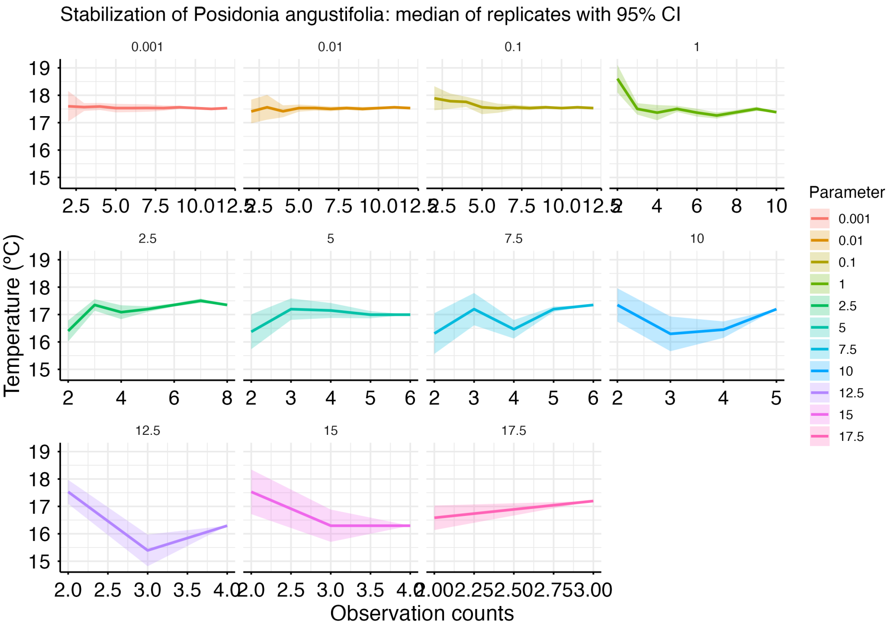
Stabilization of Posidonia angustifolia: median of replicates with 95% CI
0.001
0.01
0.1
1
2.5
5
7.5
10
12.5
15
17.5
Temperature (°C)
Observation counts
Parameter
0.001
0.01
0.1
1
2.5
5
7.5
10
12.5
15
17.5

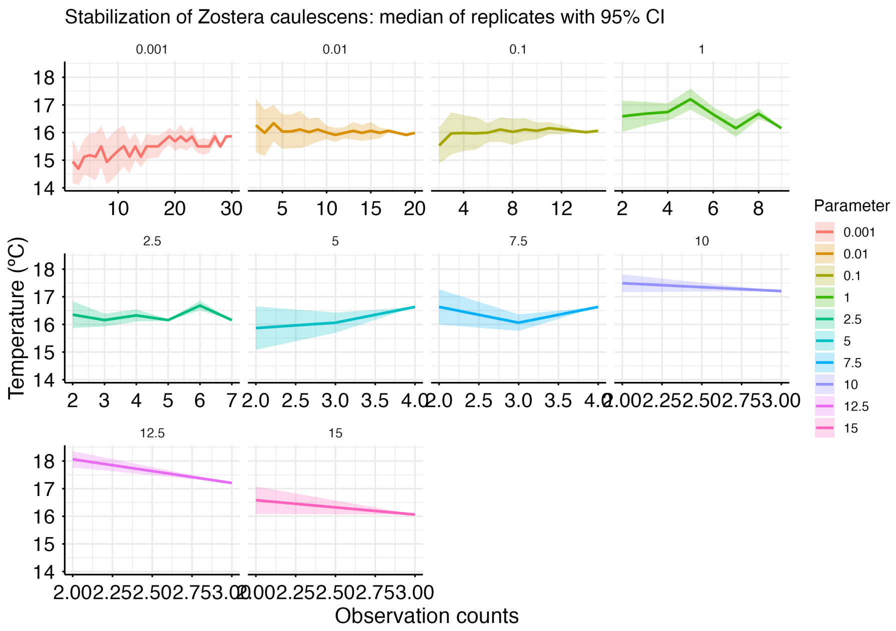
Stabilization of Zostera caulescens: median of replicates with 95% CI
0.001
0.01
0.1
1
2.5
5
7.5
10
12.5
15
Temperature (°C)
Observation counts
Parameter
0.001
0.01
0.1
1
2.5
5
7.5
10
12.5
15

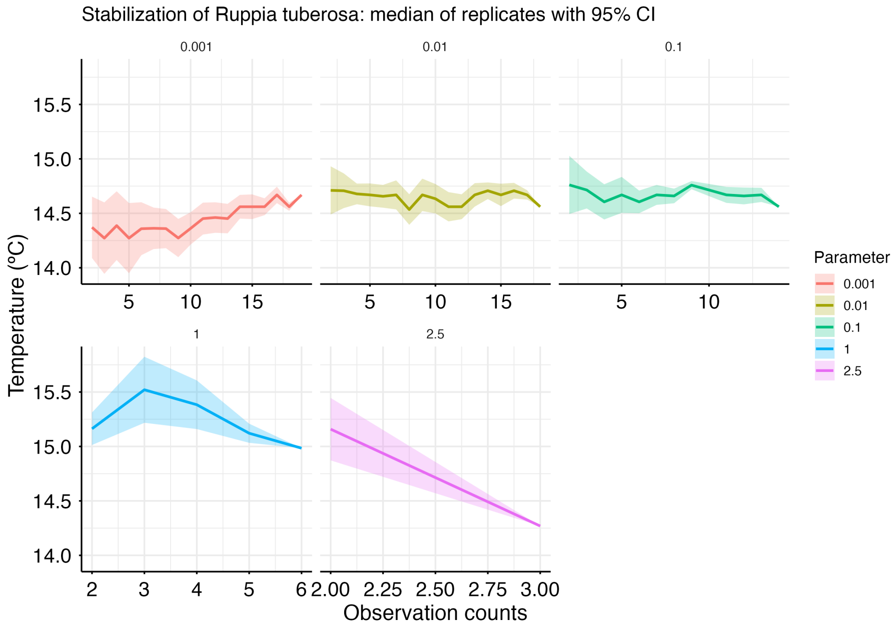
Stabilization of Ruppia tuberosa: median of replicates with 95% CI
0.001
0.01
0.1
1
2.5
Temperature (°C)
Observation counts
Parameter
0.001
0.01
0.1
1
2.5

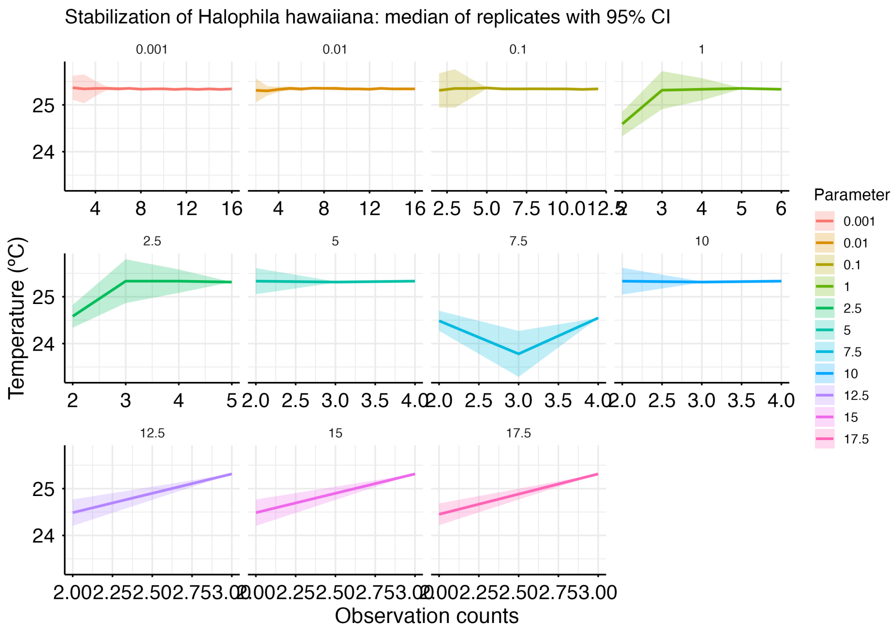
Stabilization of Halophila hawaiiana: median of replicates with 95% CI
0.001
0.01
0.1
1
2.5
5
7.5
10
12.5
15
17.5
Temperature (°C)
Observation counts
Parameter
0.001
0.01
0.1
1
2.5
5
7.5
10
12.5
15
17.5

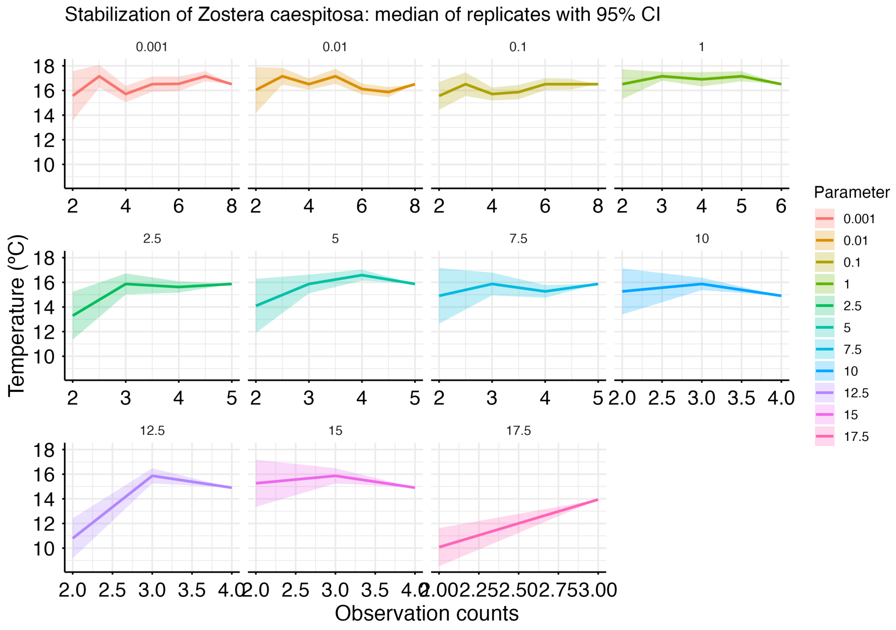
Stabilization of Zostera caespitosa: median of replicates with 95% CI
0.001
0.01
0.1
1
2.5
5
7.5
10
12.5
15
17.5
Temperature (°C)
Observation counts
Parameter
0.001
0.01
0.1
1
2.5
5
7.5
10
12.5
15
17.5

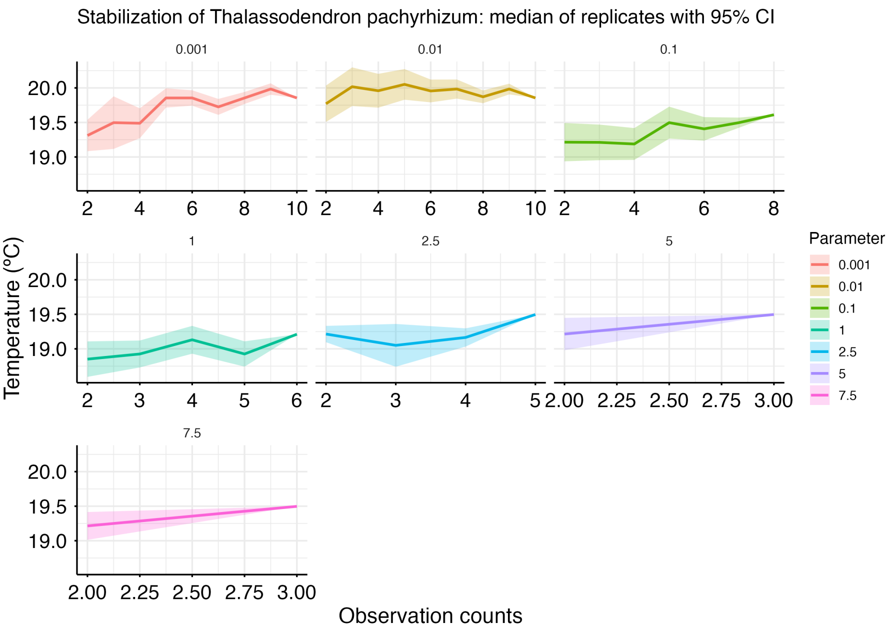
Stabilization of Thalassodendron pachyrhizum: median of replicates with 95% CI
0.001
0.01
0.1
1
2.5
5
7.5
Temperature (°C)
Observation counts
Parameter
0.001
0.01
0.1
1
2.5
5
7.5

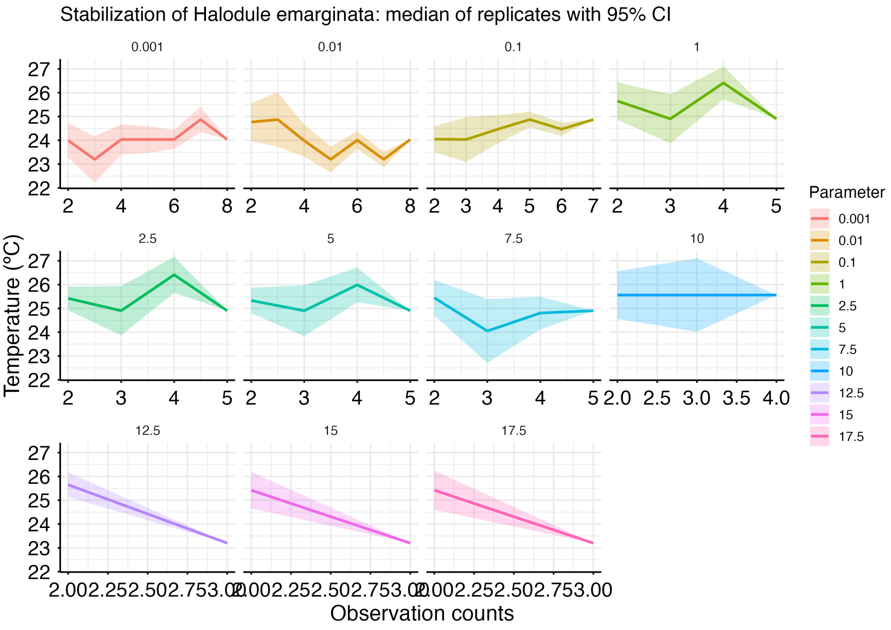
Stabilization of Halodule emarginata: median of replicates with 95% CI
0.001
0.01
0.1
1
2.5
5
7.5
10
12.5
15
17.5
Temperature (°C)
Observation counts
Parameter
0.001
0.01
0.1
1
2.5
5
7.5
10
12.5
15
17.5

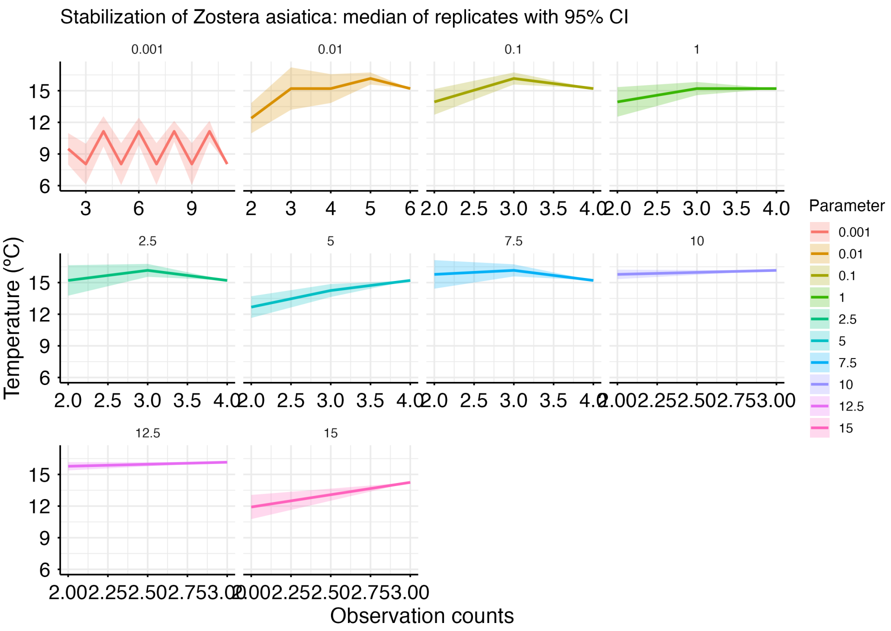
Stabilization of Zostera asiatica: median of replicates with 95% CI
0.001
0.01
0.1
1
2.5
5
7.5
10
12.5
15
Temperature (°C)
Observation counts
Parameter

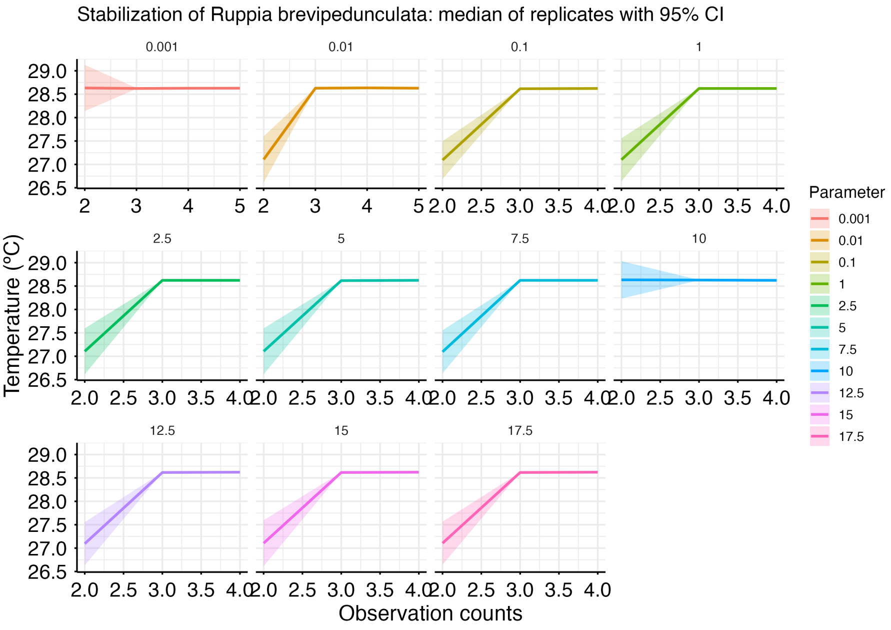
Stabilization of Ruppia brevipedunculata: median of replicates with 95% CI
0.001
0.01
0.1
1
2.5
5
7.5
10
12.5
15
17.5
Temperature (°C)
Observation counts
Parameter

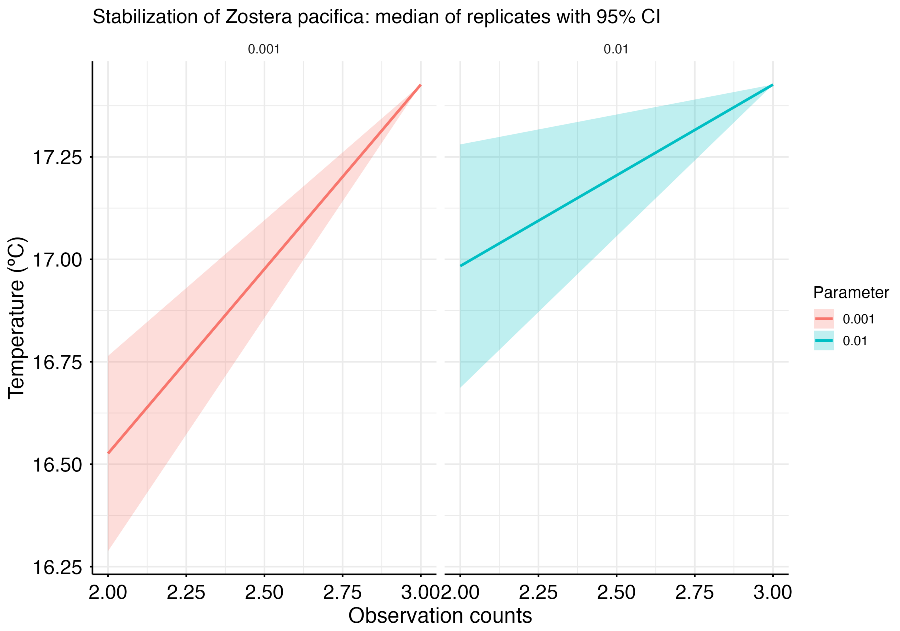
Stabilization of Zostera pacifica: median of replicates with 95% CI
0.001
0.01
17.25
17.00
16.75
16.50
16.25
Temperature (°C)
2.00
2.25
2.50
2.75
3.00
Observation counts
Parameter
0.001
0.01

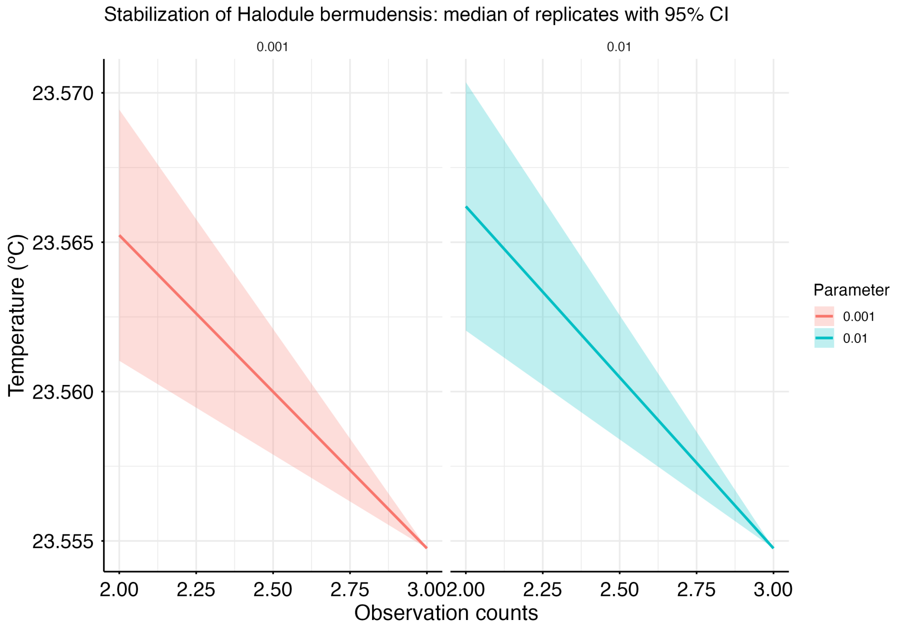
Stabilization of Halodule bermudensis: median of replicates with 95% CI
0.001
0.01
23.570
23.565
23.560
23.555
Temperature (°C)
2.00
2.25
2.50
2.75
3.00
Observation counts
Parameter
0.001
0.01

#### S7: SENSITIVITY ANALYSES

##### S7.1. Alternative range size resolutions

To evaluate the sensitivity of the framework to the spatial resolution used for calculating the range-size descriptor ($D_s$), analyses were repeated using resolutions 0.25°, 0.50°, and 0.75° (0.5º ± 50%). Fig. 4 summarises the coefficient of variation (CV) of the results in $D_s$, and LS-TP and BD-TP parameters across the three resolutions, while Fig. 5 shows the resulting SVM decision surfaces. The metric used as an input ($D_s$) varies considerably with resolution, yet the quantities of ecological interest (LS and BD parameters) remain largely unchanged.

Range size (Spearman $\rho > 0.99$), LS thinning parameter (Spearman $\rho = 1$), and BD thinning parameter estimates (Spearman $\rho > 0.98$) were highly consistent across spatial resolutions for all pairwise comparisons. As expected, $D_s$ values exhibited moderate variation between resolutions because the metric is directly derived from grid-cell size. LS parameters were identical across all resolutions (CV = 0 for all species), given its independence from $D_s$. BD-TP predicted parameters showed minimal variation for most species, with only a few outliers displaying higher sensitivity. The main driver that explains the right-skewed CV-data distribution for the BD-TP comparison is the outlier *H. tricostata*, whose $\theta_{BD}$ is capped at resolutions 0.5 and 0.75, but which is calculated as $\theta = 0.42$ for the 0.25 resolution.

DRAFT

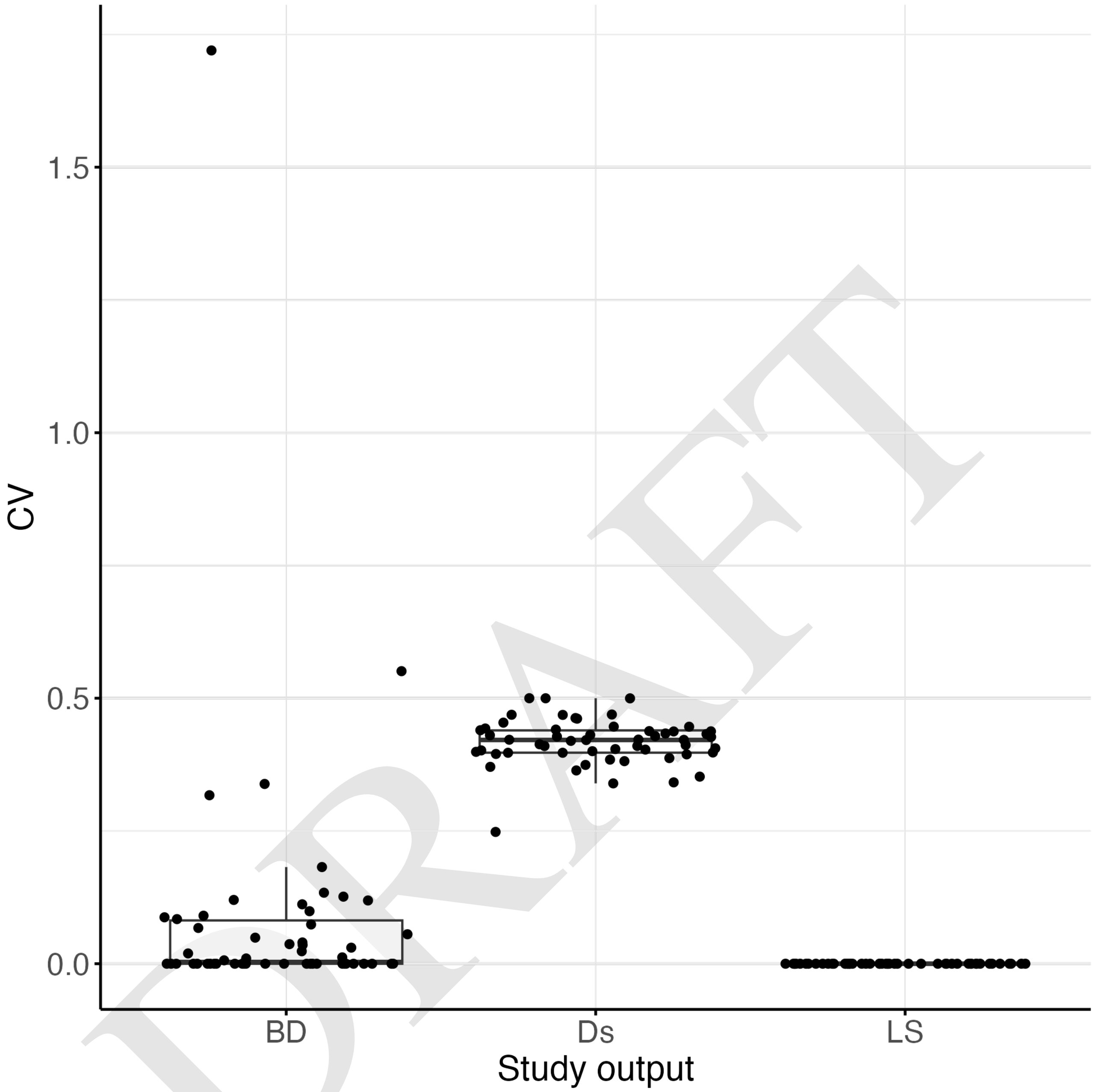

DRAFT

**Figure 4.** Distribution of the Coefficient of Variation for the study outputs (BD-TP, Ds and LS-TP) for each species (dots).

Comparison of the SVM decision hyperplanes revealed that increasing the resolution used to calculate $D_s$ primarily resulted in a vertical expansion of y-axis (i.e. larger $D_s$ values) and, thus, steeper hyperplanes, while the overall conclusion to be driven from the stability boundary (i.e. the predicted thinning parameter) remained largely unchanged. Hence, the framework is robust to the choice of spatial resolution for $D_s$ computation.

##### S7.2. Alternative tolerance settings for the stabilization

Three tolerance settings are key for the stability computations: 'tol_value', the maximum absolute difference from the final median (default = 0.3); 'tol_width', the maximum confidence interval width (default = 0.5); and 'window_run', the number of consecutive observations in which stability is kept (default = 20). A sensitivity analysis was run using deviations of ±50% from the default. The proportion of species detected as always unstable (for all parameters) was used as a comparison along the $3^3$ possible tolerance setting combinations. Results were summarised in a heatmap (Fig. 6). Only window_run affected the results

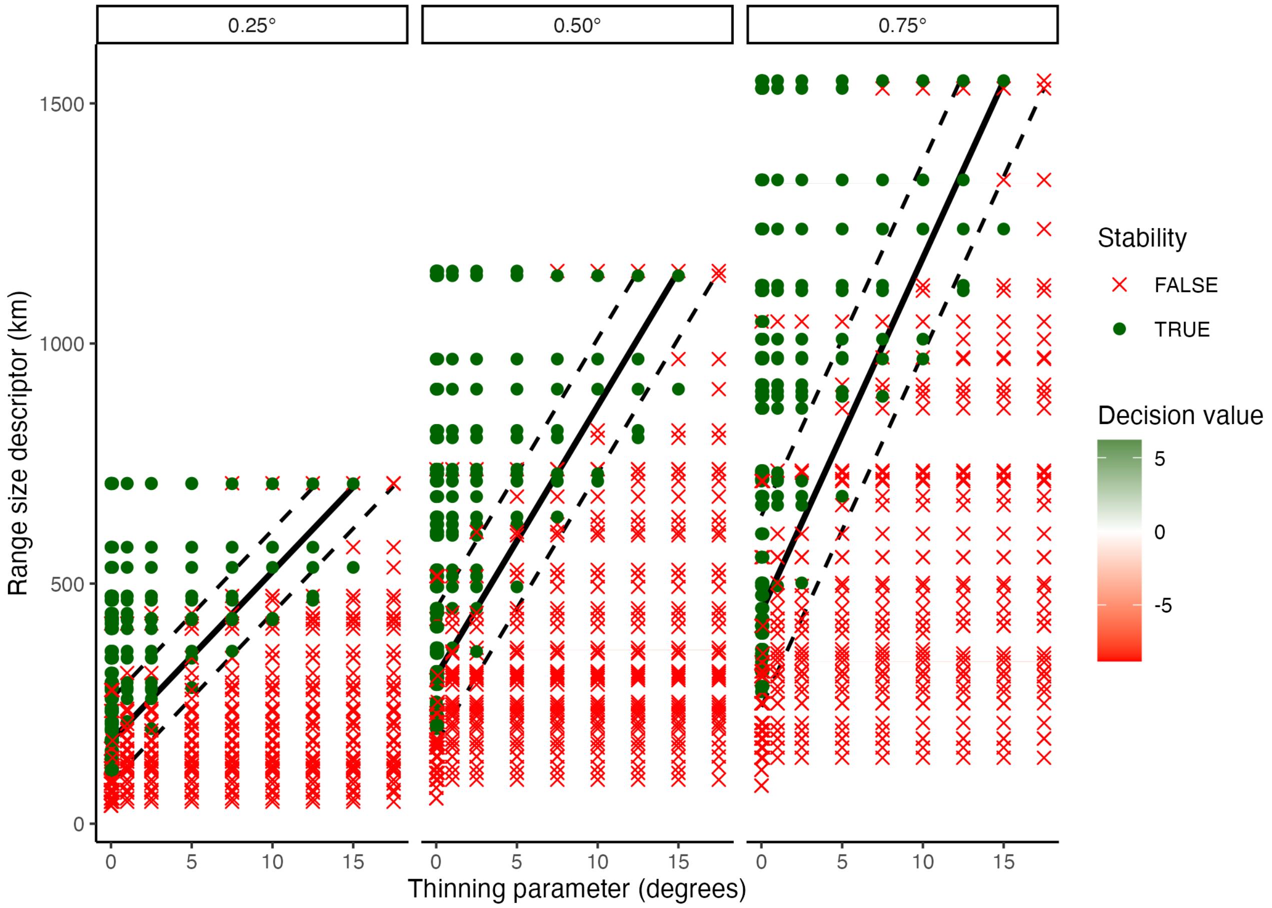


**Figure 5.** BD-TP and BD-LS results for the three tested resolutions for range size computation, rasterising original records with resolutions of 0.25, 0.5, and 0.75°.

clearly (in $\tilde{2}5\%$ from 10 to 30). Tolerance width is the second setting with the highest impact on results although the variation is less marked. Differences due to the tolerance absolute value are negligible.

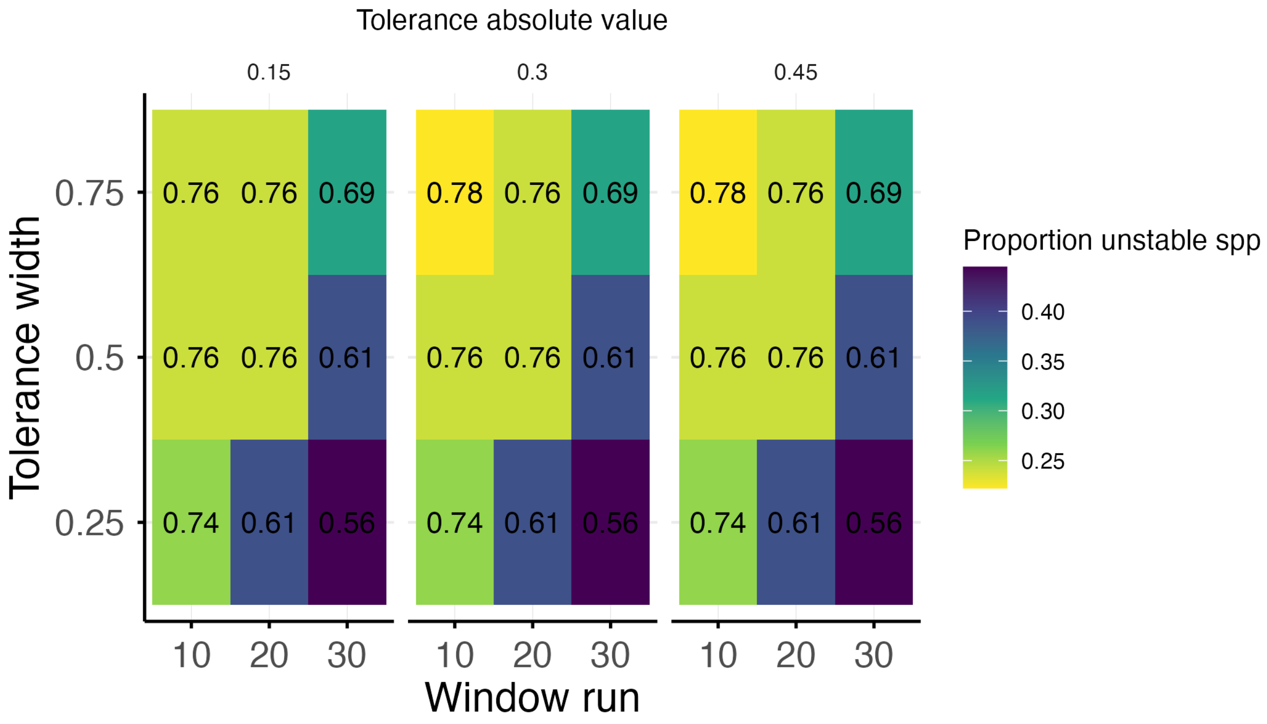


**Figure 6.** Heatmap showing the sensitivity analysis for tolerance settings. Alternative tolerance settings deviate by ±50% the default. The proportion of species that are unstable at all thinning parameters (proportion stable spp.) was used for the comparison

##### S7.3. Alternative parameter sets

In addition to the default parameter set, we tested different parameter grains, generating parameter sets with the following parameter quantities within the original range $(0, 17.5]$: 2, 4, 6, 8, 10, 12, 14, 16, 18, 20 (Table 8).

**Table 8.** The 10 parameter sets of different grains (2 to 20) used for the sensitivity analysis.

| Grain | Parameters |
|---|---|
| n_2 | [5 10] |
| n_4 | [3 6 9 12] |
| n_6 | [2.1 4.3 6.4 8.6 10.7 12.9] |
| n_8 | [1.7 3.3 5.0 6.7 8.3 10.0 11.7 13.3] |
| n_10 | [1.4 2.7 4.1 5.5 6.8 8.2 9.5 10.9 12.3 13.6] |
| n_12 | [1.2 2.3 3.5 4.6 5.8 6.9 8.1 9.2 10.4 11.5 12.7 13.8] |
| n_14 | [1 2 3 4 5 6 7 8 9 10 11 12 13 14] |
| n_16 | [0.9 1.8 2.6 3.5 4.4 5.3 6.2 7.1 7.9 8.8 9.7 10.6 11.5 12.4 13.2 14.1] |
| n_18 | [0.8 1.6 2.4 3.2 3.9 4.7 5.5 6.3 7.1 7.9 8.7 9.5 10.3 11.1 11.8 12.6 13.4 14.2] |
| n_20 | [0.7 1.4 2.1 2.9 3.6 4.3 5.0 5.7 6.4 7.1 7.9 8.6 9.3 10.0 10.7 11.4 12.1 12.9 13.6 14.3] |

Recalculating the main plot that summarises both BD-TP and LS-TP showed a similar pattern to the one predicted by the default settings, confirming our selection of 10 parameters appropriate as a default (Fig. 7).

DRAFT

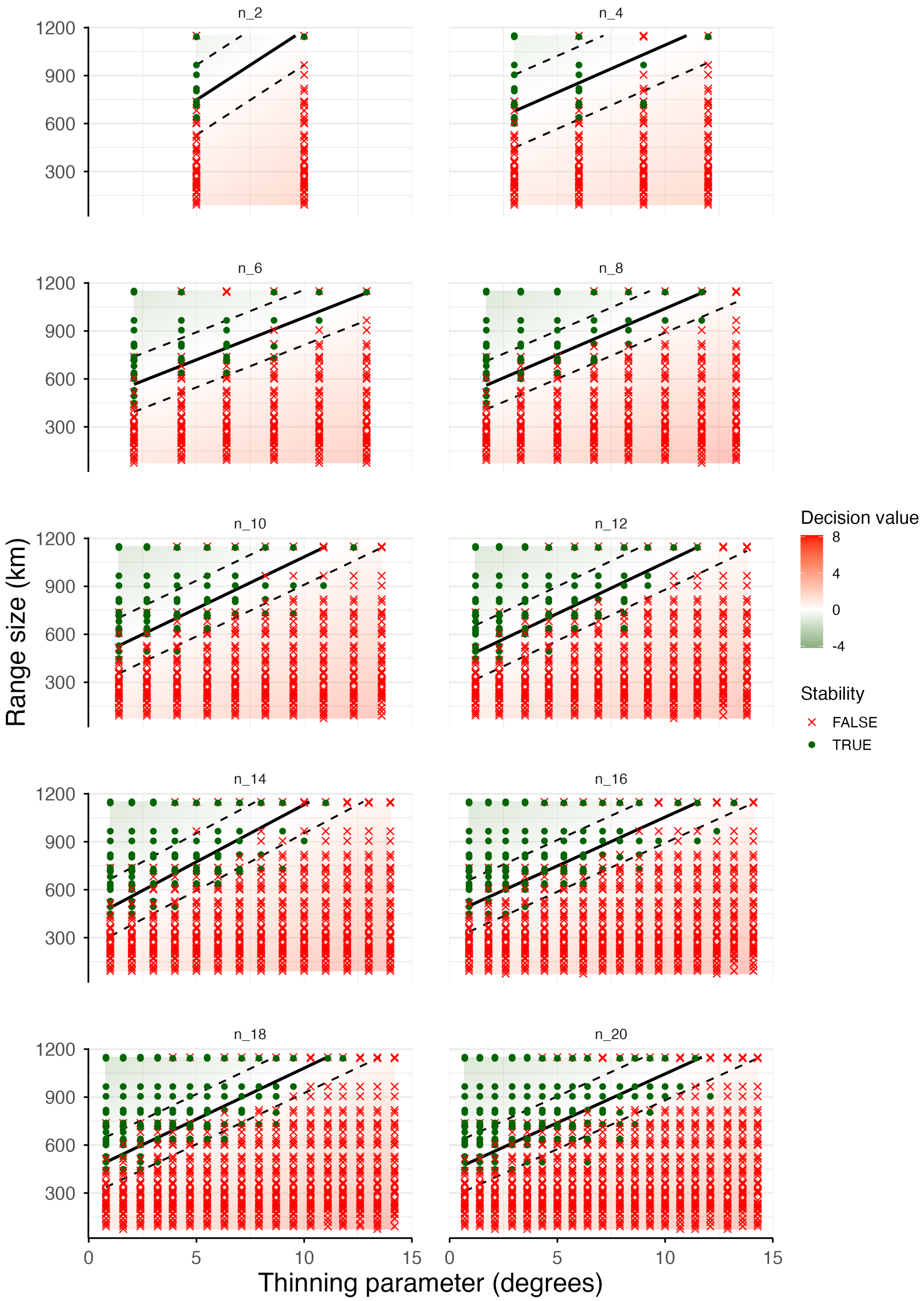


**Figure 7.** SVM plots showing the BD-TP results for different parameters grains (10 parameter sets). Thermally stable thinning parameter and species combinations are represented with green dots, whereas thermally unstable combinations are shown in red crosses. The black solid line represents the hyperplane predicted by BD-TP. Dashed lines indicate margin boundaries. The LS-TP output can be interpreted as the last stable symbol per species.

DRAFT

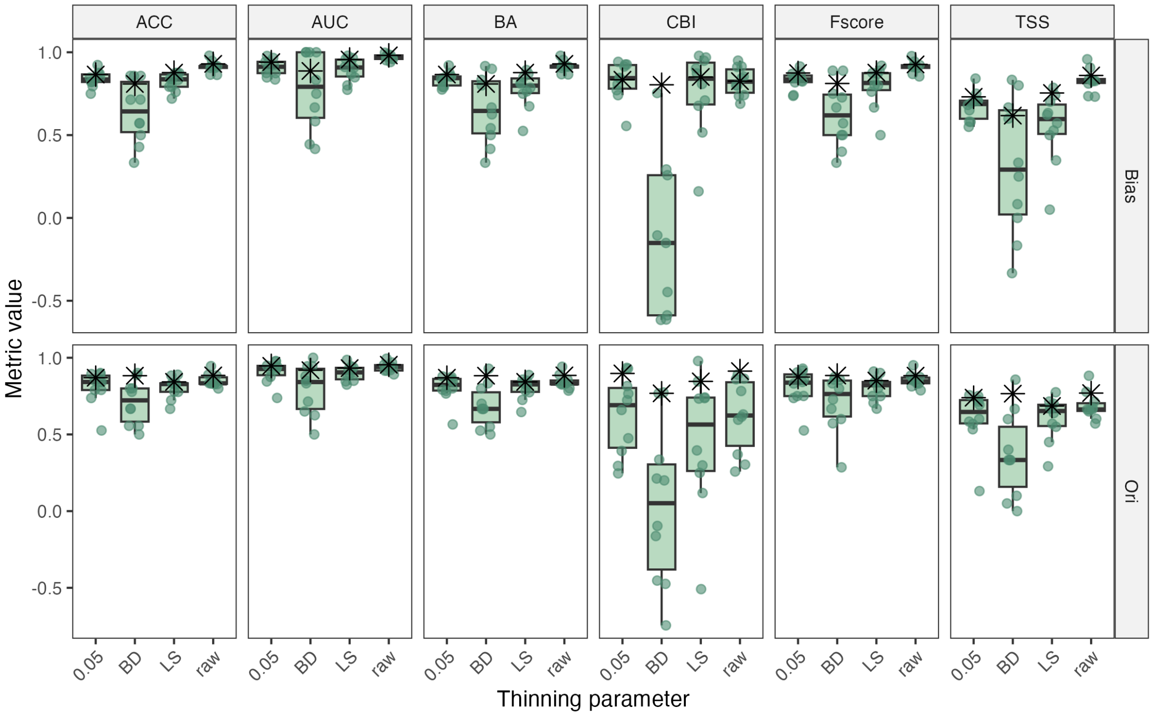


**Figure 8.** K-fold cross-validated (boxplots) and total value for evaluation metrics. Cross-validation was performed as a subset of the thinned dataset.

The standard deviation of the predicted values across cross-validation folds of the HSM provided a spatial overview of model uncertainty. HSI values were low for all cases.

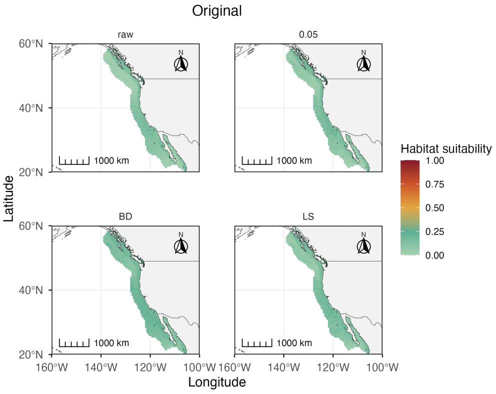


**(a)** Original occurrence dataset.

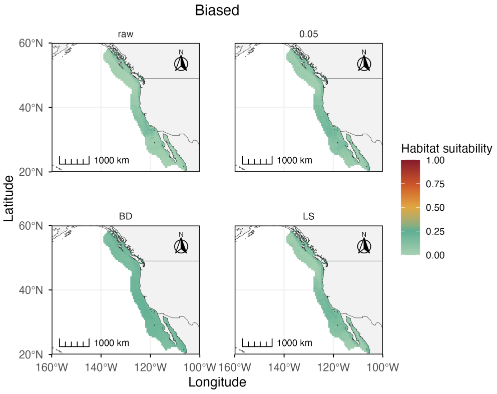


**(b)** Artificially biased occurrence dataset.

**Figure 9.** Maps showing the standard deviation of habitat suitability maps for *Phyllospadix torreyi* based on (a) the original and (b) artificially biased occurrence datasets. Habitat suitability indexes were calculated with BART and SD values are shown with a gradient fill. Both in (a) and (b), the raw data (raw) and the data thinned with 0.05 and the predicted parameters by BD and LS were inputted into the model. The predictive distribution is shown as the median.
